# The human brain mechanisms of afterimages: From networks to cortical layers

**DOI:** 10.1101/2025.08.30.673266

**Authors:** Sharif I. Kronemer, Burak Akin, Micah Holness, A. Tyler Morgan, Laurentius Huber, Paul A. Taylor, Javier Gonzalez-Castillo, Daniel A. Handwerker, Peter A. Bandettini

## Abstract

Afterimages are common visual illusions that have long attracted popular and scientific interest, yet their neural mechanisms remain little understood. We used high-spatial-resolution fMRI to investigate human whole brain and cortical layer activity in primary visual cortex (V1) linked with afterimages and perceptually similar animated images – stimuli designed to imitate the appearance of afterimages. Both afterimages and perceptually similar images engaged overlapping, widespread brain activity, particularly in visual sensory regions that follow the contralateral circuitry of the primary visual pathway. However, afterimages elicited weaker fMRI signals across the brain compared to images, except in salience network regions, where activity was enhanced for afterimages. Salience network activity could reflect conflict or perceptual reality monitoring mechanisms that detect the illusory nature of afterimages. Cortical layer-specific analyses in V1 revealed afterimages selectively engaged middle and deep cortical layers, while perceptually similar images activated middle and superficial layers. In agreement, layer-specific decoding of afterimage and mock afterimage fMRI signals was present only in deep layers. One interpretation of these V1 cortical layer findings is that afterimage perception involves feedback or top-down neural input akin to self-generated or endogenous conscious perception (e.g., visual imagery). In addition, baseline eye behavior (pupil size and blinking) and fMRI signals in arousal, salience, and visual networks differed depending on whether afterimages were perceived or not perceived. These results challenge the prevailing framing on the neurophysiological origins of afterimages as arising from either retinal *or* central neural processes. As with image perception, typical afterimages emerge by the interaction between retinal *and* central neural mechanisms. These findings on the neural processing underlying afterimages may be broadly informative of the neural mechanisms of visual consciousness.

## Introduction

Afterimages – historically known as “visual persistence”, “aftereffects”, or “ocular spectra” – are common visual illusions. Typically, afterimages are induced by a light source that is no longer physically present. For instance, when headlights from a car disappear from view at night or when looking away from an image on a computer screen, an illusory conscious vision may appear with similar visual features (e.g., shape and visual field location) as the original light source that can last for many seconds, even in total darkness or when the eyes are closed. Afterimages are perceived as either the same color or shade (positive afterimages) or a complementary color or shade (negative afterimages) as the inducing image. While afterimages are specific to vision, there are analogous forms of perseverating sensory perception in audition and somatosensation [1, 2].

Afterimages have been studied for millennia. Nevertheless, the precise neural mechanisms of afterimages remain little understood. The role of retinal signaling following photoreceptor adaptation or bleaching (e.g., after viewing a bright light) is commonly referenced [3]. Likewise, Hermann von Helmholtz characterized afterimages as “a photograph on the retina” [4]. A retinal-based mechanism is supported by the observation that afterimage position in visual space updates with gaze position [4–6]. Also supporting a retinal contribution are afterimage-linked signals or “after-responses” that are found in retinal afferents and relay cells of the cat and monkey lateral geniculate nucleus (LGN) [7, 8]. In addition, even without photoreceptor bleaching, retinal ganglion cells in monkeys have been shown to evoke a post-stimulation rebound, potentially the sourcing signal of afterimages, although this response persists beyond the typical duration of afterimages [9].

However, retinal-only explanations of afterimages are incomplete. The relative contribution of retinal versus central neural mechanisms in afterimage perception has long been debated and remains unresolved [3]. Even cerebral generation of afterimages *without* retinal input has also been considered [10]. Involvement of central neural processes in afterimage perception is supported by behavioral research highlighting that afterimages can be induced by illusory images or when afterimages appear where there was no previous light (e.g., filling-in illusions) [11, 12]. There are even reports of afterimages induced by hallucination and visual imagery [13–17]. Interocular grouping (i.e., the perceptual integration of monocular afterimages) [18], afterimage rivalry [19], and conditioned afterimages (e.g., afterimages induced by conditioned tones) are also suggestive of central neural mechanisms [18, 20, 21]. Further, the modulation of afterimage perception by eye movements is evidence for post-retinal modulation of afterimages [22]. Also highlighting the potential involvement of cortical processes in modifying the neural signals underlying afterimages are examples where afterimage perception is altered by visual illusions (e.g., illusory depth and surface slant) [23, 24].

Direct support for the role of cortex in afterimage perception include findings in non-human animals of persistent visual cortical neural activity after the offset of visual stimuli that induce afterimages [25, 26]. Also, human cortical evoked potentials have been recorded with electroencephalography during the perception of afterimages, including a link between afterimages and depressed alpha frequency band activity [27–30]. Furthermore, disrupting human occipital cortical electrophysiology with transcranial magnetic stimulation altered afterimages [31].

Localization of the brain regions associated with afterimages is suggested by the few human fMRI experiments that found afterimage-linked blood-oxygen-level-dependent (BOLD) signal in human primary visual cortex (V1), fusiform gyrus (FG), and area V8 [32–35]. Likewise, a monkey fMRI study found afterimage-linked brain responses throughout visual cortex, including V2, V3, and V4 [36]. Although, another experiment found that lesioning V4 and MT did not eliminate the perception of afterimages [37]. These findings support the involvement of the visual cortex in afterimages, indicating that visual sensory brain regions may contribute in modifying retinal inputs that drive central neural activity observed with afterimages or even generating the afterimage signal itself.

Further interpretation is limited because none of the current fMRI studies report findings outside of visual cortex and all were implemented with spatial resolution that do not allow for interpreting meso-scale dynamics, including cortical layer specific activity that can infer the direction of information processing [38]. Our goal was to advance beyond prior neuroimaging studies on afterimages to further elucidate the brain mechanisms of afterimages. To achieve this, we conducted the first human whole brain and the first human cortical layer resolution fMRI study of afterimages.

In a previous behavioral experiment [39], we gathered perceptual details (sharpness, contrast, and duration) on afterimages from healthy, adult participants. We used those reports in the current investigation to design animated, on-screen images that were perceptually similar with each participant’s afterimage experience. These tailored images we refer to as “mock afterimages” to emphasize their perceptual similarity to “real” afterimages, despite mock afterimages being physically present stimuli, while nothing was physically shown when the afterimages were perceived. In the current experiment, afterimages and mock afterimages were administered in a perception task while recording pupillometry, eye tracking, and whole brain fMRI. A subset of participants also completed an additional study session during which the perception task was administered while recording pupillometry, eye tracking, and cortical-layer resolution V1 fMRI.

We hypothesized that afterimages and perceptually similar images would evoke similar activity in visual cortex and across subcortical and association cortices. However, we postulated some differences in the afterimage and image whole brain fMRI signals, including salience network regions that may distinguish illusory afterimages from real images. Furthermore, based on previous behavioral findings linking afterimages to sensory-independent perception (e.g., hallucination and imagery) and their sensitivity to top-down effects (e.g., priming), we hypothesized that afterimages would involve V1 cortical layer activity suggestive of feedback neural signaling, whereas images would exhibit a laminar profile characteristic of feedforward signaling.

Leveraging the results from this study to elucidate the brain mechanisms of afterimages is valuable for explaining centuries of behavioral research on afterimage perception and can be broadly informative on the neural mechanisms underlying visual consciousness [40].

## Methods

*Participants.* Healthy, adult participants were recruited from the local Bethesda, Maryland, United States of America (USA) community (N = 35; males = 15; mean age = 28.03 years [standard deviation (SD) = 8.73]; mean education = 16.49 years [SD = 1.42]). All participants completed a behavioral and whole brain fMRI study session (see *Whole brain fMRI sequence* section). In addition, a subset of the participants (N = 13; males = 4; mean age = 31.54 years [SD = 10.41]; mean education = 17.69 years [SD = 2.32]) completed a layer-resolution V1 fMRI study session (see *V1 fMRI sequence* section). Recruitment was completed in accordance with the National Institute of Mental Health Institutional Review Board, and all participants gave informed consent prior to participation.

*Inclusion criteria* were: (1) between 18 and 65 years old, (2) completion of a healthy physical examination by a nurse practitioner within a year of the study sessions, (3) able to provide informed consent. *Exclusion criteria* were: (1) current or previous histories of neurologic or psychiatric disorder, (2) low vision (i.e., unable to read small text on a computer screen) that could not be corrected, (3) head injuries (e.g., loss of consciousness for > 30 minutes and three or more concussive injuries). In addition, 3 participants were excluded from analyses of the whole brain fMRI dataset due to behavioral performance (incorrect reporting during the behavioral tasks; 2 participants) or reporting that they did not see afterimages following the inducer stimulus (1 participant; see *Afterimage induction* section). In the V1 fMRI study, 1 participant was excluded from analyses because they did not complete the study session (no fMRI data was collected). Of the remaining participants, 2 participants were excluded from analyses because they did not show a V1 BOLD or VASO response for task events.

In total, *32 participants* were included in the whole brain fMRI data analysis and *10 participants* were included in the V1 fMRI data analysis. While the number of participants in the V1 fMRI dataset is typical for human cortical layer fMRI studies, the small sample size should be considered in the full context of the findings. Additional participant exclusions were made for the supplementary analyses on the eye tracking and pupillometry data and whole brain and V1 fMRI data for specific event conditions based on data quality and number of data epochs (see *Analysis Methods* section).

### Behavior

*Afterimage induction.* Afterimages were induced using a black silhouette image of a human face in frontal view (presentation duration = 4 seconds; visual angle = 4.60 x 8.47 degrees; maximum luminance = ∼4 cd/m^2^; https://creazilla.com/nodes/2524-face-silhouette; Figure 1A; Supplementary Movie 1). See [39] for full details on the inducer stimulus. All but one participant (excluded from analyses) indicated that the inducer resulted in negative afterimages that appeared as a white or light gray, blurry version of the face image and was positioned in the same on-screen location as the inducer when maintaining central fixation.

**Figure 1.**
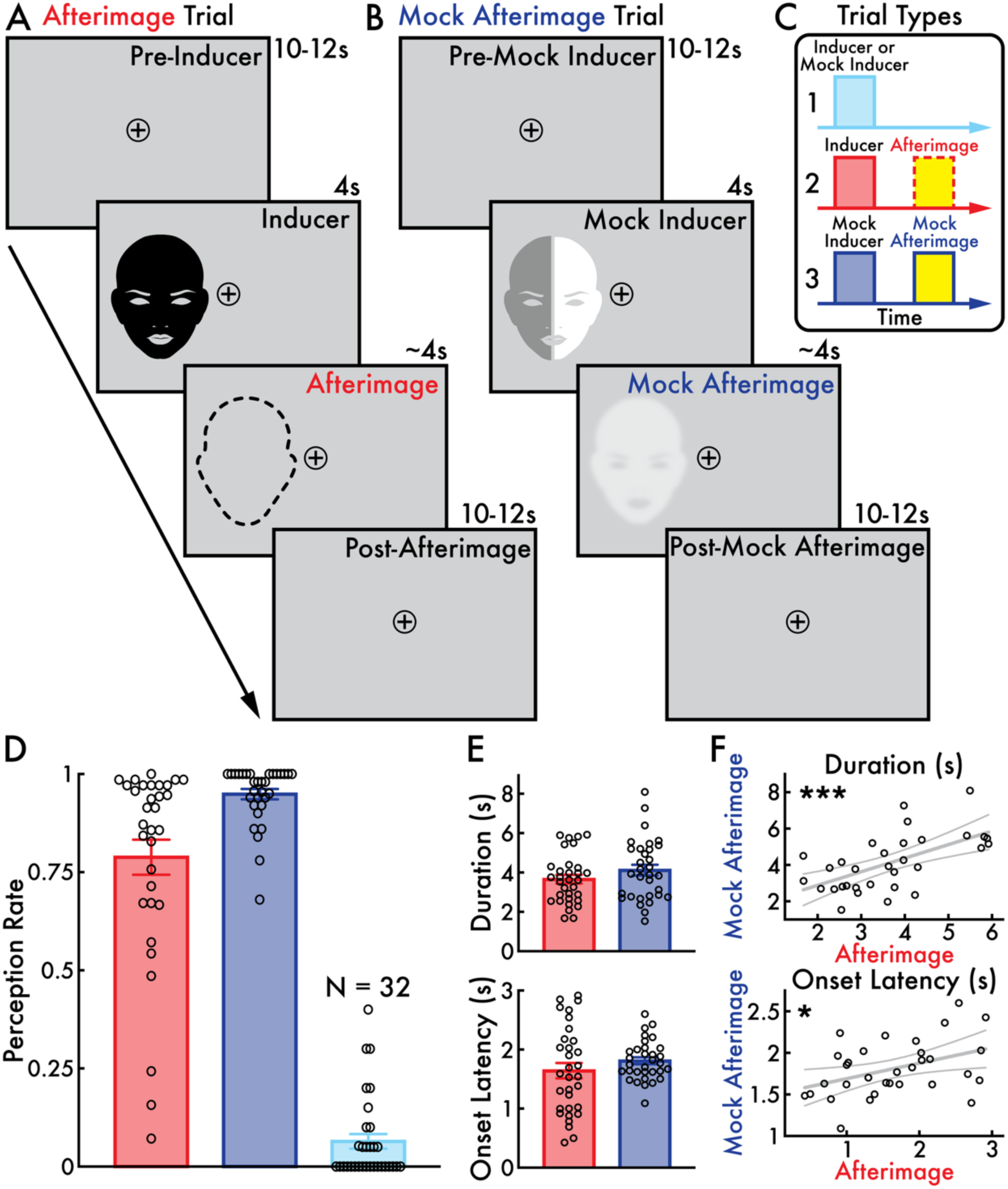
Afterimage and mock afterimage perception task and behavioral results. The afterimage and mock afterimage perception task involved three trial types **(C)**: (1) inducers *without* afterimages or mock inducers *without* mock afterimages (*inducers + mock inducers only trial*; not depicted), (2) inducers *with* afterimages (*afterimage trial*), and (3) mock inducers *with* mock afterimages (*mock afterimage trial*). The mock afterimage is an animated, on-screen image that was designed to be perceptually similar with each participant’s afterimage perception. The perceptual similarity between afterimages and mock afterimages is represented by the yellow highlight in C. Most participants believed that the mock afterimage was illusory (i.e., not a physical stimulus) because the mock afterimage resembled an afterimage (Table 1). **(A)** In afterimage trials (Supplementary Movie 1), the inducer stimulus appeared (4 seconds [s]) followed by a blank screen (jittered 10-12 s). Participants regularly reported perceiving an afterimage following the inducer, represented by a dotted outline to indicate that while the afterimage is perceived nothing physically appears on screen. **(B)** In mock afterimage trials, the mock inducer appeared (4 s; shown as a half gray and white image to represent that the mock inducer flickered between these two shades; 1 Hz). Following the disappearance of the mock inducer (post-mock inducer interval jittered 10-12 s), either a mock afterimage appeared (70% of trials; Supplementary Movie 2) or nothing appeared (blank screen; 30% of trials; Supplementary Movie 3). Participants were instructed to report the onset and offset of all afterimages and mock afterimages they perceived. **(D)** The mean perception rate with standard error of the mean (SEM) for afterimages following the inducer (red) and mock afterimages following the mock inducer when the mock afterimage was present (dark blue) and absent (light blue). Participants regularly perceived afterimages (> 0.75) and mock afterimages (> 0.90), but not when the mock inducer was presented alone (i.e., without a mock afterimage; < 0.10). **(E)** The mean afterimage and mock afterimage duration and onset latency with SEM. The afterimage and mock afterimage durations and onset latencies were not statistically different. **(F)** The correlation between the afterimage and mock afterimage durations and onset latencies with a linear regression line and 95% confidence intervals. Statistically significant positive correlations were found for both duration and onset latency (* *p* < 0.05; *** *p* < 0.001). The open circles in subplots D, E, and F represent individual participants (N = 32). Behavioral results are evaluated from the whole brain fMRI session. See the *Behavior* Analysis Methods section for full details on the mock afterimage, perception task, and behavioral analyses. The post-afterimage and mock afterimage period was inclusive of the afterimage and mock afterimage phase.

**Table 1.** Representative participant reports on perceived similarities and differences between afterimages and their perceptually similar animated images – what are referred to as “mock afterimages”. Each report (x) reflects an individual participant’s response to the post-task questionnaire (see *Post-task questionnaire* Methods section).

| *Similar* | *Different* |
| --- | --- |
| × Afterimages and mock afterimages had similar brightness, crispness, and started at the same time.<br>× Afterimages and mock afterimages looked similar, including brightness and onset time.<br>× Afterimages and mock afterimages had the same brightness and duration.<br>× Afterimages and mock afterimages clarity, onset time, and duration were similar.<br>× Afterimages and mock afterimages shape, size, position, and brightness were the same.<br>× Afterimages and mock afterimages had the same general shape and brightness.<br>× Afterimages and mock afterimages started and ended around the same time.<br>× Afterimages and mock afterimages started around the same time and had similar brightness | × Afterimages appeared more like a blob but still face-like; mock afterimages were sharper and longer lasting.<br>× Afterimages were blurrier, lasted longer, and brighter; mock afterimages had a delayed onset.<br>× Mock afterimages looked artificial; were more detailed; afterimages were blurry.<br>× Afterimages started slightly earlier and was briefer than the mock afterimages.<br>× Mock afterimages appeared for longer.<br>× Afterimages had an earlier onset; mock afterimage was brighter and more vivid.<br>× Afterimages were brighter.<br>× Mock afterimages were clearer, slower to appear, and longer lasting.<br>× Mock afterimages were sharper, clearer, and longer lasting. |

*Afterimage and image perceptual reporting task.* To access perceptual details on the perceived visual features of afterimages, previous studies have asked participants to report on the subjective appearance of their afterimages [41, 42]. Similarly, in a previous study, we developed a perceptual reporting method that asked participants to indicate three perceptual characteristics: (1) sharpness, (2) contrast, and (3) duration (Supplementary Figure 1A) [39]. The perceptual reporting task involved participants using a “controllable image” – a white version of the inducer stimulus – that they manually adjusted in real time with key presses to indicate either the sharpness or contrast and duration (i.e., contrast and duration were simultaneously reported) of their afterimages following the presentation of the inducer (see *Afterimage induction* section). The controllable image was selected to approximately match the white or light gray appearance of the afterimages.

Serving to both train participants and validate the afterimage perceptual reports, participants first matched the controllable image to on-screen images. These images were modified versions of the controllable image, which were systematically varied in sharpness, contrast, and duration to approximate the perceptual range of afterimages along these dimensions, based on pilot testing (see [39] for full details). When comparing the subjective reports with the image features (e.g., the reported sharpness of the image versus the true image sharpness), it was found that participants were accurate indicating the sharpness, contrast, and duration of the images [39]. This result validates the perceptual reporting method and supports the reliability of the afterimage reports.

*Mock afterimage.* The participants in the current experiment (see *Participants* section) are a subset of the participants reported in our previous study [39]. For each participant, we used their reports from [39] on afterimage sharpness, contrast, and duration to create a customized “mock afterimage” – a unique mock afterimage for each participant. The mock afterimage is an *on-screen*, *animated image* designed to resemble an illusory afterimage (Supplementary Movie 2). In short, the mock afterimage is a *perceptually similar* image to each participant’s afterimage. Individualized mock afterimages were created to capture the variability in afterimage experiences across participants (e.g., some participants described bright, vivid, and prolonged afterimages, while others reported dim, blurry, and brief afterimages [39]; Supplementary Figure 1C and D). Importantly, participants were not initially informed that the mock afterimage was an on-screen image. Instead, they were led to believe that these perceptions were also illusory afterimages. A post-task questionnaire was administered to assess whether the afterimage and mock afterimage appeared similar or different, and to evaluate whether they believed the mock afterimage was real or illusory (see *Post-task questionnaire* section).

The mock afterimage was a manipulated version of the controllable image, thereby matching the shape and shade of the afterimage following the inducer (see *Afterimage and image perceptual reporting task* section). Specifically, the sharpness of each mock afterimage was set to the participant’s mean reported afterimage sharpness (Supplementary Figure 1C). The sharpness of the mock afterimage was constant throughout its entire presentation. Since afterimages are often perceived with dynamic brightness or vividness, to match the change in afterimage contrast over time, the mock afterimage was animated according to how participants reported their afterimage contrast changed during its perception. The contrast over time and duration of the mock afterimage was calculated by finding the mean duration of the afterimage (see [39] for details), scaling each reported afterimage by this duration, then resampling the contrast values at 100-millisecond increments, and, finally, calculating the mean opacity at each time point (Supplementary Figure 1D). To eliminate rapid, unnatural jumps in contrast, the mock afterimage contrast-by-time array was smoothed (Savitzky-Golay filter: window size = 11 seconds; polynomial order = 1).

The mock afterimage served two main functions. First, the mock afterimage helped validate the accuracy of the afterimage perceptual reports (see *Afterimage and mock afterimage perceptual matching validation* section). Specifically, if participants were unable to distinguish between their afterimages and mock afterimages or did not realize that the mock afterimage was a physical, on-screen stimulus (i.e., believed it to be illusory), this would support that the perceptual reports used to construct the mock afterimage accurately reflected the afterimage perception.

Second, the mock afterimage served to compare physiological signals associated with perceiving physical images versus illusory afterimages. As a perceptually similar counterpart to the afterimage, the mock afterimage enabled this contrast by controlling for low-level visual feature differences (e.g., a long-lasting, bright real image versus a short, dull afterimage). By leveraging the mock afterimage, the current study examined images and afterimages that elicited similar conscious experiences yet distinguished by their origins: image perception associated with ongoing visual input and afterimage perception emerging without simultaneous visual stimulation, although perseverating retinal activity may still be present.

A key difference between the afterimage and mock afterimage is that the afterimage is proceeded by an inducer stimulus. Meanwhile, the mock afterimage can be perceived by its presentation alone (i.e., without a preceding visual input). The inducer potentially biases results because it will introduce physiological changes that are systematically present for the afterimage but absent in the mock afterimage condition. This potential confound is especially challenging in fMRI due to hemodynamic delays that result in the mixing of signals associated with closely timed events. To address this challenge, we designed a “mock inducer” stimulus that was paired with the mock afterimage to match the inducer-afterimage pair (Figure 1B; Supplementary Movie 2). The mock inducer was identical in size, shape, and presentation duration as the inducer (see *Afterimage induction* section). However, instead of a static image, the mock inducer flickered at 1 Hz between a light and dark gray shade. These visual characteristics of the mock inducer were tailored to not evoke afterimages, confirmed by pilot testing (data not shown) and the current behavioral results (see *Afterimage and mock afterimage perception task* Results section; Figure 1D; Supplementary Figure 2A).

By presenting the mock inducer immediately before the mock afterimage, we approximated the physiological signal interactions between the inducer and afterimage by mimicking this pairing with the mock inducer and mock afterimage. In addition, the mock inducer helped to conceal that the mock afterimage was an on-screen image (i.e., if the mock afterimage appeared alone, it would become apparent that it was a physical stimulus; see *Afterimage and mock afterimage perceptual matching validation* section). Whole brain and V1 analyses compared the fMRI responses associated with inducers and mock inducers to verify that they were not systematically biasing findings attributed to afterimages and mock afterimages (see *Whole brain BOLD signals* Results section; Supplementary Figures 8 and 17). These analyses emphasize that the afterimage and mock afterimage fMRI responses cannot be fully explained by differences between the inducer and mock inducer.

*Afterimage and mock afterimage perceptual matching validation.* To validate that the afterimage and mock afterimage were perceptually similar, participants completed a *perceptual matching validation task* (Supplementary Figure 1B). In this task, participants were introduced to the mock inducer (see *Mock afterimage* section). However, participants were misleadingly instructed that afterimages could appear following the mock inducer. Instead, unbeknownst to the participants, when the mock inducer disappeared, each participant’s unique mock afterimage was presented in a subset of trials. Therefore, it could appear as if the mock inducer was followed by an afterimage.

The perceptual matching validation task involved 48 trials. In 16 trials, neither the inducer nor mock inducer appeared (blank trials). For the remaining 32 trials, both the inducer and mock inducer appeared on opposites sides of the screen, while participants centrally fixated. Each trial consisted of four main task phases (Supplementary Figure 1B): (1) pre-inducer and mock inducer (6-8 seconds; not shown in Supplementary Figure 1B), (2) inducer and mock inducer (4 seconds), (3) afterimage and mock afterimage (10-12 seconds), and (4) question phases (self-paced).

During the *pre-inducer and mock inducer phase*, only a central fixation image (a plus sign inside an open circle) appeared and participants were instructed to maintain their gaze on this point. During the *inducer and mock inducer phase* both the inducer and mock inducer appeared on opposite sides of the screen, the inducer and mock inducer location counterbalanced across trials. During the *afterimage and mock afterimage phase*, the screen was blank (a solid gray background) on the side of the screen where the inducer appeared. On the side of the screen where the mock inducer appeared, the mock afterimage was presented in 17 out of 32 trials; in the remaining trials, this side of the screen was blank.

Finally, during the *question phase*, participants were asked “Where did you see an afterimage?”. Participants indicated with a keypress whether they perceived an afterimage on (1) the left or (2) right side of the screen, (3) both sides of the screen, or (4) they did not perceive any afterimages on either side of the screen. Notably, at this stage of the experiment, participants were unaware that the mock afterimage was an on-screen stimulus (see *Mock afterimage* Results section). Thereby, the instruction to report on the location of afterimages was inclusive of both afterimages and mock afterimages, as the participants believed that the mock afterimages were also illusory afterimages at this stage of the study.

*Afterimage and mock afterimage perception task.* The afterimage and mock afterimage perception task was administered with simultaneous fMRI, eye tracking, and pupillometry (see *Whole brain fMRI sequence*, *V1 fMRI sequence*, and *Eye tracking and pupillometry* sections). This task comprised of ∼10-minute blocks consisting of 28 trials each. Participants completed a total of 5 blocks per fMRI study session except for three participants in the whole brain fMRI study session: 1 participant completed only 4 blocks due to reporting discomfort and 2 participants who completed only 121 and 128 trials of the total 140 trials across 5 blocks due to the task presentation software crashing during the experiment.

Each task trial consisted of 4 phases (Figure 1A and B): (1) *pre-inducer and mock inducer* (10-12 seconds), (2) *inducer and mock inducer* (4 seconds), (3) *afterimage and mock afterimage* (variable duration of ∼4 seconds), and (4) *post-afterimage and mock afterimage phases* (10-12 seconds; the post-afterimage and mock afterimage period was inclusive of the afterimage and mock afterimage phase; i.e., the mock afterimage presentation duration was subtracted from the post-mock afterimage duration, so the cumulative duration of the mock afterimage and post-mock afterimage phases was less than or equal to 12 seconds).

The *pre-inducer and mock inducer phases* were identical and consisted of a blank gray screen with a central fixation image (a plus sign inside an open circle) that remained visible throughout all trial phases. During the *inducer and mock inducer phases*, either the inducer or mock inducer image appeared on either the right or left side of the fixation point. The inducer and mock inducer location was randomly selected on a trial-by-trial basis, while there was an equal number of presentations between the left and right sides of the screen. In the *afterimage phase*, nothing appeared (a blank screen) following the offset of the inducer (*afterimage trial*; Figure 1A). In the *mock afterimage phase*, in ∼70 percent of trials (10 out of 14 mock inducer trials per block) a mock afterimage appeared in the same on-screen location that the mock inducer was presented, while for ∼30 percent of trials (4 out of 14 mock inducer trials per block) nothing appeared (*mock afterimage trial*; Figure 1B). Finally, the *post-afterimage and mock afterimage phases* were identical and consisted of a blank gray screen with a central fixation image.

Whenever participants perceived an afterimage or mock afterimage, they were instructed to immediately indicate its onset and offset using button presses. If nothing appeared following the inducer and mock inducer, participants were instructed to withhold a response. The buttons coding for perceived onset and offset were counterbalanced across participants (condition A: button 1 = onset, button 2 = offset; condition B: button 2 = onset, button 1 = offset).

At this stage of the experiment, most participants were unaware that the mock afterimage was an on-screen stimulus. For naïve participants, they were instructed to report the onset and offset of afterimages, which was inclusive of mock afterimages because these participants believed they were also illusory afterimages. For the participants who became aware that the mock afterimage was an on-screen image either during the perceptual matching validation task (see *Afterimage and mock afterimage perceptual matching validation* section) or the perception task, they were instructed to report on the onset and offset of both afterimages and mock afterimages.

*Post-task questionnaire.* After completing the perceptual matching validation and the afterimage and mock afterimage perception tasks (i.e., after exiting the MRI scanner for the whole brain and V1 fMRI study sessions), participants were administered a post-task questionnaire. The questionnaire inquired on whether participants perceived afterimages, whether the afterimages and mock afterimages appeared similar or different, and if they noticed that the mock afterimage was an on-screen image. Also, participants were encouraged to share any notable observations regarding their afterimage and mock afterimage experiences.

### Eye tracking and pupillometry

*Eye tracking and pupillometry acquisition.* Eye behaviors are known to reflect widespread brain activity and are linked to conscious perception [43, 44]. To complement and help interpret the fMRI findings, we acquired and analyzed pupil size, blink, and microsaccade dynamics. Head-fixed, eye tracking and pupillometry were gathered during the whole brain and V1 fMRI study sessions with the EyeLink 1000 Plus system (recorded eye: right eye, except for 5 participants during the whole brain fMRI session where the left eye offered more stable eye tracking; sampling rate = 1000 Hz; MRI compatible, long-range camera; SR Research, Inc.). The tracker camera was mounted inside the MRI bore, positioned behind the participant’s head. The tracker camera was oriented to capture the participant’s eye via a mirror affixed to the MRI head coil (see *Whole brain fMRI sequence*, *V1 fMRI sequence*, and *Equipment, software, and facility* sections) – the same mirror participants used to view the projector screen displaying the task. The eye tracker camera was positioned approximately 64 cm from the participant’s eye. The head coil and foam padding used to stabilize head position also helped ensure reliable eye tracking throughout the study. Although, for a minority of participants, eye tracking and pupillometry quality was poor due to the intermittent loss of eye position.

### fMRI

*Whole brain fMRI sequence.* Whole brain fMRI was acquired on a MAGNETOM 7T Plus MRI (Siemens, Inc., Healthineers, Erlangen, Germany) with a 32-channel head coil (Nova Medical, Inc., Wilmington MA, USA). BOLD imaging was recorded with a 2D-multi-band sequence (voxel size = 1.2 millimeters [mm] isotropic; repetition time [TR] = 1000 milliseconds [ms]; echo time [TE] = 22.00 ms; flip angle = 55 degrees; slices = 80; phase encoding direction = anterior to posterior; multi-band acceleration factor = 4; field of view = 176 mm) [45]. The volume position and angle were oriented to maximize subcortical, cortical, and cerebellar coverage. This was achieved by running a localizer scan (voxel size = 0.5 mm isotropic; TR = 8.6 ms; TE = 3.69 ms; flip angle = 20 degrees; slices = 1; phase encoding direction = anterior to posterior). The experimenter used the localizer image to adjust the BOLD volume location using the MRI scanner console graphic user interphase.

Once the volume location was specified, 3^rd^ order B0-shimming was completed to optimize B0-field homogeneity within this field of view. Specifically, we reconstructed phase images from a whole brain gradient echo image. We utilized the standard Siemens algorithm to compute B0 shim values to homogenize phase across the images and repeated this process three times. Next, we manually altered the linear shim voltages to minimize line width and computed scanning frequency.

After completing the BOLD recording part of the study that was run concurrently with the perception task (see *Afterimage and mock afterimage perception task* section), a high-spatial-resolution anatomical T1-weighted whole brain image (magnetization prepared – rapid gradient echo [MP2RAGE] [46]; voxel size = 0.8 mm isotropic; TR = 4300 ms; TE = 1.99 ms; inversion time (TI) 1 = 840 ms; TI 2 = 2370 ms; flip angle 1 = 5.0 degrees; flip angle 2 = 6.0 degrees; slices = 224; phase encoding direction = anterior to posterior) was acquired to assist in the preprocessing of the fMRI data (see *Whole brain fMRI* Analysis Methods section).

*V1 fMRI sequence*. V1 fMRI was acquired on a MAGNETOM 7T Plus MRI (Siemens, Inc., Healthineers, Erlangen, Germany) with a 32-channel head coil (Nova Medical Inc., Wilmington MA, USA). Simultaneous BOLD and VASO [47] imaging was recorded with a 3D-EPI readout (voxel size = 0.8 mm isotropic; volume acquisition sampling rate = 1589 ms; pair TR = 3178 ms; TE 1 = 23.50 ms; TI 1 = 1201.3 ms; TI 2 = 2293.9 ms; variable flip angles with reference [last] = 45 degrees; slices = 18; phase encoding direction = anterior to posterior; field of view = 150 mm) [48–50]. 3^rd^ order B0-shimming was completed to optimize B0-field homogeneity in visual cortex. The slab position and angle were oriented by the experimenter to be centered within and run parallel to each participant’s calcarine sulcus along the mid-sagittal plane (Figure 5A; Supplementary Figure 14). This was achieved by running a brief (∼10 seconds) 2D inversion recovery turbo flash sequence (voxel size = 1.0 x 1.0 x 8.0 mm; TR = 4300 ms; TE = 3.17 ms; TI 1 = 840 ms; TI 2 = 2540 ms; flip angle 1 = 5 degrees; flip angle 2 = 8.0 degrees; slices = 6; phase encoding direction = anterior to posterior). Based on this localizer, the experimenters manually adjusted the position and angle of the fMRI imaging slab using the MRI graphic user interface.

After completing the BOLD and VASO recording part of the study that was run concurrently with the perception task (see *Afterimage and mock afterimage perception task* section), a high-spatial-resolution anatomical T1-weighted whole brain image was acquired (MP2RAGE; see *Whole brain fMRI sequence* section) to assist in the segmentation, columnification, and layerification of the BOLD and VASO volumes (see *V1 fMRI* Analysis Methods section). For two participants, an MP2RAGE scan was not acquired during the V1 fMRI session. Instead, their MP2RAGE images acquired during the whole brain fMRI session were used for subsequent V1 fMRI analyses.

*Equipment, software, and facility.* The behavioral tasks were run on a behavioral laptop (MacBook Pro; 13-inch; 2560 x 1600 pixels, 2019; Mac OS Catalina v10.15.7; Apple, Inc.) and with PsychoPy (version 2022.2.24; Open Science Tools Ltd [51]). In the behavioral session, participants viewed the tasks (see *Afterimage and image perceptual reporting task* and *Afterimage and mock afterimage perceptual matching validation* sections) on a VIEWPixx monitor (1920 x 1200 pixels; refresh rate = 120 Hz; VPixx Technologies, Inc.) that was mirrored to the behavioral laptop screen. During the MRI sessions, participants laid supine, and a mirror mounted on the head coil positioned above their eyes reflected task images displayed on a screen (approximately 22 x 28.5 cm) located behind their head. The viewing distance between the participants and the center of the screen was approximately 64 cm. Task images were projected onto the screen using a rear projector system (PROPixx; 1920 x 1080 pixels; refresh rate = 480 Hz; VPixx Technologies, Inc.) connected to the behavioral laptop in the control room.

In both the behavioral and fMRI study sessions, the room lighting was made consistent across all participants. Participants responded during the tasks with their right hand using a keyboard (behavioral session: directly connected to the behavioral laptop) and button box (fMRI sessions: 4 Button Inline Fiber Optic Response Pad; Current Designs, Inc.). The button box was connected to the behavioral laptop via an electronic interface (932 Interface & Power Supply; Current Designs, Inc.).

During the behavioral sessions, the experimenter sat behind participants to deliver instructions and monitor task performance, eye tracking, and pupillometry recordings. In the fMRI sessions, the experimenters sat in a control room to monitor task performance, fMRI, eye tracking, and pupillometry recordings, and communicated with and monitored the participants via an intercom unit (Siemens, Inc.).

## Analysis Methods

All analyses were completed in Analysis of Functional NeuroImages (AFNI; version 25.1.07; https://afni.nimh.nih.gov; [52, 53]), Anatomical Normalization Tools (ANTs; https://stnava.github.io/ANTs; [54]), FreeSurfer (version 7.4.1; https://surfer.nmr.mgh.harvard.edu; [55]), ITK-SNAP (version 3.8.0; https://www.itksnap.org/pmwiki/pmwiki.php; [56]), LayNii toolbox (version 2.9.0; https://github.com/layerfMRI/LAYNII; [57]), MATLAB (version R2023b; https://www.mathworks.com/products/matlab.html; MathWorks, Inc.), and Statistical Parametric Mapping (SPM; version 12; https://www.fil.ion.ucl.ac.uk/spm). Visualizations were made using MATLAB, AFNI, Illustrator (version 2025; Adobe, Inc.), and Surf Ice (version 15.4.1; https://www.nitrc.org/projects/surfice; [58]).

### Behavior

Afterimage and mock afterimage perception rate was calculated as the number of afterimages and mock afterimages perceived following the inducer and mock inducer, respectively, versus all inducer and mock inducers presentations (Figure 1D; Supplementary Figure 2A). A perceived afterimage and mock afterimage were determined if (1) the participant reported both an onset and offset time, (2) the onset response was earlier than the offset response time, and (3) a mock afterimage was presented (i.e., not a blank trial) for the mock inducer trials. A not perceived afterimage and mock afterimage was determined if the participant did not report both an onset and offset time.

The duration and onset latency of perceived afterimages and mock afterimages was calculated by taking the difference between the reported onset and offset (duration) and subtracting the time between the offset of the inducer or mock inducer and the reported onset of the afterimage or mock afterimage, respectively (latency; Figure 1E; Supplementary Figure 2B).

Wilcoxon matched-pairs signed-rank tests (two-tailed) evaluated if (1) the duration and (2) onset latency of the afterimage and mock afterimage were statistically different across participants (Figure 1E; Supplementary Figure 2B). Simple linear regression and Pearson correlation tested if there was a significant relationship in the duration and onset latency of the afterimage and mock afterimage across participants (Figure 1F; Supplementary Figure 2C). Duration and onset latency variance were calculated for all perceived afterimages and mock afterimages within participants. Wilcoxon matched-pairs signed-rank tests (two-tailed) evaluated if (1) the duration and (2) onset latency variance of the afterimage and mock afterimage were statistically different across participants (Supplementary Figure 3). For all behavioral analyses, the whole brain and V1 fMRI sessions were evaluated separately.

Finally, we evaluated within-participant consistency in the duration and onset latency of afterimages and mock afterimages across the whole brain and V1 fMRI sessions. Specifically, simple linear regression and Pearson correlation tested if there was a significant relationship in the duration and onset latency of the afterimage and mock afterimage between the study sessions (Supplementary Figure 4). Only participants who completed both the whole brain and V1 fMRI study sessions were included in the analysis (N = 12).

### Eye tracking and pupillometry

*Preprocessing.* Pupil size, blinking, and microsaccade dynamics were evaluated for each of the afterimage and mock afterimage perception task trial conditions. First, pupil size data were preprocessed to remove blinks and artifacts using a custom software. In summary, possible blink and artifactual events (e.g., lost tracking or prolonged eye closure) were identified by finding samples with atypically small pupil size (pupil size equals 0 when the pupil is entirely occluded by the eyelid or when eye tracking is lost) or rapid changes in pupil size. Candidate blink events were confirmed based on several additional criteria, including blink duration and interblink duration thresholds. All blink and artifactual event samples were removed from the pupil size data and replaced by linear interpolation. Pupil size data preprocessing and blink detection quality was confirmed with visual inspection.

Microsaccade events were identified from the gaze position data using custom software that implemented standard methods for the detection of saccades and microsaccades [59, 60]. In brief, possible saccade and microsaccade events were determined based on gaze velocity and duration, and microsaccades were distinguished from saccades by the Euclidean distance between the initial and final gaze position of the eye movement – microsaccades defined as eye movements < 1 degree of visual angle. Saccade and microsaccade detection quality was confirmed with visual inspection.

The resulting processed datasets were: (1) preprocessed pupil size, (2) binary blink (0 = blink absent; 1 = blink present), and (3) binary microsaccade (0 = microsaccade absent; 1 = microsaccade present) arrays.

*Epoch extraction.* Pupil size, blink, and microsaccade epochs were segmented for each participant. Epochs were centered on the inducer and mock inducer onset times. Epochs were categorized according to the perception task trial conditions (see *Afterimage and mock afterimage perception task* Methods section): (1) inducer *without* an afterimage, (2) inducer *with* an afterimage, (3) mock inducer *without* a mock afterimage, and (4) mock inducer *with* a mock afterimage. Next, the mean across all epochs within participant, trial condition, and eye behavior type was calculated. For participants who completed both the whole brain and V1 fMRI studies, epochs were combined between sessions within participant, trial condition, and eye behavior type and then averaged. High frequency components were removed from the participant mean pupil, blink, and microsaccade epoch timecourses by smoothing (250 ms moving average span; also, a zero-phase, zero-temporal shift low-pass filter was tested that achieved similar results [data not shown]). Finally, the participant mean epoch timecourses were baselined to the mean signal among the 1000 ms prior to the inducer and mock inducer onset.

*Group-level analyses.* The eye tracking and pupillometry data were not analyzed for 2 participants who completed the whole brain fMRI study session due to poor eye tracking. Of the remaining 30 participants, group-level epoch timecourses were calculated by first standardizing or z-scoring the pupil size, blink fraction, and microsaccade fraction values across participants and then evaluating the mean and standard error of the mean across all participants and within trial condition (Supplementary Figure 5). A minimum of 5 epochs within a trial condition was required for a participant to be included in group-level analyses for that condition. This criterion resulted in the removal of 8 participants from inducers without afterimages condition who nearly always reported an afterimage following the inducer.

Wilcoxon matched-pairs signed-rank tests (two-tailed) evaluated differences in the mean pupil size, blink fraction, and microsaccade fraction within 4 epoch intervals: (1) baseline (–2-0 seconds preceding the inducer and mock inducer), (2) inducer and mock inducer (1-3 seconds following the inducer and mock inducer onset), (3) afterimage and mock afterimage (7-9 seconds following the inducer and mock inducer onset), and (4) post-afterimage and post-mock afterimage intervals (11-13 seconds following the inducer and mock inducer onset). These intervals were selected based on visual inspection of the group averaged pupil, blink, and microsaccade timecourses that indicated these timepoints involved the maximum and stable changes across all eye behaviors. Three contrasts were analyzed: (1) the inducers *with* afterimages versus inducers *without* afterimages, (2) mock inducers *with* mock afterimages versus mock inducers *without* mock afterimages, and (3) inducers *with* afterimages versus mock inducers *with* mock afterimages (Supplementary Figure 5). Holm-Bonferroni correction was applied across all statistical tests within each eye behavior type (4 tested intervals x 3 tested condition contrasts = 12 tests per eye behavior) to control for multiple comparisons.

### Whole brain fMRI

*Preprocessing.* The whole head anatomical image (MP2RAGE; see *Whole brain fMRI sequence* section) was segmented (FreeSurfer) and skull stripped. The skull stripped, segmented anatomical image was used for standard fMRI preprocessing applied on the BOLD volumes with AFNI’s afni_proc.py [61], including slice timing correction, head motion estimation, linear affine EPI-anatomical alignment using the lpc+ZZ cost function [62], nonlinear alignment of the anatomical to template space (volumes were registered to the MNI152_2009_template; [63]), smoothing (kernel size = 2 mm), whole brain masking, and motion regression (3 rotation and 3 translation). For improved computational efficiency, the BOLD voxel resolution was rescaled from 1.2 mm to 1.5 mm isotropic. Finally, percent change BOLD was calculated for each brain voxel within fMRI runs, which corresponded with the perception task blocks (see *Afterimage and mock afterimage perception task* section) relative to the mean voxel-wise BOLD responses across all run volumes. The preprocessed data were quality checked to verify that participant motion was relatively low and did not appear correlated with the task and to confirm that the BOLD, anatomical, and template volumes were in alignment [64].

*Epoch extraction.* BOLD voxel-by-time epochs were segmented. Epochs were centered on the inducer and mock inducer stimulus onset times (see *Afterimage and mock afterimage perception task* section). Specifically, 25 volumes (25 seconds) before and after the inducer and mock inducer onset TR were extracted. This epoch interval, while exceeding the perception task trial interval (see *Afterimage and mock afterimage perception task* Methods section), was selected to account for expected hemodynamic delays of BOLD associated with task events. In rare cases, when the epoch duration exceeded the onset or offset of a run, the missing volume values were replaced with “not-a-number” (MATLAB).

Since contrasts between task conditions were made on a timepoint-by-timepoint basis, for trials with an afterimage and mock afterimage, a temporal normalization procedure was applied to account for the variable duration and onset latency among afterimages and mock afterimages. Based on the group-level onset latency and duration reports (Figure 1E; Supplementary Figure 2B), all epochs were fit to an onset latency of 2 seconds (2 TRs) and an afterimage and mock afterimage duration of 4 seconds (4 TRs). Epochs were resampled with linear interpolation to match these standard onset latency and duration intervals. For example, an epoch where an afterimage was reported as lasting for 2 seconds was upsampled to 4 seconds (i.e., the addition of two time points was made between the afterimage onset and offset volumes). Meanwhile, if the afterimage duration exceeded the standard duration, downsampling was achieved by averaging volumes adjacent in time. Temporal normalization was not applied on inducer and mock inducer intervals because they had a constant duration (4 seconds; 4 TRs).

Visual inspection of the temporal normalized data confirmed that this procedure did not introduce artifacts. As an additional confirmation, we conducted whole brain BOLD analyses (see *Whole brain spatiotemporal cluster-based permutation testing* section) *without* temporal normalization (Supplementary Figure 6). These complementary analyses affirmed our findings from the temporal normalization dataset.

Finally, all epochs within each trial condition were averaged within participant resulting in a voxel-by-time (TR) per condition BOLD volume. To increase sample size, the inducers without afterimages and mock inducers without mock afterimages epochs were combined and averaged on the epoch-level basis to define the “inducers + mock inducers only” epoch condition. Combining the inducers and mock inducers only epochs was justified following analyses that confirmed inducers and mock inducers evoked similar whole brain BOLD responses (see *Whole brain BOLD signals* Results section; Supplementary Figure 8).

*Whole brain spatiotemporal cluster-based permutation testing.* Spatiotemporal cluster-based permutation testing (two-tailed; 1000 permutations; cluster forming and cluster level *p* < 0.05; baseline interval = 5 seconds pre-inducer and mock inducer onset) was applied on the whole brain BOLD data focusing on the 0-25 seconds post-inducer and mock inducer onset. Full details on the permutation method are available at [65]. Briefly, the cluster-based permutation analysis built a null distribution by random permutation of the trial conditions and a cluster-forming statistic based on spatial and temporal adjacency defined as when statistically significant voxels of the same sign are adjacently located in space or time (i.e., voxels that are active over consecutive time points, active voxels that neighbor other active voxels at individual time points, or both). Spatiotemporal clusters were then identified in the non-permutated data and evaluated against the null distribution to find statistically significant spatiotemporal clusters. The final output of this analysis was the signed, statistically significant voxels at each queried timepoint. To improve computational efficiency, only gray matter voxels were tested, and the BOLD volume spatial resolution was downsampled to 3.0 mm isotropic.

Cluster-based permutation analysis results were visualized by plotting the whole brain *t*-value map with sub-threshold *t*-value voxels shown with transparency and above threshold voxels highlighted with a black outline (e.g., Figure 2A, C, and E) [66, 67]. Brain surface visualization was made with Surf Ice by projecting voxel values to a cortical surface mesh (BrainMesh_ICBM152; e.g., Figure 2B, D, and F). Only voxels that were found statistically significant by cluster-based permutation testing were projected to the surface.

**Figure 2.**
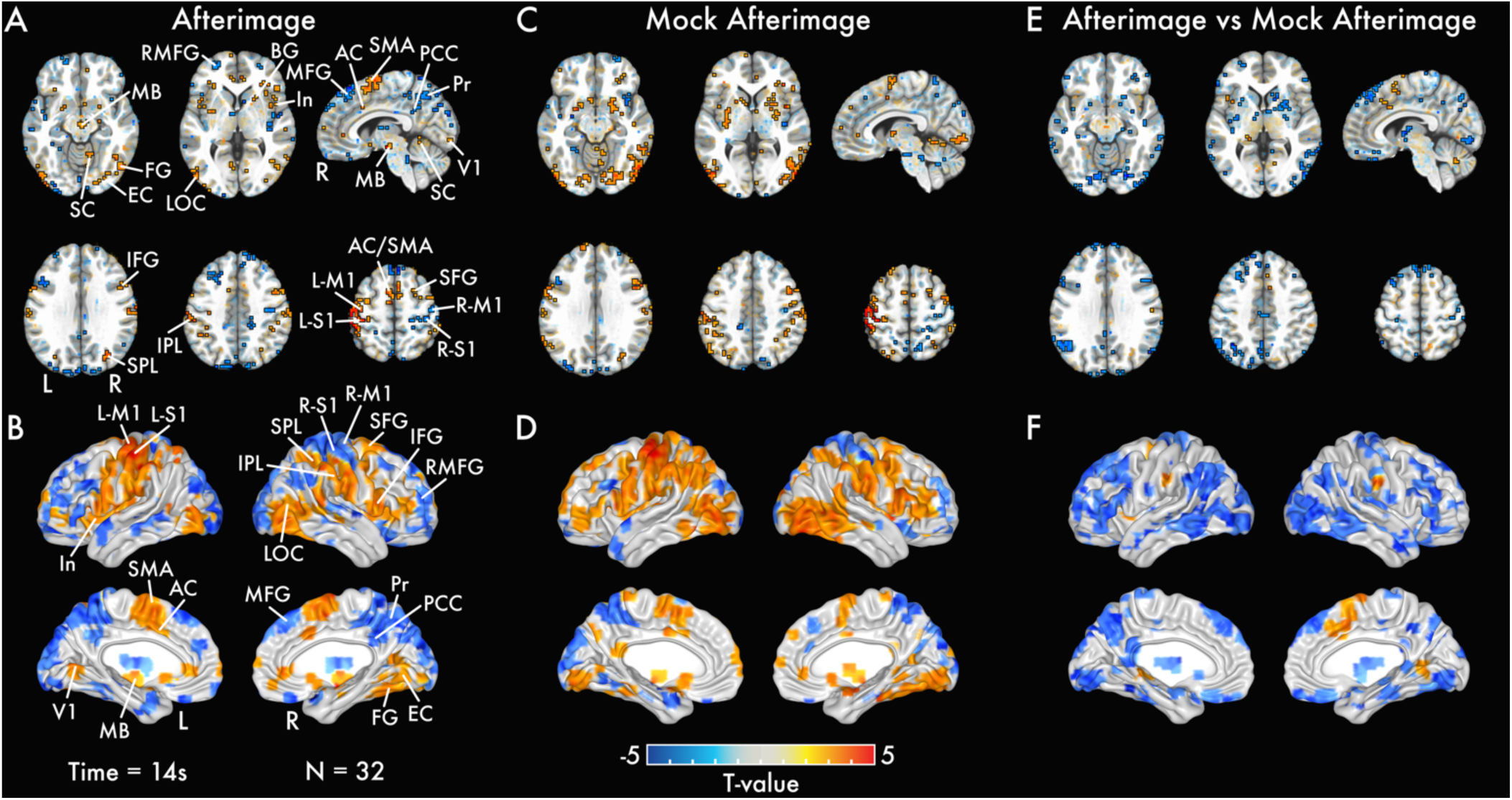
Whole brain volume and surface BOLD activation maps. Statistically significant *t*-values are highlighted with a black outline, while sub-threshold regions are shown with transparency at time point 14 seconds (s) after the inducer and mock inducer onset – the approximate peak response time for afterimages and mock afterimages – for the following statistical contrasts: **(A)** inducers *with* afterimages versus inducers + mock inducers only conditions (*afterimage*), **(C)** mock inducers *with* mock afterimages versus inducers + mock inducers only conditions (*mock afterimage*), and **(E)** inducers *with* afterimages versus mock inducers *with* mock afterimages conditions (*afterimage vs mock afterimage*). Brain surface visualization of *t*-values from statistically significant voxels are shown in **(B)**, **(D)**, and **(F)** for the same contrasts highlighted in the volume activation maps. See Supplementary Slides 1, 2, and 3 for volume activation maps for additional time points (0 to 25 s after the inducer and mock inducer onset). Anterior cingulate cortex (AC); basal ganglia (BG); extrastriate cortex (EC); fusiform gyrus (FG); inferior frontal gyrus (IFG); inferior parietal lobule (IPL); insula (In); lateral occipital cortex (LOC); left primary motor cortex (L-M1); left primary somatosensory cortex (L-S1); medial frontal gyrus (MFG); midbrain (MB); rostral middle frontal gyrus (RMFG); right primary motor cortex (R-M1); right primary somatosensory cortex (R-S1); precuneus (Pr); primary visual cortex (V1); posterior cingulate cortex (PCC); superior cerebellum (SC); superior frontal gyrus (SFG); supplementary motor area (SMA); superior parietal lobule (SPL).

Three main tests were explored: (1) inducers *with* afterimages versus inducers + mock inducers only (unilateral and bilateral visual field analyses), (2) mock inducers *with* mock afterimages versus inducers + mock inducers only (unilateral and bilateral visual field analyses), and (3) inducers *with* afterimages versus mock inducers *with* mock afterimages (unilateral visual field analyses only). In unilateral analyses, epochs corresponding to left versus right visual field inducer or mock inducer presentations were evaluated separately. Independently testing the left and right-sided stimuli presentations was done to assess contralateral and ipsilateral signaling dynamics. In bilateral analyses, epochs corresponding to left and right visual field inducer and mock inducer presentations were evaluated together.

Additional analyses were explored included: (1) inducers *with* afterimages versus inducers *without* afterimages, (2) mock inducers *with* mock afterimages versus mock inducers *without* mock afterimages, (3) inducers *without* afterimages versus baseline, (4) mock inducers *without* mock afterimages versus baseline, and (5) inducers *without* afterimages versus mock inducers *without* mock afterimages. 14 participants were excluded from these analysis because of an insufficient number of inducers without afterimages epochs (minimum threshold of 5 epochs per condition). For consistency, the same participants were excluded from the mock inducers with and without mock afterimages analyses. Note that only 8 participants were removed by this exclusion criterion in the eye tracking and pupillometry data analysis because epochs were combined within participant between the whole brain and V1 fMRI sessions.

*between model analysis.* To assess the statistical robustness of the whole brain cluster-based permutation analyses, a standard fMRI general linear model (GLM) analysis was conducted (AFNI 3dDeconvolve). In summary, beta coefficients were estimated for task events by building a model for each participant with task condition regressors (inducer, mock inducer, afterimage, and mock afterimage onsets), an eye blink onset regressor, and motion regressors (3 rotation and 3 translation parameters). Next, the voxel-time beta coefficient volumes for the afterimage and mock afterimage were analyzed across participants with cluster-based permutation analysis (baseline interval = 8 seconds at the end of the modeled timecourse which was determined by visual inspection to be an interval that followed the afterimage and mock afterimage response; see *Whole brain spatiotemporal cluster-based permutation testing* section).

The beta coefficient cluster-based permutation analysis results were visualized by plotting the whole brain *t*-value map with sub-threshold *t*-value voxels shown with transparency and above threshold voxels highlighted with a black outline (Supplementary Figure 7A, C, and E) [66, 67]. Brain surface visualization was made with Surf Ice by projecting voxel values to a cortical surface mesh (BrainMesh_ICBM152; Supplementary Figure 7B, D, and F). Only voxels that were found statistically significant by cluster-based permutation testing were projected to the surface.

Three tests were explored corresponding with the spatiotemporal cluster-based permutation testing analyses: (1) afterimages versus baseline, (2) mock afterimages versus baseline, and (3) afterimages versus mock afterimages (Supplementary Figure 7). The beta coefficient volumes were downsampled (3.0 mm isotropic) to match the spatial resolution of the cluster-based permutation testing analyses. 7 participants were excluded from the GLM analysis because they did not have reliable pupillometry data in one or more task blocks (e.g., eye tracking was intermittently lost during the study session). High quality pupillometry data is necessary to evaluate blinks (see *Eye tracking and pupillometry* section), thereby a blink regressor could not be specified in the GLM model for these participants. In total, 25 participants were included in the whole brain GLM analysis.

*K-means clustering.* K-means clustering was evaluated on the 25 seconds following the inducer and mock inducer onset using correlation as the distance measure. K-values between 2 and 5 were tested (Supplementary Figure 11). K-means cluster silhouette scores (i.e., for each voxel, the ratio of its mean distance to voxels in its own cluster versus voxels in its nearest other cluster), BOLD timecourses, and brain maps were evaluated to interpret cluster results. Only a subset of whole brain voxels was included in the k-means clustering analysis. Voxels were included in k-means clustering if they showed statistically significant BOLD activity from either the afterimage and mock afterimage condition analyses (Figure 2A and C; see *Whole brain BOLD cluster-based permutation testing* section) at any time within the 19-second period from the afterimage and mock afterimage onset, regardless to whether the BOLD signal change was positive or negative. To correspond with the spatial resolution of the whole brain statistical analyses (see *Whole brain spatiotemporal cluster-based permutation testing* section), k-means clustering was also applied on the 3.0 mm isotropic downsampled BOLD volumes. All colored areas in Figure 3A and Supplementary Figures 11A and 12A and B represent the voxels included in the k-means analysis (i.e., all non-colored areas were excluded from analyses).

**Figure 3.**
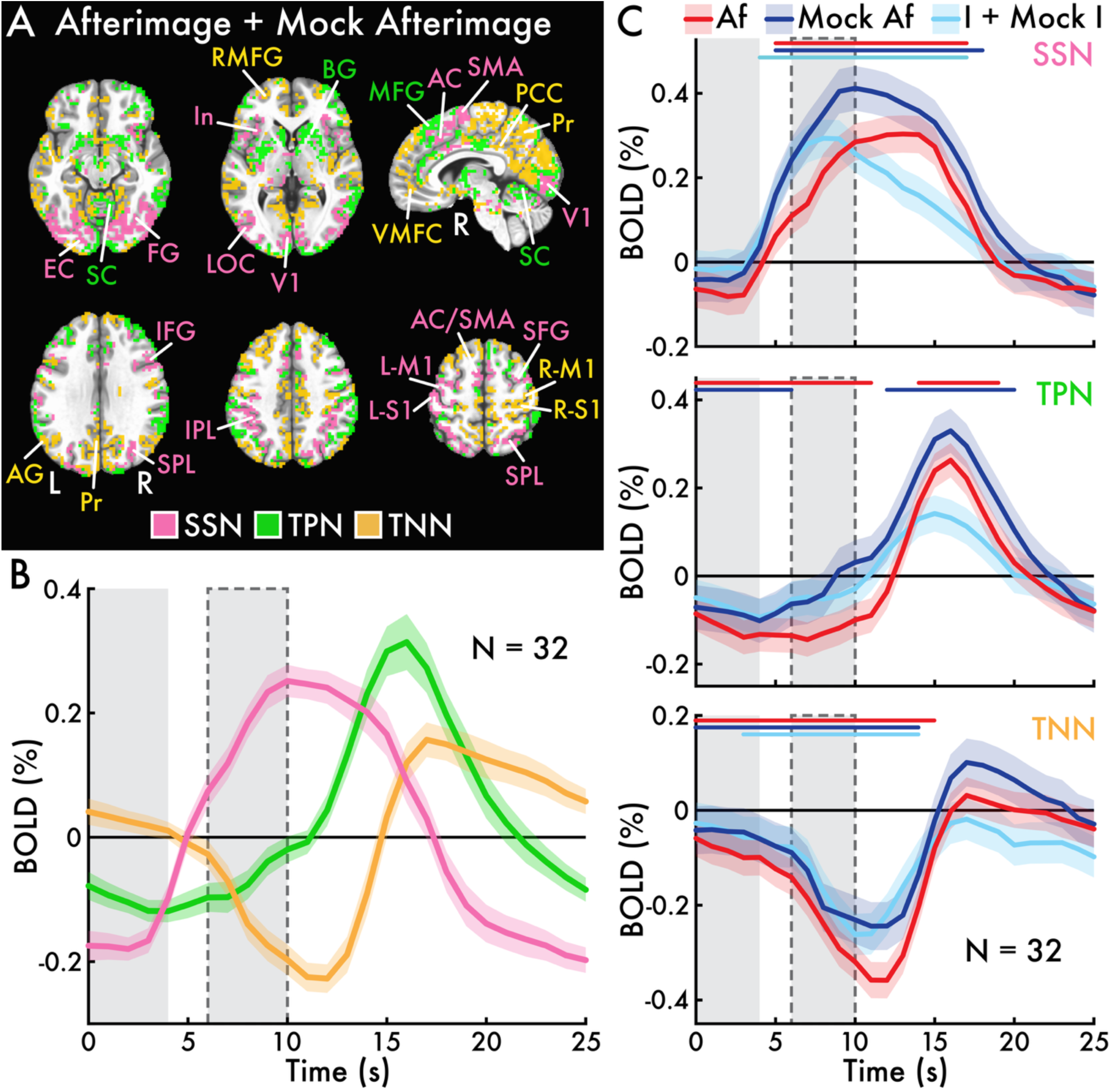
Whole brain BOLD spatiotemporal clustering and network timecourses. **(A)** K-means clusters (k = 3) of the average responses between the inducers *with* afterimages and mock inducers *with* mock afterimages conditions (*afterimage + mock afterimage*) for gray matter brain regions that showed statistically significant change in either the inducers *with* afterimages or mock inducers *with* mock afterimages conditions. The clusters corresponded with established functional brain networks: (1) *salience and sensory* (SSN), (2) *task positive* (TPN), and (3) *task negative* (TNN) networks. The network labels approximate the functional role of the brain regions grouped within each cluster. Major anatomical regions are labelled and color-coded by network. **(B)** The mean percent change blood-oxygen-level-dependent (BOLD; %) timecourses with standard error of the mean (SEM; shaded area) across all voxels within the SSN, TPN, and TNN. **(C)** The average BOLD response with SEM (shaded area) within the SSN, TPN, and TNN networks for the main trial conditions of the perception task (Figure 1C): (1) inducers *with* afterimages (*Af*), (2) mock inducers *with* mock afterimages (*Mock Af*), and (3) inducers *without* afterimages and mock inducers *without* mock afterimages conditions (inducers + mock inducers only; *I + Mock I*). Statistically significant intervals are highlighted with a horizontal bar above each timecourse, color-coded by condition. In B and C, the solid gray bars highlight the inducer and mock inducer period, while the solid gray bar with a dotted outline highlights the approximate afterimage and mock afterimage period. Angular gyrus (AG); anterior cingulate cortex (AC); basal ganglia (BG); extrastriate cortex (EC); fusiform gyrus (FG); inferior frontal gyrus (IFG); insula (In); lateral occipital cortex (LOC); left primary motor cortex (L-M1); left primary somatosensory cortex (L-S1); medial frontal gyrus (MFG); right primary motor cortex (R-M1); right primary somatosensory cortex (R-S1); precuneus (Pr); primary visual cortex (V1); posterior cingulate cortex (PCC); inferior parietal lobule (IPL); rostral middle frontal gyrus (RMFG); superior cerebellum (SC); superior frontal gyrus (SFG); supplementary motor area (SMA); superior parietal lobule (SPL); ventral medial prefrontal cortex (VMFC).

Clustering was performed on the group-averaged signal from both the inducers *with* afterimages and mock inducers *with* mock afterimages conditions (afterimage + mock afterimage; Figure 3; Supplementary Figure 11). For this analysis, signals between conditions were first averaged within participant and then averaged across all participants. This approach allowed for the identification of a common set of brain networks that could be compared across task conditions. The afterimage + mock afterimage analysis was justified after observing similar whole brain results for the inducers *with* afterimages and mock inducers *with* mock afterimages conditions (see *Whole brain BOLD signals* Results section; Figure 2), and that separately clustering on the inducers *with* afterimages and mock inducers *with* mock afterimages conditions resulted in similar cluster profiles (Supplementary Figure 12A and C versus B and D).

All voxels grouped into a common cluster were assigned a constant value to visualize the cluster spatial extent (Figure 3A; Supplementary Figures 11A and 12A and B). K-means cluster BOLD activity timecourses were evaluated by averaging the percent change BOLD signal across all voxels within each cluster at each time point (Figure 3B; Supplementary Figures 11B and 12C and D).

In addition, BOLD timecourses were calculated for each task condition – afterimage, mock afterimage, and inducers + mock inducers only (Figure 1C) – using each k-means cluster as a network region of interest. This was achieved by averaging the percent change BOLD signal across all voxels within each cluster separately for each task condition (Figure 3C). These cluster-based, task condition timecourses were statistically evaluated against a null response across all time points using temporal cluster-based permutation testing (two-tailed; 1000 permutations; cluster forming and cluster level *p* < 0.05; [68]).

*Regions of interest.* Regions of interest (ROIs) analyses were performed to evaluate BOLD signal responses over time across task conditions in task-engaged brain areas. ROIs were selected based on whole brain BOLD analyses and k-means clustering that highlighted sensory, salience, and motor network regions as responsive to afterimages and mock afterimages (Figure 2 and 3; see *Whole brain BOLD signals* and *Whole brain functional networks* Results sections). From these areas, 6 representative ROIs were selected: (1) LGN, (2) V1, (3) FG, (4) anterior cingulate cortex (AC), (5) insula (In), and (6) left primary motor cortex (M1) (Figure 4A). ROI analyses were conducted on the original, non-downsampled BOLD volumes with a spatial resolution of 1.5 mm isotropic.

**Figure 4.**
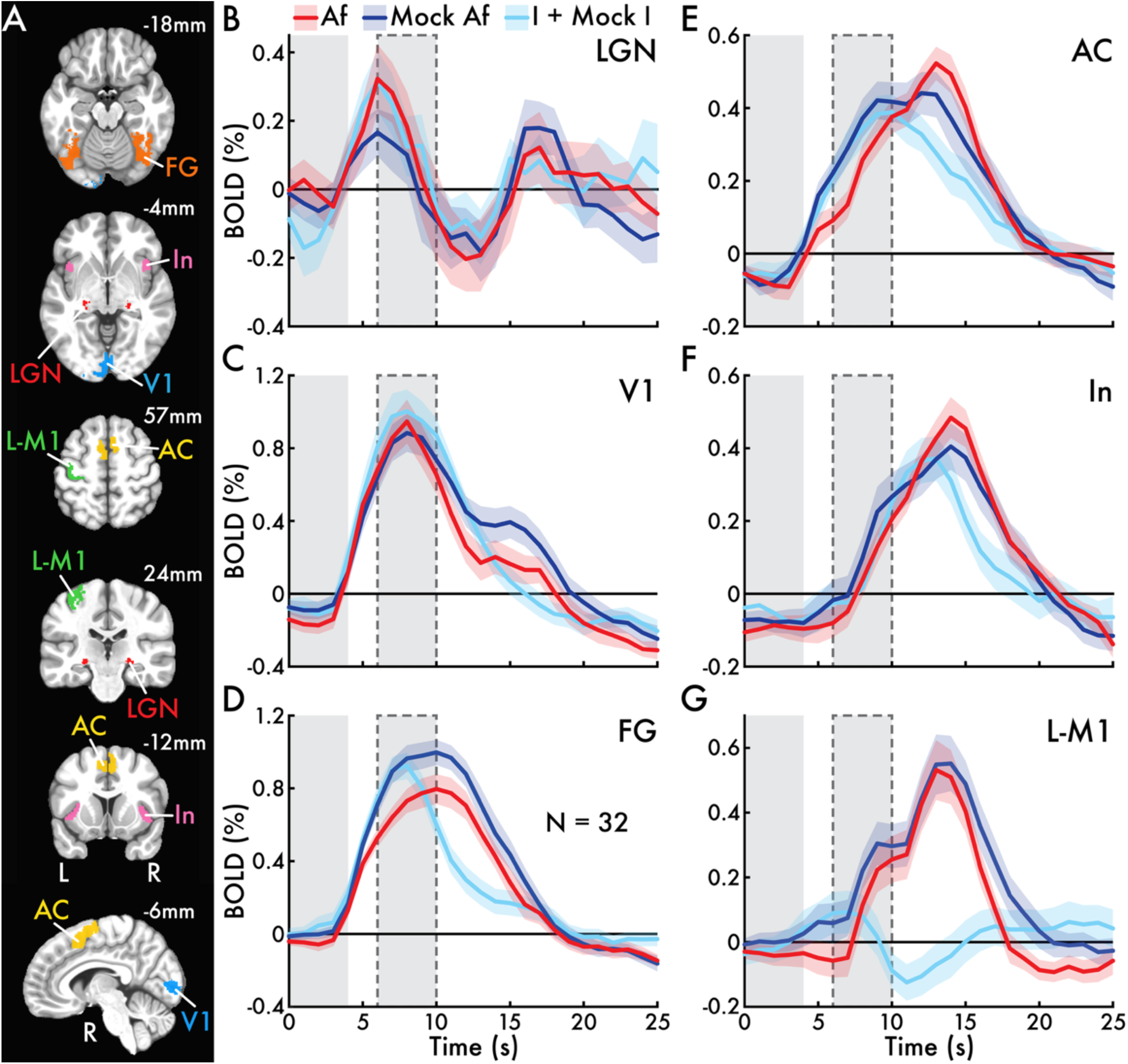
Sensory and salience network regions of interest BOLD timecourses. Regions of interest (ROIs) were selected based on their inclusion in the sensory and salience network (Figure 3A). **(A)** Visualizations of the ROIs with Montreal Neurological Institute coordinates in millimeters (mm), indicating their position in the axial, coronal, and sagittal planes. **(B)** Lateral geniculate nucleus (LGN), (**C**) primary visual cortex (V1), **(D)** fusiform gyrus (FG), **(E)** anterior cingulate (AC), **(F)** insula (In), and **(G)** left primary motor cortex (L-M1) mean blood-oxygen-level-dependent (BOLD) timecourses with standard error of the mean (shaded area) were calculated across participants (N = 32) for the main trial conditions of the perception task (Figure 1C): (1) inducers *with* afterimages (*Af*), (2) mock inducers *with* mock afterimages (*Mock Af*), and (3) inducers *without* afterimages and mock inducers *without* mock afterimages conditions (inducers + mock inducers only; *I + Mock I*). In all BOLD timecourse subplots, the gray bar between 0 and 4 seconds (s) highlights the inducer and mock inducer period. The solid gray bar with a dotted outline between 6 and 10 s highlights the approximate afterimage and mock afterimage period.

**Figure 5.**
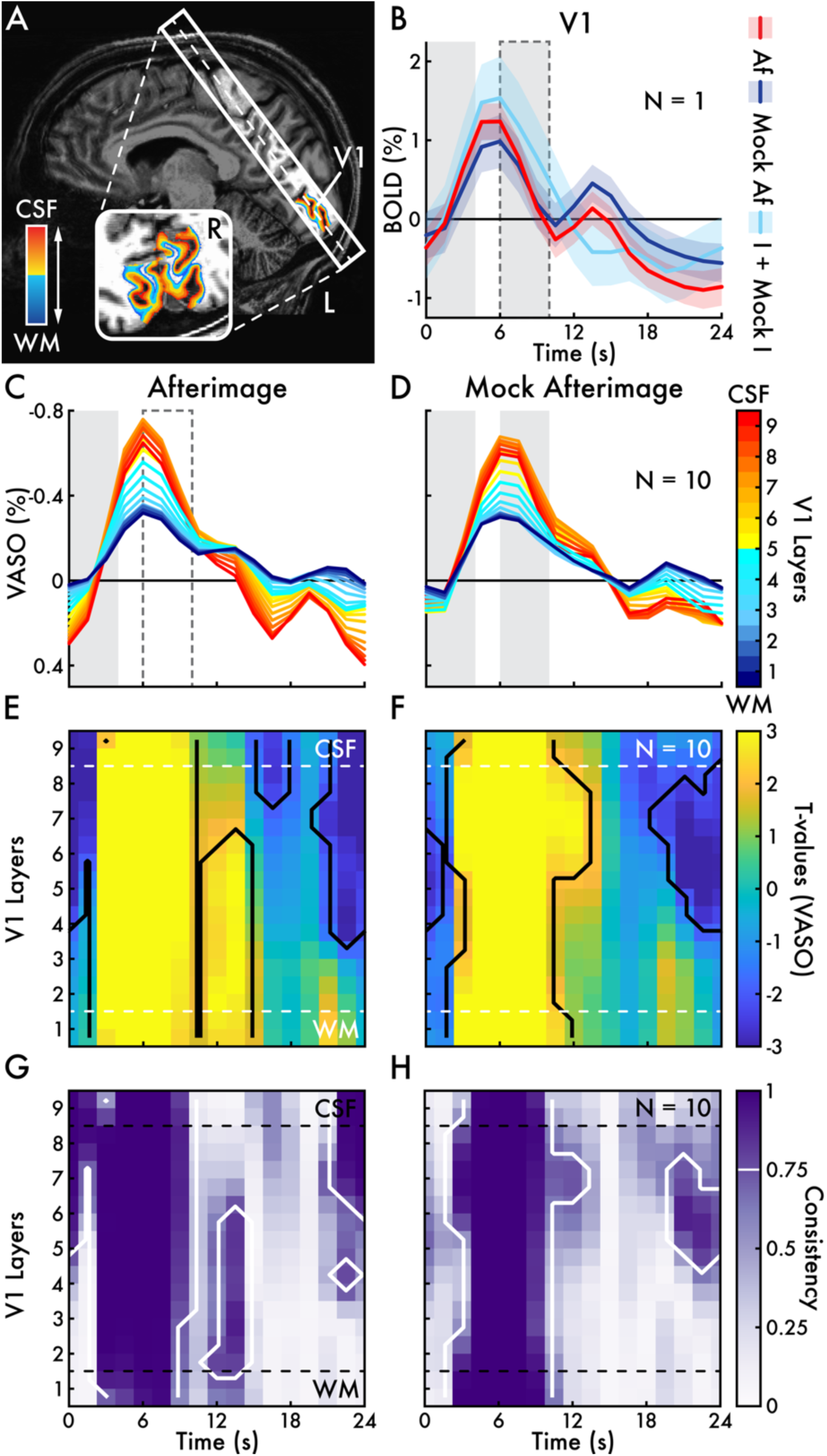
Primary visual cortex layer-dependent VASO activity. **(A)** Representative participant sagittal and axial anatomical brain images and fMRI slab highlighting the layer segmentation in primary visual cortex (V1; dark red = superficial cortical layers near cerebrospinal fluid [CSF]; dark blue = deep cortical layers near the cortical white matter [WM]). The dotted line in the sagittal view approximates the position of the coronal image. **(B)** The mean percent change blood-oxygen-level-dependent (BOLD; %) timecourses with standard error of the mean (shaded area) for all voxels in the V1 region of interest for the participant highlighted in A across the main trial conditions of the perception task (Figure 1C): (1) inducers *with* afterimages (*Af*), (2) mock inducers *with* mock afterimages (*Mock Af*), and (3) inducers *without* afterimages and mock inducers *without* mock afterimages conditions (inducers + mock inducers only; *I + Mock I*). These timecourses resemble those in V1 from group-level whole brain BOLD analyses (Figure 4C). The percent change vascular space occupancy (VASO; %) is shown across V1 cortical layers for **(C)** inducers *with* afterimages (*afterimage*) and **(D)** mock inducers *with* mock afterimages (*mock afterimage*). The percent change VASO signal was analyzed and shown inverted (negative is up) to enhance clarity and coherence with the BOLD signal. In B, C, and D, the gray bar between 0 and 4 seconds (s) highlights the inducer and mock inducer period. The dotted open bar in C, solid gray bar in D, and solid gray bar with a dotted outline in B between 6 and 10 s highlights the approximate afterimage and mock afterimage period. Subplots **(E)** and **(F)** show the *t*-values for cortical layer-time cluster-based permutation testing. Statistically significant layer-time clusters are outlined with a solid black line. Subplots **(G)** and **(H)** show consistency maps from bootstrap analyses (5000 resamples) of the cortical layer-time cluster-based permutation testing (see *V1 fMRI* Analysis Methods section). All layer-time clusters with a consistency proportion > 0.75 (i.e., the layer-time cluster was statistically significant for more than 75% of bootstrap iterations) are outlined with a solid white line.

The LGN voxels were defined based on each participant’s estimated LGN area according to anatomical whole brain segmentation completed during whole brain fMRI preprocessing (see Whole brain fMRI *Preprocessing* Methods section). To allow for group-level analyses, the participant-level LGN ROIs were normalized to a common template space and LGN voxels were combined across participants to define a group LGN region. For the remaining ROIs, voxels were selected according to the Montreal Neurological Institute Glasser atlas [69].

To identify the most task-engaged voxels within the sensory and left M1 ROIs, within-ROI functional localization was performed. This was achieved by running spatiotemporal cluster-based permutation testing (two-tailed; 1000 permutations; cluster forming and cluster level *p* < 0.05; test interval = 0-12 seconds) that evaluated *left* versus *right visual field* inducers + mock inducers only epochs for the LGN, V1, and FG ROIs. This analysis evaluated contralateral versus ipsilateral activity independent of motor response (i.e., participants did not make a button press for the inducers + mock inducers only condition). The contralateral relationship between visual field and visual sensory regions meant that the left versus right visual field contrast would highlight *right* hemispheric regions, and vice versa.

For the left M1 ROI, a statistical contrast between the afterimage + mock afterimage (averaged between conditions within participants) versus the inducers + mock inducers only condition was evaluated because the latter did not involve button presses, while the former included right-handed responses to indicate the onset and offset of afterimages and mock afterimages. Thereby, this contrast was sensitive to task motor responses.

For all ROI functional localization analyses, a voxel was included in the ROI if it met 2 criteria: (1) it was statistically significant at any time point between 0-12 seconds after the inducer or mock inducer onset, and (2) its average voxel *t*-value across those significant time points was *greater than 0*, indicating a tendency toward increased BOLD activity. Voxels that did not meet these criteria were discarded from the functionally localized ROIs. Finally, timecourses were visualized within each ROI by taking the mean percent change BOLD response across all selected ROI voxels and subtracted from the mean signal in the 5 seconds preceding the inducer and mock inducer onset. Each trial condition was evaluated separately and the mean timecourses are shown with standard error of mean calculated across participants (Figure 4).

### V1 fMRI

Analysis of the layer-resolution V1 fMRI data involved several processing stages detailed below. A schematic summary of the analysis pipeline is depicted in Supplementary Figure 13.

*Preprocessing.* V1 fMRI data acquisition involved simultaneous collection of BOLD and VASO images using interleaved TRs (see *V1 fMRI sequence* section). The VASO volumes were initially acquired as unprocessed cerebral blood volume-weighted or “nulled” signals. The final VASO signal was derived from these nulled images as an output of the processing procedure.

Each run consisted of 500 volumes that were sorted into 2 image sets: 250 BOLD volumes and 250 nulled volumes. The first 3 volumes of each set were removed to allow magnetization to reach steady state. Based on the remaining 247 volumes within each image set, 3 reference volumes were created: (1) BOLD, (2) nulled, and (3) T1-reference volumes. The BOLD and nulled reference volumes were calculated by averaging the volumes from the *first* fMRI run. The T1-reference volume was calculated by dividing the nulled reference image by the BOLD reference image, resulting in a T1-weighted contrast image used for registering the functional volumes to anatomical space. The first run was used to generate these reference images because it was least affected by motion artifacts.

Subsequently, the BOLD and nulled volumes were motion corrected with non-linear alignment to their respective reference images (ANTs). To reduce thermal noise, NOise Reduction with DIistribution Corrected (NORDIC) principal component analysis was implemented separately on the motion-corrected nulled and BOLD volumes [70–72]. BOLD and nulled volumes were temporally upsampled from 247 to 494 volumes using linear interpolation. The VASO signal was then calculated by performing a volume-by-volume division of the nulled signals from the BOLD signals [49]. Finally, BOLD and VASO percent signal change was calculated for each run relative to the mean signal of each voxel across all volumes within a run.

*Layerification & columnification.* A critical stage in layer-resolution fMRI analysis involves segmenting gray matter voxels within the ROI into layers and columns – a process known as “layerification” and “columnification”. Importantly, in this context, “layers” and “columns” do not correspond to precise anatomical or functional boundaries in neural tissues such as the 6 histological layers of human cortical gray matter or V1 ocular dominance columns. Rather, layers and columns refer to spatial segmentations of the gray matter that are aligned with these anatomical orientations. When spatial resolution is sufficiently high, these segmentations support inferences of the functional properties that are characteristic of the underlying neural tissue architecture.

Layer and column segmentation was performed for each participant, beginning with bias field correction on the whole brain anatomical image (MP2RAGE; SPM). Brain tissue segmentation was then performed (FreeSurfer) and mapped to the cortical surface (AFNI’s Surface Mapping tool; https://afni.nimh.nih.gov/Suma). The pial, gray, and white matter surfaces were extracted for each hemisphere and combined into a gray matter rim volume (Supplementary Figure 13B). Manual corrections were made to address segmentation errors (e.g., inclusion of the sagittal sinus in the gray matter rim or “kissing gyri” where adjacent gyri lack visible cerebral spinal fluid [CSF] separation). These edits were made using a digital pad (Wacom Intuos CTL4100; Wacom Co., Ltd; ITK-SNAP). The corrected gray matter rim was then upsampled using linear interpolation to a high-spatial-resolution grid (0.27 mm isotropic) to refine tissue boundaries and match the spatial resolution of the T1 reference volume.

The gray matter rim was next registered to the functional space (BOLD and VASO volumes) using the bias field corrected anatomical image and the T1 reference volume (ANTs). To enhance registration accuracy, a manually drawn occipital cortex mask based on the T1 reference volume was used to constrain the registration. The mask included visual cortex gray and white matter but avoided the upper and lower 2 slices of the acquired volume because of the risk for slab misalignment.

Finally, cortical layers and columns were segmented (LayNii toolbox). Cortical layers were defined by parcellating the gray matter rim into 5 equal-volume parcels, with 2 additional layers of equal thickness (0.3 mm) added just above and below the gray matter boundary (LN2_LAYERS; Supplementary Figure 13C). This resulted in a 9-layer padded gray matter rim (i.e., the upper and lower layers extended into white matter and CSF) that was used in subsequent layer-level epoch extraction (see details below). Cortical columns were defined by parcellating the padded gray matter rim across the entire slab into 5000 approximately equal-volume compartments (LN2_COLUMNS; Supplementary Figure 13D).

*Volume-level epoch extraction.* Perception task event epochs (see *Afterimage and mock afterimage perception task* Methods section) were segmented across the entire V1 fMRI acquisition volume. First, task event times (e.g., inducer stimulus onset and button presses) were identified for each trial and categorized according to the 3 main task conditions of interest (Figure 1C): (1) afterimage, (2) mock afterimage, and (3) inducers + mock inducers only. Next, events times were affiliated with their nearest TR to account for the interleaved acquisition of BOLD and VASO volumes. Therefore, each event time was separately matched to its nearest BOLD and VASO TR, which could correspond to different volumes in each image series. Epochs were then extracted from the percent change volumes for both BOLD and VASO. Epochs were centered on the nearest TR to the inducer and mock inducer onset and included 20 TRs (∼32 seconds) before and after the center TR to account for hemodynamic delays.

Some VASO epochs were observed to include large, artifactual spikes. These epochs were identified by computing the first derivative of the average VASO signal across all voxels within the V1 ROI (see *V1 Region of Interest* section). Any time point at or after the onset of the inducer or mock inducer where the first derivative value exceeded 5 percent (this threshold selected based on visual inspection), independent to signal direction, was marked as artifactual, and the corresponding epoch was excluded from further analysis. Across participants, no more than 20 percent of VASO epochs were excluded per task condition based on this criterion. No BOLD epochs were removed. Finally, a separate set of BOLD inducers + mock inducers only condition epochs were spatially smoothed (gaussian blur = 2.0 mm full width at half maximum) to assist in V1 ROI functional localization (see *V1 Region of Interest* section).

A supplementary analysis of eye blink V1 cortical layer responses was performed to account for the contribution of blinking in these results (Supplementary Figure 20). This was achieved by identifying blinks from the pupillometry data (see *Eye tracking and pupillometry* section). To isolate blink-specific activity from task events, blinks were selected for analysis if they occurred during task interstimulus intervals (> 3 TRs or ∼4.5 seconds from the onset of the inducers or mock inducers) and blinks with a minimum interblink interval of 4 TRs (∼6 seconds), so that multiple blinks occurring within a short period were not analyzed as independent events. These parameters resulted in most participants contributing > 100 blinks for analysis. Subsequently, the same epoch extraction approach was implemented as with the main task events, including affiliating the blink onset times with their nearest BOLD and VASO TRs, segmenting percent change BOLD and VASO volume epochs 20 TRs (∼32 seconds) before and after the blink onset TR, and rejecting blink VASO epochs that included artifactual spikes.

*V1 region of interest.* The V1 ROI was functionally localized for each participant based on the BOLD responses to the inducers + mock inducers only condition. First, spatially smoothed BOLD epochs *without* afterimages or mock afterimages were grouped by left versus right visual field presentation location. Smoothed BOLD signals were used to facilitate visual inspection of the V1 activation area (Supplementary Figure 13E). Next, all inducers + mock inducers only epochs were averaged separately within the left and right visual field conditions. The peak activation time in contralateral V1 linked to the inducer and mock inducer presentation was determined by visual inspection. At this peak time point, the boundaries of the V1 activation area in the contralateral visual cortex were manually delineated (AFNI). The resulting left and right V1 ROIs were then registered to the gray matter rim (see *Layerification & columnification* section) by resampling them into the layer and column voxel grid space (0.27 mm isotropic). To ensure an approximately equal representation of cortical layers within the V1 ROIs, voxels were related to the gray matter rim columns. If any manually drawn voxel overlapped with a column, all voxels within that column were included in the ROI. Finally, manual corrections were made to exclude any erroneously included columns from the contralateral hemisphere. The resulting left and right V1 ROIs were used for subsequent layer-dependent analyses (Supplementary Figure 13F; see Supplementary Figure 14 for the combined left and right V1 ROI from all participants included in the V1 fMRI analysis).

*Layer-level epoch extraction.* Epochs were initially segmented from the entire V1 fMRI acquisition volume (see *Volume-level epoch extraction* section). To enable layer-dependent analysis, these volume-level epochs were resampled into the layer and column grid space (0.27 mm isotropic). Within the V1 ROIs (see *V1 regions of interest* section), the mean BOLD and VASO signals were calculated separately for each cortical layer at each time point. This was achieved by finding the layer-specific voxels within the *contralateral* V1 ROI for each visual field condition (i.e., the right hemisphere ROI voxels were analyzed for epochs corresponding to the left visual field stimulus presentation, and the left hemisphere ROI voxels for the right visual field presentation; Supplementary Figure 13G). The signals from the contralateral voxels were then averaged across all contralateral conditions to provide a bilateral activation profile based on contralateral response dynamics.

*V1 region of interest BOLD timecourses.* The mean V1 ROI BOLD timecourses for the inducers *with* afterimages, mock inducers *with* mock afterimages, and inducers + mock inducers only conditions were visualized for 1 representative participant by taking the mean BOLD response across all V1 ROI voxels (Figure 5B). The standard error of the mean was calculated across epochs within each condition.

*V1 spatiotemporal cluster-based permutation testing.* To statistically assess V1 layer-dependent activity over time, spatiotemporal cluster-based permutation testing was conducted (two-tailed; 1000 permutations; cluster forming and cluster level *p* < 0.05; baseline interval = 19 TRs [∼28 seconds] pre-inducer and mock inducer onset, which is the maximum duration to the onset of the preceding trial inducer or mock inducer) focusing on the 0-20 TRs (∼32 seconds) post-inducer and mock inducer onset. The analysis followed a similar approach to the whole brain cluster-based permutation analyses (see *Whole brain spatiotemporal cluster-based permutation testing* section) with one key difference: in the layer-dependent analysis, spatial adjacency was defined based on neighboring cortical layers.

Analyses were performed across participants using their mean BOLD and VASO signals within each cortical layer of the V1 ROI. VASO layer signal timecourses were filtered using a zero-phase, zero-temporal shift low-pass filter to remove signal spikes observed across voxels and time, yet results were similar with and without filtering. To enhance spatial sensitivity in the cluster-based permutation testing, the layer-resolved BOLD and VASO signals were upsampled from 9 to 18 layers using linear interpolation, although comparable results were observed with and without upsampling. For visual interpretability, the sign of the VASO signal was flipped (i.e., multiplied by negative 1), as decreases in VASO signal correspond to increases in BOLD signal.

Two main tests were conducted separately for BOLD and VASO signals: (1) inducers *with* afterimages versus baseline and (2) mock inducers *with* mock afterimages versus baseline (VASO: Figure 5; BOLD: Supplementary Figure 15). We also conducted several supplementary analyses separately on the BOLD and VASO signals: (1) inducers + mock inducers only versus baseline (Supplementary Figure 16), (2) inducers only versus mock inducers only (Supplementary Figure 17), (3) inducers *with* afterimages versus mock inducers *with* mock afterimages, (4) inducers *with* afterimages versus inducers + mock inducers only, (5) mock inducers *with* mock afterimages versus inducers + mock inducers only (Supplementary Figures 18 and 19), and (6) eye blinks versus baseline (Supplementary Figure 20),

*Bootstrapping V1 spatiotemporal cluster-based permutation testing.* To assess the statistical robustness of the V1 layer-dependent activity identified by spatiotemporal cluster-based permutation testing, bootstrapping analysis was performed. A resampled dataset (resampling with replacement) among the analyzed participants was formed (N = 10) and spatiotemporal cluster-based permutation testing was evaluated on the resampled dataset. This process was repeated for 5000 iterations. Across each iteration, statistically significant layer-time clusters were tracked. A final statistical robustness or *consistency map* was generated by finding the proportion of iterations where the layer-time clusters were found statistically significant. The consistency maps (e.g., Figure 5G and H) are shown with contour lines surrounding layer-time clusters that were found statistically significant in > 75 percent of bootstrap iterations, highlighting the most statistically robust layer-time clusters.

*V1 cortical layer decoding.* Decoding analyses were conducted to complement the group-level assessment of the layer-specific signals linked with afterimages and mock afterimages (Figure 6). Previous research has demonstrated that decoding approaches can resolve layer-specific activity at the event level in layer-resolution fMRI (e.g. [73]). Observing common V1 cortical layer dynamics in both participant-level and event-level analyses would strengthen confidence in these results.

**Figure 6.**
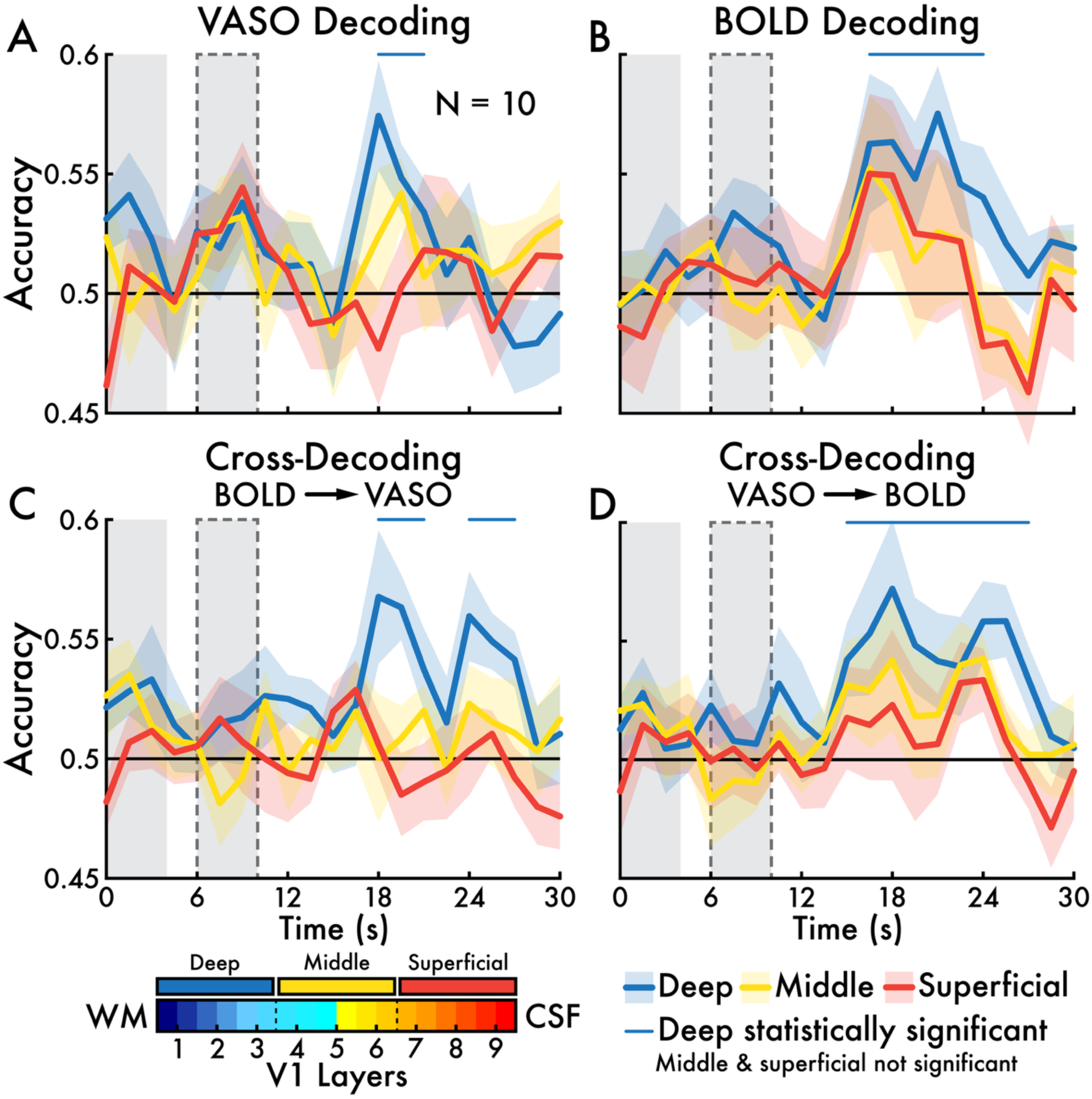
Afterimage versus mock afterimage VASO and BOLD V1 cortical layer decoding. Classifiers were trained to predict inducers *with* afterimages (*afterimage*) versus mock inducers *with* mock afterimages (*mock afterimage*) trials on a timepoint-by-timepoint basis within each participant (N = 10). Vascular space occupancy (VASO) and blood-oxygen-level-dependent (BOLD) classifier were trained independently. Classification was tested using “deep”, “middle”, and “superficial” regions of interest (ROI; VASO: **A**; BOLD: **B**; see *V1 fMRI* Analysis Methods section). The mean prediction accuracy rate across all participants at each time point following the inducer and mock inducer onset (0 seconds [s]) is shown colored with the tested layer ROI with standard error of the mean (shaded area). Modality cross-decoding classifier accuracy performance is shown for **(C)** training on BOLD and testing on VASO and (**D**) training on VASO and testing on BOLD. Statistically significant above chance (0.5) decoding performance is highlighted with a horizontal line in the color corresponding with the layer ROI (there was no significant middle or superficial decoding or cross-decoding). In all subplots, the gray bar between 0 and 4 s highlights the inducer and mock inducer period. The solid gray bar with a dotted outline between 6 and 10 s highlights the approximate afterimage and mock afterimage period. Cerebrospinal fluid (CSF); white matter (WM).

Within each participant, linear support vector machine (SVM) classifiers were trained to predict inducer *with* afterimage versus mock inducer *with* mock afterimage trials on a timepoint-by-timepoint basis. A total of 20 TRs (∼30 seconds) following the inducer and mock inducer onset were evaluated. At each timepoint, the classifier was repeated across 10 iterations of 10-fold cross-validation, with random train-test splits generated for each iteration to reduce dependence on a single partition. Balanced prediction accuracy was computed per iteration by averaging class-specific accuracies (i.e., true positives divided by the total trials for each class). Final decoding accuracy at each timepoint (TR) was defined as the mean balanced accuracy across all iterations. Statistically significant decoding across participants was assessed using temporal cluster-based permutation testing (one-tailed; 1000 permutations; cluster forming and cluster level *p* < 0.05), identifying timepoints with above chance prediction accuracy (0.5). Decoding was performed separately for the VASO and BOLD datasets (Figure 6A and B). As in the group-level analyses, the VASO signal was flipped (multiplied by –1) to match the BOLD signal directionality.

Modality cross-decoding analyses were also conducted to examine shared information between the VASO and BOLD signals. Classifiers were trained on one modality and tested on the other (Figure 6C: trained on BOLD and tested on VASO; Figure 6D: trained on VASO and tested on BOLD) using the same classification procedure described above, but without iterative cross-validation (training and testing datasets were independent). Cross-decoding performance was evaluated with temporal cluster-based permutation testing (one-tailed; 1000 permutations; cluster forming and cluster level *p* < 0.05).

To assess layer-specific contributions, VASO and BOLD decoding and cross-decoding classification was performed within layer-defined ROIs. Prior to ROI definition, the layer dimension was upsampled from 9 to 18 layers to correspond with the group-level analyses (see *V1 spatiotemporal cluster-based permutation testing* section). A 3-partition ROI set grouped the 18 individual layers into “deep” (layers 1-6), “middle” (layers 7-12), and “superficial” (layers 13-18) compartments. Each layer within each compartment contributed a single value towards classification at each timepoint. Alternative layer ROI schemes were tested, including performing decoding on each layer independently. However, we report decoding and cross-decoding on the 3-partition ROI set – deep, middle, and superficial – because it was most interpretable and suitable for statistical testing.

## Results

### Afterimage and image perceptual reporting task and post-task questionnaire

Participants reported on the sharpness, contrast, and duration of their afterimages during the afterimage and image perceptual reporting task (Supplementary Figure 1A) [39]. These perceptual details were used to create mock afterimages: animated, images designed to appear as each participant’s afterimage. Thus, the mock afterimage served as a perceptual control tailored to each participant. Across participants, the mock afterimages had a mean sharpness of 14.5 pixels (SD = 4.22; 0 = no blurring; 25 = maximum blurring), a mean maximum contrast of 0.20 (SD = 0.055; 0 = no contrast; 1 = full contrast), and a mean duration of 6.13 seconds (SD = 1.51; Supplementary Figure 1C).

Importantly, participants were not told that the mock afterimage was an on-screen image. However, 6 participants (18.75 percent) reported noticing that the mock afterimage was an on-screen image (e.g., 1 participant explaining that when they moved their eyes away from fixation, the mock afterimage remained in place on screen, whereas afterimages followed their gaze position). The remaining participants believed the mock afterimage was an illusory percept until they were debriefed at the end of their final study session. Overall, participants reported in the post-task questionnaire that the afterimage and mock afterimage shared similar visual perceptual attributes, although some participants reported differences (Table 1). Also, 5 participants described distortions and transformations in their afterimages, including the appearance of facial expressions, resemblance to familiar faces, and changes in shape (Table 2).

**Table 2.** Participant reports on afterimages appearing to distort or transform. Each report (x) reflects an individual participant’s response to the post-task questionnaire (see *Post-task questionnaire* Methods section).

| *Afterimage distortions and transformations* |  |
| --- | --- |
| × | Afterimage turned into a demon and crunched its face; one time the afterimage had red lips. |
| × | Afterimages morphed; the afterimage face looked 80 years old, while the inducer face appeared young; alien or ghostly appearing afterimages; afterimage eyebrows changed shape; afterimage resembled a familiar face or cartoon character. |
| × | Afterimages appeared like a skull. |
| × | The outline of the afterimage sometimes shifted its position. |
| × | Afterimage would disappear in sequential sections; the chin portion of the afterimage face disappearing before the eyes. |

### Afterimage and mock afterimage perception task

Participants completed a perception task in which they experienced illusory afterimages and perceptually similar, on-screen images or mock afterimages (see *Afterimage and mock afterimage perception task* Methods section). The perception task was administered with simultaneous pupillometry, eye tracking, and fMRI recordings. The following sections detail the results of these measurements in relation to task events, with a particular focus on afterimages and mock afterimages.

*Behavior.* During the whole brain fMRI study session, participants reliably reported afterimages following the inducers (mean perception rate = 0.79; SD = 0.25) but rarely reported afterimages after the mock inducers when no mock afterimage was physically presented (mean perception rate = 0.064; SD = 0.11; Figure 1D). This confirms that the mock inducers alone did not elicit afterimages. However, when mock afterimages were presented following the mock inducers, participants reported perceiving them at a high rate (mean perception rate = 0.95; SD = 0.075). Similar afterimage and mock afterimage perception rates were observed during the V1 fMRI study session (afterimage mean perception rate = 0.83; SD = 0.12; mock afterimage mean perception rate = 0.94; SD = 0.095; mock afterimage mean perception rate following mock inducers without mock afterimages = 0.05; SD = 0.07; Supplementary Figure 2).

During the whole brain fMRI study session, the duration of afterimages (mean = 3.66 seconds; SD = 1.25) and mock afterimages (mean = 4.12 seconds; SD = 1.55), and the onset latency of afterimages (mean = 1.64 seconds; SD = 0.74) and mock afterimages (mean = 1.81 seconds; SD = 0.33) were not statistically different (*p* > 0.05; Figure 1E). Similar durations and onset latencies were reported during the V1 fMRI study session (afterimage mean duration = 4.23 seconds; SD = 1.52; mock afterimage mean duration = 3.77 seconds; SD = 1.11; afterimage mean onset latency = 1.51 seconds; SD = 0.67; mock afterimage mean onset latency = 1.84 seconds; SD = 0.31; Supplementary Figure 2). Within participant variance of afterimage and mock afterimage duration and onset latency was statistically greater for afterimages versus mock afterimages in both the whole brain and V1 fMRI study sessions, except the onset latency was not significant for the V1 fMRI session (Supplementary Figure 3).

In addition, in the whole brain fMRI session, there was a statistically significant correlation across participants for the duration (*r* = 0.60; *p* = 0.0003) and onset latency (*r* = 0.41; *p* = 0.019) between afterimages and mock afterimages, suggesting shared temporal perceptual features within participants (Figure 1F). In the V1 fMRI session, a statistically significant correlation was found for duration (*r* = 0.62; *p* = 0.037) but not onset latency (*p* > 0.05; Supplementary Figure 2C).

For participants who completed both the whole brain fMRI and V1 fMRI study sessions (N = 12), afterimage temporal measures were significantly correlated between sessions for both onset latency (*r* = 0.93; *p* = 9.03×10^−6^) and duration (*r* = 0.72; *p* = 0.0089; Supplementary Figure 4A and B).

Similarly, mock afterimage temporal measures were significantly correlated between sessions for both onset latency (*r* = 0.78; *p* = 0.0026) and duration (*r* = 0.96; *p* = 6.85×10^−7^; Supplementary Figure 4C and D). The correlation in the mock afterimage onset latency and duration between study sessions is not surprising given that participants were presented with the same mock afterimage during both the whole brain and V1 fMRI sessions. In contrast, the correlation in afterimage onset latency and duration between study sessions suggests that afterimage perception may reflect a stable individual trait that persists over at least several days, corresponding to the interval between the fMRI study sessions.

*Eye behavior responses.* During the baseline interval prior to the inducer stimulus onset (–2-0 seconds), pupil size was significantly smaller for inducer trials *without* versus *with* afterimages (*p* = 0.0053; Supplementary Figure 5A). This finding is complemented by whole brain fMRI results that show brain state prior to the inducer onset distinguishes between inducer trials *with* and *without* afterimages (Supplementary Figure 9A and B; Supplementary Slides 4; see *Whole brain BOLD signals* section). A decrease in microsaccade fraction for inducer trials *without* versus *with* afterimages did not survive correction for multiple comparisons (*p* = 0.025). There were no significant differences in the baseline interval for mock inducer trials *with* versus *without* mock afterimages.

During the inducer and mock inducer presentation interval (0-4 seconds), pupil size and blink fraction exhibited distinct patterns of change. The mock inducer produced two troughs and peaks in pupil size, blink fraction, and microsaccade fraction corresponding with its flashing presentation (Supplementary Figure 5B and C). Notably, eye behaviors during the mock inducer presentation showed little to no difference between mock inducer trials *with* and *without* mock afterimages (Supplementary Figure 5B). In contrast, differences were observed during the inducer interval between inducer trials *with* and *without* afterimages (Supplementary Figure 5A). Specifically, pupil size was larger (*p* = 0.0013) and blink fraction was smaller (*p* = 0.0019) for inducers that led to an afterimage compared to those that did not. An increase in microsaccade rate also appeared for inducer trials *with* versus *without* afterimages and mock inducer trials *with* and *without* mock afterimages, but these differences did not survive correction for multiple comparisons (inducer trial: *p* = 0.013; mock inducer trial: 0.0047). This result is linked with distinct inducer related whole brain fMRI responses for inducer trials *with* versus *without* afterimages (Supplementary Slides 4; see *Whole brain BOLD signals* section). Together, these findings suggest that physiological state both before and during the inducer presentation is related to whether an afterimage is perceived.

During the afterimage and mock afterimage interval (7-9 seconds), trials with afterimages and mock afterimages exhibited a decrease in blink fraction (inducer trials *with* versus *without* afterimages: *p* = 0.0042; mock inducer trials *with* versus *without* mock afterimages: *p* = 8×10^−8^; Supplementary Figure 5A and B). An increase in microsaccade rate for mock inducer trials *with* versus *without* mock afterimages did not survive correction for multiple comparisons (*p* = 0.015).

During the post-afterimage interval (11-13 seconds), inducer trials *with* afterimages had a smaller pupil size (*p* = 7×10^−4^), increased blink fraction (*p* = 9×10^−6^), and decreased microsaccade fraction (*p* = 4×10^−4^) relative to inducer trials *without* afterimages (Supplementary Figure 5A). During the post-mock afterimage interval, mock inducer trials *with* mock afterimages had an increased blink fraction (*p* = 7×10^−5^) relative to mock inducer trials *without* mock afterimages (Supplementary Figure 5B). Also, there was an increase in microsaccade fraction for mock inducer trials *with* versus *without* mock afterimages, but this difference did not survive correction for multiple comparisons (*p* = 0.0081).

Comparison between the inducers *with* afterimages and mock inducers *with* mock afterimages conditions revealed differences in pupil size for the inducer and mock inducer (*p* = 2×10^−9^) and post-afterimage and mock afterimage intervals (*p* = 4×10^−4^; Supplementary Figure 5C). Also, microsaccade fraction showed a difference between conditions for the inducer and mock inducer interval but did not survive correction for multiple comparisons (*p* = 0.006; Supplementary Figure 5C).

*Whole brain BOLD event-related signals.* Whole brain fMRI analyses revealed that afterimages and mock afterimages recruited widespread subcortical and cortical BOLD signal increases, independent of inducer and mock inducer driven responses, including in visual (e.g., LGN, V1, FG and LOC), salience (e.g., AC and In), and motor network regions (e.g., supplementary motor area [SMA], basal ganglia, and left M1; Figure 2A, B, C, and D; Supplementary Slides 1 and 2). There were also increases in visual attention and decision making frontal and parietal regions, including the superior parietal lobule, inferior parietal lobule, superior frontal gyrus, and inferior frontal gyrus. The engagement of these areas could be explained by task demands (e.g., responding to the onset and offset of afterimages and mock afterimages). Supporting this interpretation, these possible task-related regions, including motor areas did not differ between afterimage and mock afterimage conditions (Figure 2E and F).

When unilateral visual field analyses were performed (i.e., independently analyzing the left versus right visual field stimulus presentation trials), similar areas of activation were observed as in the bilateral analyses (Supplementary Figure 10). However, a key difference was that visual sensory regions (e.g., V1, FG, and LOC) showed contralateral BOLD signal increases relative to the visual field presentation location, while the ipsilateral areas generally exhibited no change or signal decreases. Meanwhile, for both the afterimage and mock afterimage conditions, higher order association, salience, and motor network regions (e.g., left M1) were retinotopically invariant.

Contrasting the afterimage and mock afterimage BOLD signals revealed broad decreases for afterimages relative to mock afterimages (Figure 2E and F; Supplementary Slides 3). In short, while afterimages and mock afterimages shared similar areas of activation, the magnitude of the signal was significantly reduced for afterimages relative to mock afterimages across many of these overlapping brain networks. These decreases were predominately present in visual and frontal cortices, particularly middle frontal gyrus. However, two notable salience network regions revealed *greater* BOLD signals for afterimages versus mock afterimages: AC and In.

Building confidence in the unilateral whole brain BOLD results, similar activation patterns were observed when applying a general linear model analysis that accounted for head movement and blinking (Supplementary Figure 7). Common findings between the analysis methods include widespread cortical and subcortical activation for afterimages and mock afterimages and reduced signals for afterimages versus mock afterimages, except for increases in salience network regions.

Finally, differences in afterimage and mock afterimage BOLD signals cannot be explained by activity from inducers and mock inducers. This was directly tested by contrasting BOLD responses to inducers and mock inducers. This comparison revealed only small differences at the peak inducer response time (6 seconds post-inducer and mock inducer onset) and the peak afterimage and mock afterimage response time (14 seconds post-inducer and mock inducer onset; Supplementary Figure 8).

*Whole brain functional networks.* Functional clustering on afterimage and mock afterimage BOLD dynamics was performed to organize whole brain activity patterns and summarize signal trends (see *K-means clustering* Methods section). K-means clustering (k = 3) resulted in clusters that approximately corresponded with established functional brain networks (Figure 3A; Supplementary Figures 11A and 12A and B): (1) sensory and salience (SSN), (2) task-positive (TPN), and (3) task-negative (TNN) networks. The network names were assigned based on the known functional roles of the major brain regions grouped within each network. Among other regions, the SSN includes V1, extrastriate cortex, FG, lateral occipital cortex, superior frontal gyrus, AC, and In; the TPN includes the medial frontal gyrus (pre-SMA), basal ganglia, and right superior cerebellum; and the TNN includes the angular gyrus, ventral medial prefrontal cortex, precuneus, and posterior cingulate cortex (PCC) – all areas affiliated with the default mode network. Importantly, these cluster labels are approximations, as some regions do not strictly align with canonical network definitions (e.g., the left M1 was grouped within the SSN, while right M1 was grouped within TNN, corresponding to right-handed button presses made during the perception task).

The average BOLD timecourses for the SSN, TPN, and TNN clarified their functional identity based on temporal dynamics relative to task events (Figure 3B; Supplementary Figure 12C and D). The SSN exhibited an early BOLD increase, peaking ∼11 seconds post-inducer and mock inducer onset. The TNN showed an inverse pattern to the SSN, with an initial decrease reaching a nadir at ∼12 seconds, followed by an increase between 15 and 25 seconds. Finally, the TPN had a late BOLD increase, peaking at ∼15 seconds. The SSN response timing suggests that these areas are sensitive to afterimages and the task stimuli (inducer, mock inducer, and mock afterimage). In contrast, the late TPN response suggests sensitivity to the task behavior (i.e., participants indicating the onset and offset of perceived afterimages and mock afterimages).

This interpretation is supported by additional analyses that used each network as a ROI to visualize BOLD signal trends across task conditions and within each network. In the SSN, the afterimage, mock afterimage, and inducers + mock inducers only conditions all shared similar onset latencies but diverged in their peak response time: the inducers + mock inducers only condition exhibited an early peak response ∼6 seconds post-inducer and mock inducer onset, while the afterimage and mock afterimage conditions showed later peaks between 10-15 seconds (Figure 3C). Also, the response magnitude in the SSN was reduced for afterimages compared to mock afterimages, consistent with the observed whole brain BOLD signal magnitudes (Figure 2E and F; Supplementary Slides 3).

In the TPN, afterimage and mock afterimage conditions showed similar BOLD timecourses, while the inducers + mock inducers only condition did not elicit a significant response (Figure 3C). This result aligns with the behavior during the perception task: button presses were made for the afterimage and mock afterimage conditions only, while no response was made for the inducers + mock inducers only trials. These results further highlight that the TPN was specifically responsive to task behavior.

In the TNN, all conditions produced a similar pattern of change: a decrease in BOLD signal, reaching a nadir between 10-13 seconds post-inducer and mock inducer onset (Figure 3C). The presence of this response for the inducers + mock inducers only condition suggests it is not solely driven by task demands, rather it may reflect network inhibitions or switches [74]. Notably, a late BOLD signal rebound was observed in the TNN, potentially reflecting recovery following visual stimulation, afterimage perception, and task-engagement.

To assess the robustness of the SSN, TPN, and TNN networks, k-means clustering was also tested for larger k-values of 4 and 5. Cluster silhouette values deteriorated for k-values greater than 4 (Supplementary Figure 11C). However, the SSN was largely stable across k-values, apparent from consistent network brain maps and BOLD timecourses (Supplementary Figure 11A and B). Meanwhile, with larger k-values, the TPN and TNN were fragmented, highlighted by distinct, temporally shifted cluster BOLD timecourses.

*Sensory and salience network regions of interests.* Within network analyses focused on the SSN because it was the most responsive brain network to both afterimages and mock afterimages (Figure 3C). We selected 6 ROIs clustered within SSN (Figure 4A; see *Regions of interest* section): (1) LGN, (2) V1, (3) FG, (4) AC, (5), In, and (6) left M1 (L-M1). In L-M1, the BOLD signal showed the expected response: late increases in activity of equal magnitude for afterimages and mock afterimages, with no response for the inducers + mock inducers only condition (Figure 4G). There were two distinct phases in the afterimage and mock afterimage L-M1 timecourses, likely corresponding to the two button presses made by participants indicating the onset and offset of perceived afterimages and mock afterimages.

The visual sensory ROIs (LGN, V1, and FG; Figure 4B, C, and D) exhibited robust early responses across all conditions, with an onset latency of ∼4 seconds after the inducer or mock inducer onset. However, only the afterimage and mock afterimage conditions showed a second distinct response, while the inducers + mock inducers only condition returned to baseline after an initial peak. In the LGN and V1, this second, afterimage and mock afterimage-linked response appeared as a distinct peak at ∼15-17 seconds. Notably, in V1, the afterimage and mock afterimage peaks were reduced in magnitude compared to the earlier inducer and mock inducer-related peaks. In FG, instead of two peaks, a prolonged, unimodal response was observed, peaking between ∼7-11 seconds. Consistent with the whole brain results (Figure 2E and F), the afterimage response was weaker in all visual sensory ROIs relative to the mock afterimage.

In the salience network ROIs (AC and In), all trial conditions revealed increases in BOLD activity. However, the peak time differed: the inducers + mock inducers only condition peaked ∼9-12 seconds post-inducer and mock inducer onset, while the afterimage and mock afterimage conditions peaked between ∼13-15 seconds (Figure 4E and F). Furthermore, as observed in the whole brain results, afterimages evoked a stronger BOLD response than the mock afterimages in AC and In.

In summary, the ROI BOLD timecourses complement and expand on the whole brain BOLD activation maps by showing the full temporal response in areas of known activity. For LGN, V1, and L-M1, bimodal BOLD responses were present, likely reflecting both stimulus processing and task-related motor activity. For FG, AC, and In, unimodal BOLD responses were present, possibly reflecting sustained or integrative processing (e.g., overlapping contributions from both the inducer and afterimage, and the mock inducer and mock afterimage).

*Whole brain states conducive for afterimage perception.* Afterimages were often but not always perceived following the inducer (Figure 1D; Supplementary Figure 2A). Some participants nearly always perceived an afterimage following the inducer, while approximately a third of participants reported seeing no afterimage in more than ∼25 percent of inducer presentations. We explored if brain activity prior to and during the inducer presentation played a role in whether an afterimage was subsequently perceived.

Whole brain BOLD signal analyses tested differences between inducer trials *with* and *without* afterimages in a subgroup of participants (N = 18). These participants were selected because they both perceived and did not perceive afterimages following the inducer, with a minimum of 5 epochs of each trial type. Notably, during the interval responsive to the period immediately preceding and during the inducer presentation, BOLD decreases were observed, particularly in visual cortex and the salience network, and BOLD increases in midbrain for inducers with afterimages relative to inducers without subsequent afterimages (Supplementary Figure 9A and B; Supplementary Slides 4). Meanwhile, mock inducer trials *with* versus *without* mock afterimages in the same participants revealed little to no difference at the same time points (Supplementary Figure 9C and D; Supplementary Slides 5), evidencing that baseline brain states influence the perception of afterimages but not mock afterimages.

These findings are linked with the differences in pupil size and blink fraction (and possibly microsaccades) for inducer trials *with* and *without* afterimages preceding (pupil) and during (pupil and blink) the inducer presentation (Supplementary Figure 5A; see *Eye behavior responses* section). Importantly, these systematic differences in eye behaviors were largely absent for mock inducers *with* versus *without* mock afterimages, except for possibly microsaccades during the mock inducer period, corresponding with the absence of BOLD signal differences (Supplementary Figure 5B). This distinct pattern of eye behavior for inducer trials *with* versus *without* afterimages could contribute to the baseline brain states. However, the precise interaction among eye behavior, brain state, and afterimage perception is unknown.

*V1 cortical layer-dependent activity.* The whole brain BOLD results along with previous fMRI studies (e.g., [34]), support that afterimages activate V1. However, common activation of V1 does not necessarily imply that afterimages and images share neural information flow. Feedforward and feedback neural signaling are associated with distinct layer-specific activity patterns, detectable using high-spatial-resolution fMRI capable of resolving meso-scale activity [49]. To investigate neural signaling patterns in V1 for afterimages and mock afterimages, we collected high-spatial-resolution BOLD and VASO fMRI data in V1 while participants completed the perception task (*V1 fMRI sequence* Methods sections).

First, we confirmed that layer-resolution BOLD signals in V1 produced responses across afterimage, mock afterimage, and inducers + mock inducers only conditions that closely matched the timecourses observed in the whole brain BOLD V1 ROI analysis (Figure 4C versus Figure 5B). This replication suggests consistency between the whole brain and V1 datasets.

Subsequently, we analyzed event-related, layer-dependent VASO and BOLD responses in V1. VASO offers higher spatial specificity than BOLD, especially in layer analyses, due to its reduced sensitivity to large vessels positioned along the cortical surface [38]. Therefore, participant-level VASO results are emphasized (Figure 5; Supplementary Figures 16A, C, and E, 17A, C, and E, 18, and 20A, C, and E), though layer-dependent BOLD signals are also reported (Supplementary Figures 15, 16B, D, and F, 17B, D, and F, 19, and 20B, D, and F). However, layer-specific afterimage versus mock afterimage event-level decoding analyses found correspondence between VASO and BOLD, thereby both modalities are highlighted in these findings (Figure 6).

During the inducer and mock inducer period, VASO increases were synchronized across V1 cortical layers, with the largest signal change located in middle and superficial layers, peaking ∼4-6 seconds after onset (Figure 5C, D, E, and F; Supplementary Figure 16A and C). A similar pattern was observed in BOLD (Supplementary Figures 15A, B, C, and D, and 16B and D). Contrasting the inducers and mock inducers revealed no significant difference in VASO, while there was *more* BOLD activity in middle and superficial layers for inducers relative to mock inducers approximately between ∼9-11 seconds (Supplementary Figure 17). These results affirm that the afterimage and mock afterimage period (∼11-15 seconds) does not include inducer and mock inducers signal differences that could systemically bias the afterimage and mock afterimage evoked responses. Consistency maps support the robustness of the inducer and mock inducer VASO (Figure 5G and H; Supplementary Figures 16E and 17E) and BOLD (Supplementary Figure 15E and F, 16F, and 17F) responses.

In correspondence, there was no significant VASO or BOLD decoding between inducer *with* afterimage and mock inducer *with* mock afterimage trials during the timepoints responsive to the inducer and mock inducer events (0-12 seconds; Figure 6A, B, C, and D). This suggests common layer-specific activity between the inducer and mock inducer.

During the afterimage and mock afterimage period, cortical layer differences were observed in VASO. Participant averaged VASO percent change V1 cortical layer timecourses reveal a *deep* afterimage and *superficial* mock afterimage layer responses (∼12-15 seconds; Figure 5C and D). This layer-specific dynamic between conditions was further supported by layer-time statistical testing that highlights that afterimages elicited *middle* and *deep* layer VASO activity, while mock afterimages drove *middle* and *superficial* layer VASO activity (Figure 5E, F, G, and H). Contrasting the VASO responses for afterimages and mock afterimages reinforces the deep layer response for afterimages (Supplementary Figure 18A). Furthermore, comparing the afterimages and mock afterimages independently with the inducers + mock inducers only condition highlights the unique deep and superficial layer responses for afterimages and mock afterimages, respectively (Supplementary Figure 18B and C). Although, none of these contrast analyses revealed statistically significant layer-time clusters (Supplementary Figure 18D, E, F, G, H, and I).

Meanwhile, the participant averaged BOLD percent change V1 cortical layer timecourses showed a consistent superficial layer response bias for both afterimages and mock afterimages (Supplementary Figure 15A and B). Layer-time statistical testing showed only significant superficial layer BOLD activity during the afterimage and mock afterimage responsive period for afterimages, while all layers were shown active for mock afterimages (Supplementary Figure 15C and D). However, the consistency maps suggest that the BOLD layer-time statistical findings lack statistical robustness (Supplementary Figure 15E and F).

The afterimage specific deep V1 cortical layer response was further supported by significant VASO and BOLD decoding between inducer *with* afterimage and mock inducer *with* mock afterimage trials only during the timepoints most responsive to afterimages and mock afterimages (∼15-25 seconds; Figure 6A and B). There was no significant decoding in the middle or superficial layers. Also, highlighting the correspondence between the VASO and BOLD decoding findings, signal modality cross-decoding was statistically significant in only deep layers during timepoint responsive to afterimages and mock afterimages (Figure 6C and D). The lack of correspondence between the VASO and BOLD participant-level, voxelwise results may be related to VASO showing more spatial sensitivity [75]. However, decoding methods may normalize against these biases, explaining why the VASO and BOLD event-level results were concordant.

Finally, the known relationship between afterimages and blinking, and our findings that blink rate was reduced during the inducer period for trials *with* versus *without* afterimages, encouraged examining the V1 cortical layer responses for blinks. This analysis could indicate how blink related responses may account for systematic differences in the V1 fMRI layer findings. Blinks resulted in a consistent pattern of change in VASO and BOLD V1 cortical layer activity: an initial increase across all layers followed by a prolonged decrease across all layers, with the superficial layers showing the largest dynamic range in signal change (Supplementary Figure 20). These blink evoked responses are not suggestive that systematic differences in blink rate between afterimage and mock afterimage trials might explain the main layer-dependent responses reported.

## Discussion

The precise neural mechanisms of afterimages are unknown, and the involvement of brain processes in afterimage perception is debated. To advance knowledge on the neurophysiological basis of afterimages, we examined the human whole brain and V1 cortical layer fMRI responses, along with eye behaviors during afterimage perception and while viewing “mock afterimages” – animated, on-screen images that were tailored to resemble each participant’s afterimage.

Our results are summarized by 4 main findings: (1) afterimages engaged widespread cortical and subcortical activity, particularly across visual sensory regions (Figure 2 and 3); (2) the magnitude of the afterimage fMRI signal was *weaker*, especially in visual sensory and lateral frontal regions compared to perceptually similar images (Figure 2, 3, and 4); (3) the magnitude of the afterimage fMRI signal was *greater* in salience network regions (AC and In) compared to perceptually similar images (Figure 2 and 4); and (4) afterimages involved activity in *deep* and *middle* cortical layers of V1, whereas perceptually similar images showed activity in *middle* and *superficial* layers, suggesting that afterimage and image perception rely on partly distinct neural information flow dynamics in V1 (Figure 5 and 6).

Additional observations include that afterimages maximally activated the contralateral visual cortex relative to the visual field in which the afterimage appeared (Supplementary Figure 10). This pattern of response mirrors those of images and shows that afterimage neural signaling abide by the contralateral circuitry of the primary visual pathway.

Furthermore, we found differences in baseline fMRI activity between trials in which an afterimage was perceived versus not perceived, including fMRI signal *decreases* in visual cortex and *increases* in salience and arousal network regions (AC, In, and midbrain; Supplementary Figure 10; Supplementary Slides 4). These findings support that preceding brain state contributes to whether an afterimage is subsequently perceived. Likewise, previous behavioral findings show that attentional and conscious states prior and during afterimage perception influences their conscious experience [76–78]. Further, the influence of neurophysiological state, which can be dynamic on a moment-by-moment basis or sustained state differences (e.g., as a result of individual trait or disease), may help explain why afterimages are found to be more perceptually heterogenous within and across individuals compared to suprathreshold images, as we report in the current study (Figure 1; Supplementary Figure 1, 2, and 3; also see [39]). In support, we found that the afterimage onset latency and duration were strongly correlated across individuals between the whole brain and V1 fMRI study sessions, which were completed days apart (Supplementary Figure 4A and B).

Linked to the baseline brain activity findings is that eye behaviors before (pupil size) and during (pupil size and blinking) the inducer presentation interval – the stimulus administered to evoke afterimages – differed between trials when an afterimage was perceived versus not perceived. However, the relationship between baseline brain states and eye behaviors associated with afterimage perception remains unresolved. We also found that blinking and microsaccades responded similarly to afterimages and perceptually similar images, although this common response may reflect task demands (Supplementary Figure 5).

An unexpected outcome from this investigation was that some participants reported that their afterimages would occasionally transform and distort (e.g., appearing as a recognizable face or making facial expressions; Table 2). This could be related to top-down effects that alter the appearance of afterimages. In support, previous behavioral research finds that afterimages are sensitive to top-down modification, including by priming and perceptual priors [42, 79]. These results correspond with the afterimage-specific deep V1 cortical layer findings (Figure 5 and 6), which could suggest higher-order cortical, top-down inputs that modify afterimage perception.

### Do afterimage and image perception share brain mechanisms?

Afterimages engaged similar brain regions as perceptually similar images, including V1, FG, and LOC. However, our results also indicate that the neural processes underlying afterimages and images differ. A major difference was reduced fMRI signal magnitude for afterimages compared to images across the entire brain, especially in visual network regions (Figures 2, 3, and 4). This finding aligns with a previous study reporting better fMRI-based classification performance for images versus afterimages, which may be explained by weaker afterimage signals [35]. Importantly, the afterimage-linked fMRI signals were weaker, despite the images closely resembling afterimages (Table 1). Notably, many participants were convinced that these images were also illusory afterimages.

Another difference was in the pattern of V1 cortical layer activity. Although both afterimages and images activated V1, layer-dependent analyses revealed that afterimages selectively engaged deep and middle layers, whereas images engaged middle and superficial layers of V1 (Figure 5). This result was further supported by successful VASO and BOLD decoding and modality cross-decoding of afterimage and mock afterimage trials in deep but not middle and superficial layers (Figure 6).

The middle layer of V1 receives feedforward input, while superficial and deep layers are primarily associated with feedback signaling [80, 81]. In addition, distinct neural activity dynamics in superficial versus deep layers of V1 suggest different types of feedback [82, 83]. One hypothesis is that superficial layers involve externally generated feedback (e.g., during visual perception), while deep layers involve internally generated feedback (e.g., during visual imagery) [84]. This framework is supported by findings that visual perceptual illusions engage superficial cortical layers, while visual imagery recruit deep layers [85, 86]. Therefore, neural information flow may be unique among different categories of conscious perception (e.g., different illusions involving distinct neural information flow properties).

Correspondingly, the observed V1 cortical layer activity suggests that afterimages involve internally generated feedback at the earliest stage of visual cortical processing. Thereby, the neural mechanisms of afterimages in V1 may overlap with visual imagery and other self-generated percepts (e.g., hallucinations and dreams). This conclusion is corroborated by behavioral evidence that the vividness of afterimages and visual imagery are related [39, 87, 88]. Relatedly, it has been reported that visual imagery can induce afterimages [13, 16].

Another framework for interpreting these V1 cortical layer results is *predictive coding theory,* which proposes that feedforward and feedback pathways convey distinct information about how well the brain’s models of the sensory world align with sensory input. When these internal models are *congruent* with input, feedback signaling dominates in deep cortical layers, while when *incongruent*, feedforward signaling is more prominent in superficial layers [89]. For example, percepts elicited by unexpected stimuli generate a large prediction error (incongruent) because top-down predictions are weak or absent, thereby superficial activations dominate. By contrast, percepts from self-generated sources, such as imagery, are associated with a strong prior and therefore a top-down dominate regime. Consistent with this framework, mock afterimages show superficial responses, while afterimages activated deep V1 cortical layers. Under predictive coding theory, this pattern suggests that afterimage perception depends on a strong prior, linking afterimages with self-generated percepts.

Finally, the differences observed between afterimages and images, such as in frontal cortical and salience areas, including AC, In, and middle frontal gyrus may reflect *conflict* and *perceptual reality monitoring*: neural processes that distinguish real from illusory or self-generated conscious experiences (e.g., seeing versus imagining) [90–93]. Our findings may partly reflect neural mechanisms that detect signal features that conflict with ongoing visual input and distinguish illusory or internally generated visual percepts from those elicited by physically present images (i.e., detecting mismatch between sensory input and brain models).

While participants were not asked to explicitly differentiate between afterimages and images, and most participants were convinced that the perceptually similar images were illusory, it still may be that participants engaged in covert monitoring of the percept identity or subliminal neural processes that distinguished afterimages from mock afterimages. These monitoring mechanisms are hypothesized to involve feedback processing, thus another framework for interpreting the deep V1 cortical layer activity observed for afterimages and other self-generated percepts (e.g., filling-in illusions; [94]).

### Alternative explanations

We attribute our whole brain and V1 cortical layer findings primarily to the brain mechanisms underlying afterimage and image conscious perception, and possibly to the contribution of conflict and perceptual reality monitoring processes. However, alternative explanations for the main experimental findings should also be considered, particularly in light of several limitations of this study.

First, despite our attempt to perceptually match images with afterimages, perceptual differences between afterimages and mock afterimages remained for a subset of participants (Table 1). In addition, afterimages showed greater temporal variability than mock afterimages in both their perceived onset following the inducer and duration. Thereby, afterimage and image brain signals may differ partly because of their distinct perceptual characteristics.

Nonetheless, we argue that any perceptual differences between afterimages and mock afterimages were likely small, especially compared to studies that examine distinct categories of conscious perception (e.g., image perception versus imagery). Also, it is notable that although perceptual differences between the inducer and mock inducer are likely *greater* than between the afterimage and mock afterimage, whole brain and V1 layer responses were *more* similar between inducers and mock inducers than between afterimages and mock afterimages (e.g., Figure 2E and F versus Supplementary Figure 8E and F). Furthermore, no significant decoding was observed in V1 during the time interval responsive to the inducer and mock inducer, suggesting shared layer-specific activity despite differences in their visual features (Figure 6). Therefore, perceptual differences between afterimages and mock afterimages likely have limited influence on our findings, which instead reflect distinct neural signal characteristics and processing patterns.

Second, the inducer and mock inducer introduce brain responses that overlap with afterimage and image-evoked neural activity. This challenge is further exacerbated by the delayed hemodynamic response measured with fMRI. Although fMRI cannot fully dissociate temporally adjacent, time-locked events, comparing different trial conditions (e.g., inducers *with* and *without* afterimages) allows us to isolate signal variations that specifically emerge when an afterimage is perceived. Also, we found that inducers and mock inducers presented without afterimages or mock afterimages, respectively, produced similar responses (Supplementary Figures 8 and 17), and there was no statistically significant V1 decoding of inducers versus mock inducers (Figure 6). Together, these results mitigate concerns that differences in brain activity between afterimages and mock afterimages are driven by the inducer and mock inducer.

Finally, the button presses participants made to indicate the onset and offset of afterimages and images make it difficult to disentangle perceptual versus report-dependent neural activity. Furthermore, the timing of these responses may have systematically differed between the afterimage and image conditions (e.g., due to distinct internal thresholds for reporting perceived onset and offset). As a result, the relationship between behavioral responses and afterimages may differ from between behavioral responses and images, potentially contributing to condition-specific brain activity, even outside of motor planning and execution brain regions. To reduce reporting confounds, future studies should explore covert markers of afterimage perception in the absence of overt report [43]. Supporting this possibility, we found that pupil, blink, and microsaccade responses differed between inducers *with* versus *without* afterimages (Supplementary Figure 5). This suggests that it may be possible to predict when afterimages are perceived on a trial-by-trial basis based on eye behavior, as has been previously achieved for images [65].

### Are afterimages retinal or central?

The neurophysiological origins of afterimages are often framed as arising from either retinal *or* central neural processes [3]. This framing posits a false dichotomy. Much like image perception, typical afterimages involve both retinal *and* central neural mechanisms. Supporting this integrated perspective, the current results revealed widespread subcortical and cortical activity linked with afterimages, and baseline brain states conducive for afterimage perception. These afterimage-linked brain signals may originate in the retina but undergo post-retinal signal modulation that shapes the afterimage experience.

An additional nuance is that there are different categories of afterimages, such as positive and negative afterimages, and less explored subtypes, including conditioned afterimages or imagery-induced afterimages [13–16, 21]. Each category of afterimage may involve a unique balance of retinal and central contributions. For example, if imagery-induced afterimages are taken at face value, they exemplify pure cortically generated afterimages – afterimages in the absence of any retinal input. Meanwhile, afterimages resulting from retinal photoreceptor bleaching (e.g., after viewing a bright light) may be best characterized by perseverating retinal activity that drives a strong feedforward input. In short, the balance between retinal and central contributions in afterimage perception may be distinct according to afterimage type, just as different categories of illusions involve distinct neural mechanisms.

## Conclusions

Afterimages are a long-standing source of popular and scientific interest, yet there are few studies that directly investigate the neural processes involved in afterimage perception. In this experiment, we recorded human whole brain and V1 cortical layer fMRI signals and eye behaviors linked with afterimages and perceptually similar images. Our results show that afterimage and image perception involve overlapping, widespread brain activity. Yet, the afterimage signals were also distinct from those of images, including reduced signal magnitude across the brain, except in salience network regions where the fMRI signal was stronger for afterimages. We also observed that baseline brain activity differed whether an afterimage was subsequently perceived or not perceived, a finding linked to baseline differences in pupil size and blinking. Furthermore, afterimages selectively engaged middle and deep cortical layers of V1, while images recruited middle and superficial layers, suggesting distinct patterns of neural signaling. Correspondingly, decoding between afterimage and mock afterimage trials was only successful in deep V1 cortical layers. These neuroimaging findings help explain behavioral evidence that link afterimages with visual imagery and reveal that afterimages are influenced by neurophysiological state and top-down effects. While typical afterimages likely originate from retinal activity, these results provide compelling evidence for the role of central neural processing in afterimage perception. This motivates future investigation into the brain mechanisms of afterimages, with implications for advancing the broader goal of elucidating the neural mechanisms of visual consciousness.

## Code and Data Availability

Data and code will be available upon publication.

## Author Contributions

SIK was involved in conceptualization, methodology, software, validation, formal analysis, investigation, data curation, writing – original draft, visualization, supervision, and project administration; BA was involved in software, validation, formal analysis, writing – original draft, and visualization; MH was involved in software, formal analysis, investigation, and writing – review & editing; ATM was involved in conceptualization, methodology, investigation, and writing – review & editing; LH was involved in methodology, software, formal analysis, investigation, and writing – review & editing; PAT was involved in software, formal analysis, writing – review & editing, and visualization; JG-C was involved in software, formal analysis, writing – review & editing, and supervision; DAH was involved in conceptualization, methodology, formal analysis, writing – review & editing, and supervision; PAB was involved in conceptualization, methodology, writing – review & editing, supervision, project administration, and funding acquisition.

## Supporting information

Supplementary Slides 1

Supplementary Slides 2

Supplementary Slides 3

Supplementary Slides 4

Supplementary Slides 5

## Acknowledgements

This research was made possible by the support of the National Institute of Mental Health Intramural Research Program (ZIAMH002783, ZICMH002884, ZICMH002888). The study was completed in compliance with the National Institutes of Health Clinical Center protocol ID 93-M-0170 (ClinicalTrials.gov ID: NCT00001360). This work utilized the computational resources of the NIH HPC Biowulf cluster (https://hpc.nih.gov). Thank you to Marco Barilari, Gang Chen, Josh Faskowitz, and Francisco Pereira for helpful conversations regarding the analysis of the fMRI data. Thank you to Chung Kan for training and guidance in MRI acquisition. Thank you Rüdiger Stirnberg for sharing the 3D-EPI sequence used in the layer-fMRI VASO experiments. Thank you to the OP4 Behavioral Health Clinic staff for their support with patient admission and physical exams. This research was supported by the Intramural Research Program of the National Institutes of Health (NIH). The contributions of the NIH authors were made as part of their official duties as NIH federal employees, are in compliance with agency policy requirements, and are considered Works of the United States Government. However, the findings and conclusions presented in this paper are those of the authors and do not necessarily reflect the views of the NIH or the U.S. Department of Health and Human Services.

---

**Supplementary Movie 1.** *Afterimage trial*. An inducer stimulus appears (4 seconds) on the right side of the screen followed by a blank screen (Figure 1A). Participants often reported a negative afterimage appearing in the same location that the inducer was presented while maintaining central fixation (Figure 1D; Supplementary Figure 2A).

**Supplementary Movie 2.** *Mock afterimage trial*. A mock inducer stimulus appears (4 seconds) on the right side of the screen followed by a mock afterimage (∼4 seconds; Figure 1B). Each participant was shown an individualized mock afterimage that were perceptually similar with their afterimage perception following the inducer (Supplementary Figure 1D).

**Supplementary Movie 3.** *Mock inducer without a mock afterimage trial*. A mock inducer stimulus appears (4 seconds) on the right side of the screen followed by a blank screen. Participants often reported that the mock inducer did not produce an afterimage (Figure 1D; Supplementary Figure 2A).

**Supplementary Slides 1.** *Afterimage whole brain volume BOLD activation maps*. Statistically significant *t*-values are highlighted with a black outline, while sub-threshold regions are shown with transparency at time point 0 to 25 seconds (s) after the inducer onset for inducers *with* afterimages versus inducers + mock inducers only conditions.

**Supplementary Slides 2.** *Mock afterimage whole brain volume BOLD activation maps*. Statistically significant *t*-values are highlighted with a black outline, while sub-threshold regions are shown with transparency at time point 0 to 25 seconds (s) after the mock inducer onset for mock inducers *with* mock afterimages versus inducers + mock inducers only conditions.

**Supplementary Slides 3.** *Afterimage versus mock afterimage whole brain volume BOLD activation maps*. Statistically significant *t*-values are highlighted with a black outline, while sub-threshold regions are shown with transparency at time point 0 to 25 seconds (s) after the inducer and mock inducer onset for inducers *with* afterimages versus mock inducers *with* mock afterimages conditions.

**Supplementary Slides 4.** *Inducers with versus without afterimages whole brain volume BOLD activation maps.* Statistically significant *t*-values are highlighted with a black outline, while sub-threshold regions are shown with transparency at time point –5 before to 25 seconds (s) after the inducer onset for inducers *with* afterimages versus inducers *without* afterimages conditions.

**Supplementary Slides 5.** *Mock inducers with versus without mock afterimages whole brain volume BOLD activation maps.* Statistically significant *t*-values are highlighted with a black outline, while sub-threshold regions are shown with transparency at time point –5 before to 25 seconds (s) after the inducer onset for mock inducers *with* mock afterimages versus mock inducers *without* mock afterimages conditions.

**Supplementary Figure 1.**
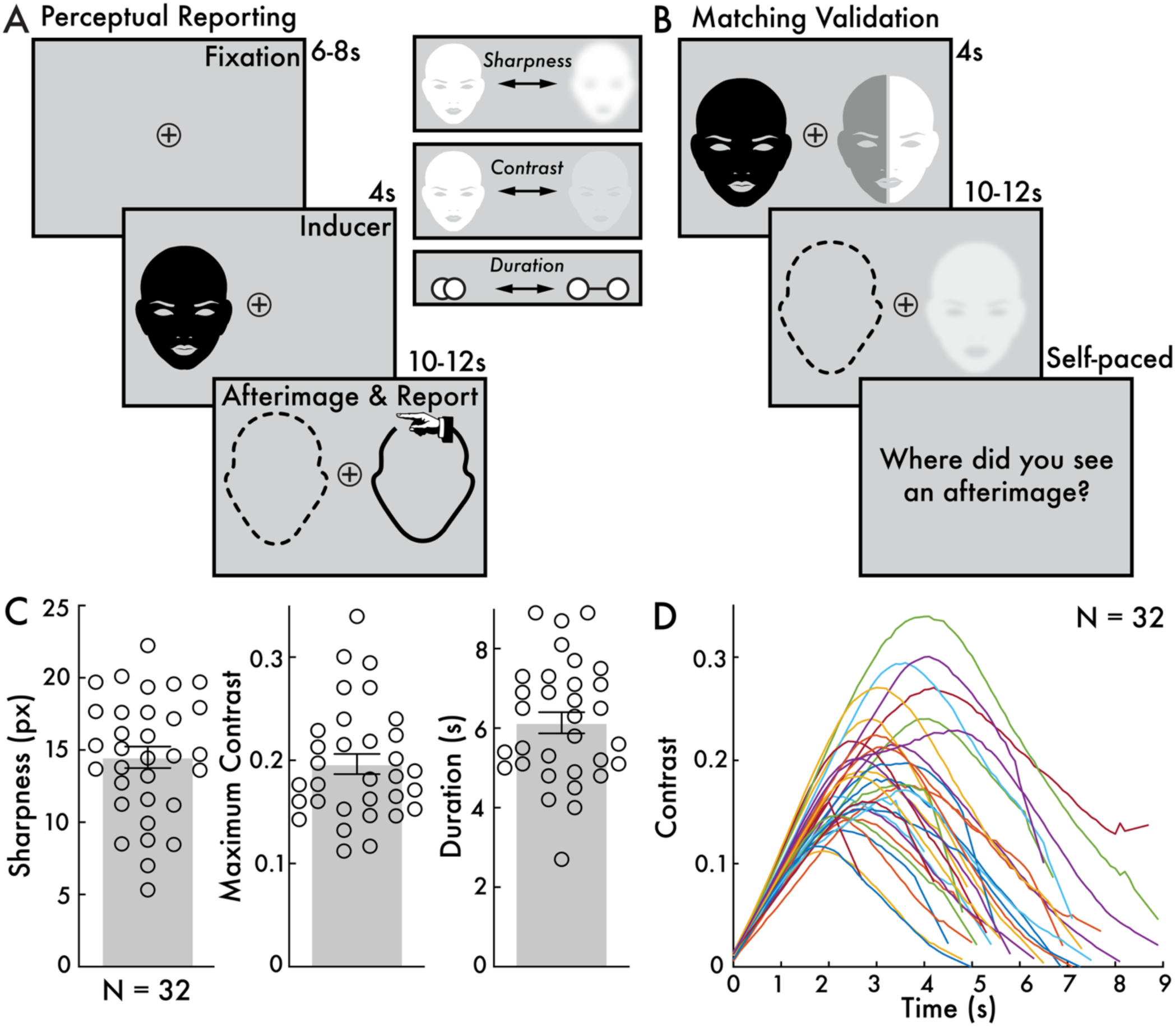
Perceptual reporting task, perceptual matching validation task, and mock afterimage parameters. **(A)** The afterimage and image perceptual reporting task involved participants indicating their perceived sharpness, contrast, and duration of afterimages and images using a controllable image (depicted with a hand icon to represent that participants could manipulate its appearance using key presses). The afterimage is depicted with a dotted outline to highlight that the screen remained blank when afterimages were perceived. For full details on the perceptual reporting task see [39]. **(B)** The matching validation task was administered to evaluate the efficacy of perceptual similarity between the afterimage and mock afterimage. In a subset of trials, the afterimage and mock afterimage were perceived side-by-side. Subsequently, participants were asked to report where they perceived an afterimage. Participants were not initially informed that the mock afterimage was an on-screen stimulus, thereby the instruction to report seen afterimages was inclusive of both afterimages and mock afterimages. **(C)** The sharpness, maximum contrast, and duration of the mock afterimages across participants. Each open circle represents individual participants. Larger sharpness values correspond with blurrier perception. **(D)** The contrast by time trend for each participant’s mock afterimage. When the mock afterimage was presented, it was animated to change its contrast over time, following these trend lines. Sharpness was kept constant throughout the mock afterimage presentation. See *Supplementary Movie 2* for an example mock afterimage.

**Supplementary Figure 2.**
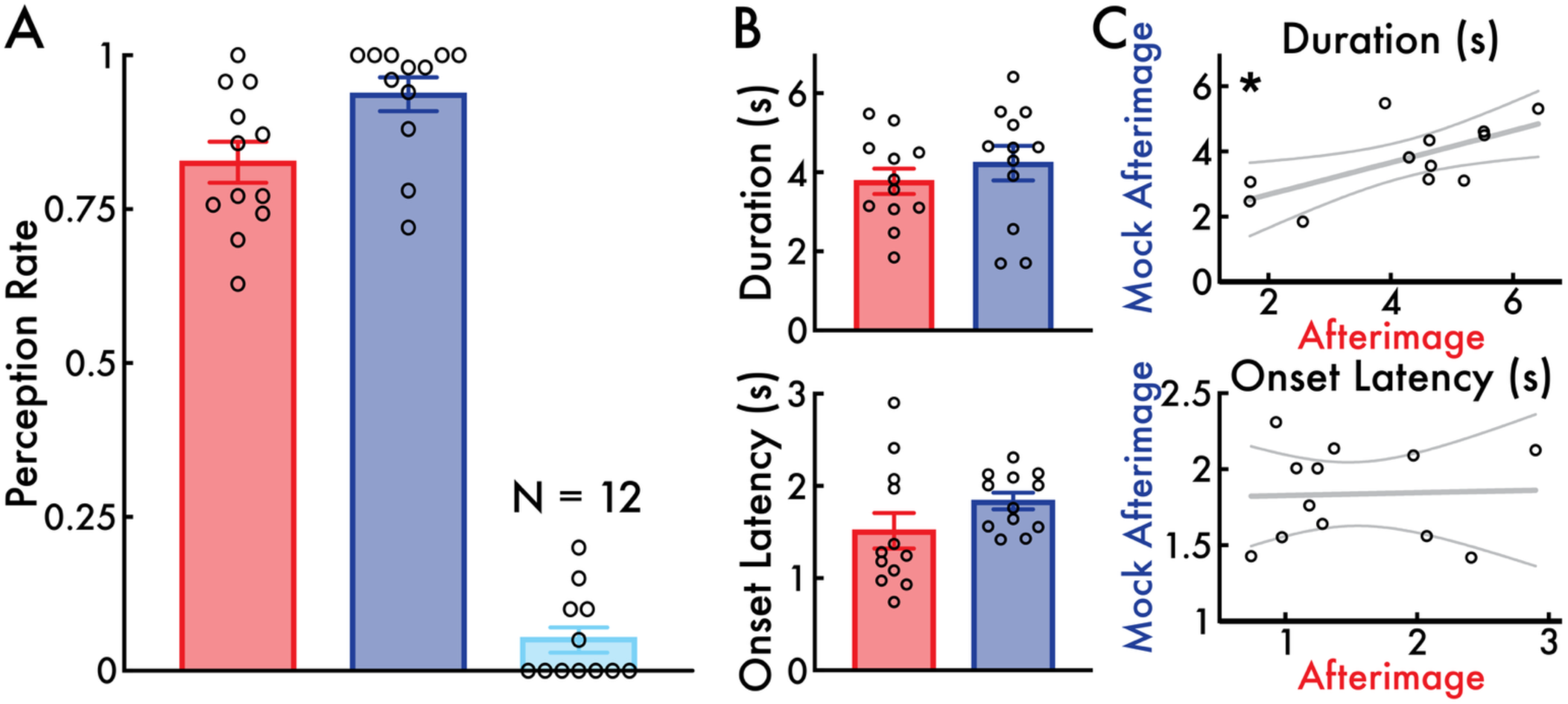
V1 fMRI session behavioral results. **(A)** The mean perception rate with standard error of the mean (SEM) for afterimages following the inducer (red) and mock afterimages following the mock inducer when the mock afterimage was present (dark blue) and absent (light blue). Participants regularly perceived afterimages (> 0.75) and mock afterimages (> 0.90), but not when the mock inducer was presented alone (i.e., without a mock afterimage; < 0.10). **(B)** The mean afterimage and mock afterimage duration and onset latency with SEM. The afterimage and mock afterimage durations and onset latencies were not statistically different. **(C)** The correlation between the afterimage and mock afterimage durations and onset latencies with a linear regression line and 95% confidence intervals. Statistically significant positive correlations were found for duration (* *p* < 0.05) but not onset latency. The open circles represent individual participants (N = 12). See the *Behavior* Analysis Methods section for full details on the behavioral analyses.

**Supplementary Figure 3.**
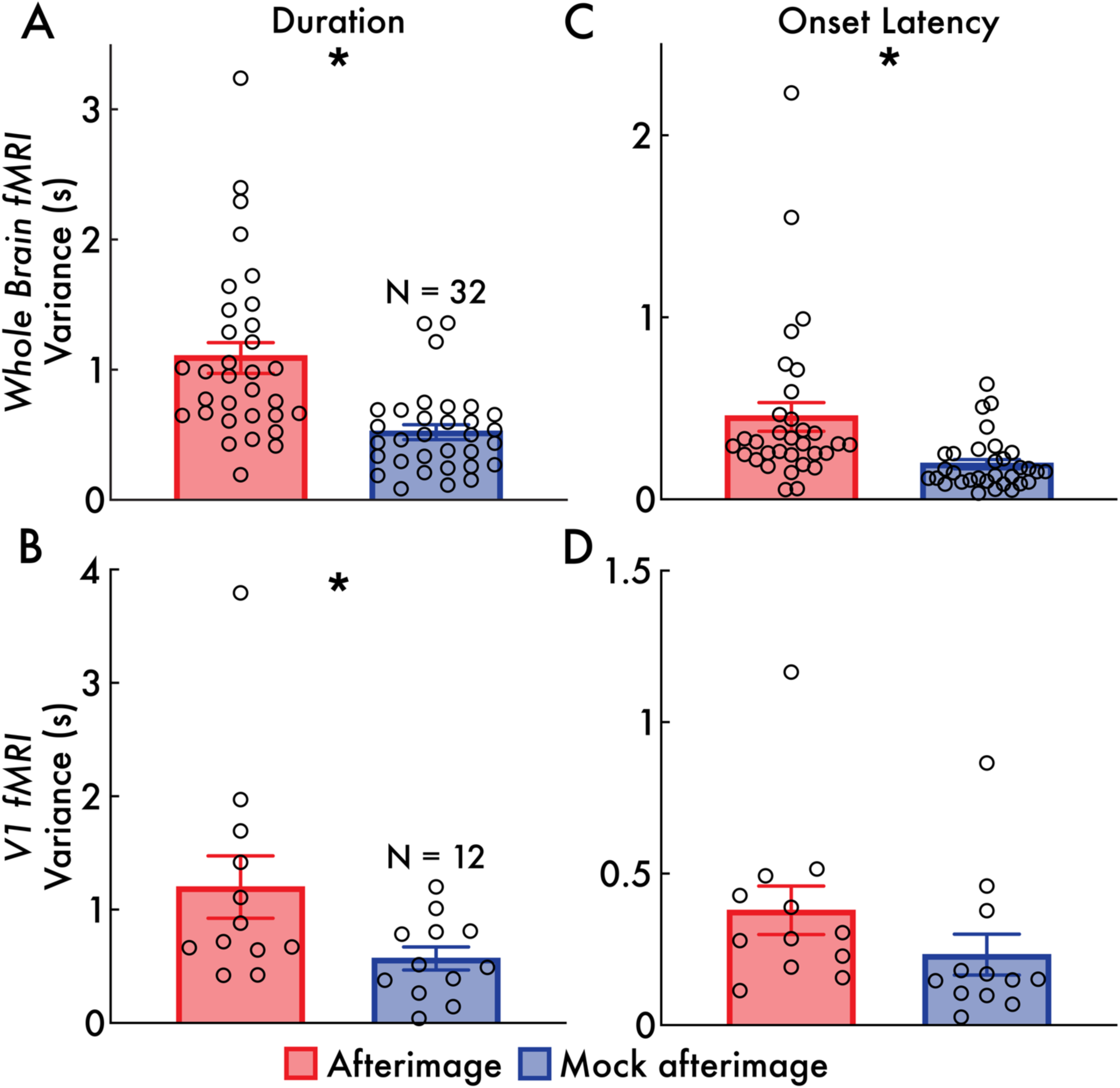
Within participant variance for reported afterimage and mock afterimage duration and onset latency. Duration variance (seconds [s]) for afterimages (red) and mock afterimages (blue) during **(A)** the whole brain fMRI and **(B)** V1 fMRI study sessions. Onset latency variance for afterimages and mock afterimages during **(C)** the whole brain fMRI and **(D)** V1 fMRI study sessions. Variance is calculated within participant and across all reported perceived afterimages and mock afterimages. In all subplots, the bar height indicates the mean variance, error bars indicate the standard error of the mean, and the open circles indicate individual participants (whole brain fMRI: N = 32; V1 fMRI: N = 12). Statistically significant differences between afterimage and mock afterimage variances are shown with an asterisk (* *p* < 0.05).

**Supplementary Figure 4.**
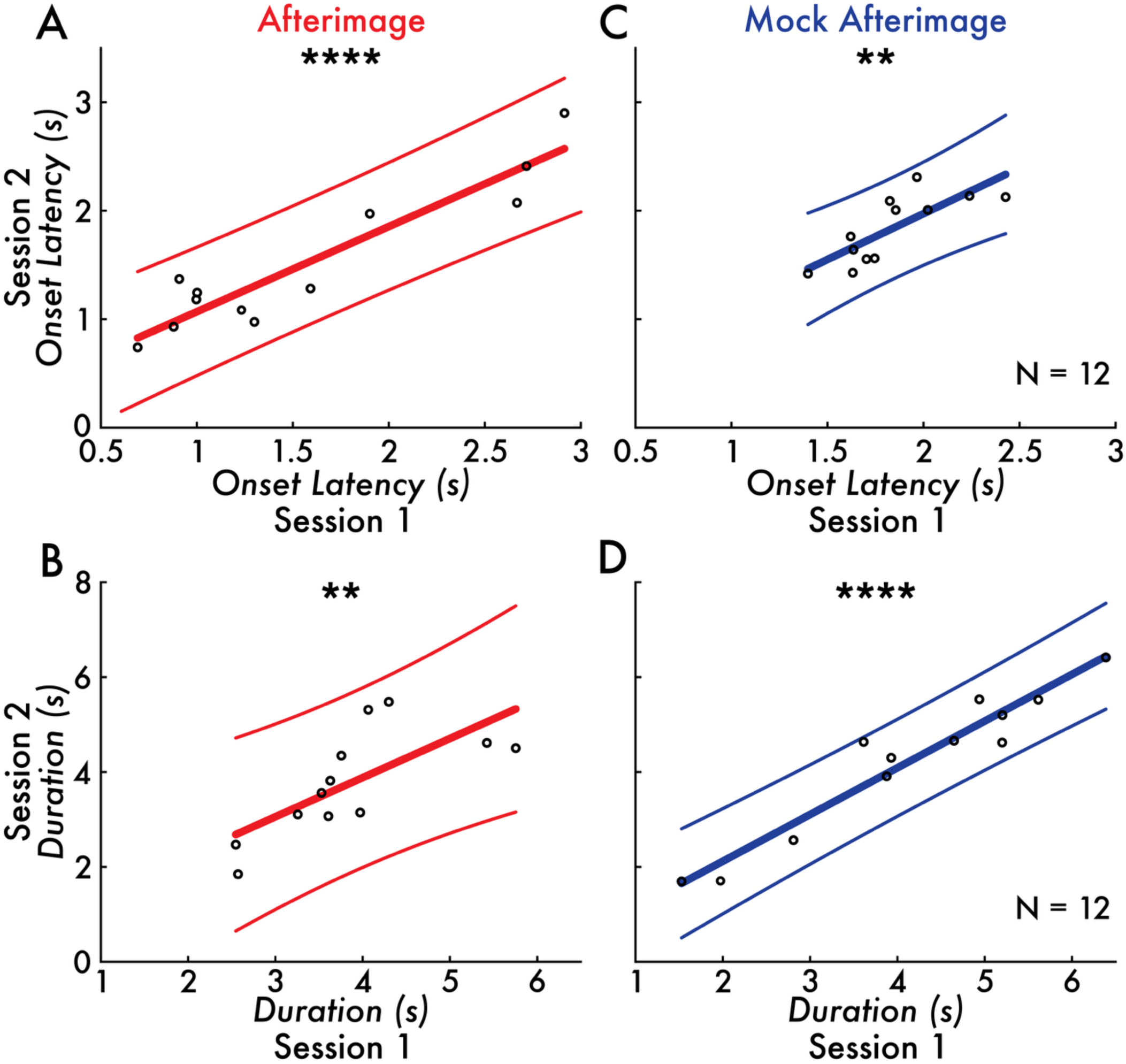
Correlation of afterimage and mock afterimage onset latency and duration across whole brain and V1 fMRI study sessions. The correlation between session 1 (whole brain fMRI) and 2 (V1 fMRI) afterimage **(A)** onset latency and **(B)** duration and mock afterimage **(C)** onset latency and **(D)** duration with a linear regression line and 95% confidence intervals. Statistically significant positive correlations between session 1 and 2 were found for both afterimage and mock afterimage onset latency and duration (** *p* < 0.01; **** *p* < 0.0001). The unit of the onset latency and duration is seconds (s). The open circles represent individual participants (N = 12).

**Supplementary Figure 5.**
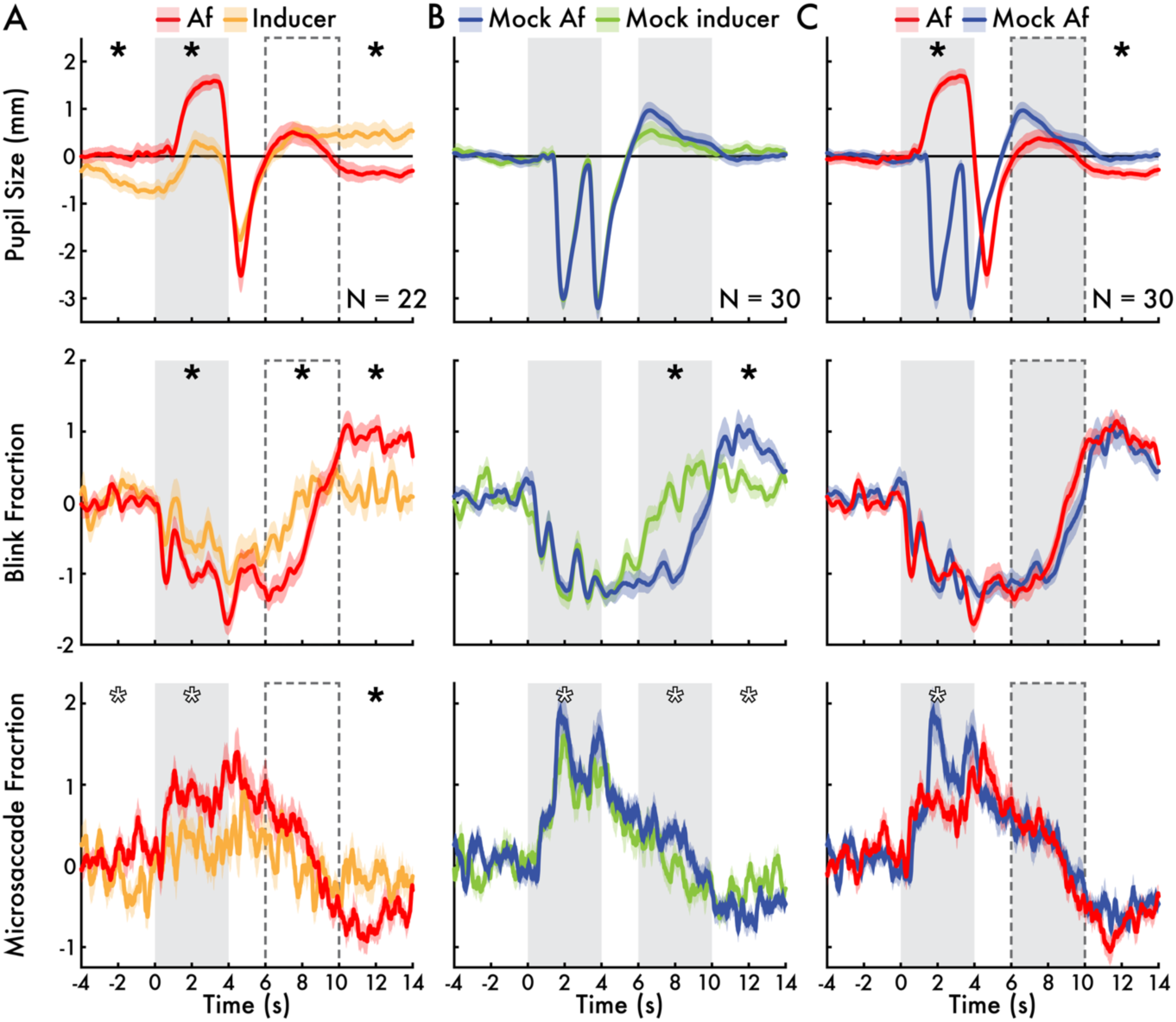
Perception task-evoked pupil size, blink, and microsaccade timecourses. **(A)** Inducers *with* afterimages versus inducers *without* afterimages conditions. **(B)** Mock inducers *with* mock afterimages versus mock inducers *with* mock afterimages conditions. **(C)** Inducers *with* afterimages versus mock inducers *with* mock afterimages conditions. The peaks and troughs during the mock inducer period correspond with the contrast flickering of the mock inducer, designed to suppress afterimages. In all subplots, the gray bar between 0 and 4 seconds (s) highlights the inducer and mock inducer period. The dotted open bar in A, solid gray bar in B, and solid gray bar with a dotted outline in C between 6 and 10 s highlight the approximate afterimage and mock afterimage period. The thicker solid timecourses indicate the mean pupil size in millimeters (mm), blink fraction (the proportion of epochs with a blink at any given timepoint), and microsaccade fraction (the proportion of epochs with a microsaccade at any given timepoint) with the standard error of the mean (shaded area). Statistically significant differences between the depicted conditions in mean eye behavior responses within the inducer and mock inducer (0-4 s), afterimage and mock afterimage (6-10 s), and post-afterimage and post-mock afterimage (10-14 s) intervals are indicated with asterisks (* *p* < 0.05; see *Eye tracking and pupillometry* Analysis Methods section; open asterisks indicate uncorrected *p* < 0.05 but did not survive correction for multiple comparisons). All values were standardized (z-scored) across participants (see *Eye tracking and pupillometry* Analysis Methods section).

**Supplementary Figure 6.**
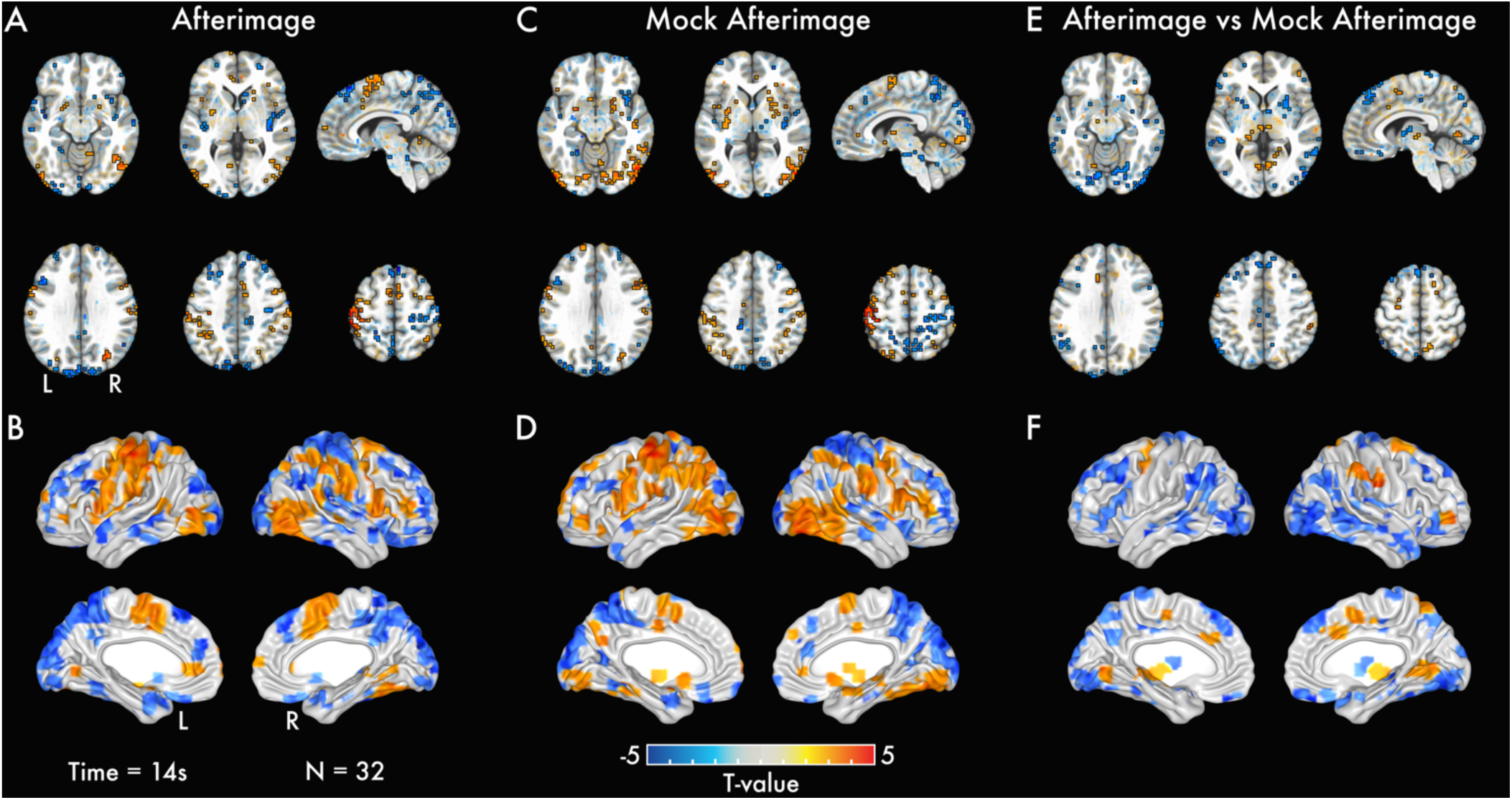
Whole brain volume and surface BOLD activation maps – no temporal normalization. Statistically significant *t*-values are highlighted with a black outline, while sub-threshold regions are shown with transparency at time point 14 seconds (s) after the inducer and mock inducer onset – the approximate peak response time for afterimages and mock afterimages – for the following statistical contrasts: **(A)** inducers *with* afterimages versus inducers + mock inducers only conditions (*afterimage*), **(C)** mock inducers *with* mock afterimages versus inducers + mock inducers only conditions (*mock afterimage*), and **(E)** inducers *with* afterimages versus mock inducers *with* mock afterimages conditions (*afterimage vs mock afterimage*). Brain surface visualization of *t*-values from statistically significant voxels are shown in **(B)**, **(D)**, and **(F)** for the same contrasts highlighted in the volume activation maps. These results correspond with those presented in Figure 2 and Supplementary Slides 1, 2, and 3, but without applying a temporal normalization preprocessing procedure (see *Whole brain fMRI* Analysis Methods section). The consistency between the results with and without temporal normalization supports that these findings are not dependent on the temporal normalization preprocessing procedure.

**Supplementary Figure 7.**
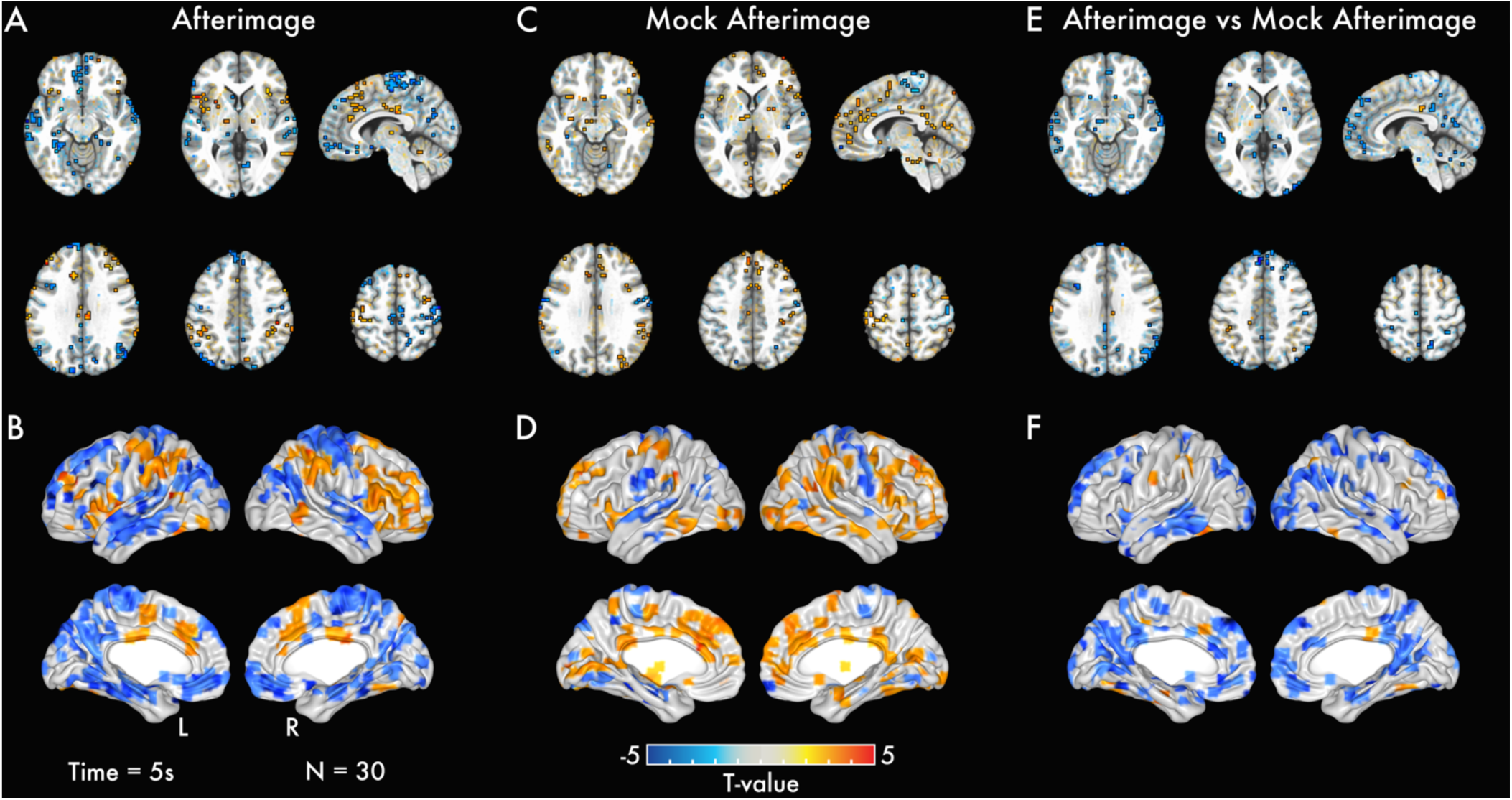
Whole brain volume and surface BOLD activation maps – general linear model. Statistically significant *t*-values are highlighted with a black outline, while sub-threshold regions are shown with transparency at time point 5 seconds (s) after afterimage and mock afterimage – the approximate peak response time for afterimages and mock afterimages – for the following statistical contrasts: **(A)** inducers *with* afterimages versus baseline (*afterimage*), **(C)** mock inducers *with* mock afterimages versus baseline (*mock afterimage*), and **(E)** inducers *with* afterimages versus mock inducers *with* mock afterimages conditions (*afterimage vs mock afterimage*). Brain surface visualization of *t*-values from statistically significant voxels are shown in **(B)**, **(D)**, and **(F)** for the same contrasts highlighted in the volume activation maps. The consistency between the general linear model (GLM) and cluster-based permutation testing results (Figure 2 and Supplementary Figure 6) supports that these findings are not dependent on the group-level analysis approach or nuisance variables accounted for in the GLM (e.g., blinking; see *Whole brain fMRI* Analysis Methods section).

**Supplementary Figure 8.**
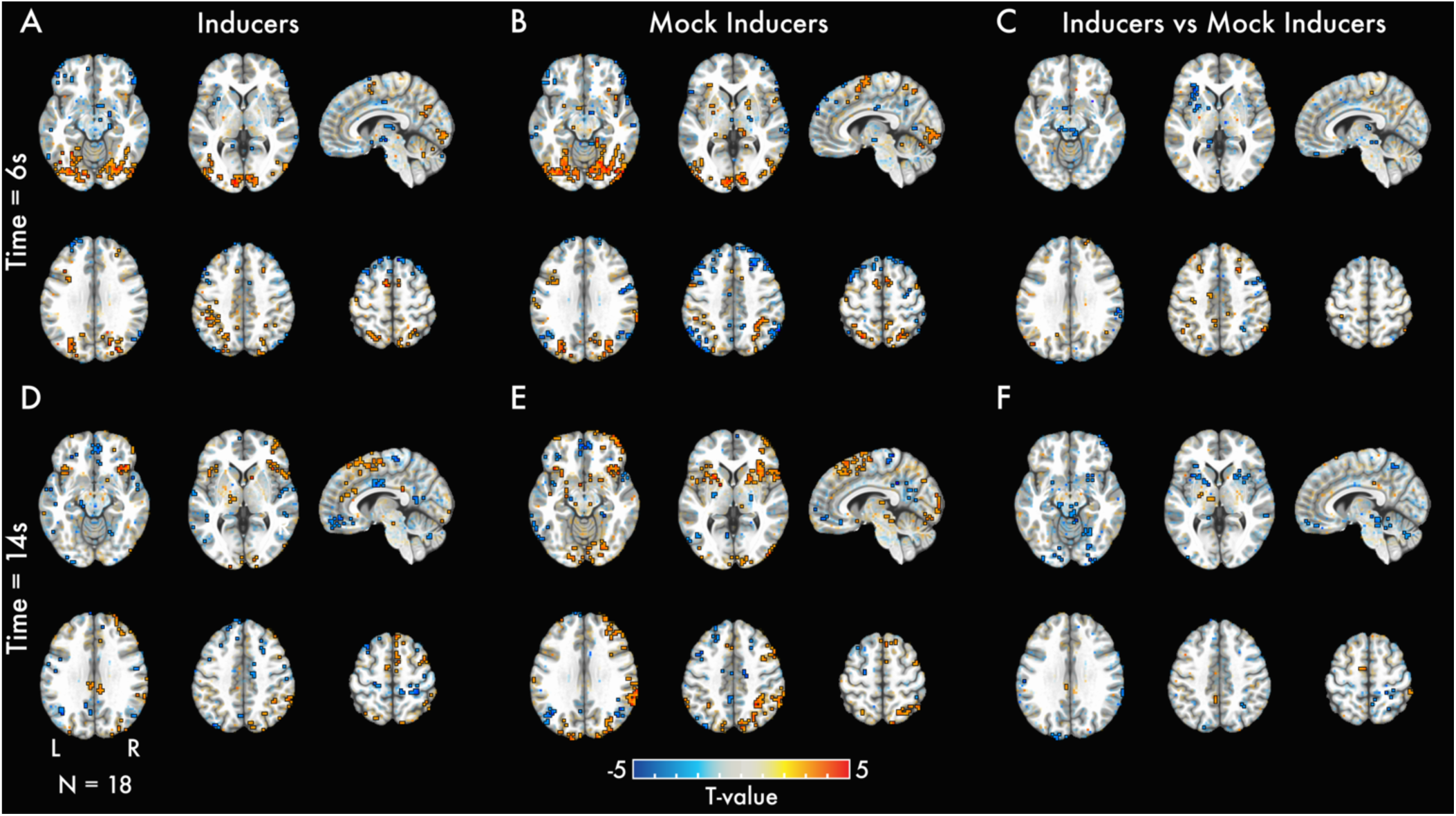
Inducers versus mock inducers whole brain volume BOLD activation maps. Statistically significant *t*-values are highlighted with a black outline, while sub-threshold regions are shown with transparency at representative time points (A, B, C) 6 seconds (s) – the approximate peak response time for inducers and mock inducers – and (D, E, F) 14 s – the approximate peak response time for afterimages and mock afterimages – after the inducer and mock inducer onset. Results are depicted for **(A**, **D)** inducers *without* afterimages, **(B**, **E)** mock inducers *without* mock afterimages, and **(C**, **F)** inducers *without* afterimages versus mock inducers *without* mock afterimages.

**Supplementary Figure 9.**
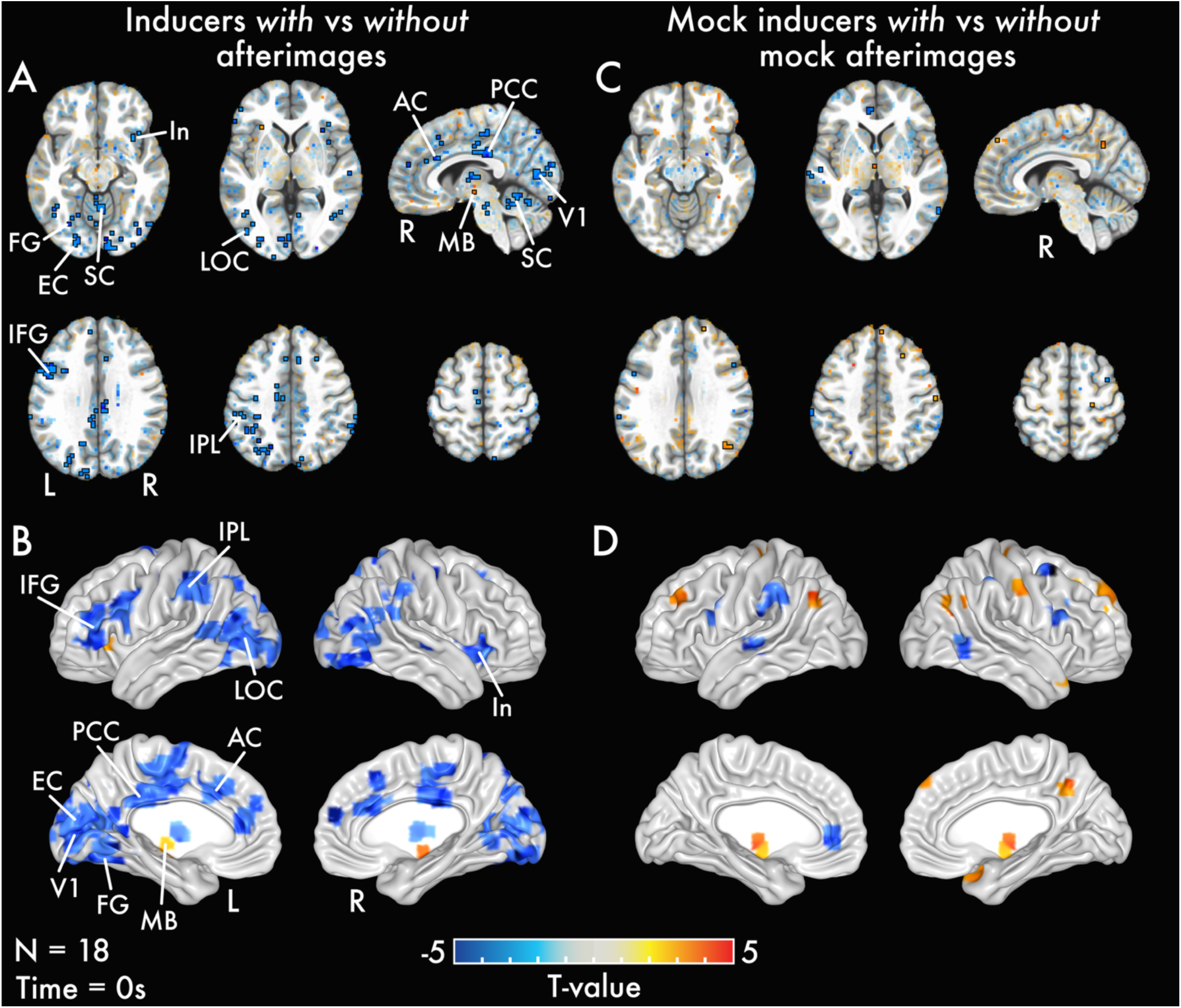
Inducers with versus without afterimages and mock inducers with versus without mock afterimages whole brain volume BOLD activation maps. Statistically significant *t*-values are highlighted with a black outline and sub-threshold regions are shown with transparency at representative time point 0 seconds (s) from the mock inducer onset. Results are depicted for **(A**, **B)** inducers *with* afterimages versus inducers *without* afterimages and **(C**, **D)** mock inducers *with* mock afterimages versus mock inducers *without* mock afterimages. See Supplementary Slides 4 and 5 for volume activation maps for additional time points. Anterior cingulate cortex (AC); extrastriate cortex (EC); fusiform gyrus (FG); inferior frontal gyrus (IFG); inferior parietal lobule (IPL); insula (In); lateral occipital cortex (LOC); midbrain (MB); posterior cingulate cortex (PCC); primary visual cortex (V1); superior cerebellum (SC); superior parietal lobule (SPL).

**Supplementary Figure 10.**
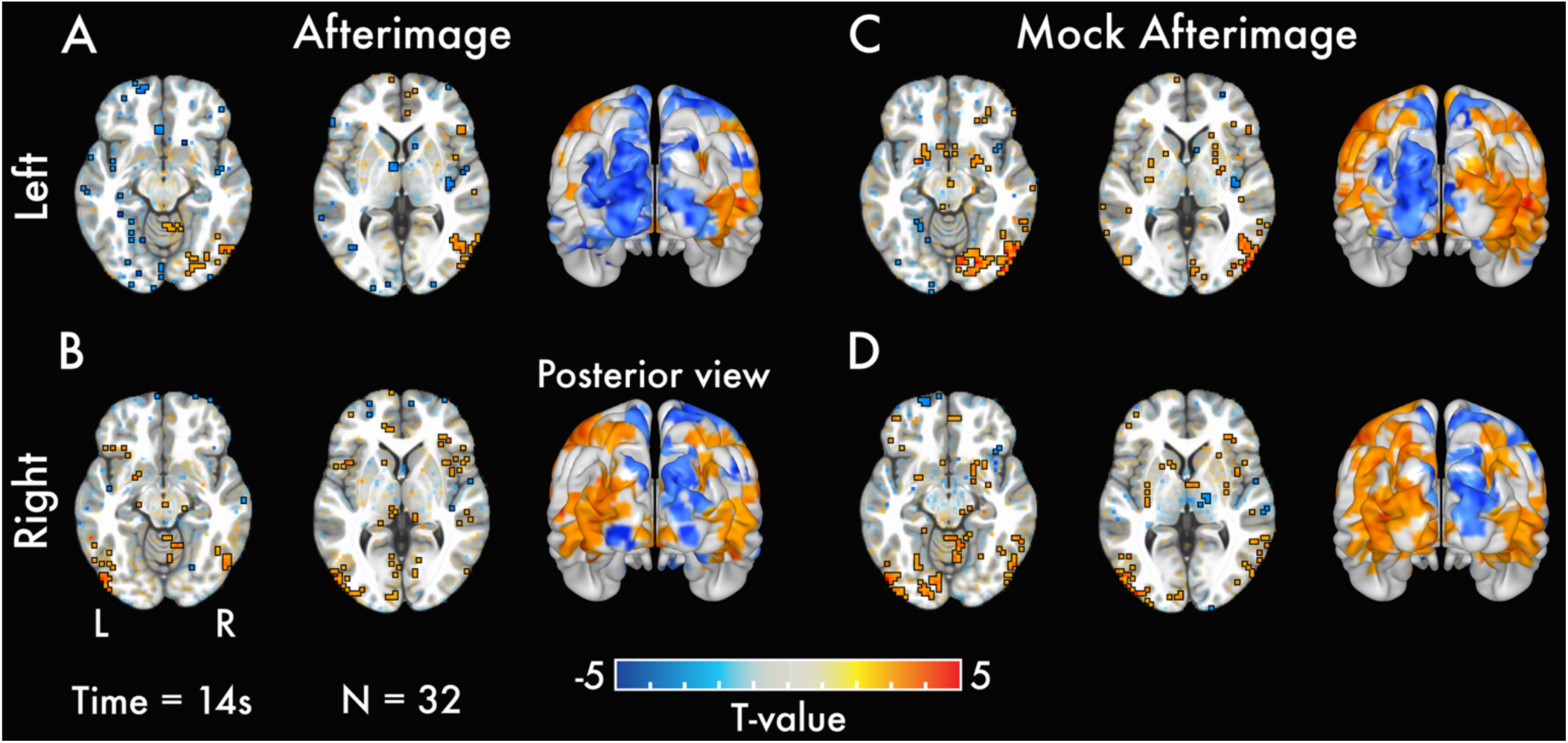
Left versus right visual field afterimage and mock afterimage whole brain volume BOLD activation maps. Statistically significant *t*-values are highlighted with a black outline, while sub-threshold regions are shown with transparency at representative time point 14 seconds (s) after the inducer and mock inducer onset – the approximate peak response time for afterimages and mock afterimages. Results are depicted for **(A)** left and **(B)** right visual field inducers *with* afterimages versus inducers + mock inducers only conditions (*afterimage*), and **(C)** left and **(D)** right visual field mock inducers *with* mock afterimages versus inducers + mock inducers only conditions (*mock afterimage*). Brain surface visualization of *t*-values from statistically significant voxels are shown in a posterior view to highlight visual cortex responses.

**Supplementary Figure 11.**
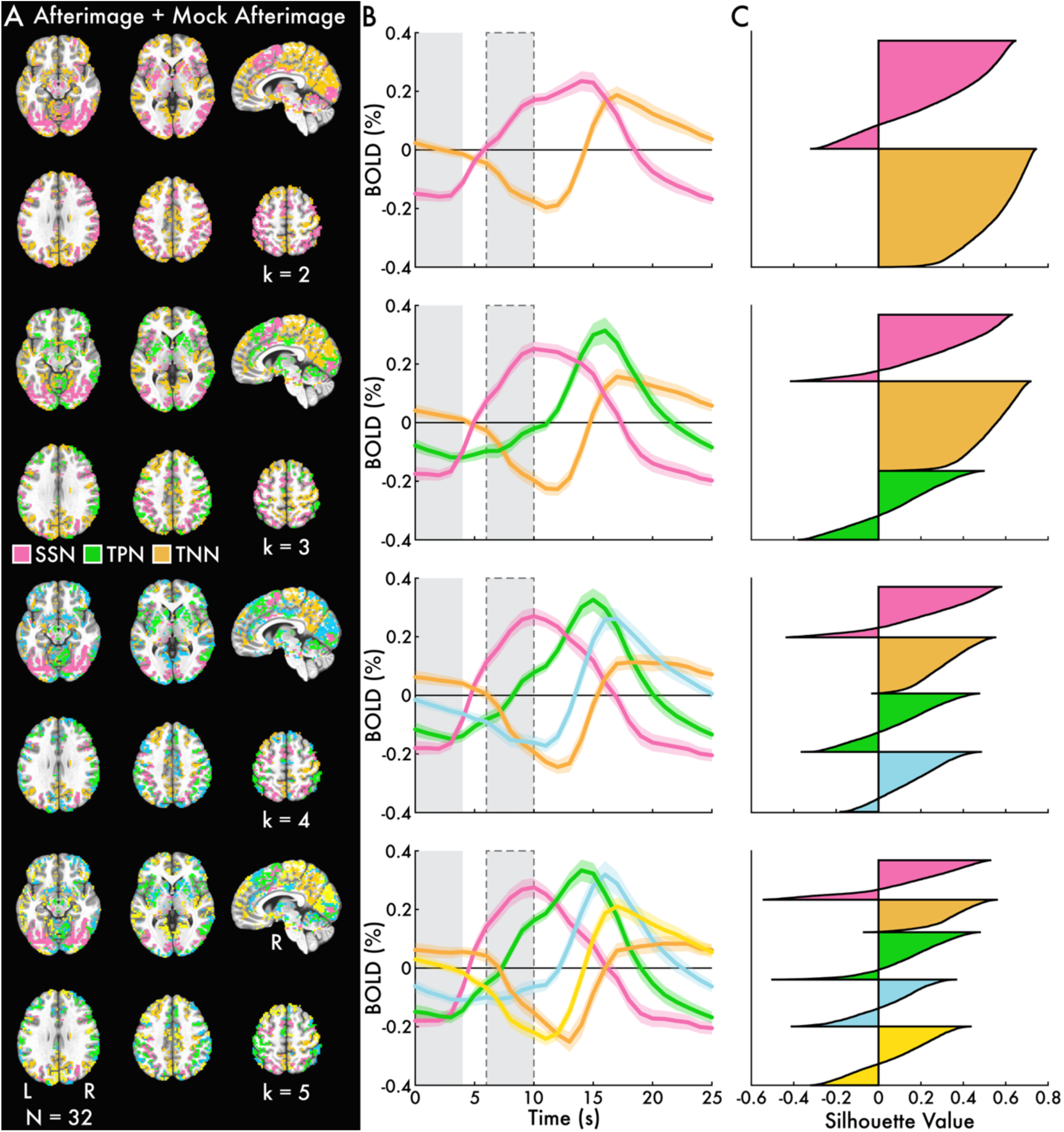
Whole brain BOLD spatiotemporal clustering brain maps, network timecourses, and silhouette values. K-means clusters (k = 2, 3, 4, and 5) computed for the average responses between the inducers *with* afterimages and mock inducers *with* mock afterimages conditions (*afterimage + mock afterimage*) for gray matter brain regions that showed statistically significant change in either the inducers *with* afterimages or mock inducers *with* mock afterimages conditions. Subplots depict **(A)** brain maps, **(B)** the mean percent change blood-oxygen-level-dependent (BOLD; %) timecourses with standard error of the mean (shaded area) across all voxels within the k-means clusters, and **(C)** silhouette values for all voxels within the k-means clusters. In B, the solid gray bar between 0 and 4 seconds (s) highlights the inducer and mock inducer period, while the solid gray bar with a dotted outline between 6 and 10 s highlights the approximate afterimage and mock afterimage period. The clusters for k = 3 corresponded with established functional brain networks: (1) *salience and sensory* (SSN), (2) *task positive* (TPN), and (3) *task negative* (TNN) networks (Figure 3A and B). The network labels approximate the functional role of the brain regions grouped within each cluster.

**Supplementary Figure 12.**
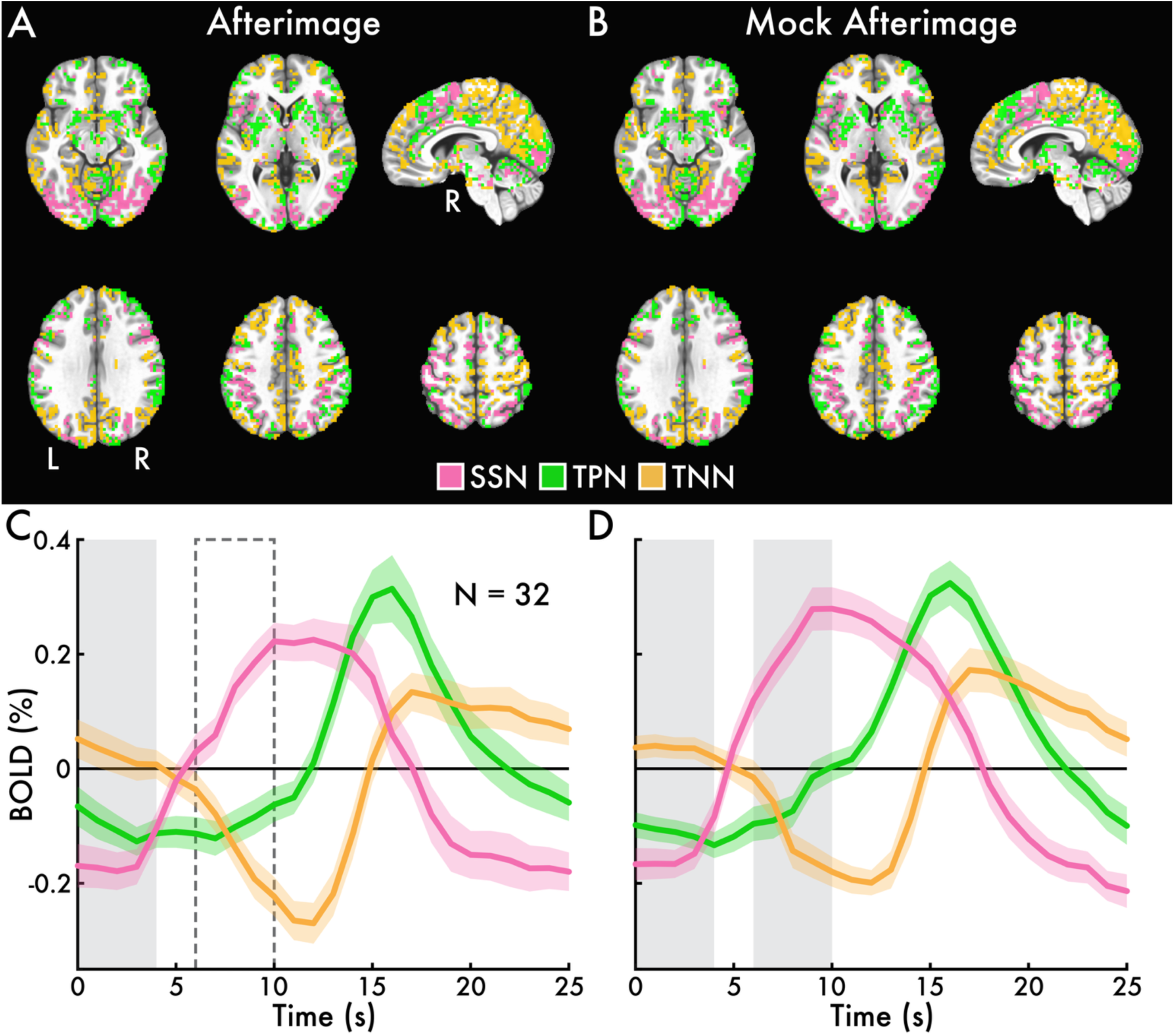
Afterimage and mock afterimage whole brain BOLD spatiotemporal clustering and network timecourses. K-means clusters (k = 3) of **(A)** the inducers *with* afterimages and **(B)** mock inducers *with* mock afterimages responses for gray matter brain regions that showed statistically significant change in either the inducers *with* afterimages or mock inducers *with* mock afterimages conditions. The clusters corresponded with established functional brain networks: (1) *salience and sensory* (SSN), (2) *task positive* (TPN), and (3) *task negative* (TNN) networks. The network labels approximate the functional role of the brain regions grouped within each cluster. The mean percent change blood-oxygen-level-dependent (BOLD; %) timecourses with standard error of the mean (shaded area) across all voxels within the SSN, TPN, and TNN for **(C)** the inducers *with* afterimages and **(D)** mock inducers *with* mock afterimages conditions. In B and D, the solid gray bar between 0 and 4 seconds (s) highlights the inducer and mock inducer period, while the dotted open bar in B and the solid gray bar in D between 6 and 10 s highlights the approximate afterimage and mock afterimage period.

**Supplementary Figure 13.**
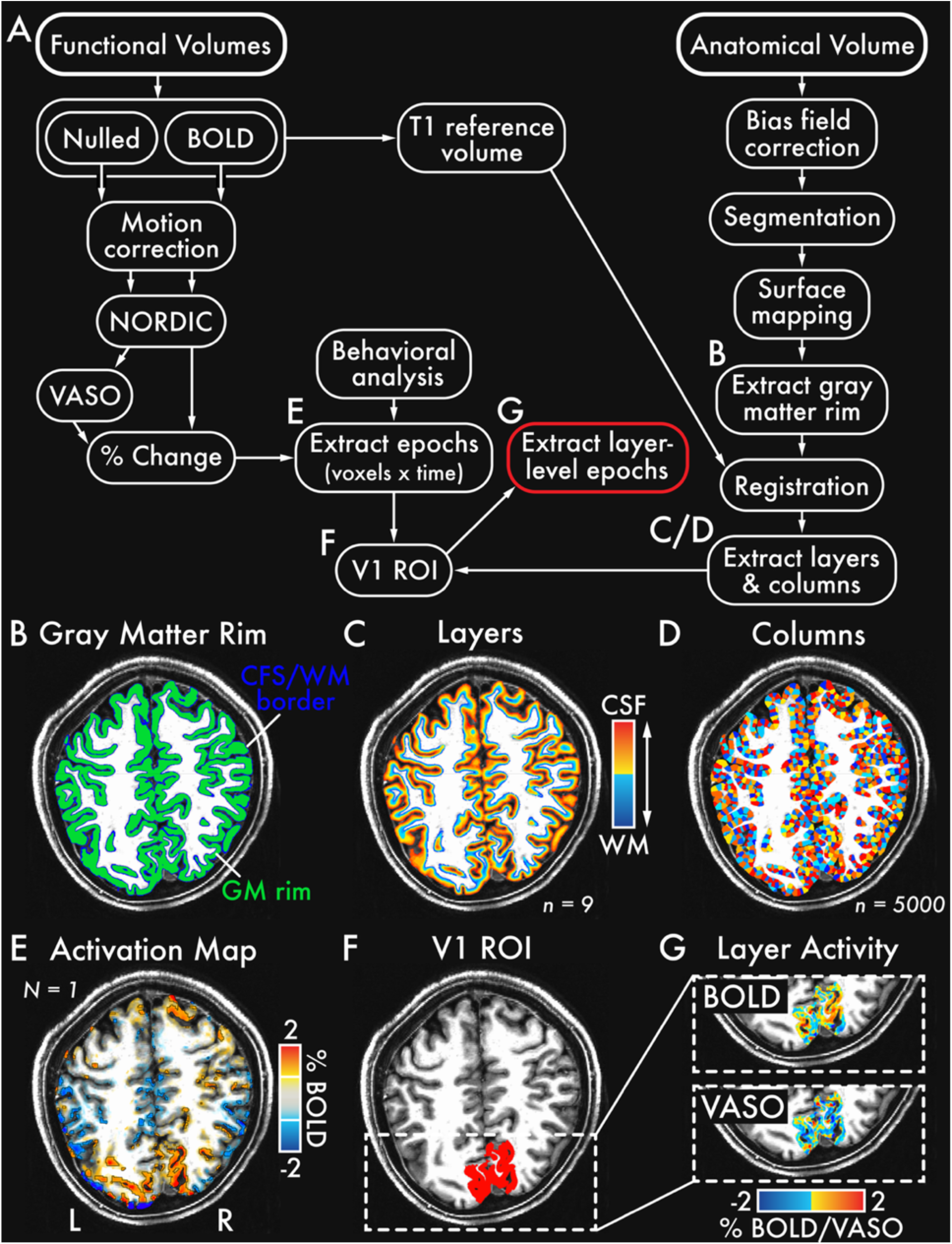
Layer resolution BOLD and VASO data processing and analysis summary. **(A)** Schematic of the layer resolution processing and analysis pipeline. Example images from a representative participant (same participant highlighted in Figure 5A and B) at different stages of the analysis. **(B)** Extracted cortical gray matter rim with the white matter (WM) and cerebral spinal fluid (CFS) borders. **(C)** Segmented layers (n = 9) within the gray matter rim (warmer colors indicate layers nearer to the CSF [superficial surface]; cooler colors indicate layers nearer to the WM [deep surface]). **(D)** Segmented columns (n = 5000) within the gray matter rim. **(E)** Percent change blood-oxygen-level-dependent (BOLD; %) activation map at 4 seconds after inducer and mock inducer onset for the inducers + mock inducers only condition. **(F)** Primary visual cortex (V1) region of interest (ROI) defined according to the activation map and cortical columns. **(G)** Percent change BOLD and vascular space occupancy (VASO) layer-resolution activity in the V1 ROI (highlighted in red to indicate the final output from the processing and analysis pipeline). See the *V1 fMRI sequence* and *V1 fMRI* Methods sections for full details on the layer resolution data acquisition, processing, and analysis.

**Supplementary Figure 14.**
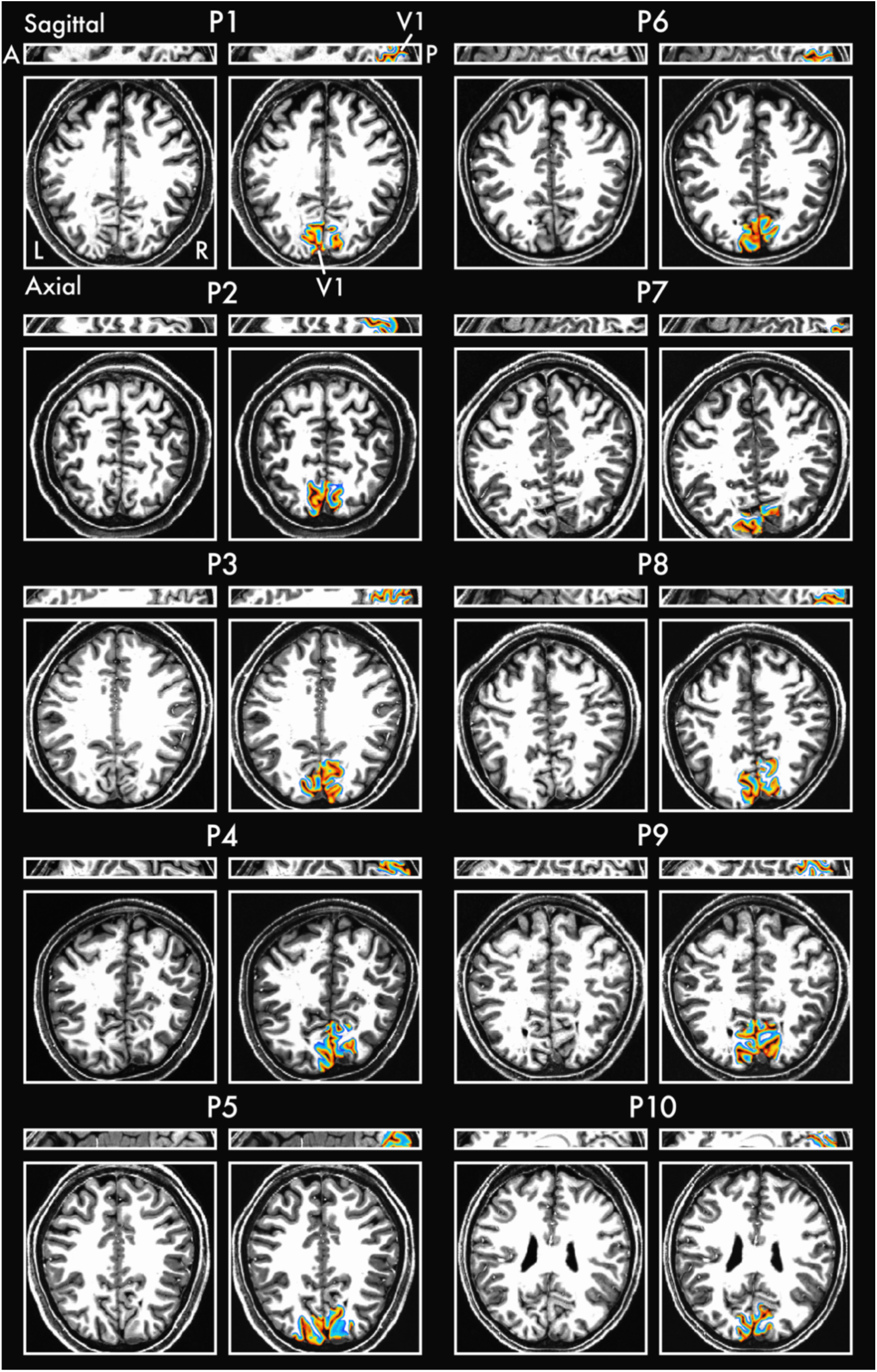
Anatomical brain image and V1 region of interest for all participants included in the layer-resolution fMRI analysis. Sagittal and axial plane visualization of anatomical brain images with (right) and without (left) the V1 region of interest (ROI) overlayed (see *V1 fMRI* Analysis Methods section for details for how the V1 ROI was defined). The V1 ROI shows the segmented layers (n = 9) within the gray matter rim (warmer colors indicate layers nearer to the CSF [superficial surface]; cooler colors indicate layers nearer to the WM [deep surface]).

**Supplementary Figure 15.**
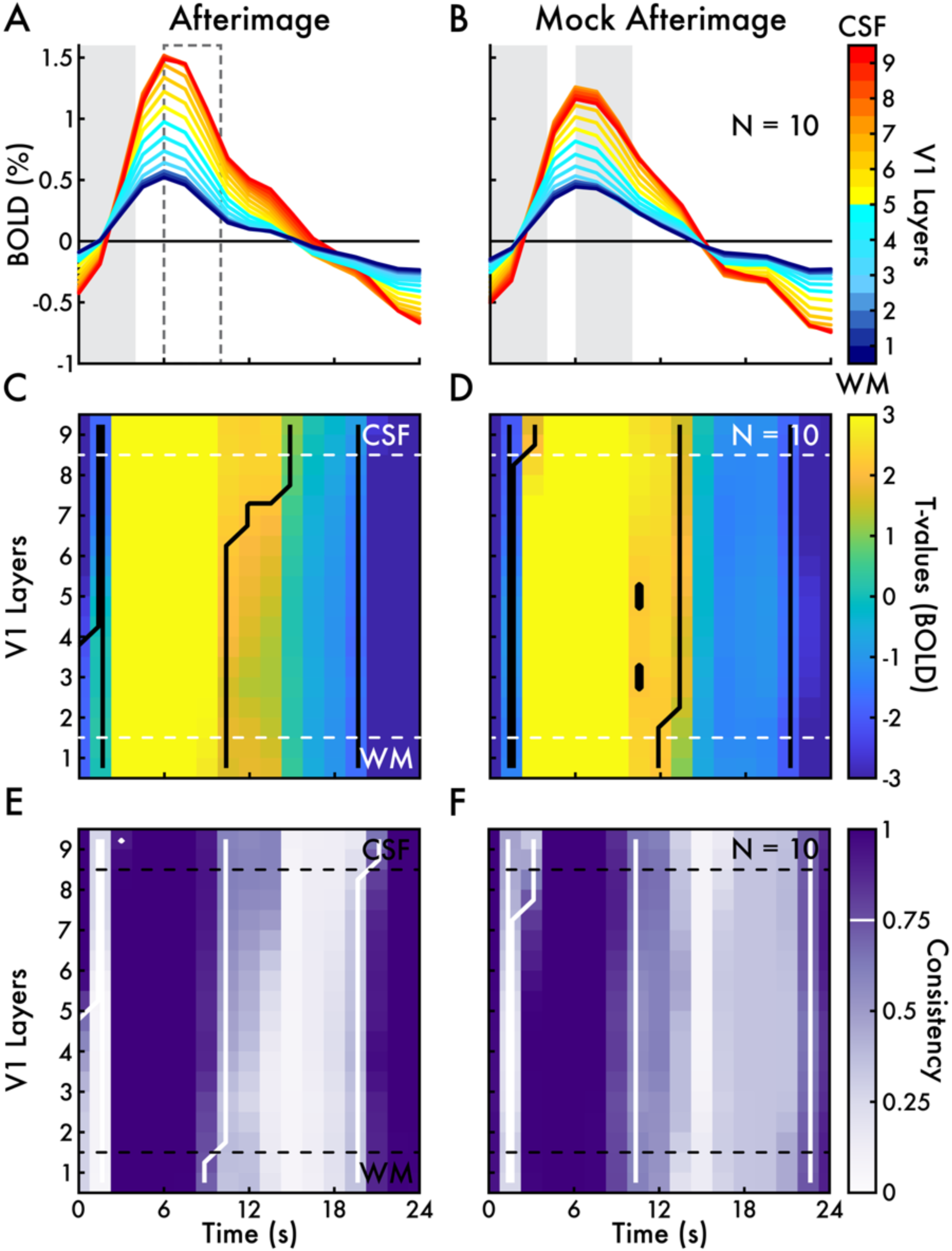
Afterimage and mock afterimage primary visual cortex layer-dependent BOLD activity. The percent change blood-oxygen-level-dependent (BOLD; %) across cortical layers for **(A)** inducers *with* afterimages (*afterimage*) and **(B)** mock inducers *with* mock afterimages (*mock afterimage*) conditions. In A and B, the gray bar between 0-4 seconds (s) indicates the inducer and mock inducer period. The dotted open bar in A and solid gray bar in B between 6 and 10 s highlight the approximate afterimage and mock afterimage period. Subplots **(C)** and **(D)** show the *t*-values for cortical layer-time cluster-based permutation testing. Statistically significant layer-time clusters are outlined with a solid black line. Subplots **(E)** and **(F)** show consistency maps from bootstrap analyses (5000 resamples) of the cortical layer-time cluster-based permutation testing (see *V1 fMRI* Analysis Methods section). All layer-time clusters with a consistency proportion > 0.75 (i.e., the layer-time cluster was statistically significant for more than 75% of bootstrap iterations) are outlined with a solid white line. Cerebrospinal fluid (CSF); white matter (WM).

**Supplementary Figure 16.**
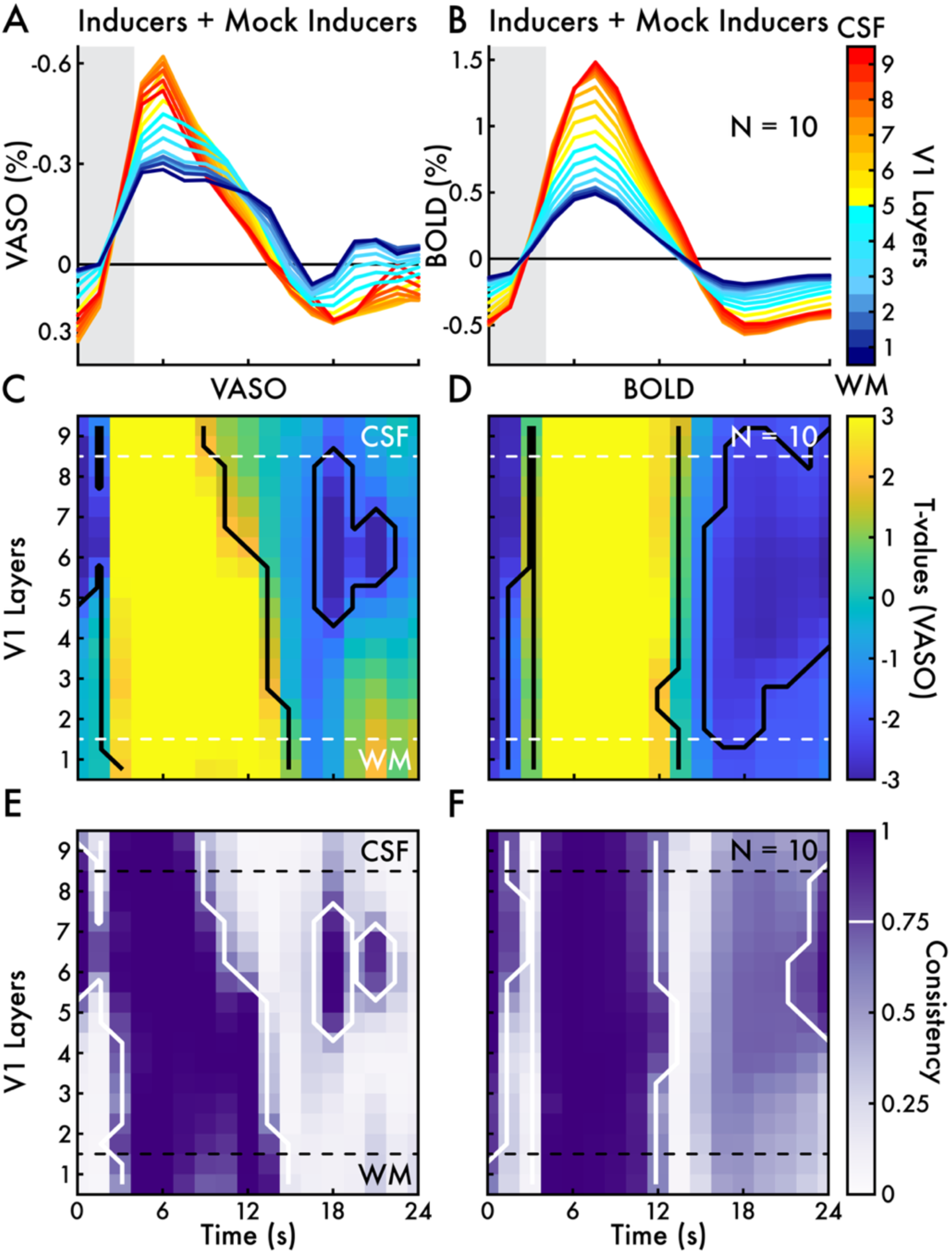
Inducers + mock inducers only condition primary visual cortex layer-dependent VASO and BOLD activity. The percent change **(A)** vascular space occupancy (VASO) and **(B)** blood-oxygen-level-dependent (BOLD; %) shown across cortical layers. In A and B, the gray bar between 0 and 4 seconds (s) highlights the inducer and mock inducer period. *T*-values from cortical layer-time cluster-based permutation testing for **(C)** VASO and **(D)** BOLD. Statistically significant layer-time clusters are outlined with a solid black line. Subplots **(E)** and **(F)** show consistency maps from bootstrap analyses (5000 resamples) of the cortical layer-time cluster-based permutation testing (see *V1 fMRI* Analysis Methods section). All layer-time clusters with a consistency proportion > 0.75 (i.e., the layer-time cluster was statistically significant for more than 75% of bootstrap iterations) are outlined with a solid white line. The percent change VASO signal was analyzed and shown inverted (negative is up) to enhance clarity and coherence with the BOLD signal. Cerebrospinal fluid (CSF); white matter (WM).

**Supplementary Figure 17.**
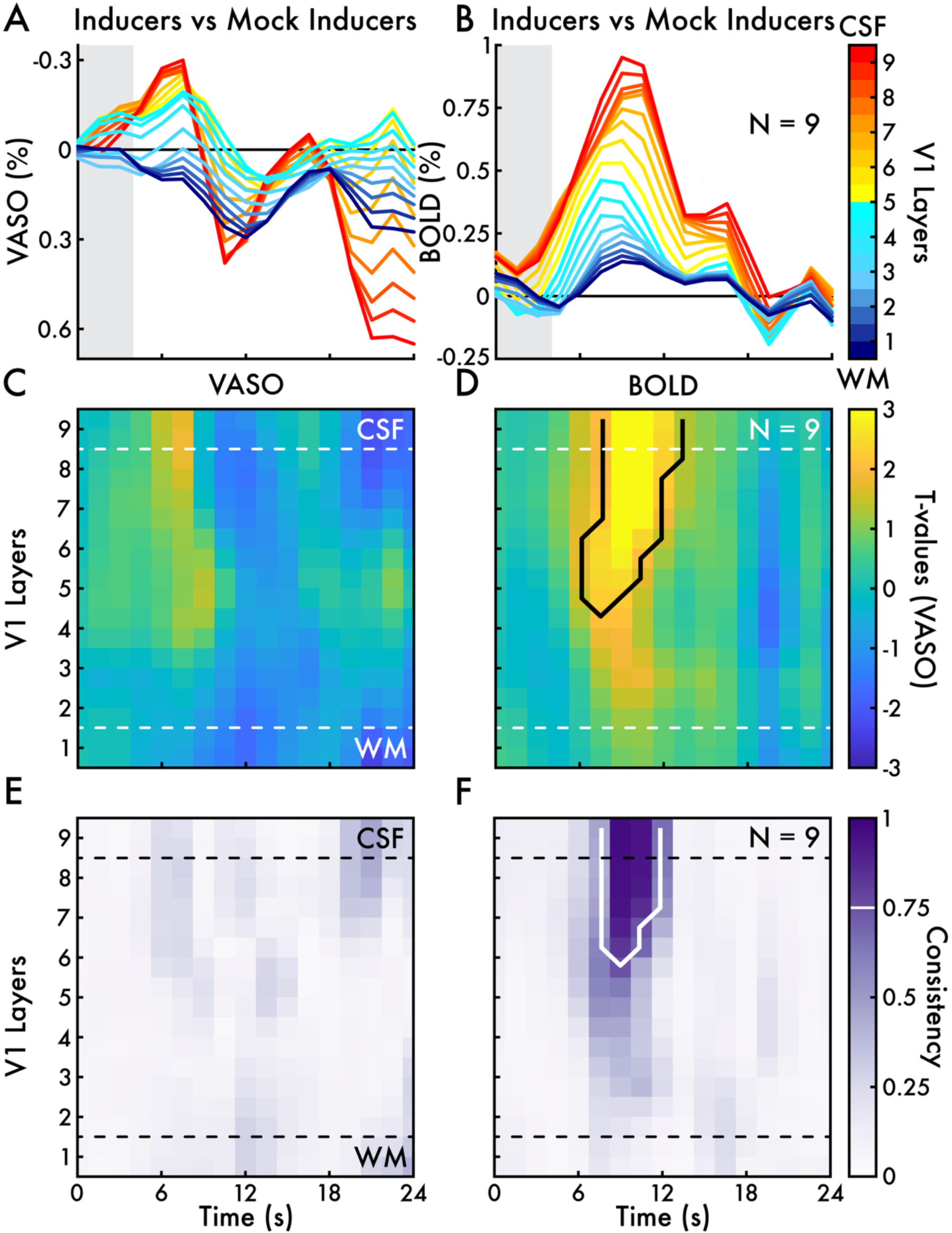
Inducers only versus mock inducers only condition primary visual cortex layer-dependent VASO and BOLD activity. The percent change **(A)** vascular space occupancy (VASO) and **(B)** blood-oxygen-level-dependent (BOLD; %) shown across cortical layers. In A and B, the gray bar between 0 and 4 seconds (s) highlights the inducer and mock inducer period. *T*-values from cortical layer-time cluster-based permutation testing for **(C)** VASO and **(D)** BOLD. Statistically significant layer-time clusters are outlined with a solid black line. Subplots **(E)** and **(F)** show consistency maps from bootstrap analyses (5000 resamples) of the cortical layer-time cluster-based permutation testing (see *V1 fMRI* Analysis Methods section). All layer-time clusters with a consistency proportion > 0.75 (i.e., the layer-time cluster was statistically significant for more than 75% of bootstrap iterations) are outlined with a solid white line. The percent change VASO signal was analyzed and shown inverted (negative is up) to enhance clarity and coherence with the BOLD signal. Cerebrospinal fluid (CSF); white matter (WM).

**Supplementary Figure 18.**
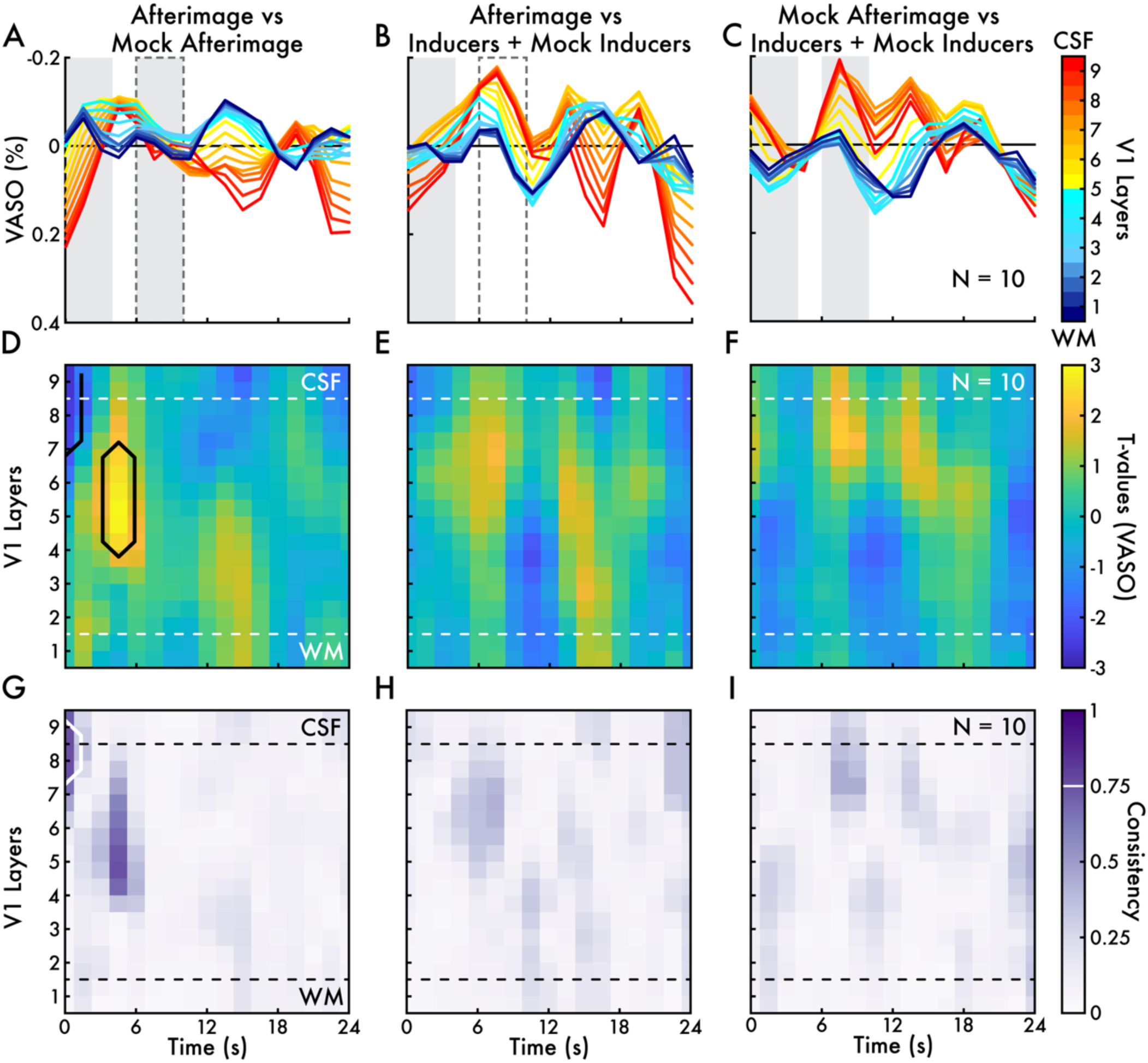
Primary visual cortex layer-dependent VASO activity. The percent change vascular space occupancy (VASO; %) across cortical layers for **(A)** inducers *with* afterimages versus mock inducers *with* mock afterimages (*afterimage vs mock afterimages*), **(B)** inducers *with* afterimages versus inducers + mock inducers only (*afterimage vs inducers + mock inducers*), and **(C)** mock inducers *with* mock afterimages versus inducers + mock inducers only (*mock afterimage vs inducers + mock inducers*). In A, B, and C, the gray bar between 0 and 4 seconds (s) highlights the inducer and mock inducer period. The solid gray bar with a dotted outline in A, dotted open bar in B, solid gray bar in C between 6 and 10 s highlights the approximate afterimage and mock afterimage period. *T*-values from cortical layer-time cluster-based permutation testing for **(D)** afterimage vs mock afterimages, **(E)** afterimage vs inducers + mock inducers, and **(F)** mock afterimage vs inducers + mock inducers. Statistically significant layer-time clusters are outlined with a solid black line. Subplots **(G)**, **(H)**, and **(I)** show consistency maps from bootstrap analyses (5000 resamples) of the cortical layer-time cluster-based permutation testing (see *V1 fMRI* Analysis Methods section). All layer-time clusters with a consistency proportion > 0.75 (i.e., the layer-time cluster was statistically significant for more than 75% of bootstrap iterations) are outlined with a solid white line. The percent change VASO signal was analyzed and shown inverted (negative is up) to enhance clarity and coherence with the BOLD signal. Cerebrospinal fluid (CSF); white matter (WM).

**Supplementary Figure 19.**
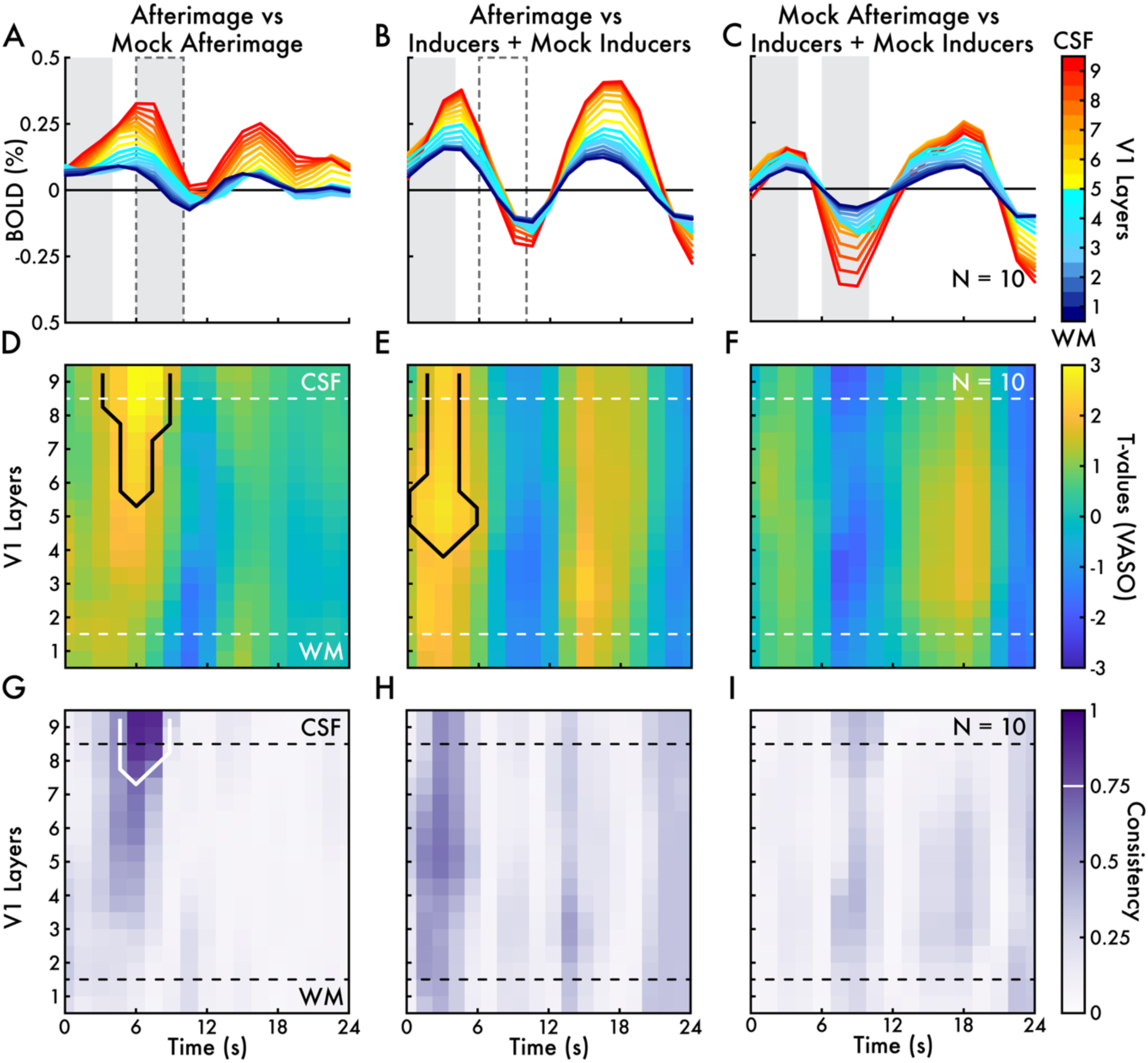
Primary visual cortex layer-dependent BOLD activity. The percent change blood-oxygen-level-dependent (BOLD; %) across cortical layers for **(A)** inducers *with* afterimages versus mock inducers *with* mock afterimages (*afterimage vs mock afterimages*), **(B)** inducers *with* afterimages versus inducers + mock inducers only (*afterimage vs inducers + mock inducers*), and **(C)** mock inducers *with* mock afterimages versus inducers + mock inducers only (*mock afterimage vs inducers + mock inducers*). In A, B, and C, the gray bar between 0 and 4 seconds (s) highlights the inducer and mock inducer period. The solid gray bar with a dotted outline in A, dotted open bar in B, solid gray bar in C between 6 and 10 s highlights the approximate afterimage and mock afterimage period. *T*-values from cortical layer-time cluster-based permutation testing for **(D)** afterimage vs mock afterimages, **(E)** afterimage vs inducers + mock inducers, and **(F)** mock afterimage vs inducers + mock inducers. Statistically significant layer-time clusters are outlined with a solid black line. Subplots **(G)**, **(H)**, and **(I)** show consistency maps from bootstrap analyses (5000 resamples) of the cortical layer-time cluster-based permutation testing (see *V1 fMRI* Analysis Methods section). All layer-time clusters with a consistency proportion > 0.75 (i.e., the layer-time cluster was statistically significant for more than 75% of bootstrap iterations) are outlined with a solid white line. Cerebrospinal fluid (CSF); white matter (WM).

**Supplementary Figure 20.**
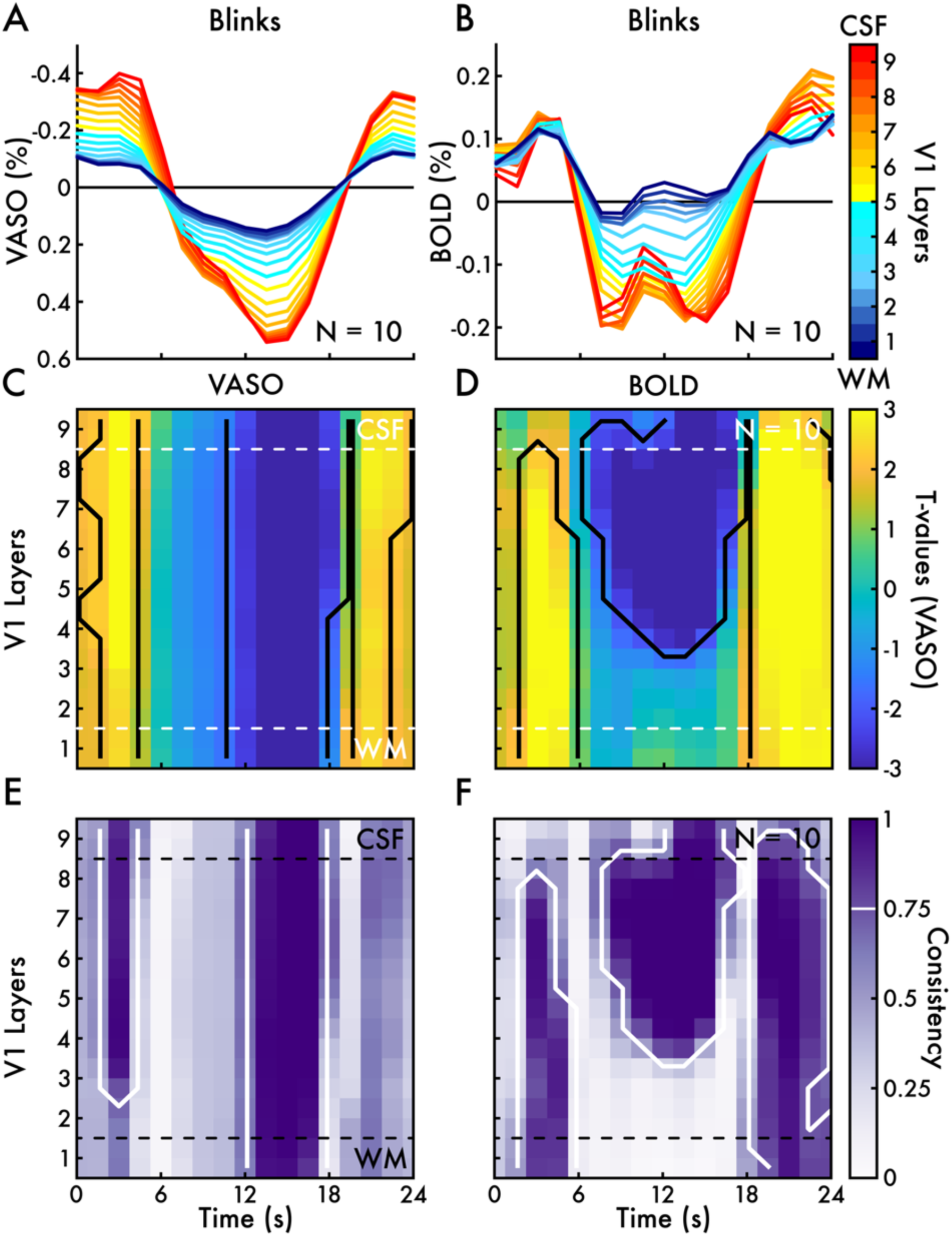
Eye blink primary visual cortex layer-dependent VASO and BOLD activity. The percent change **(A)** vascular space occupancy (VASO) and **(B)** blood-oxygen-level-dependent (BOLD; %) shown across cortical layers. *T*-values from cortical layer-time cluster-based permutation testing for **(C)** VASO and **(D)** BOLD. Statistically significant layer-time clusters are outlined with a solid black line. Subplots **(E)** and **(F)** show consistency maps from bootstrap analyses (5000 resamples) of the cortical layer-time cluster-based permutation testing (see *V1 fMRI* Analysis Methods section). All layer-time clusters with a consistency proportion > 0.75 (i.e., the layer-time cluster was statistically significant for more than 75% of bootstrap iterations) are outlined with a solid white line. The percent change VASO signal was analyzed and shown inverted (negative is up) to enhance clarity and coherence with the BOLD signal. Cerebrospinal fluid (CSF); white matter (WM).

