## Supplementary Slides 2 for "The human brain mechanisms of afterimages: From networks to cortical layers"

*Kronemer et al., 2025*

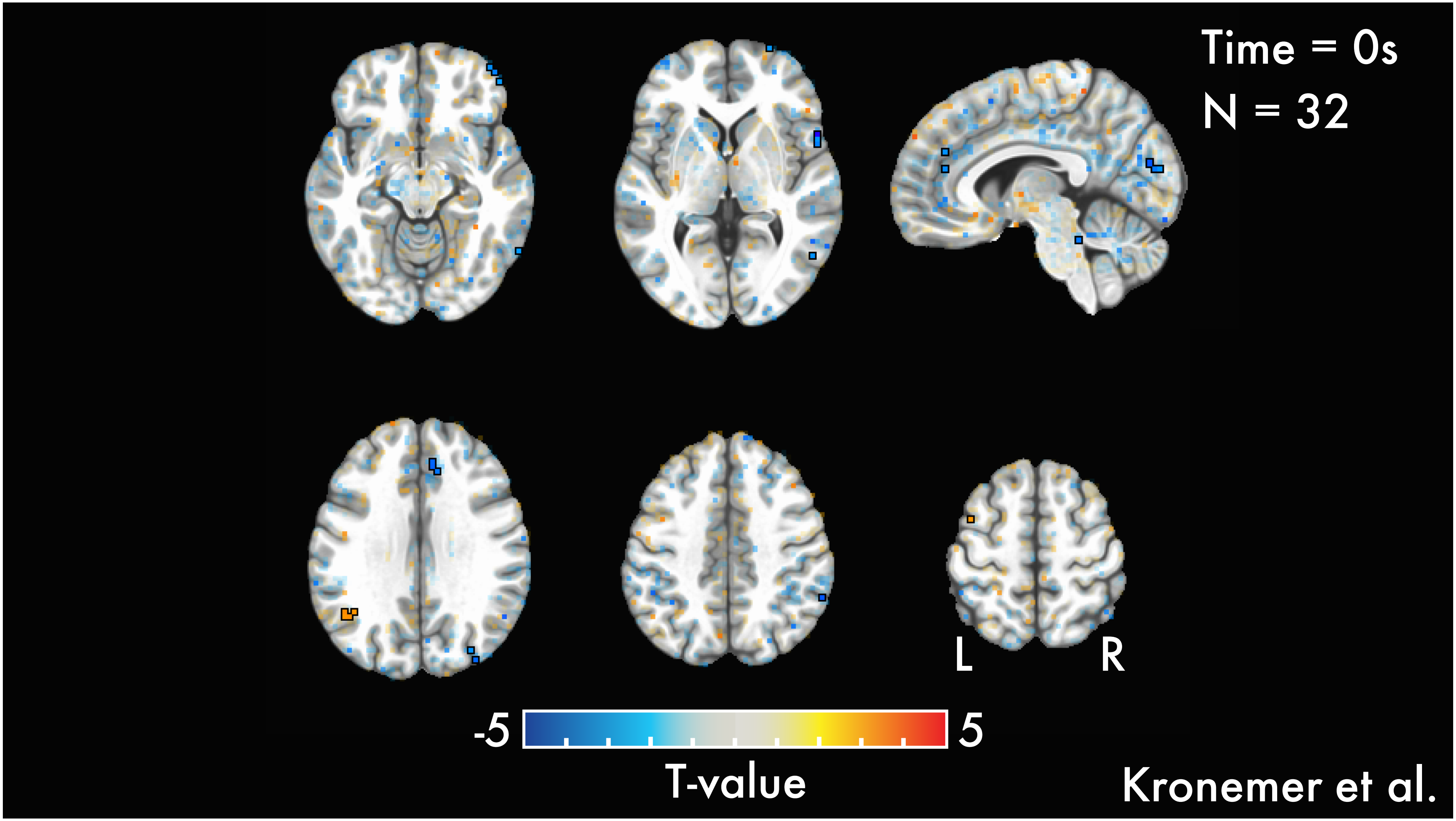
Time = 0s
N = 32
L
R
-5
5
T-value
Kronemer et al.

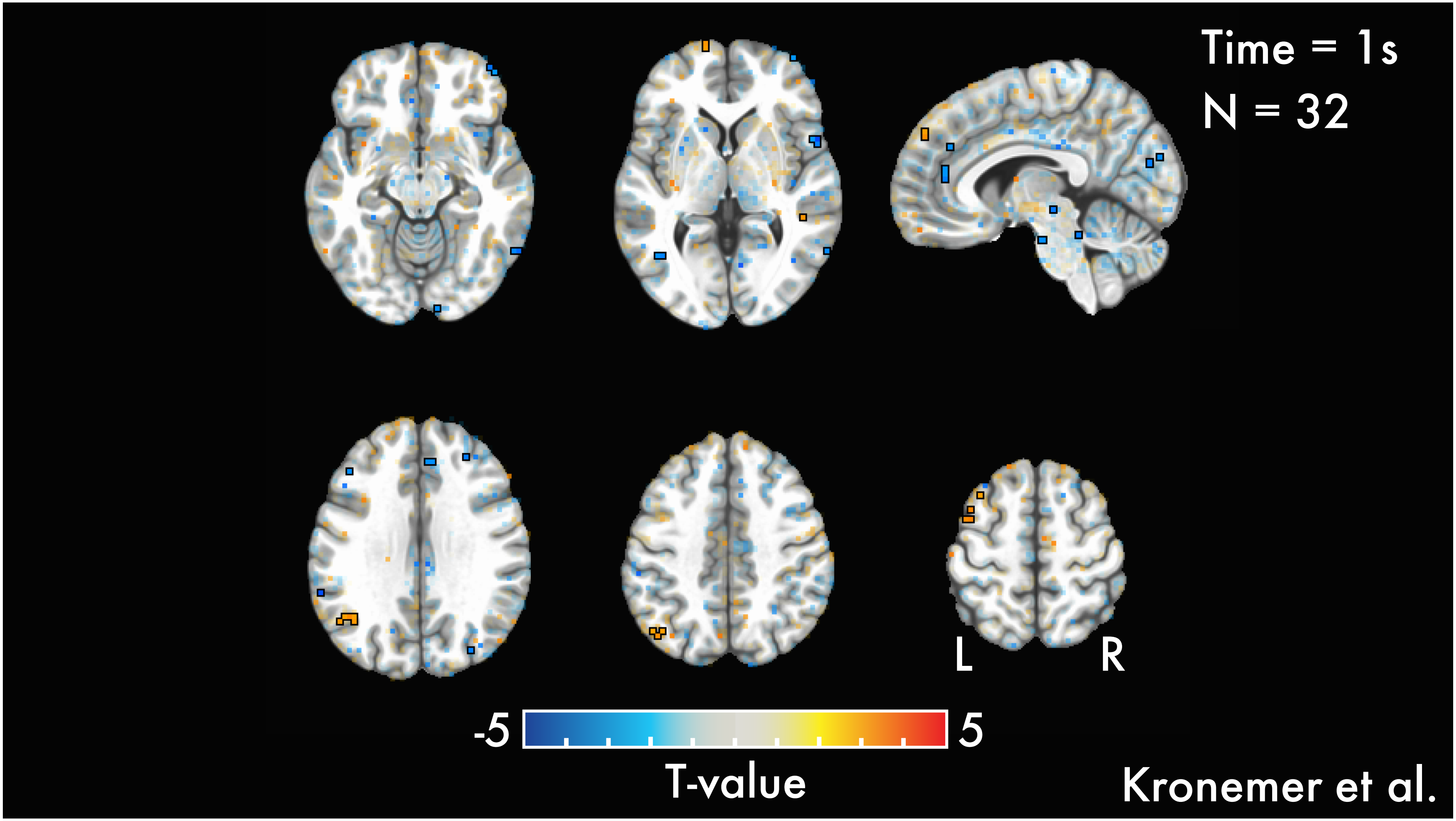

Time = 1s
N = 32
L
R
-5
5
T-value
Kronemer et al.

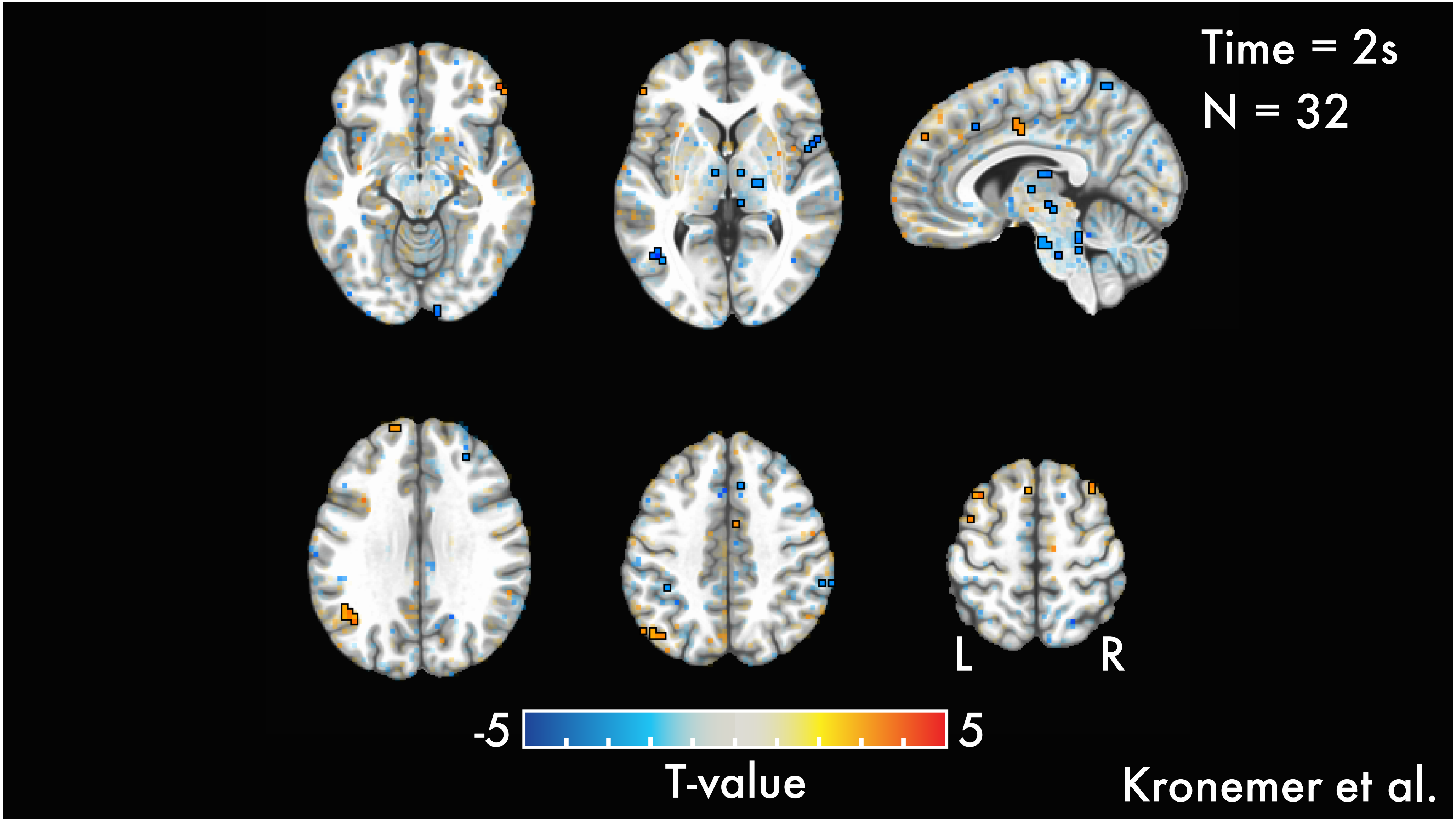

Time = 2s
N = 32
L
R
-5
5
T-value
Kronemer et al.

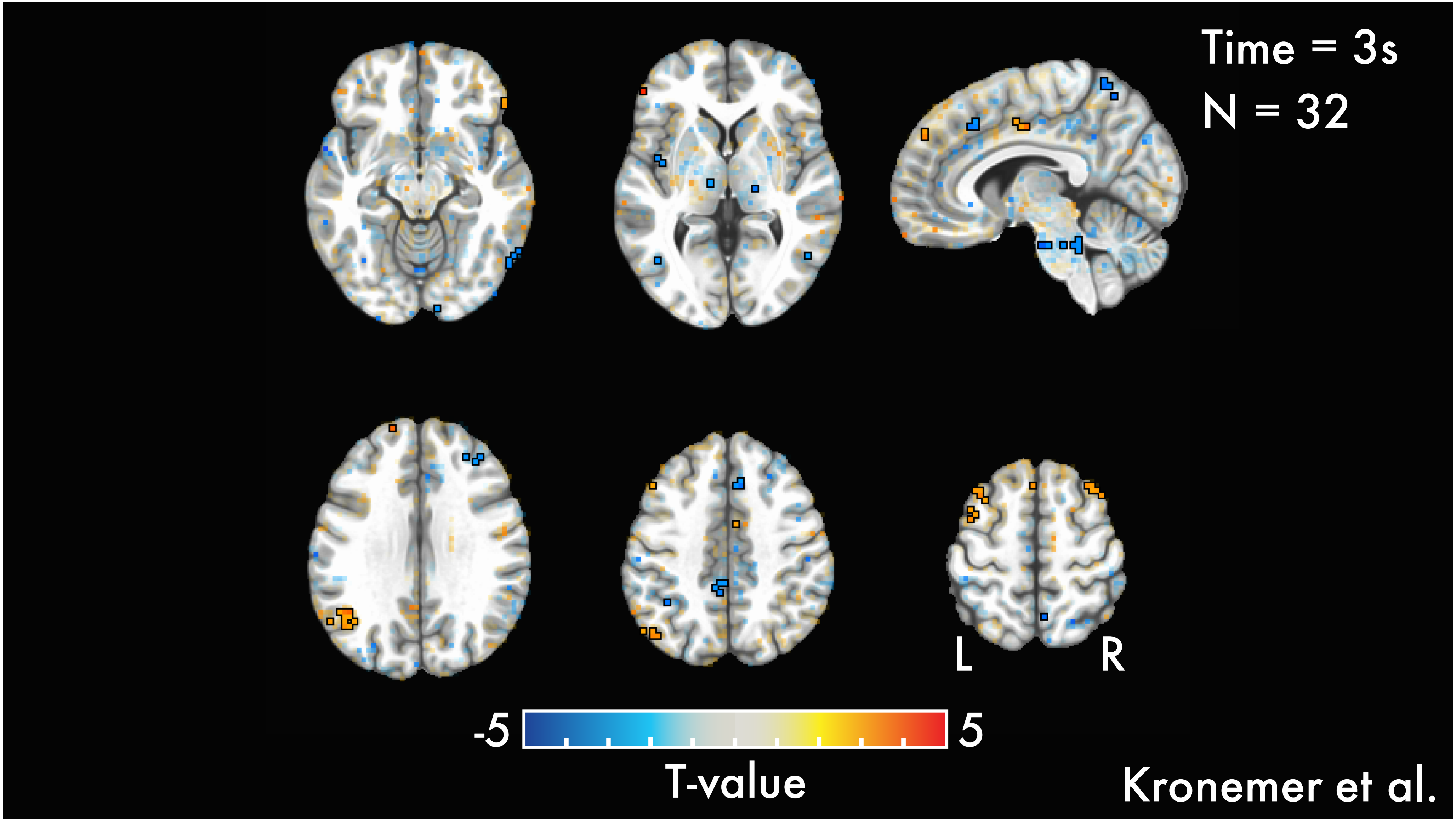

Time = 3s
N = 32
L
R
-5
5
T-value
Kronemer et al.

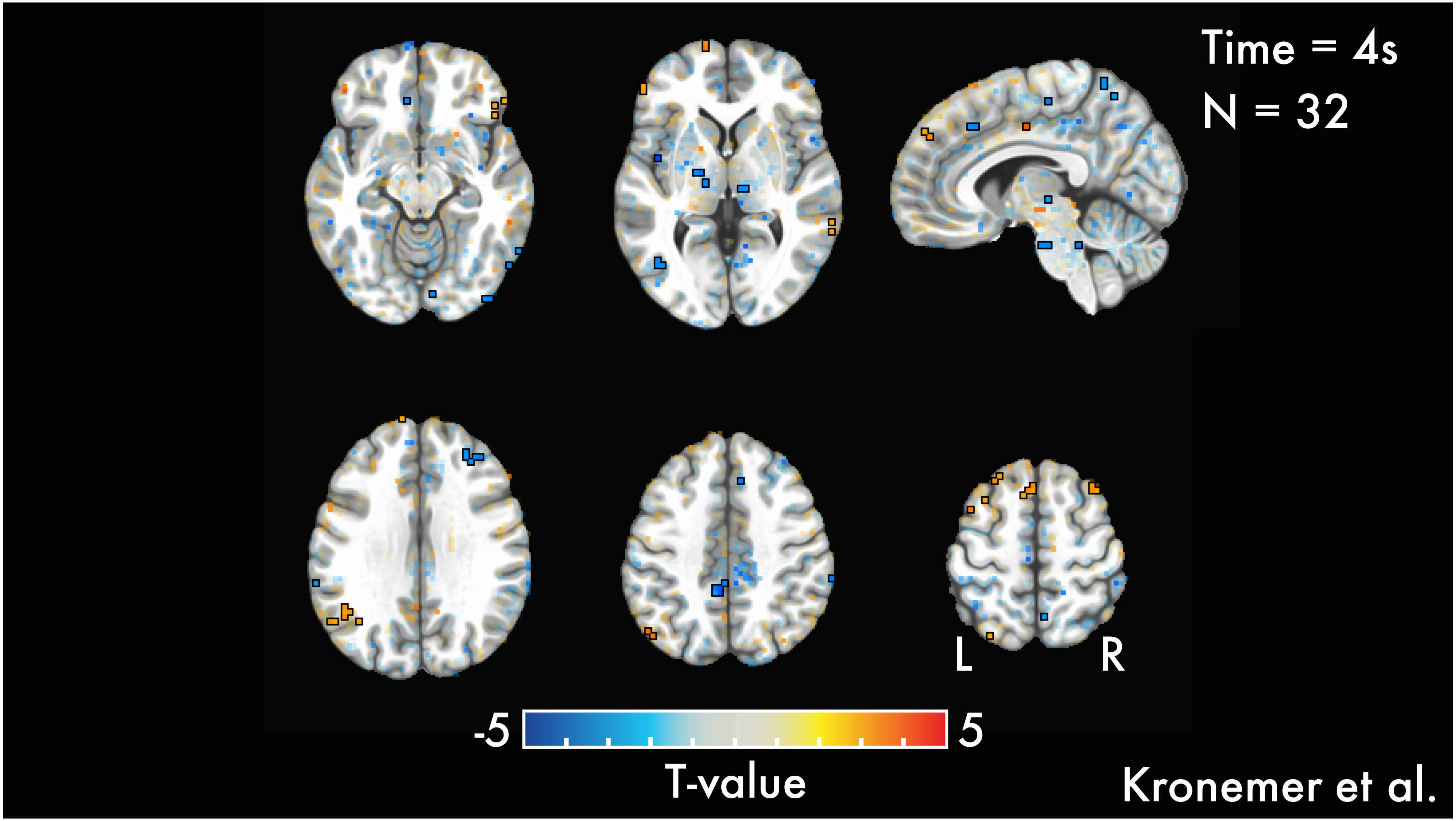
Time = 4s
N = 32
L
R
-5
5
T-value
Kronemer et al.

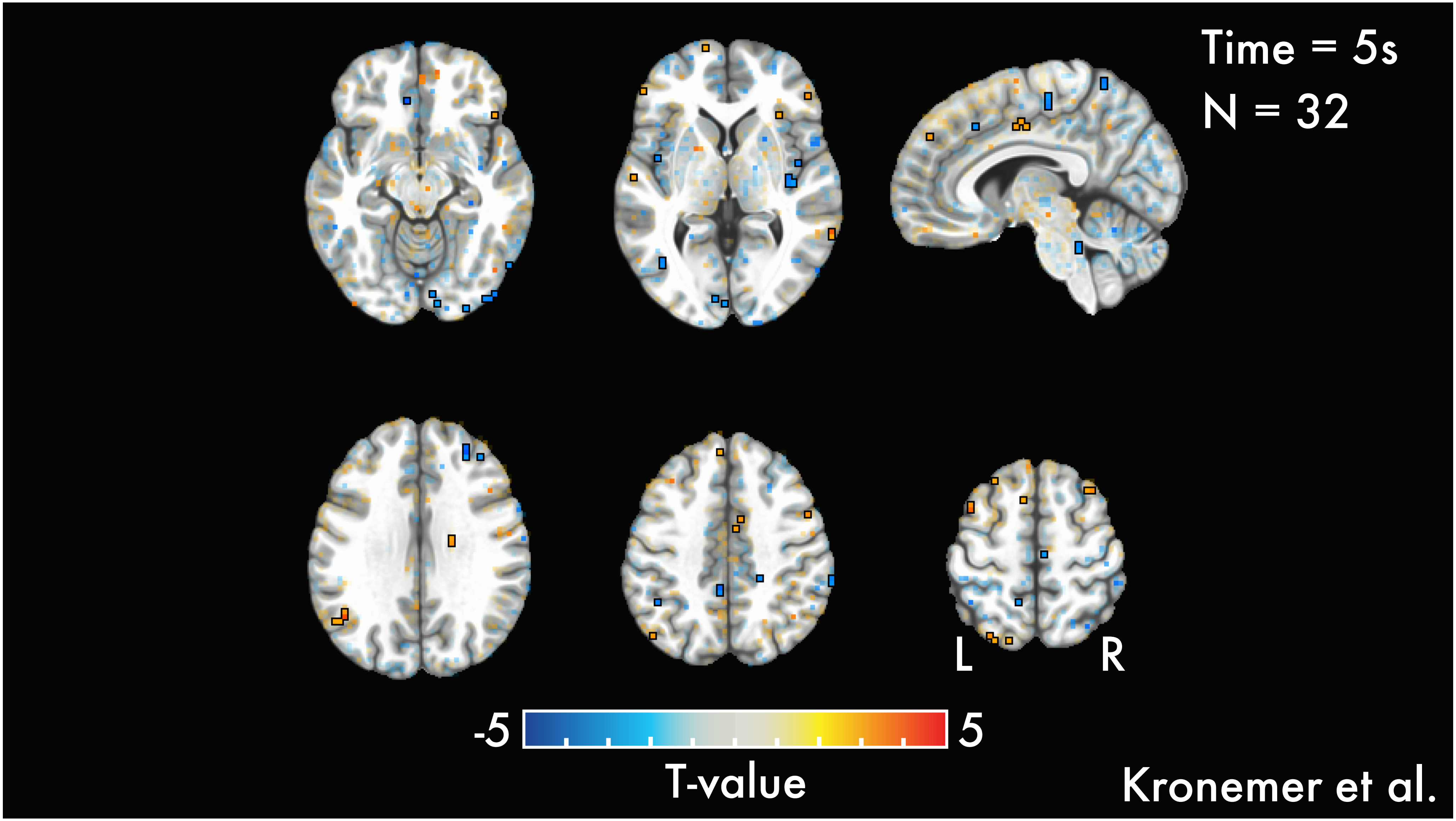
Time = 5s
N = 32
L
R
-5
5
T-value
Kronemer et al.

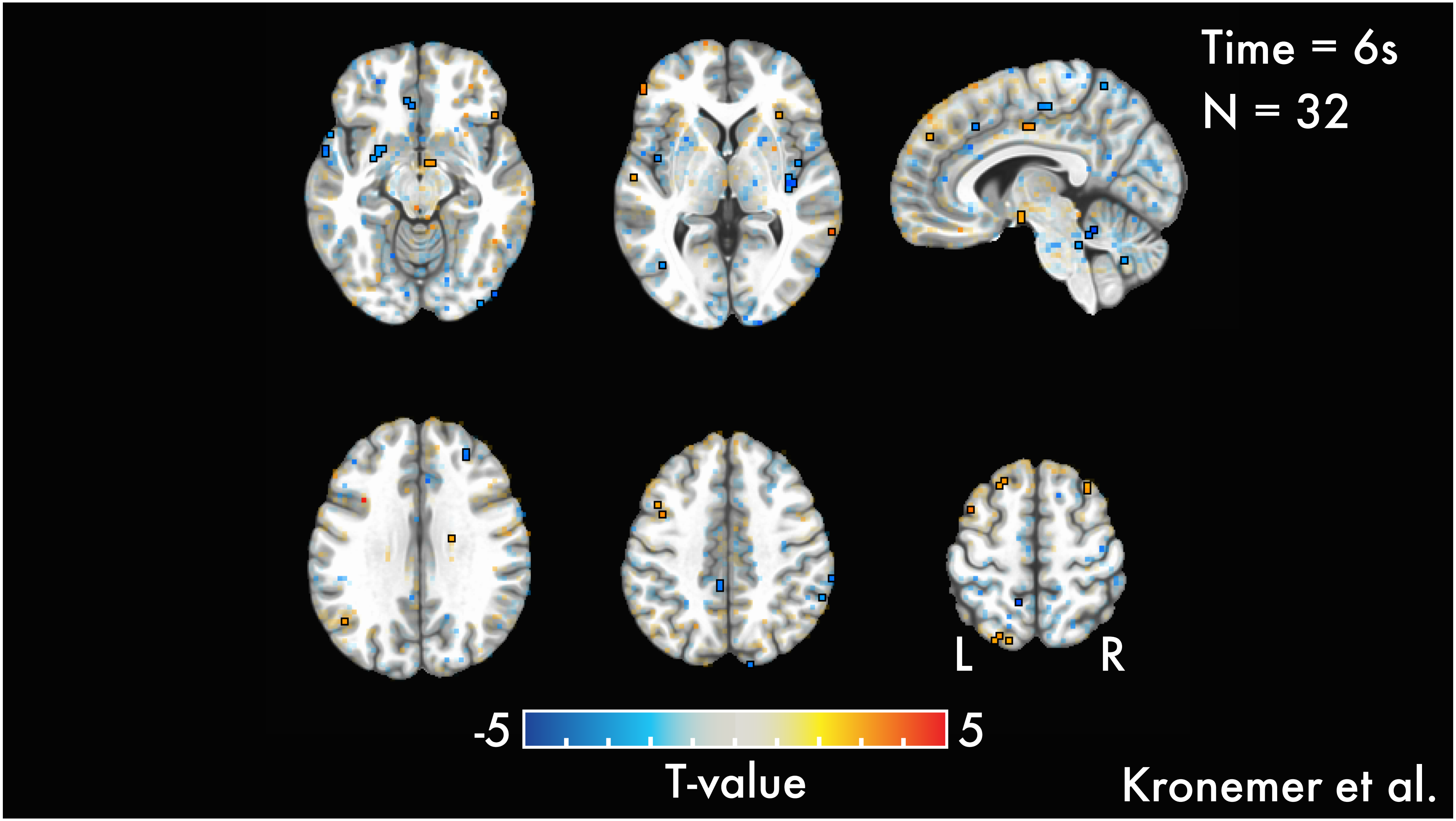
Time = 6s
N = 32
L
R
-5
5
T-value
Kronemer et al.

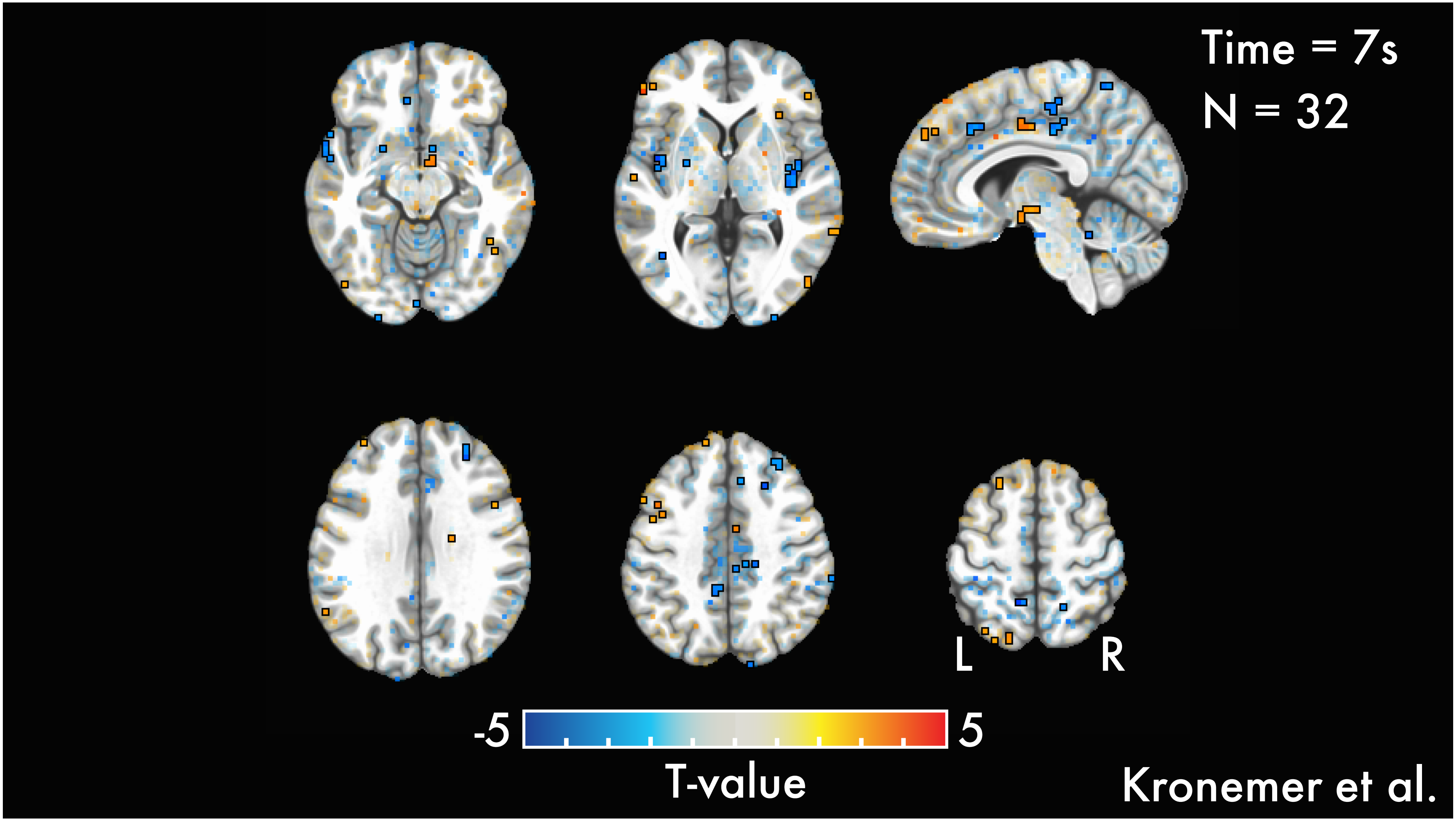

Time = 7s
N = 32
L
R
-5
5
T-value
Kronemer et al.

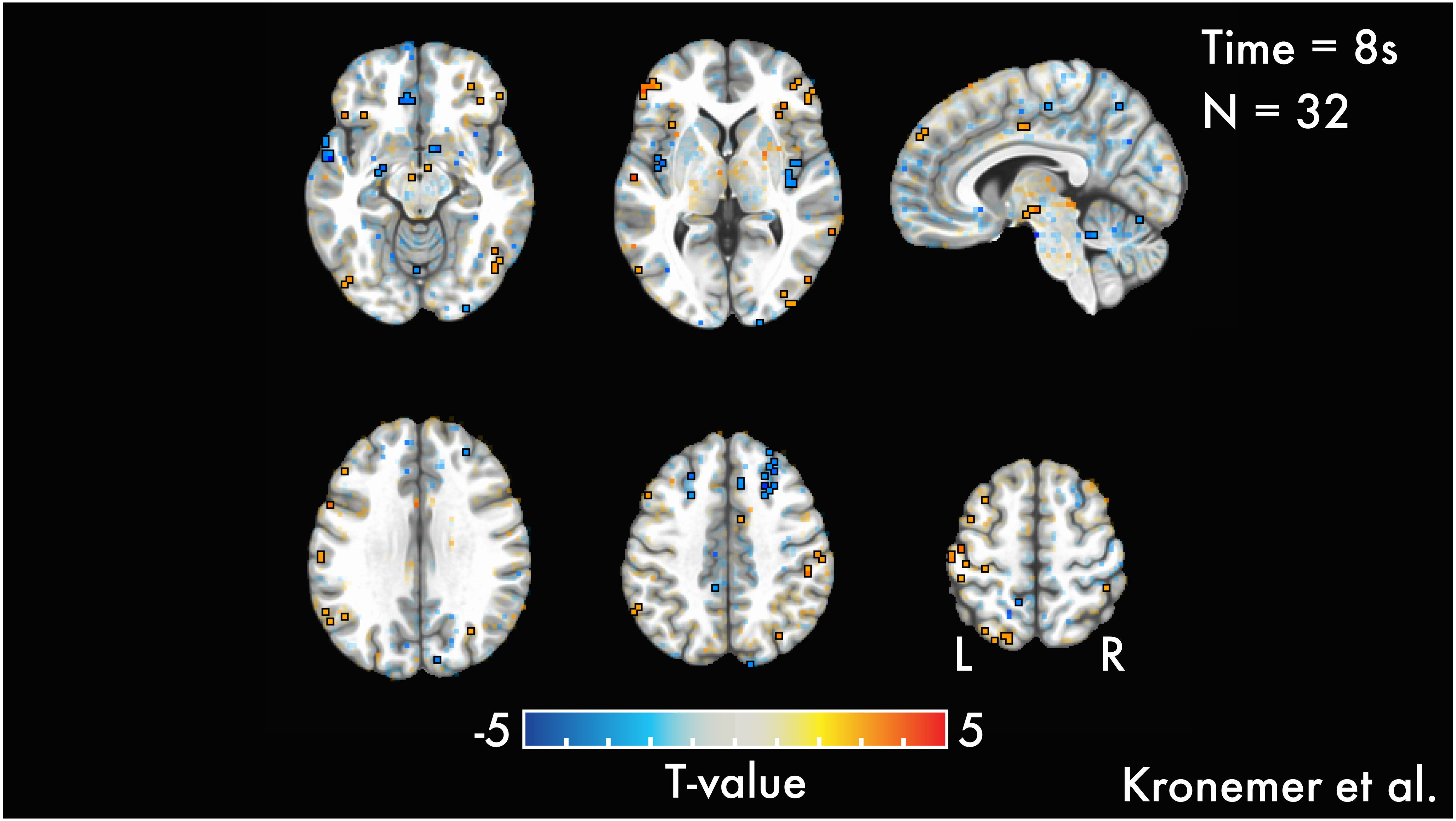
Time = 8s
N = 32
L
R
-5
5
T-value
Kronemer et al.

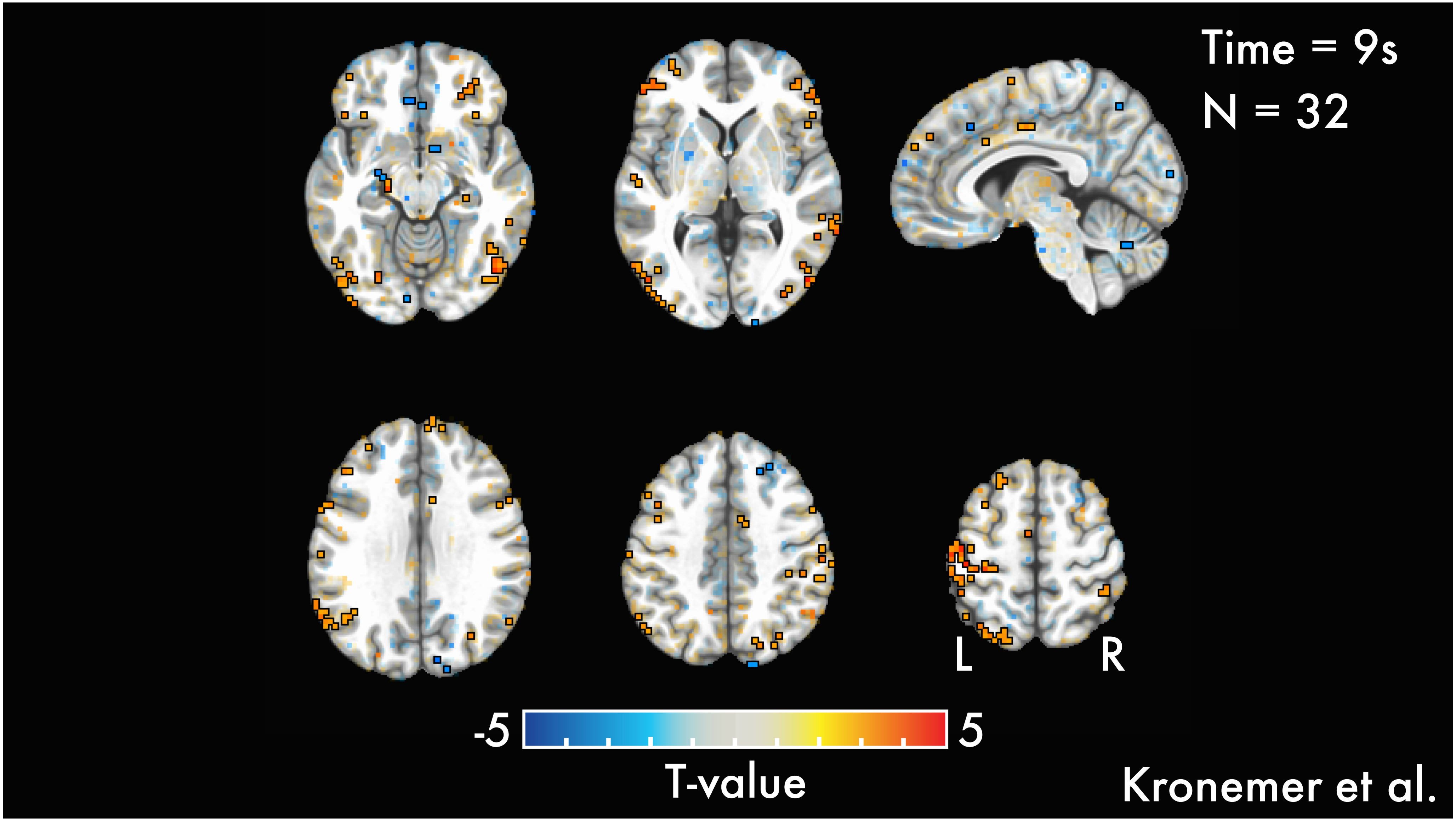

Time = 9s
N = 32
L
R
-5
5
T-value
Kronemer et al.

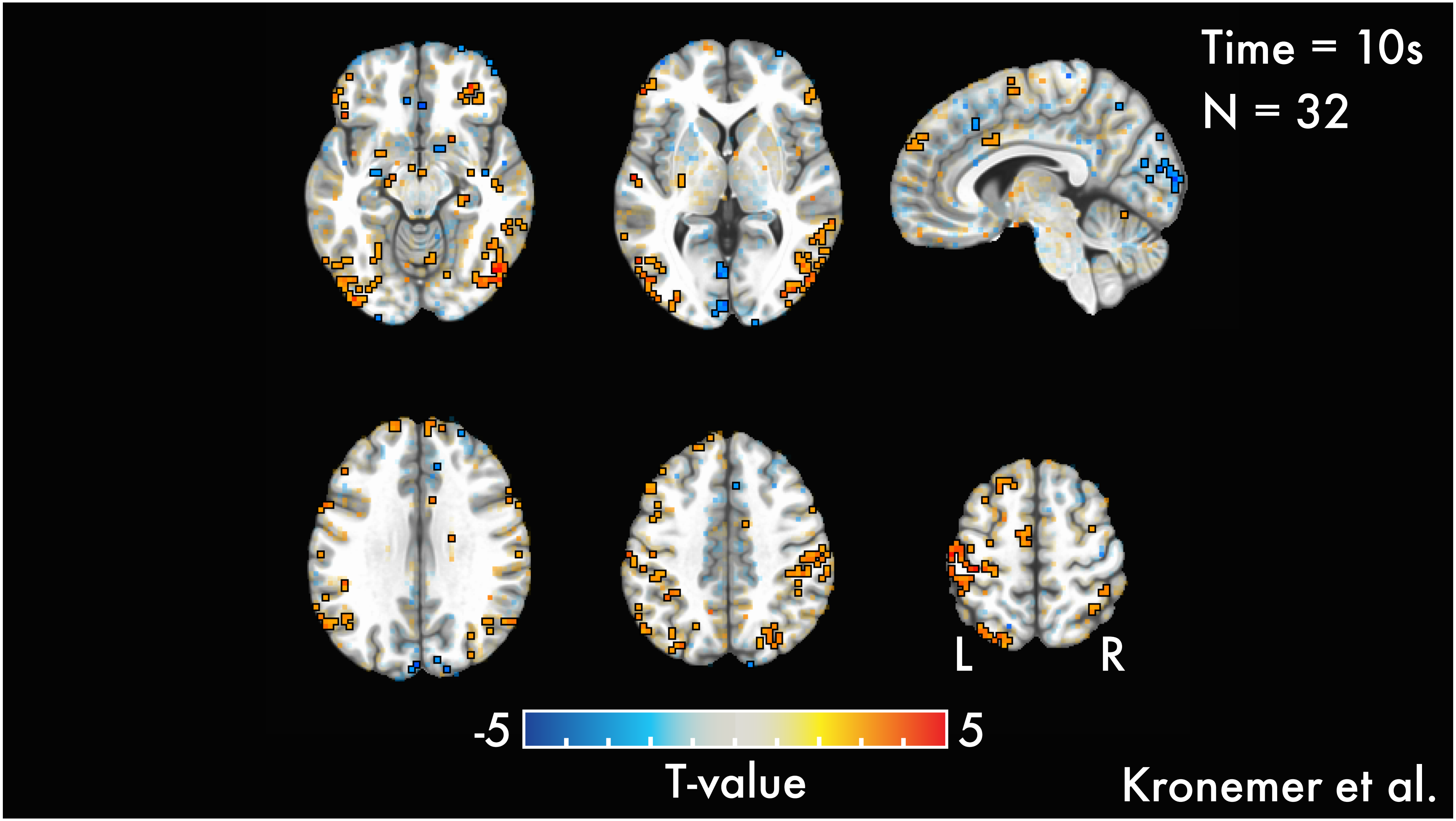
Time = 10s
N = 32
L
R
-5
5
T-value
Kronemer et al.

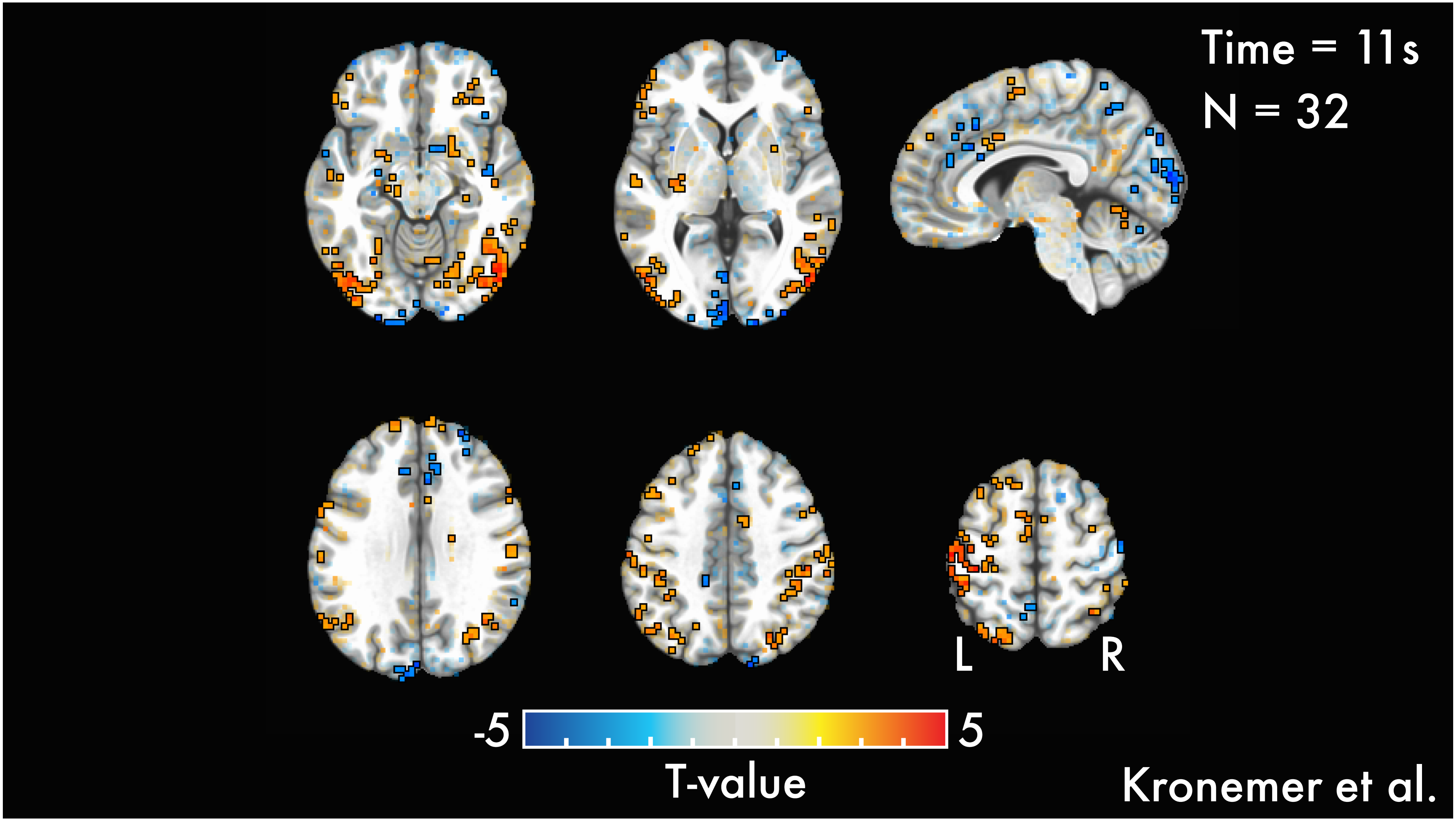

Time = 11s
N = 32
L
R
-5
5
T-value
Kronemer et al.

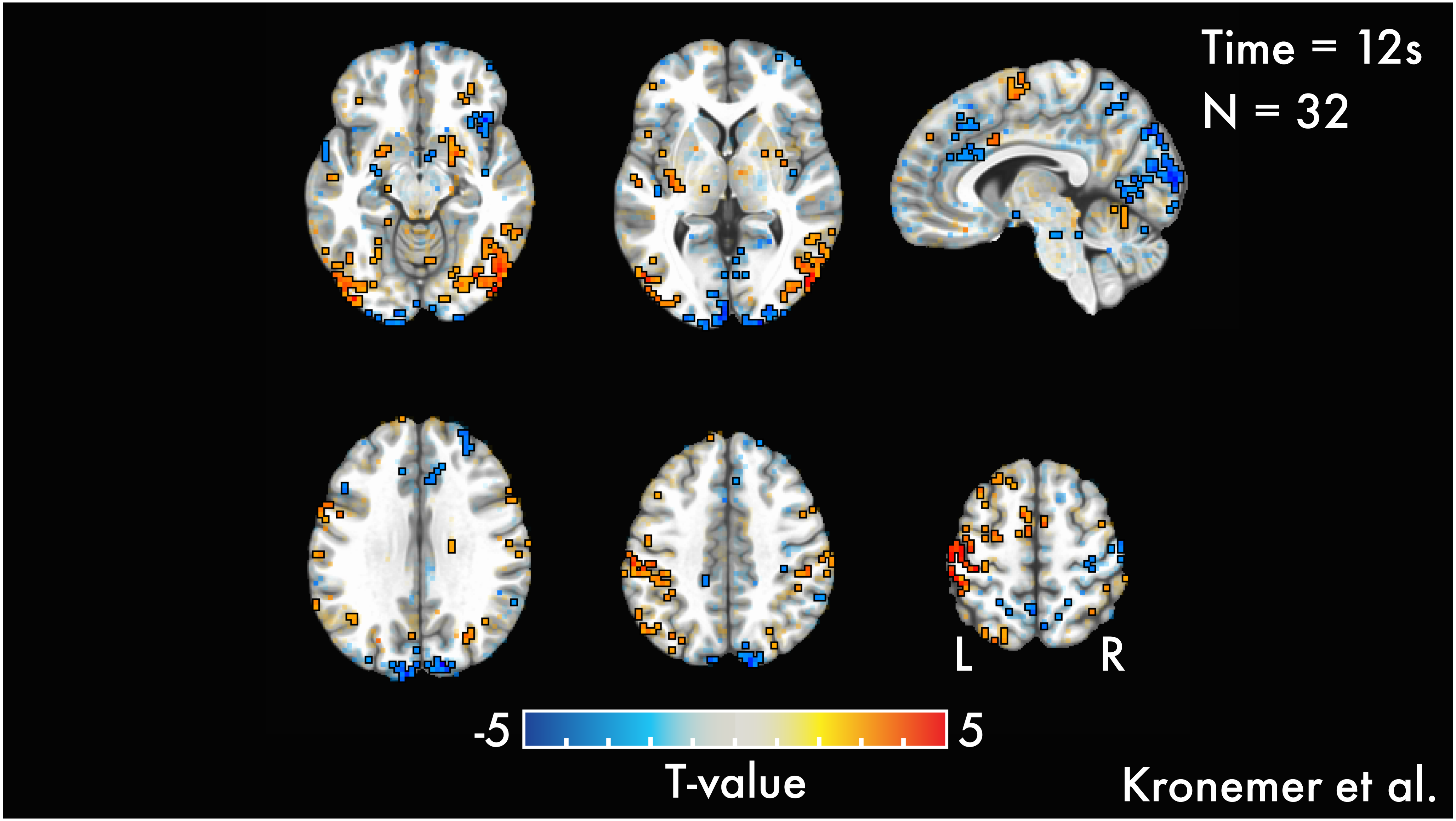

Time = 12s
N = 32
L
R
-5
5
T-value
Kronemer et al.

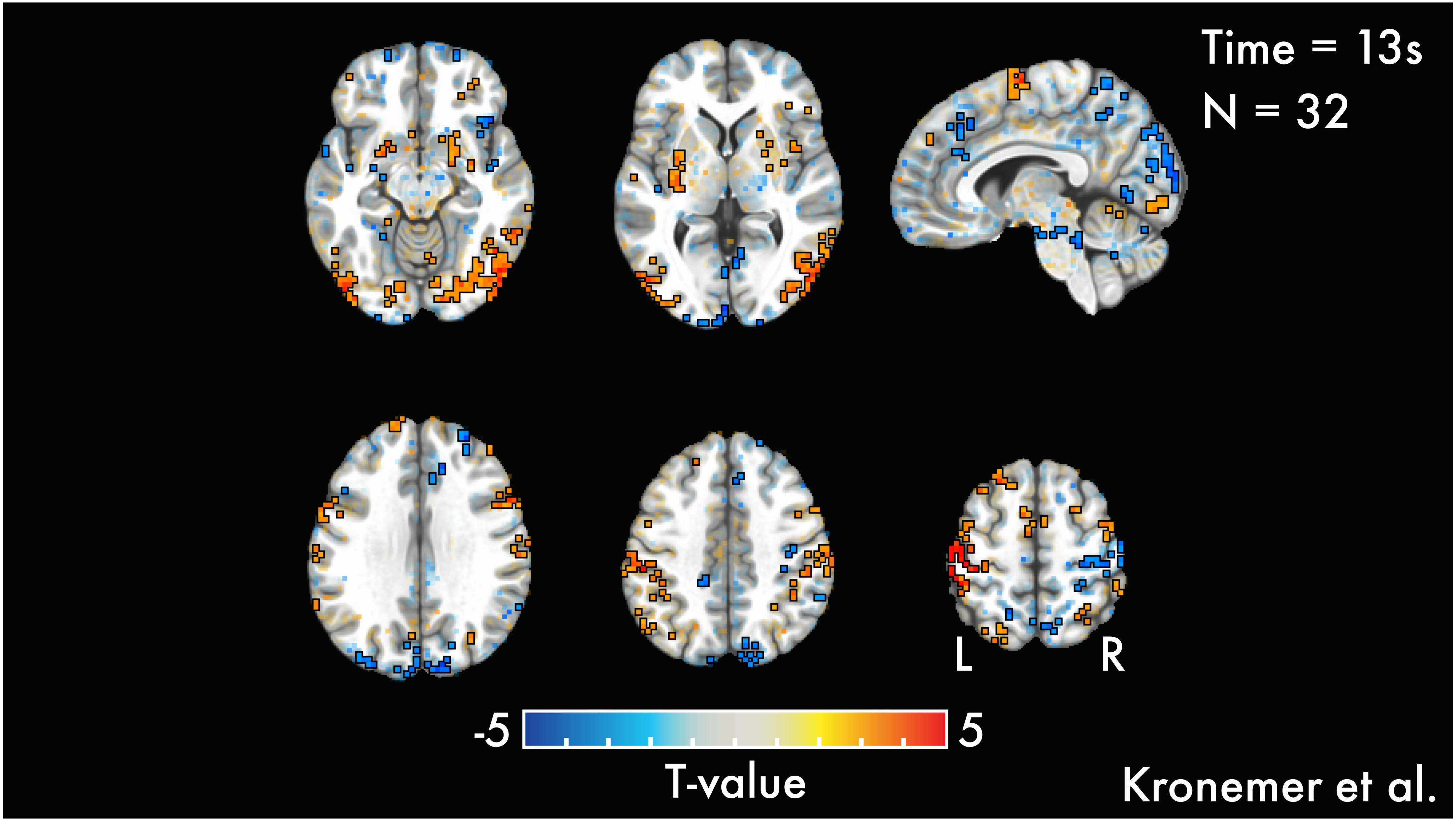

Time = 13s
N = 32
L
R
-5
5
T-value
Kronemer et al.

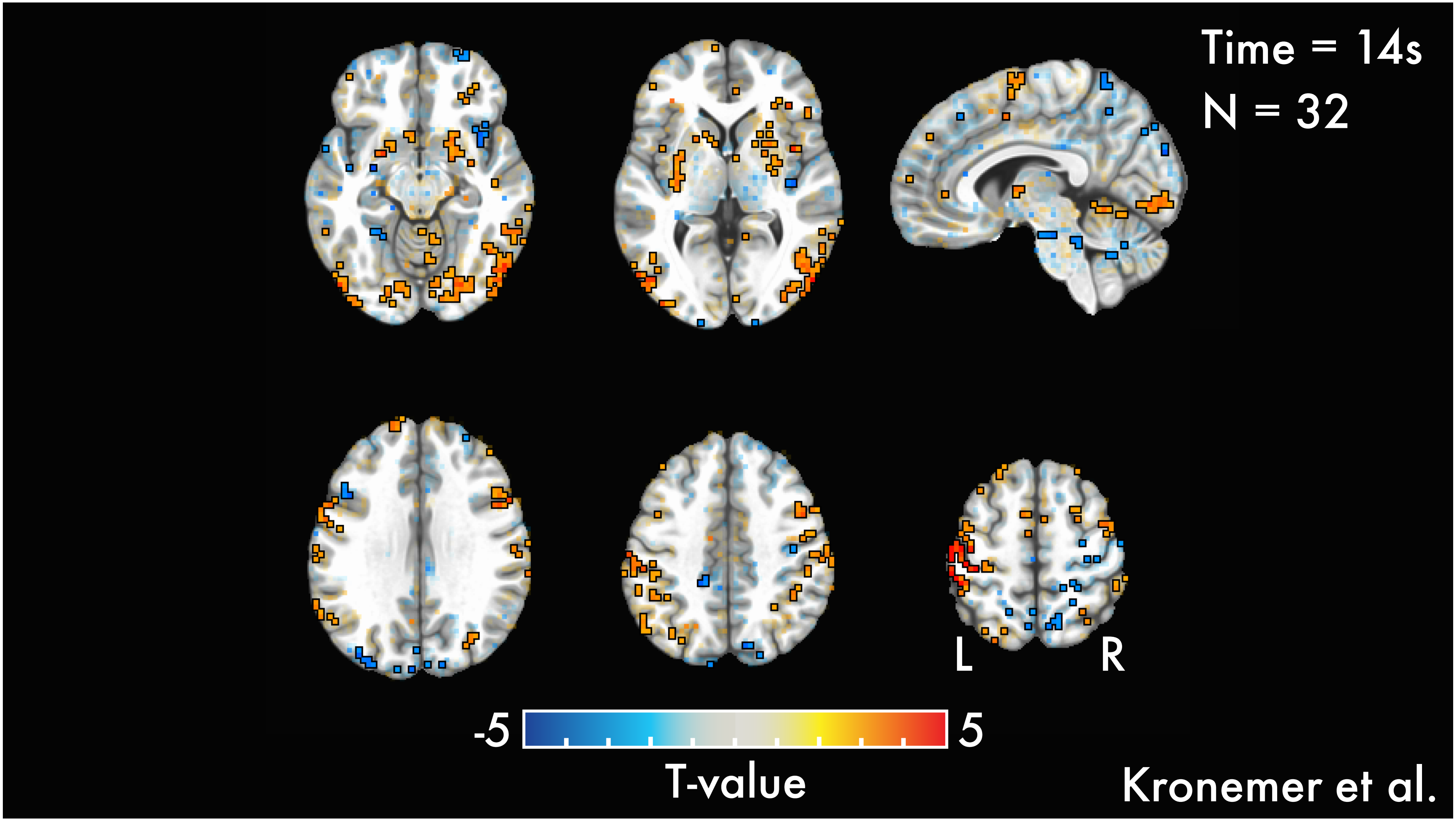
Time = 14s
N = 32
L
R
-5
5
T-value
Kronemer et al.

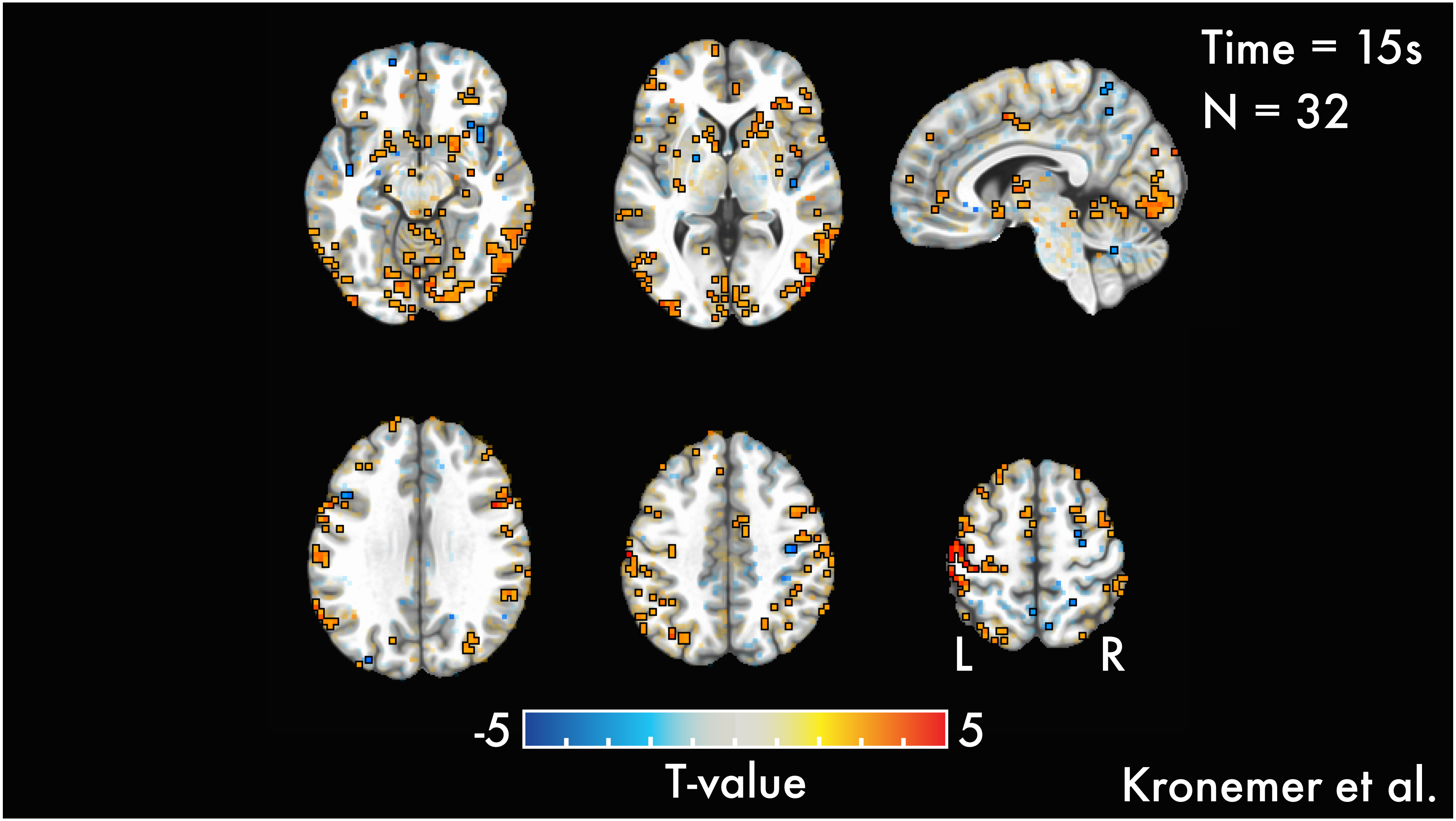
Time = 15s
N = 32
L
R
-5
5
T-value
Kronemer et al.

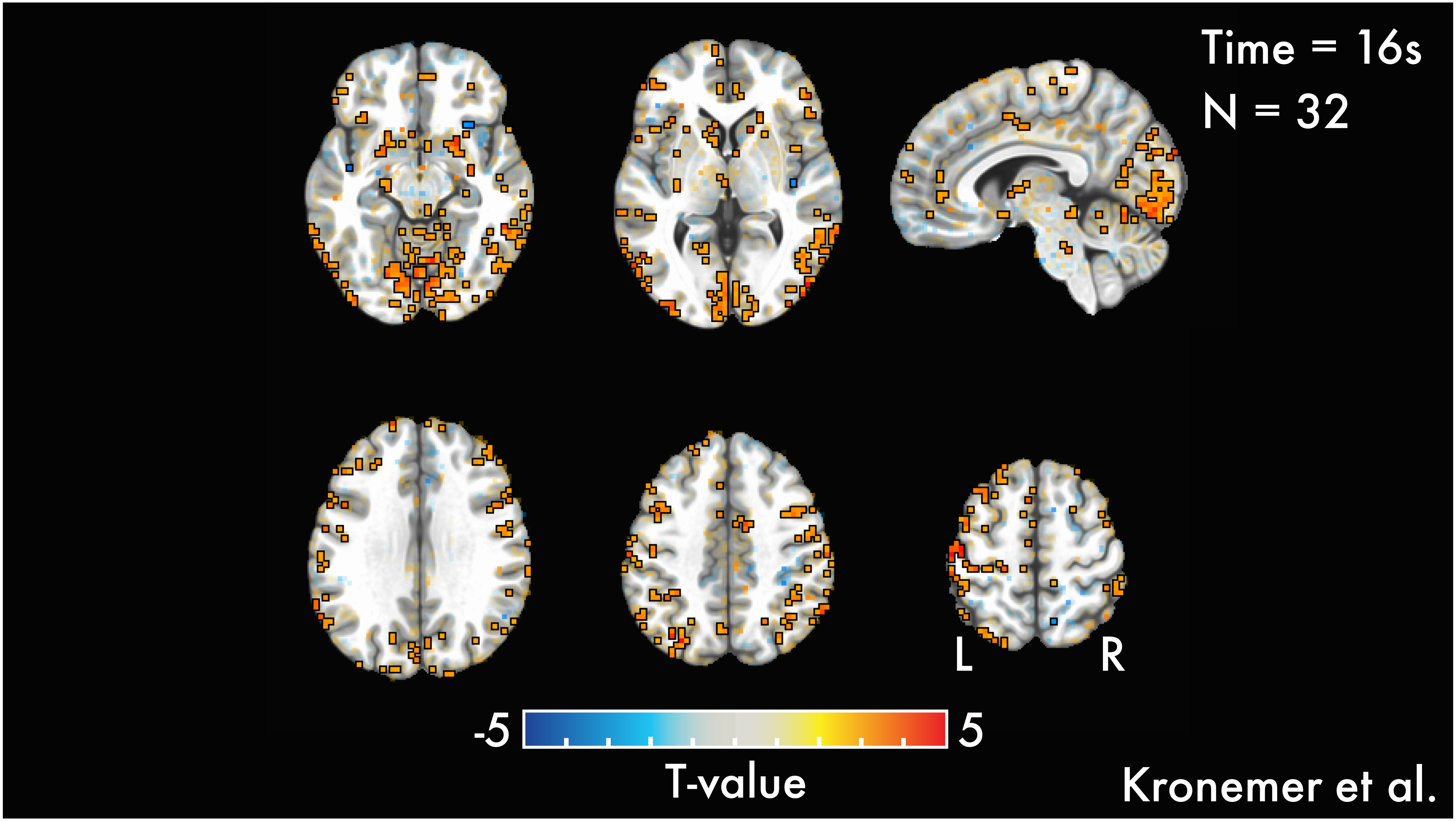
Time = 16s
N = 32
L
R
-5
5
T-value
Kronemer et al.

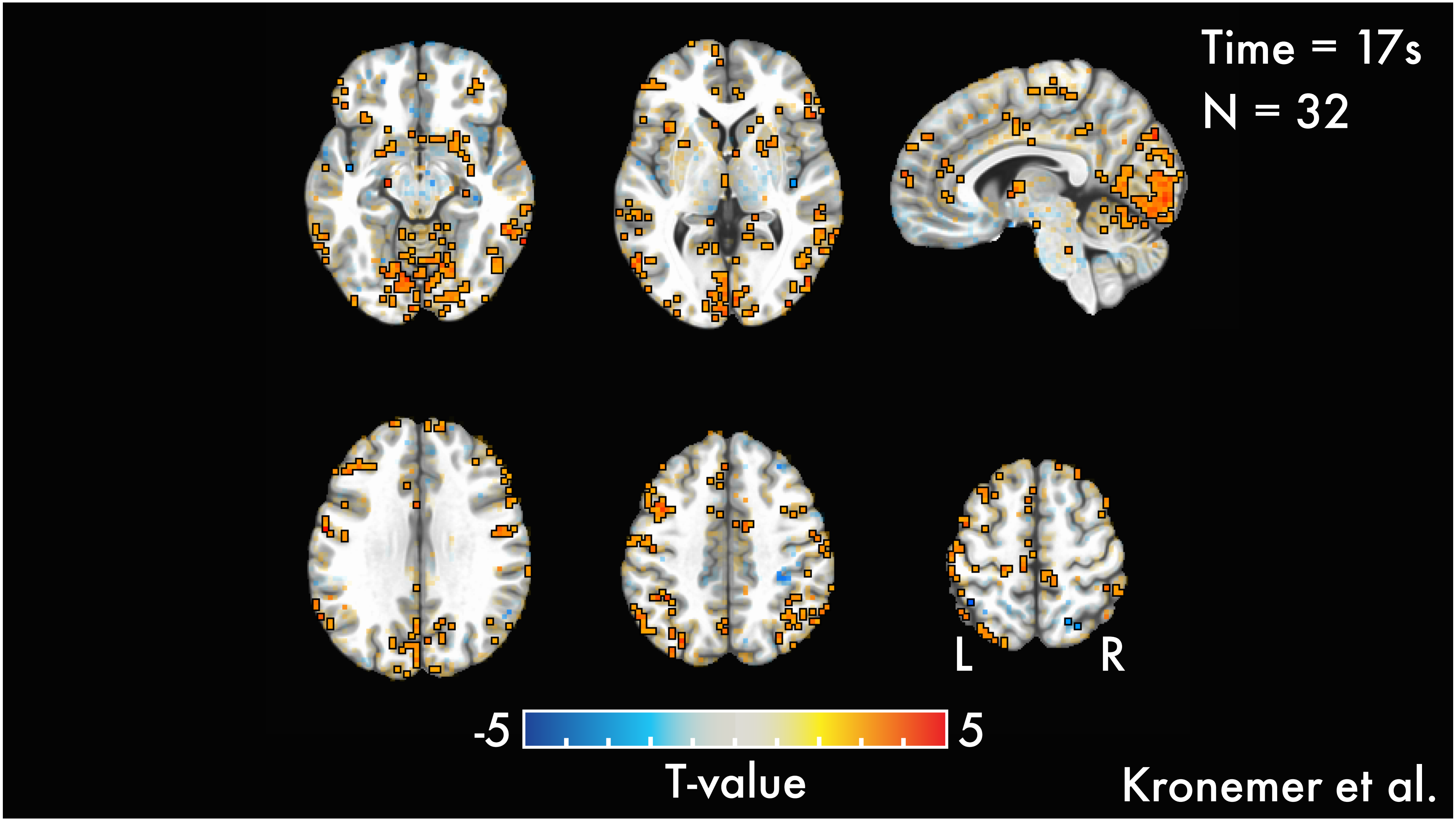

Time = 17s
N = 32
L
R
-5
5
T-value
Kronemer et al.

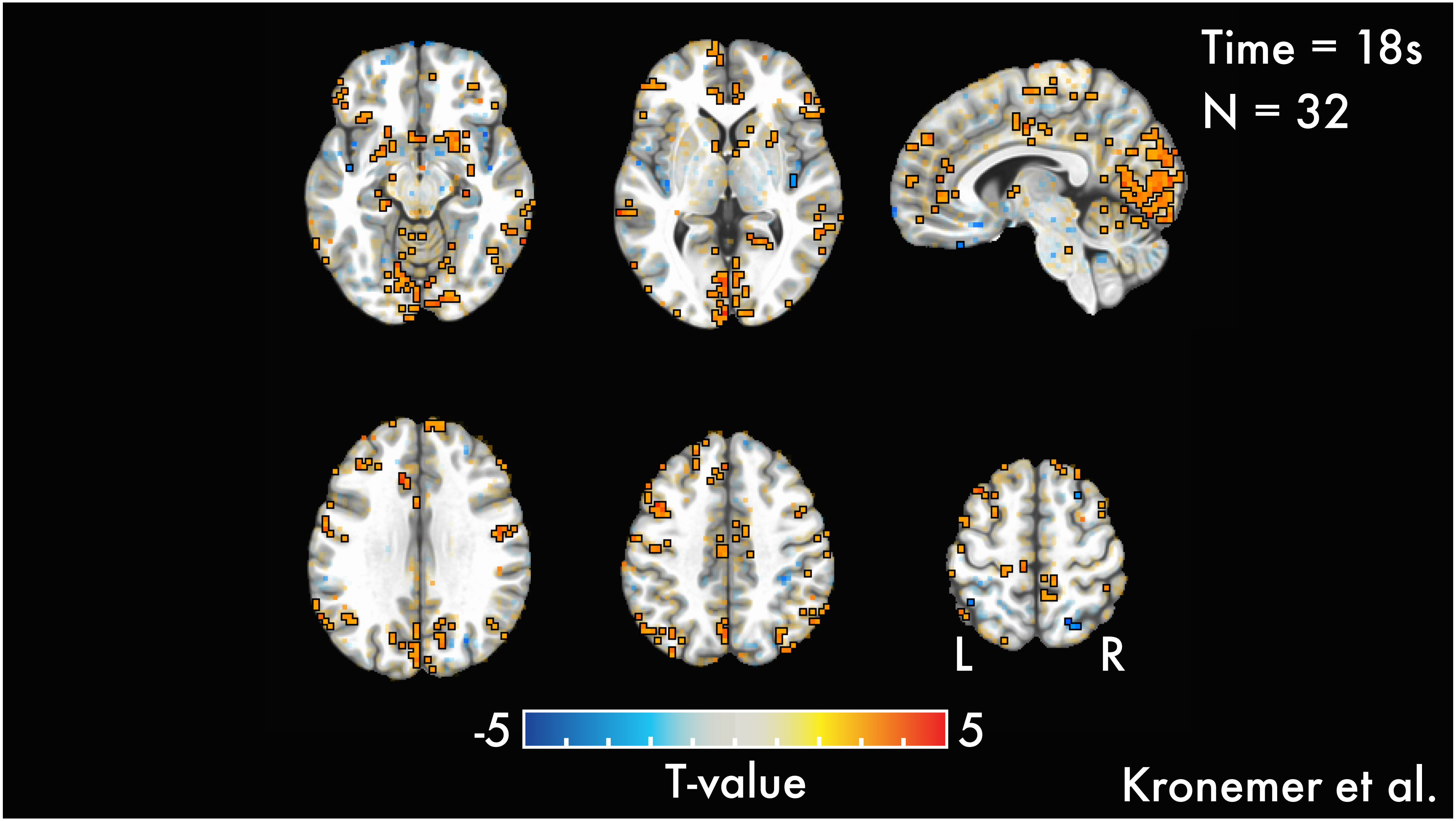

Time = 18s
N = 32
L
R
-5
5
T-value
Kronemer et al.

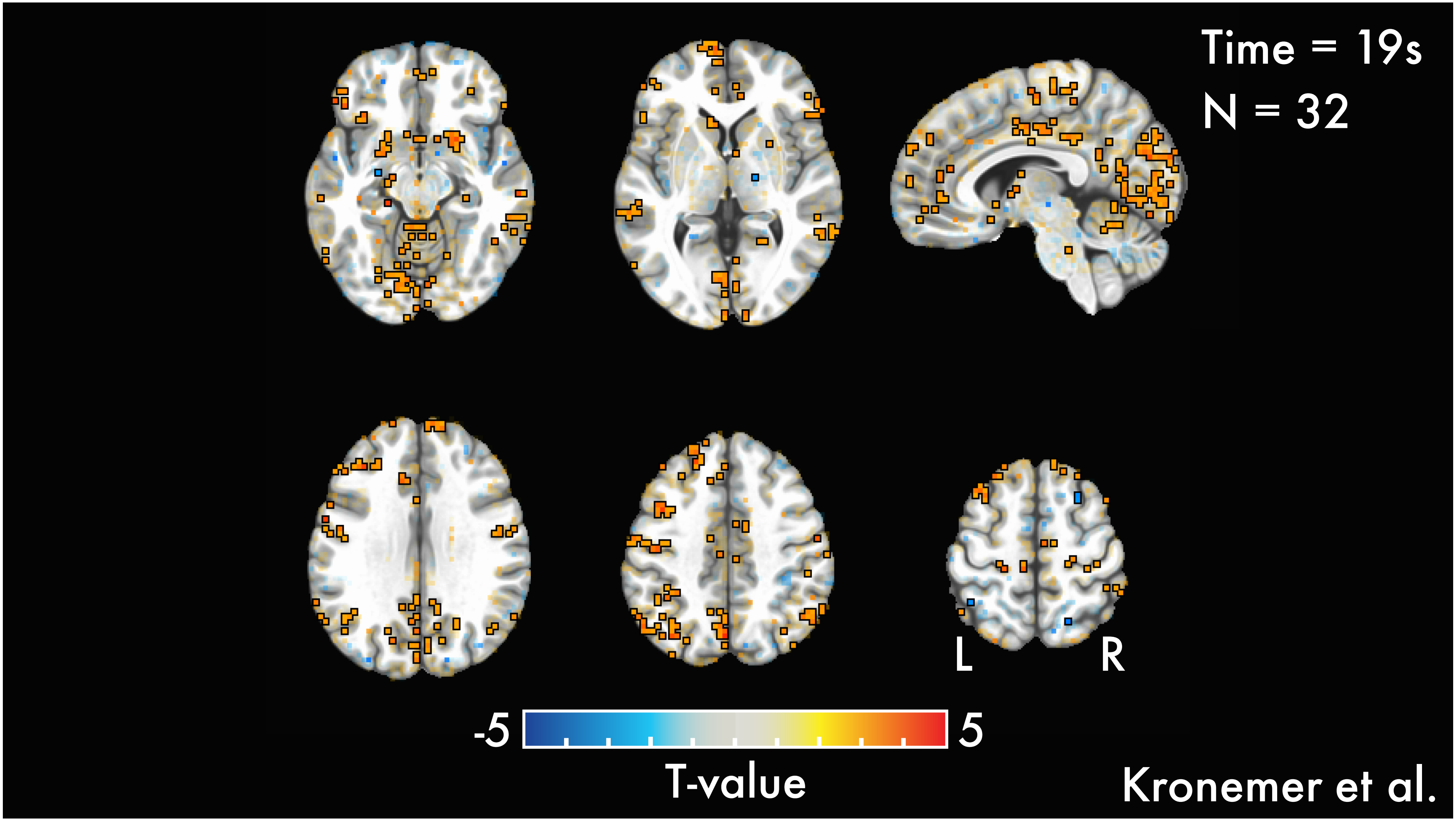
Time = 19s
N = 32
L
R
-5
5
T-value
Kronemer et al.

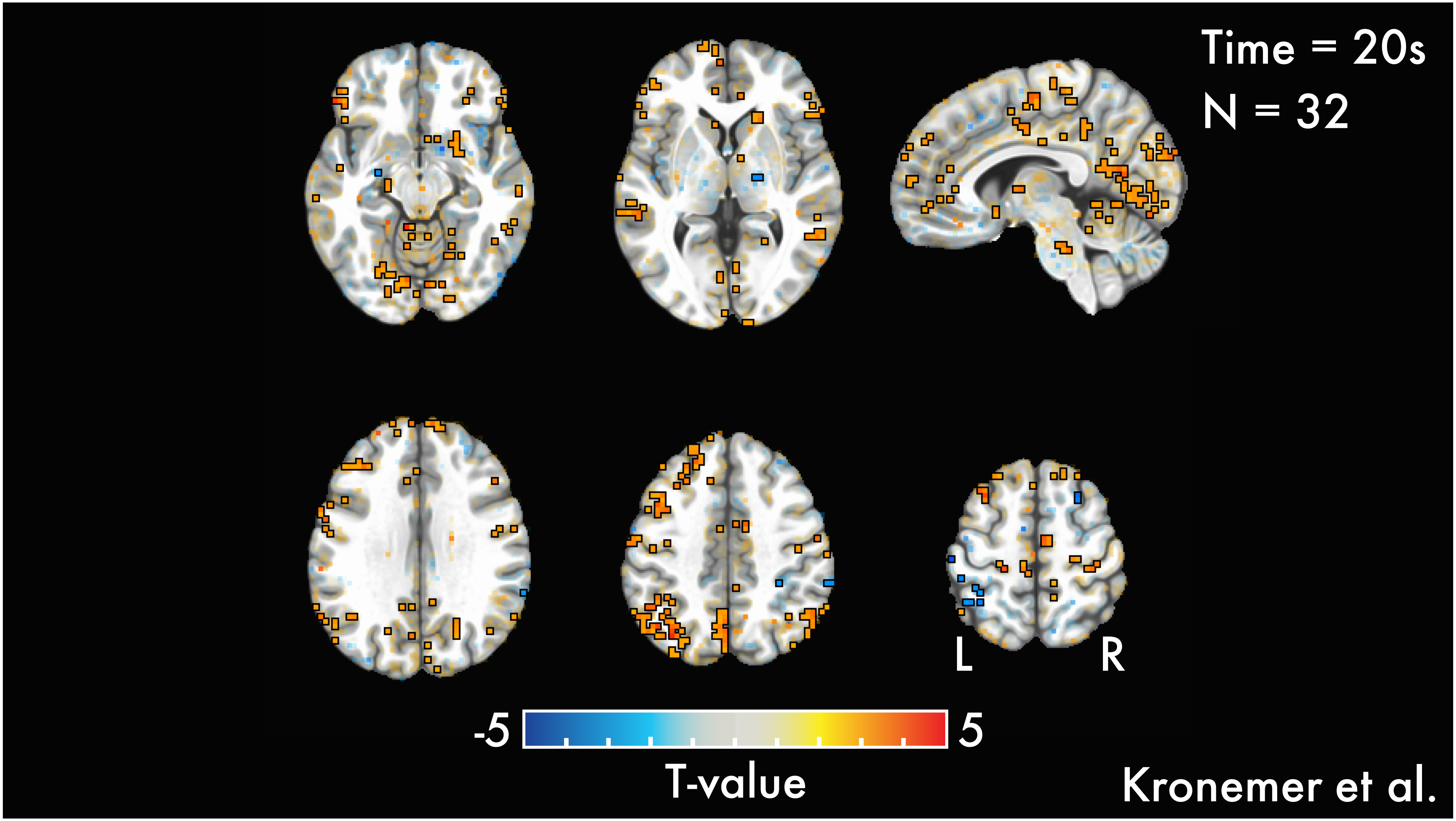

Time = 20s
N = 32
L
R
-5
5
T-value
Kronemer et al.

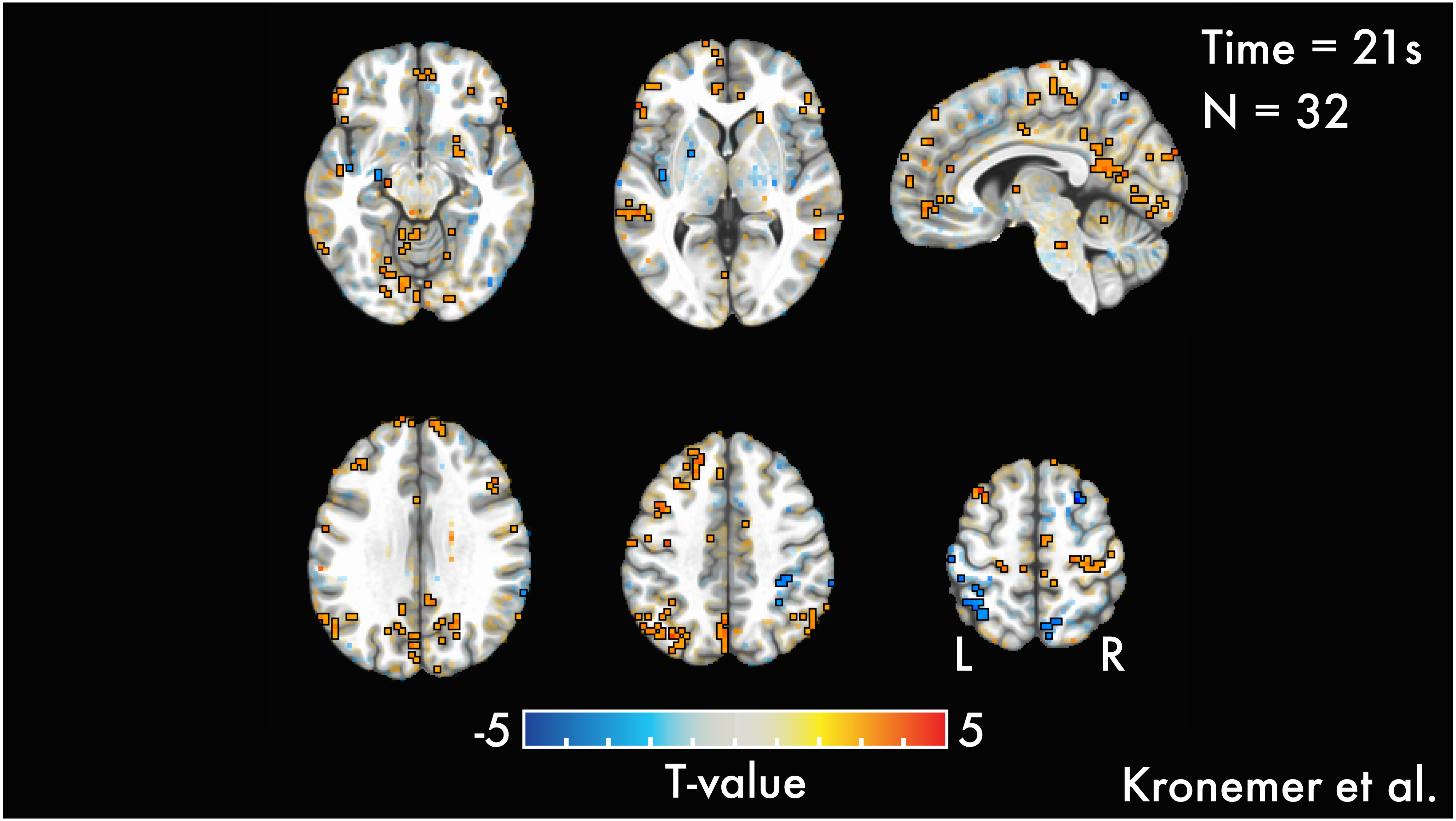

Time = 21s
N = 32
L
R
-5
5
T-value
Kronemer et al.

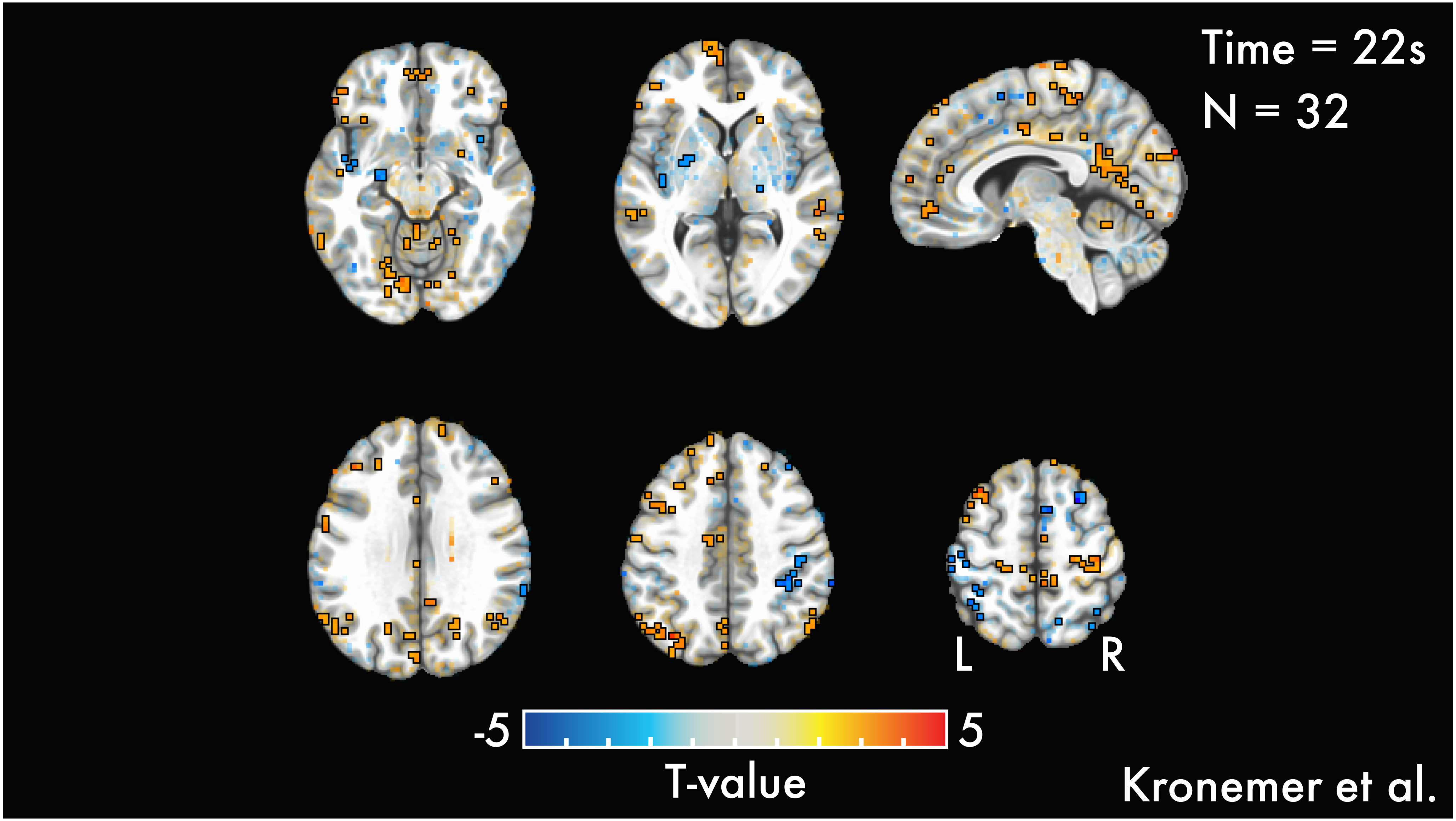
Time = 22s
N = 32
L
R
-5
5
T-value
Kronemer et al.

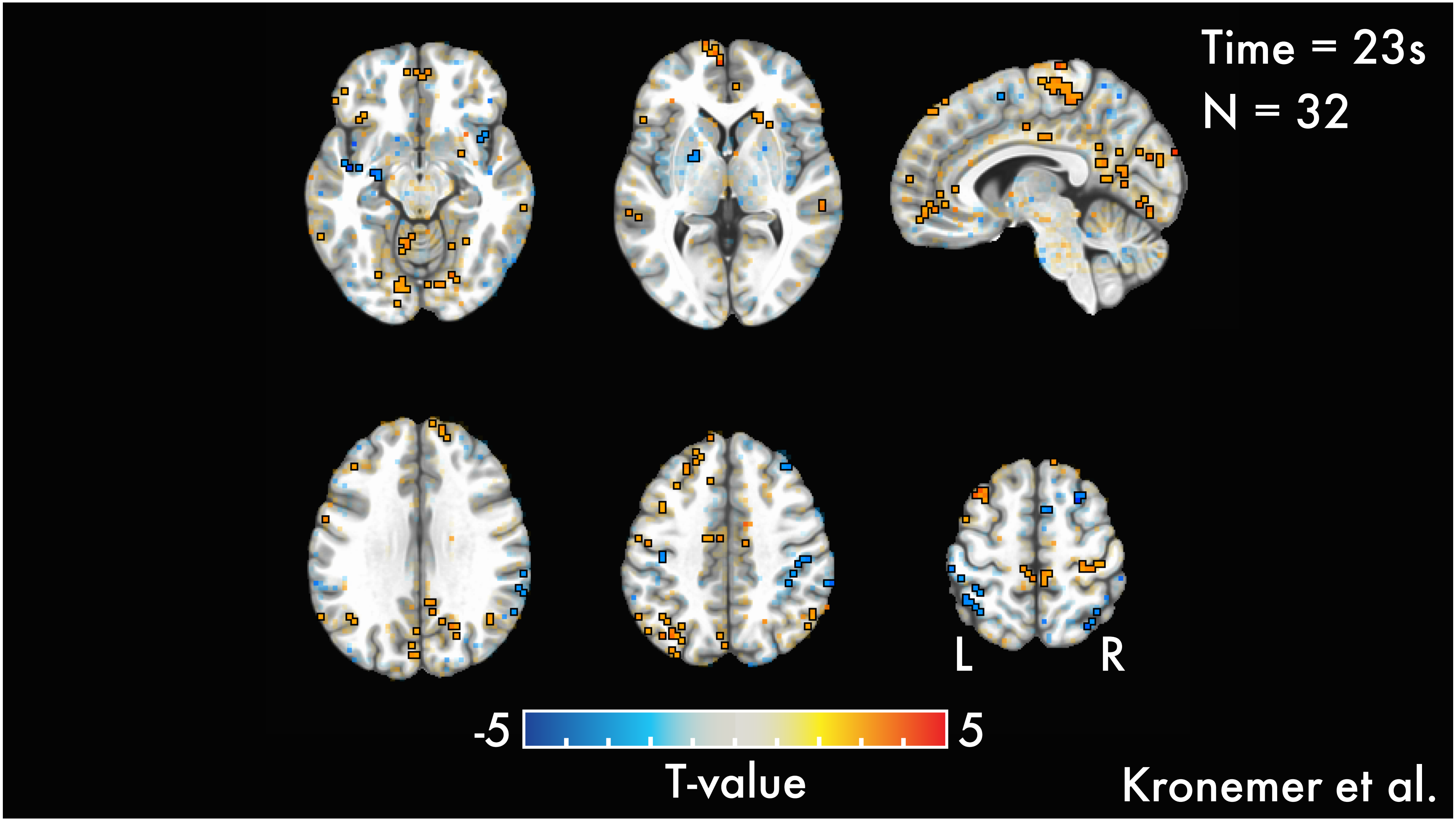
Time = 23s
N = 32
L
R
-5
5
T-value
Kronemer et al.

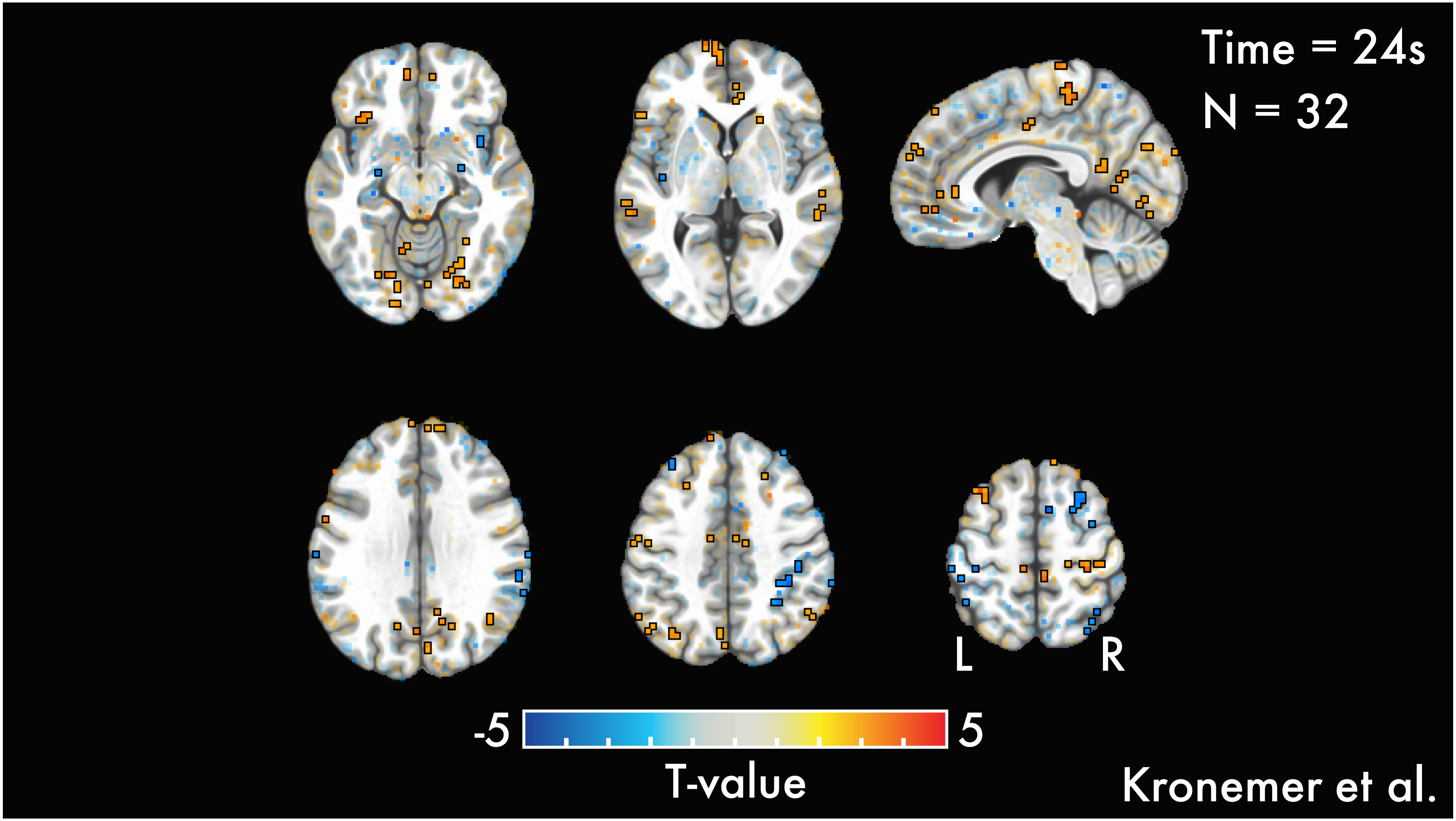
Time = 24s
N = 32
L
R
-5
5
T-value
Kronemer et al.

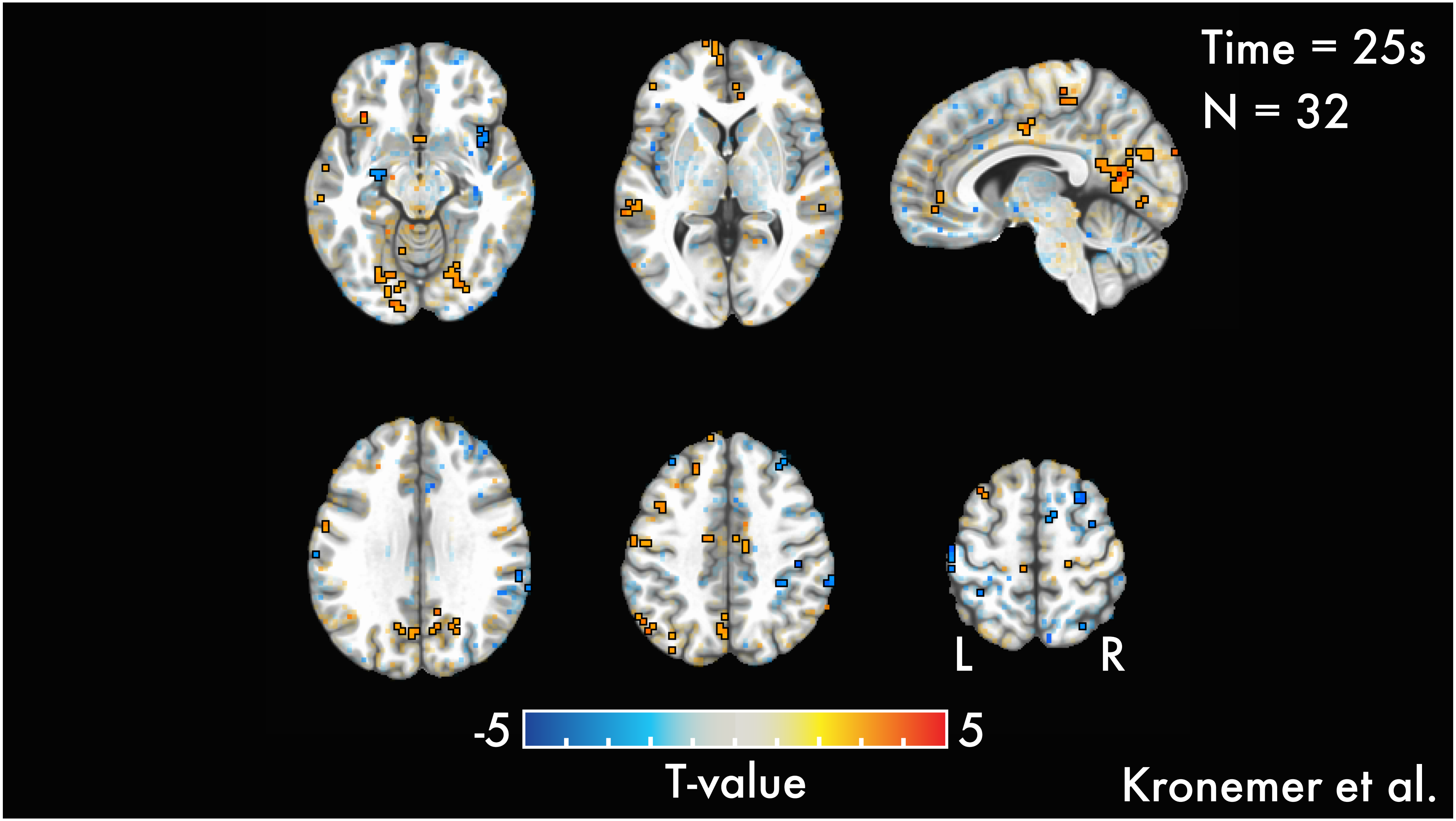
Time = 25s
N = 32
L
R
-5
5
T-value
Kronemer et al.
