## Supplementary Slides 5 for "The human brain mechanisms of afterimages: From networks to cortical layers"

### *Mock Inducers with versus without Mock Afterimages*

*Kronemer et al., 2026*

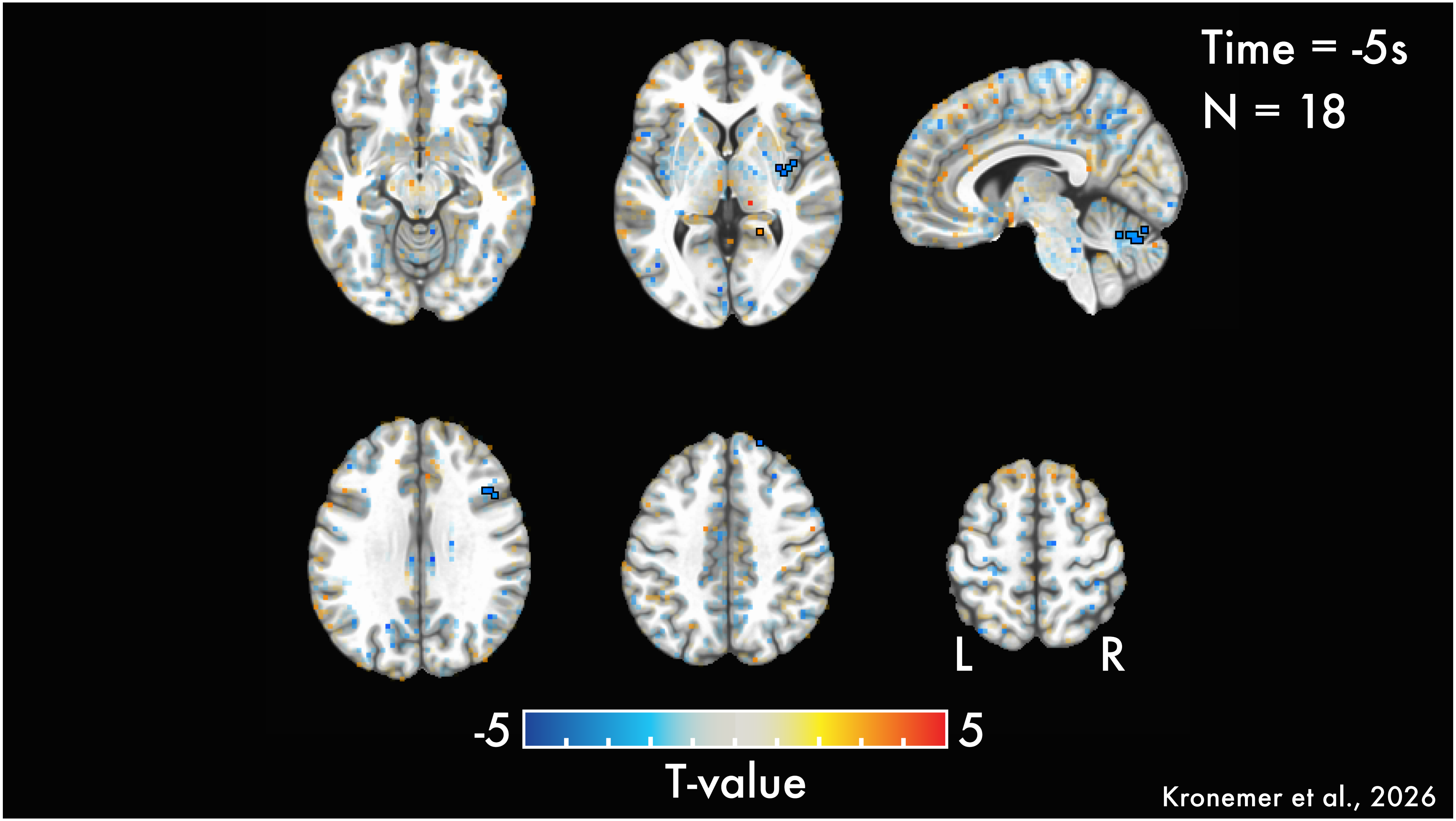
Time = -5s
N = 18
L
R
-5
5
T-value
Kronemer et al., 2026

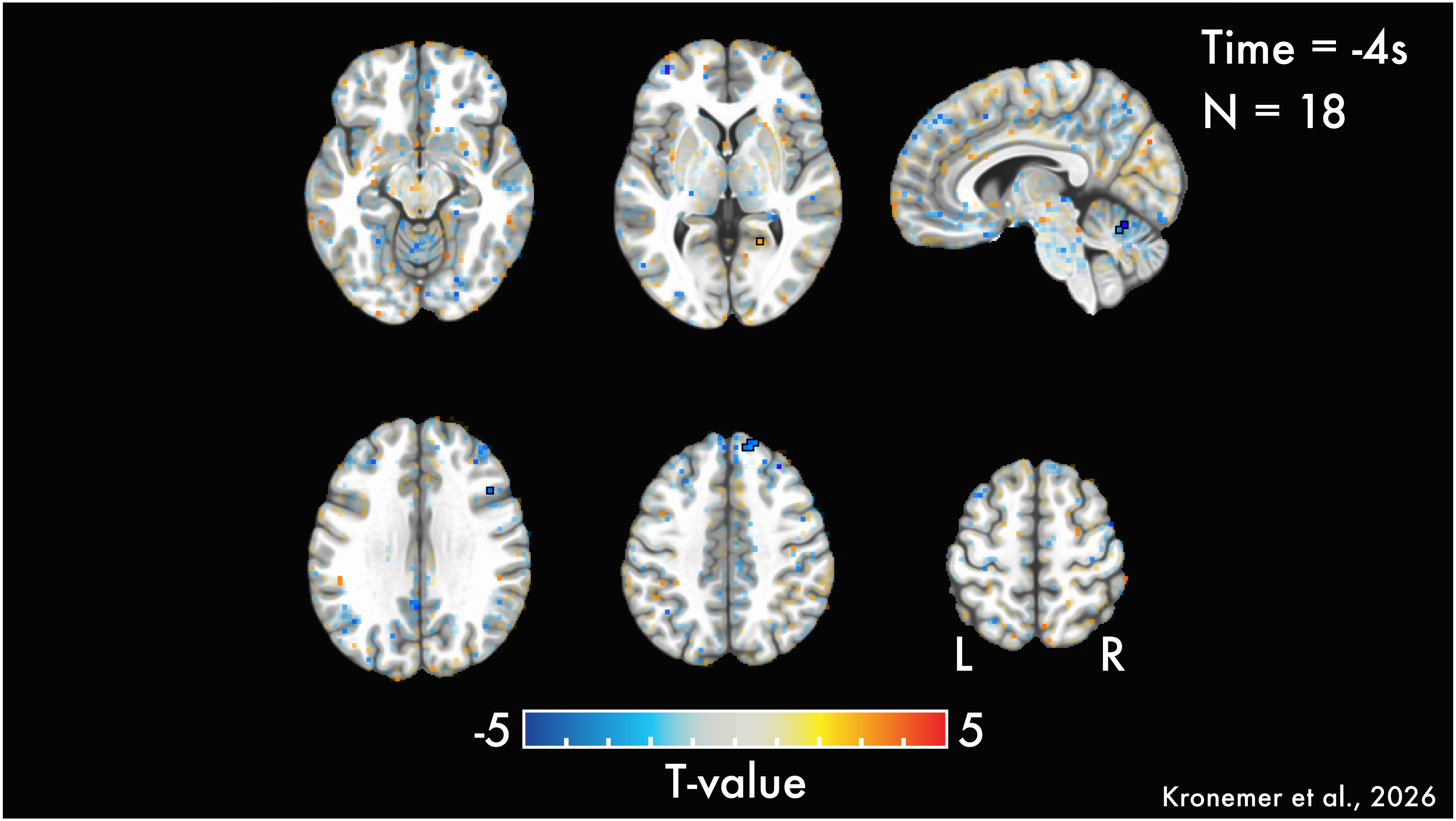

Time = -4s
N = 18
L
R
-5
5
T-value
Kronemer et al., 2026

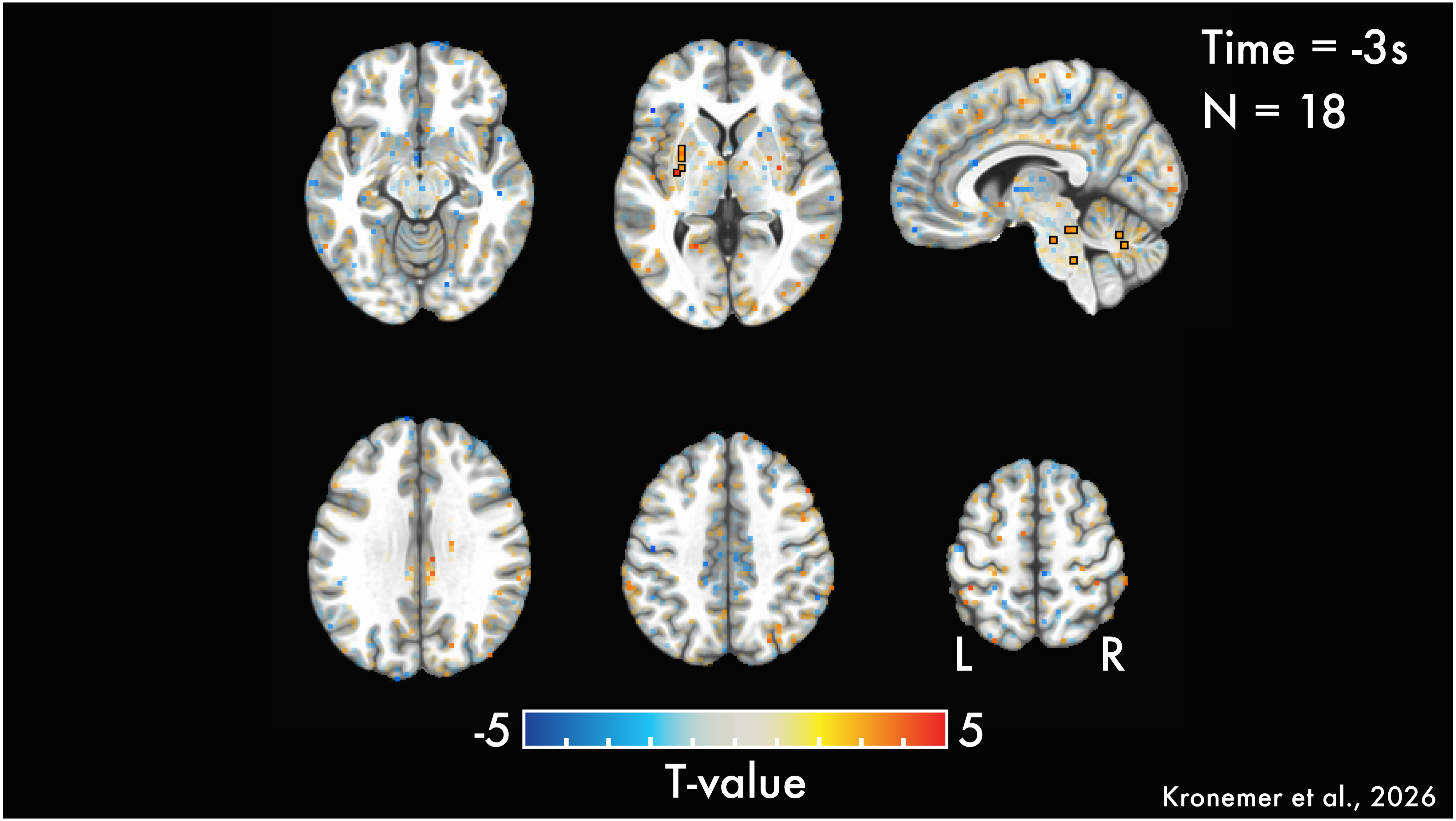
Time = -3s
N = 18
L
R
-5
5
T-value
Kronemer et al., 2026

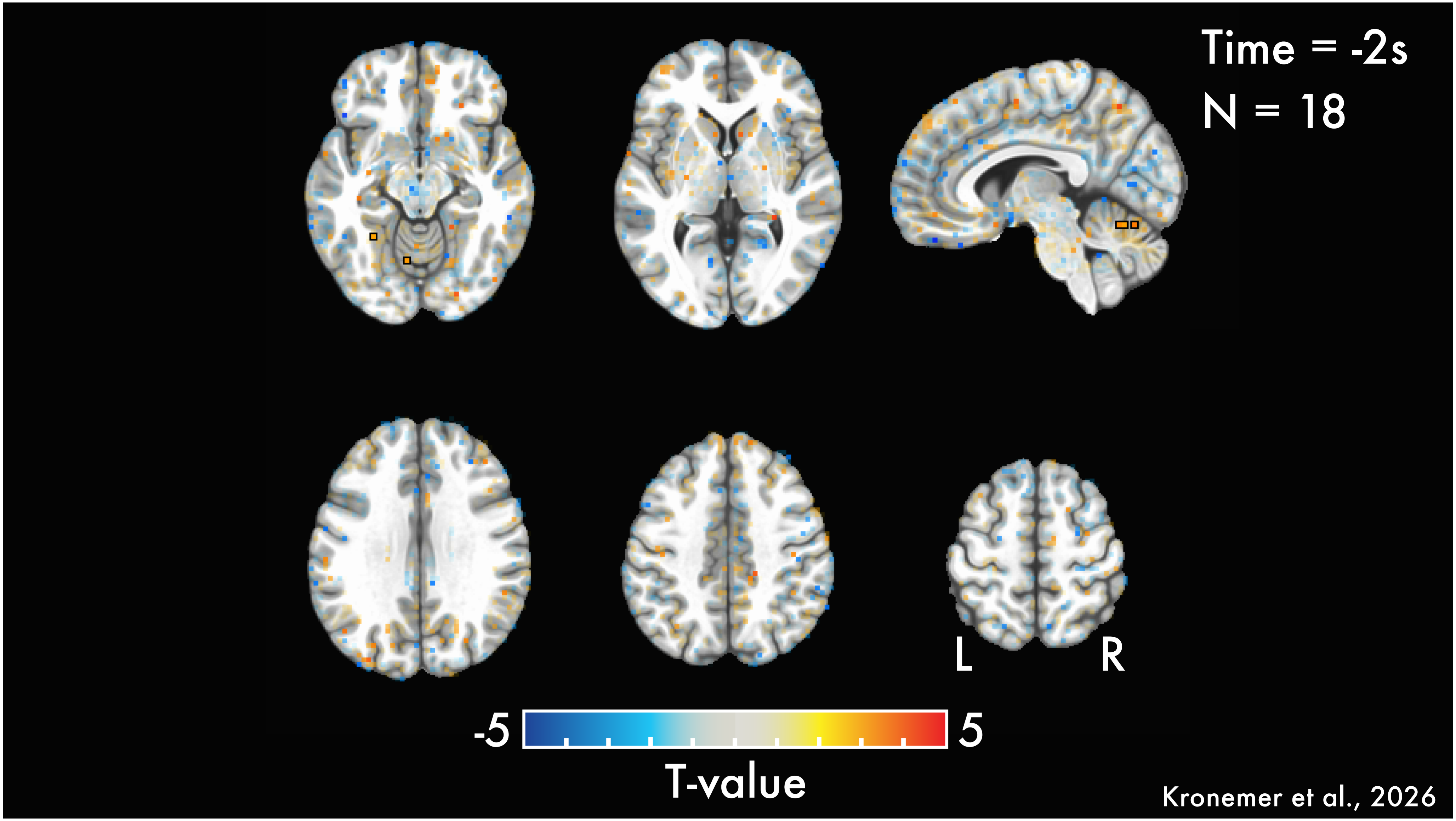

Time = -2s
N = 18
L
R
-5
5
T-value
Kronemer et al., 2026

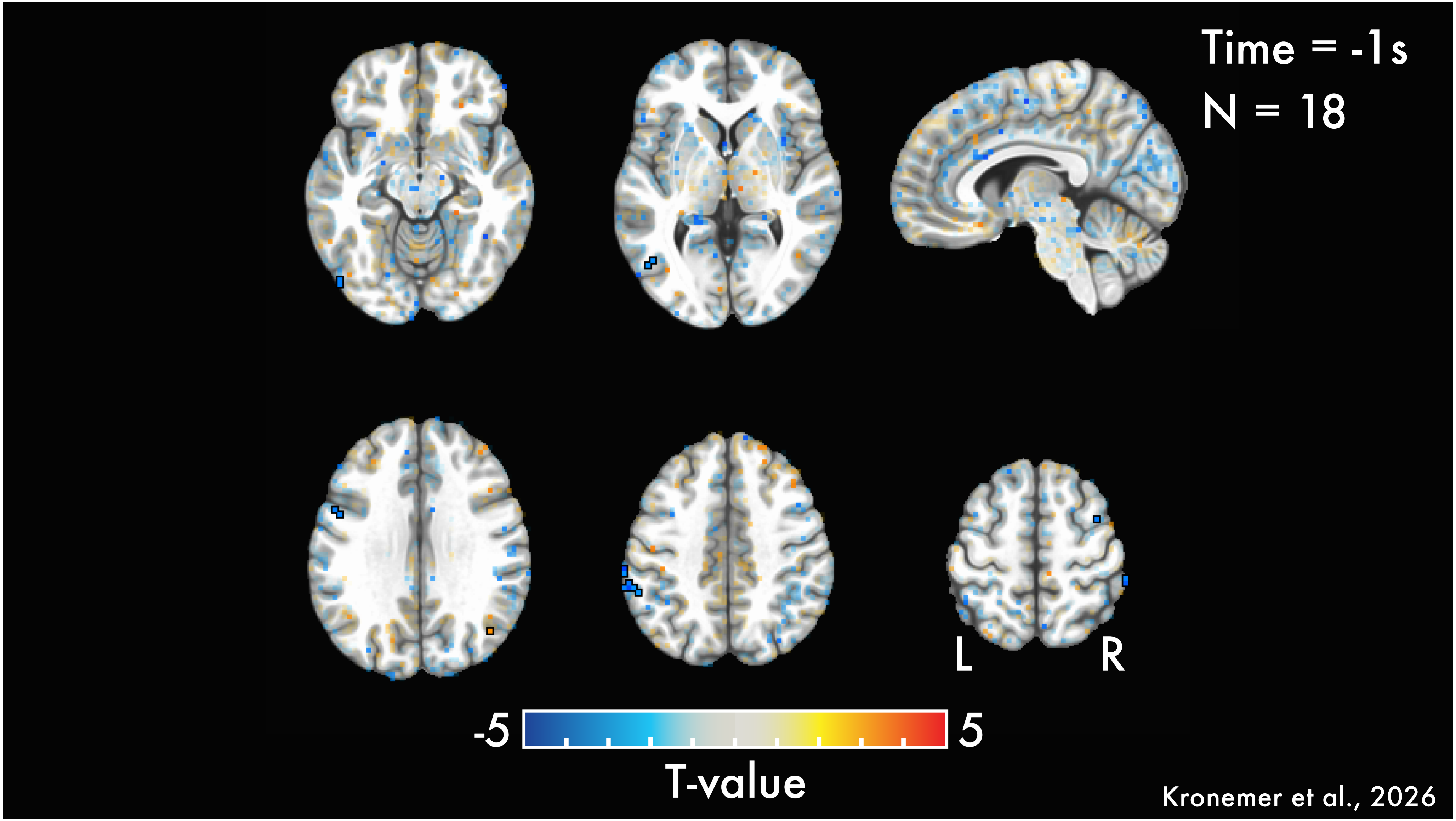
Time = -1s
N = 18
L
R
-5
5
T-value
Kronemer et al., 2026

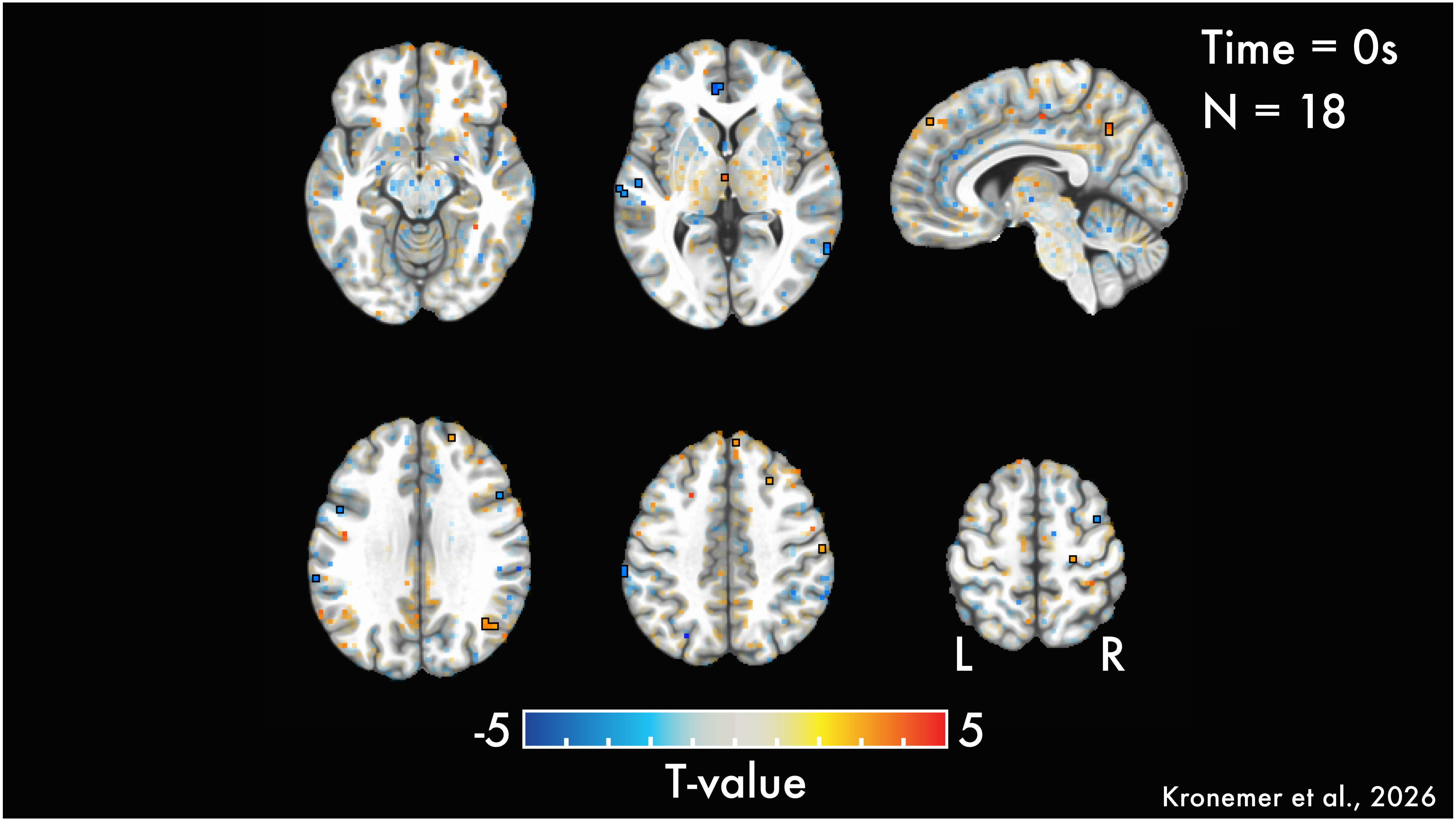
Time = 0s
N = 18
L
R
-5
5
T-value
Kronemer et al., 2026

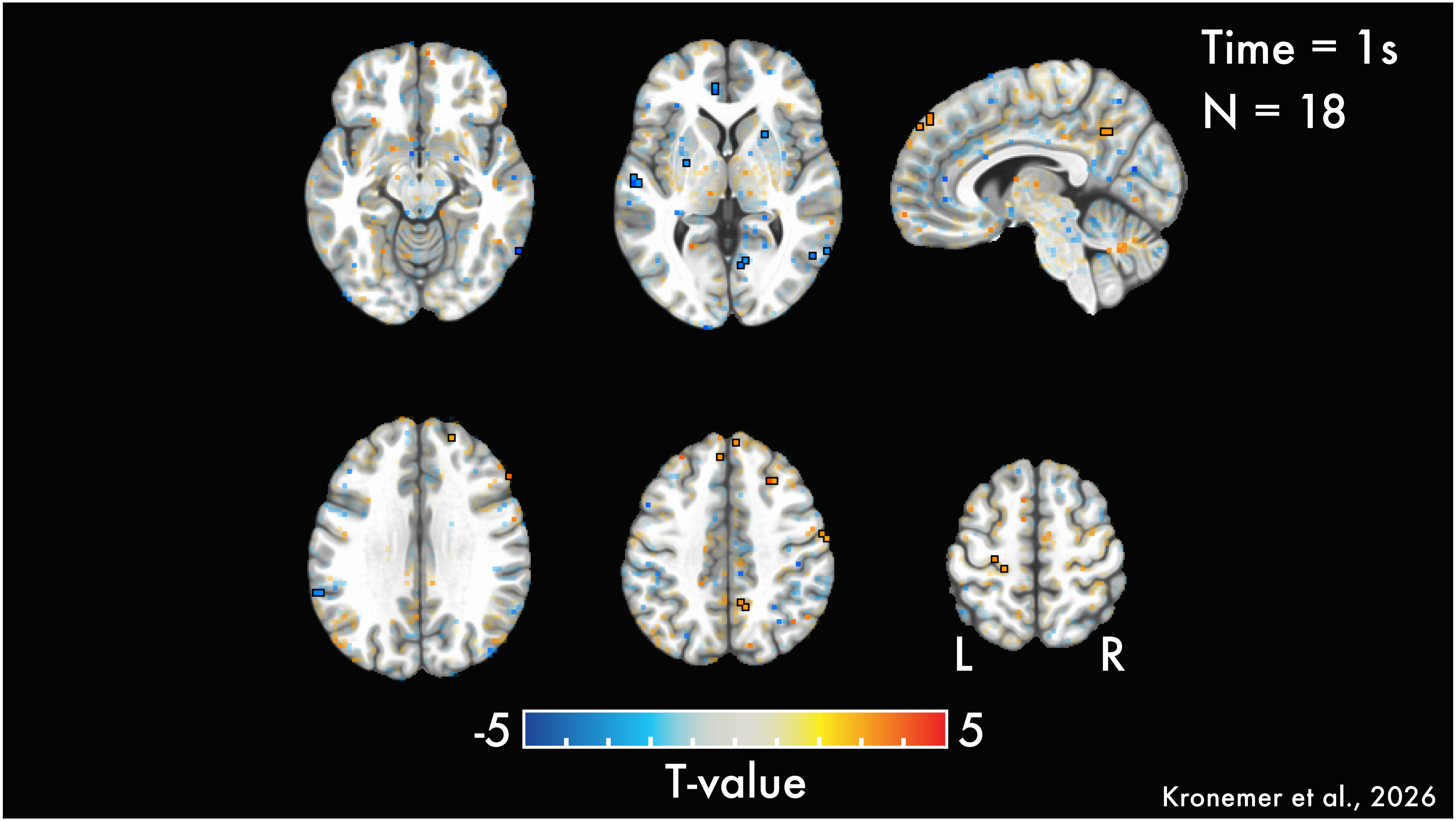

Time = 1s
N = 18
L
R
-5
5
T-value
Kronemer et al., 2026

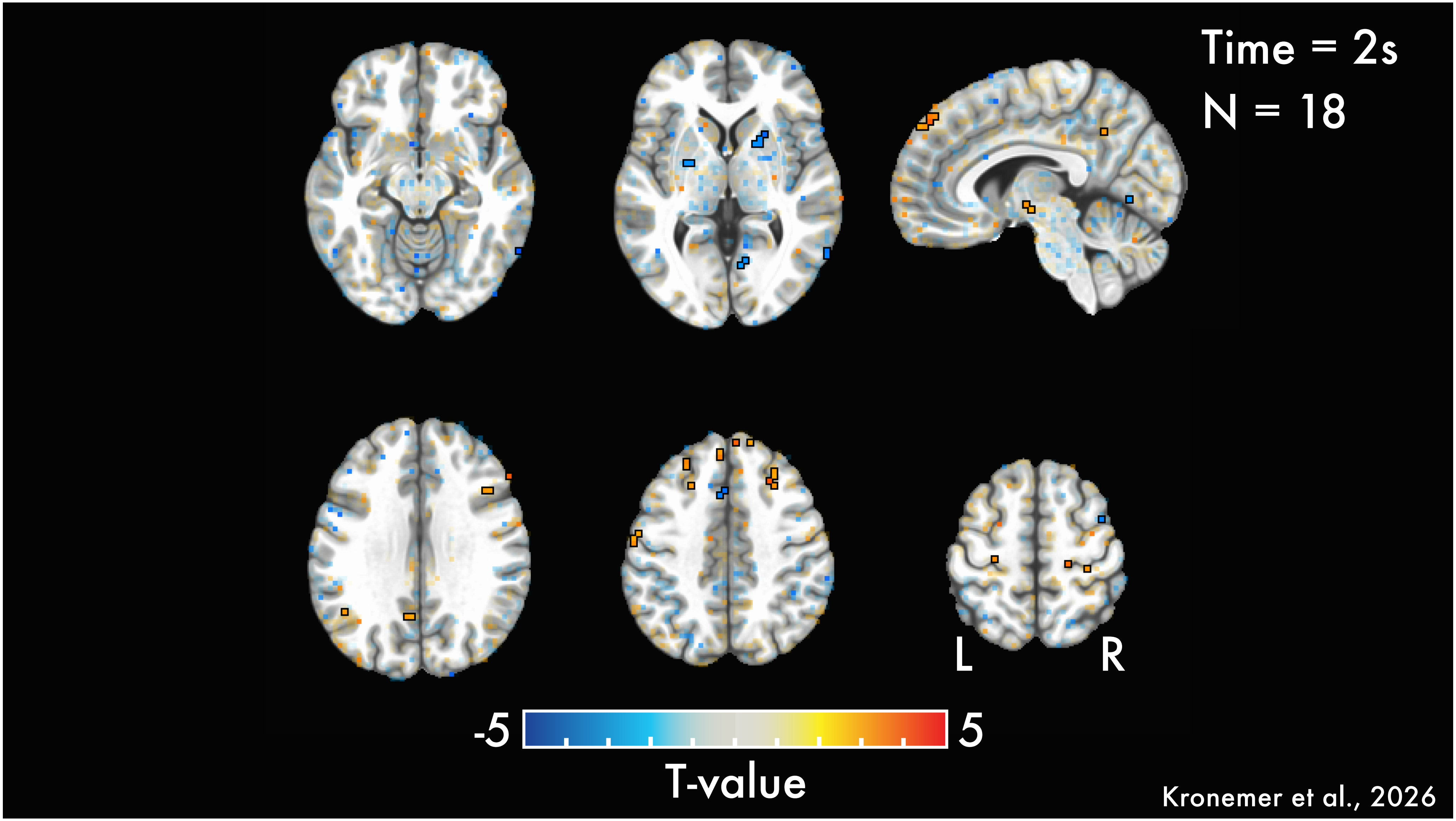

Time = 2s
N = 18
L
R
-5
5
T-value
Kronemer et al., 2026

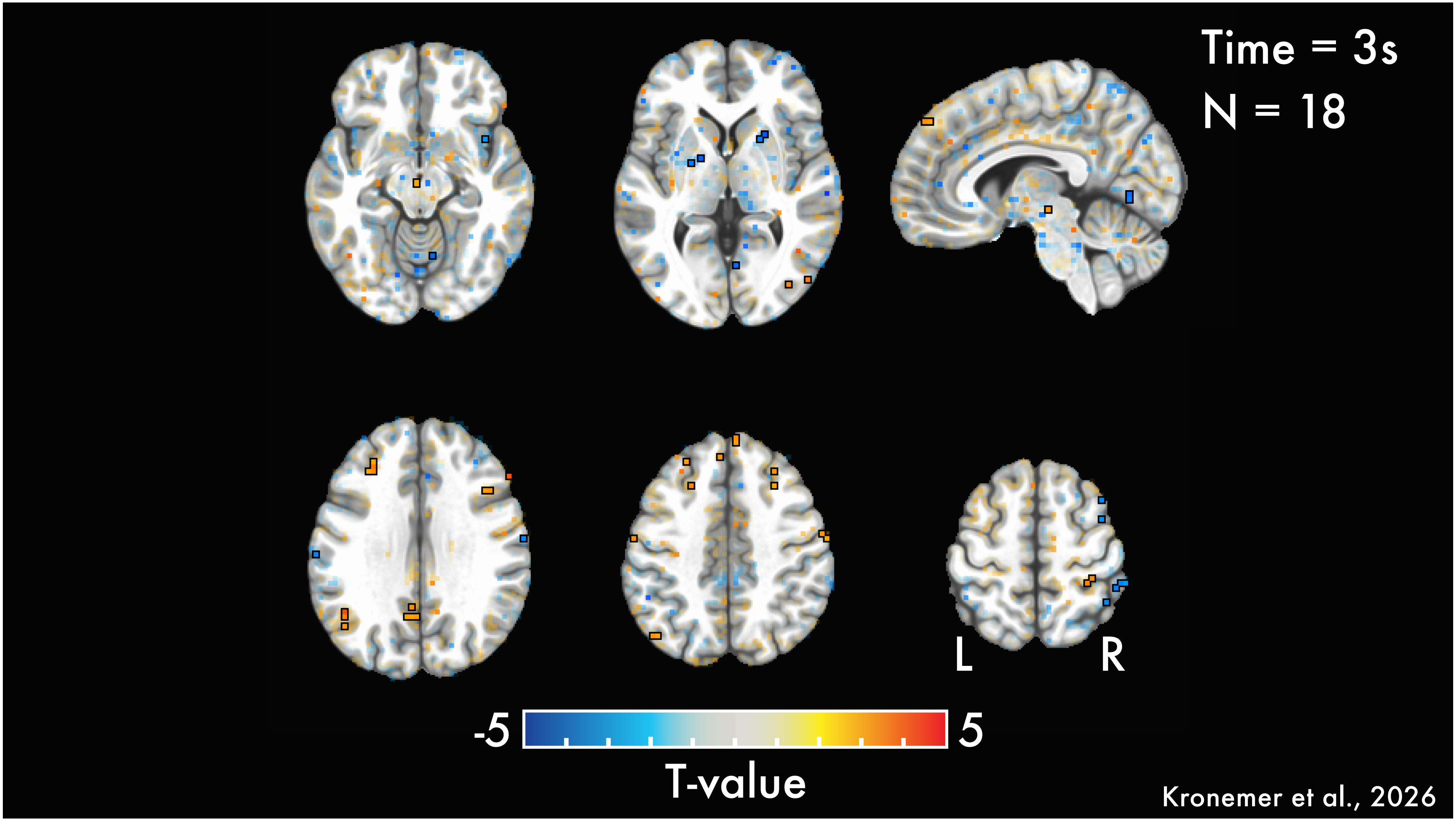
Time = 3s
N = 18
L
R
-5
5
T-value
Kronemer et al., 2026

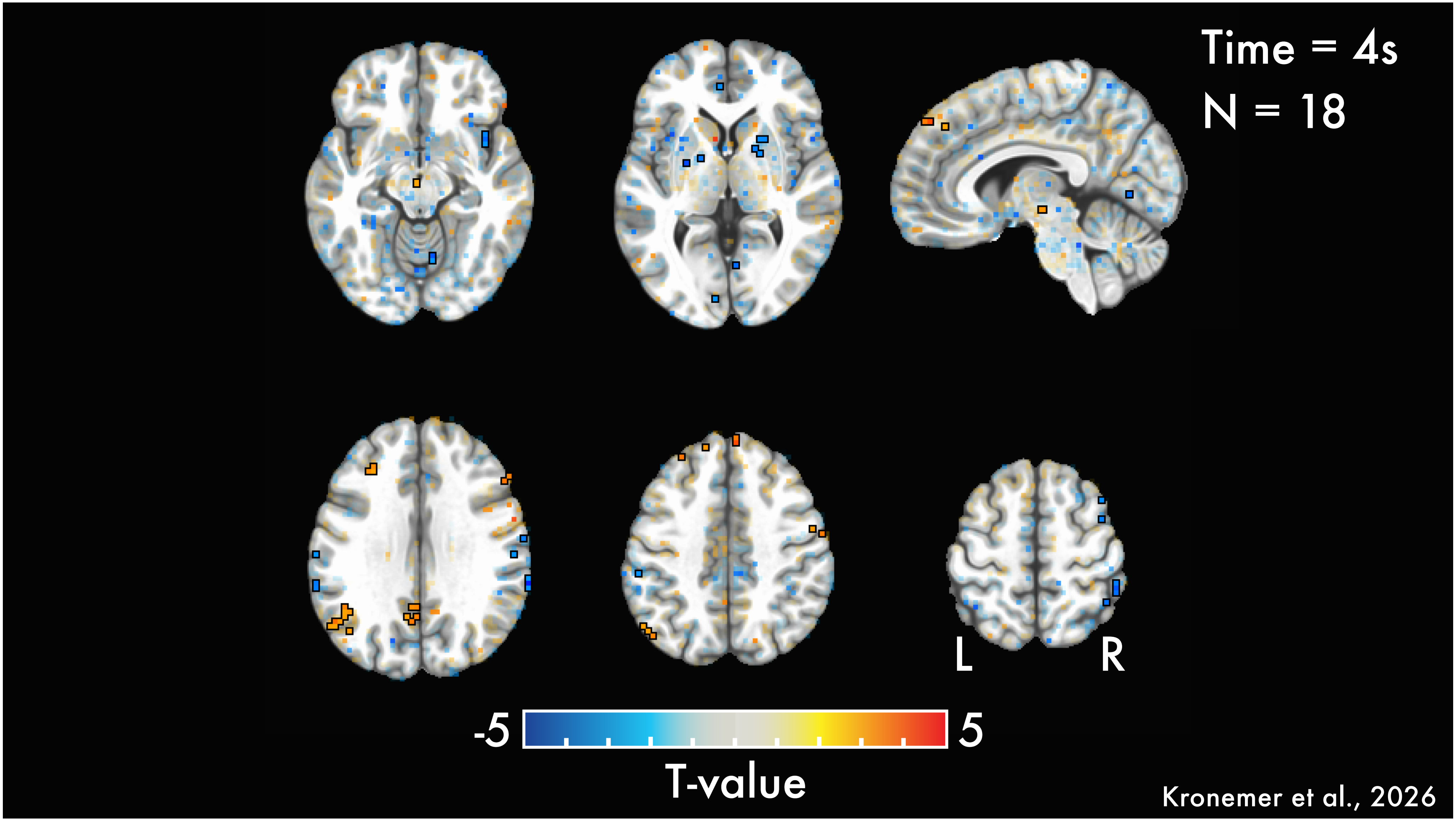

Time = 4s
N = 18
L
R
-5
5
T-value
Kronemer et al., 2026

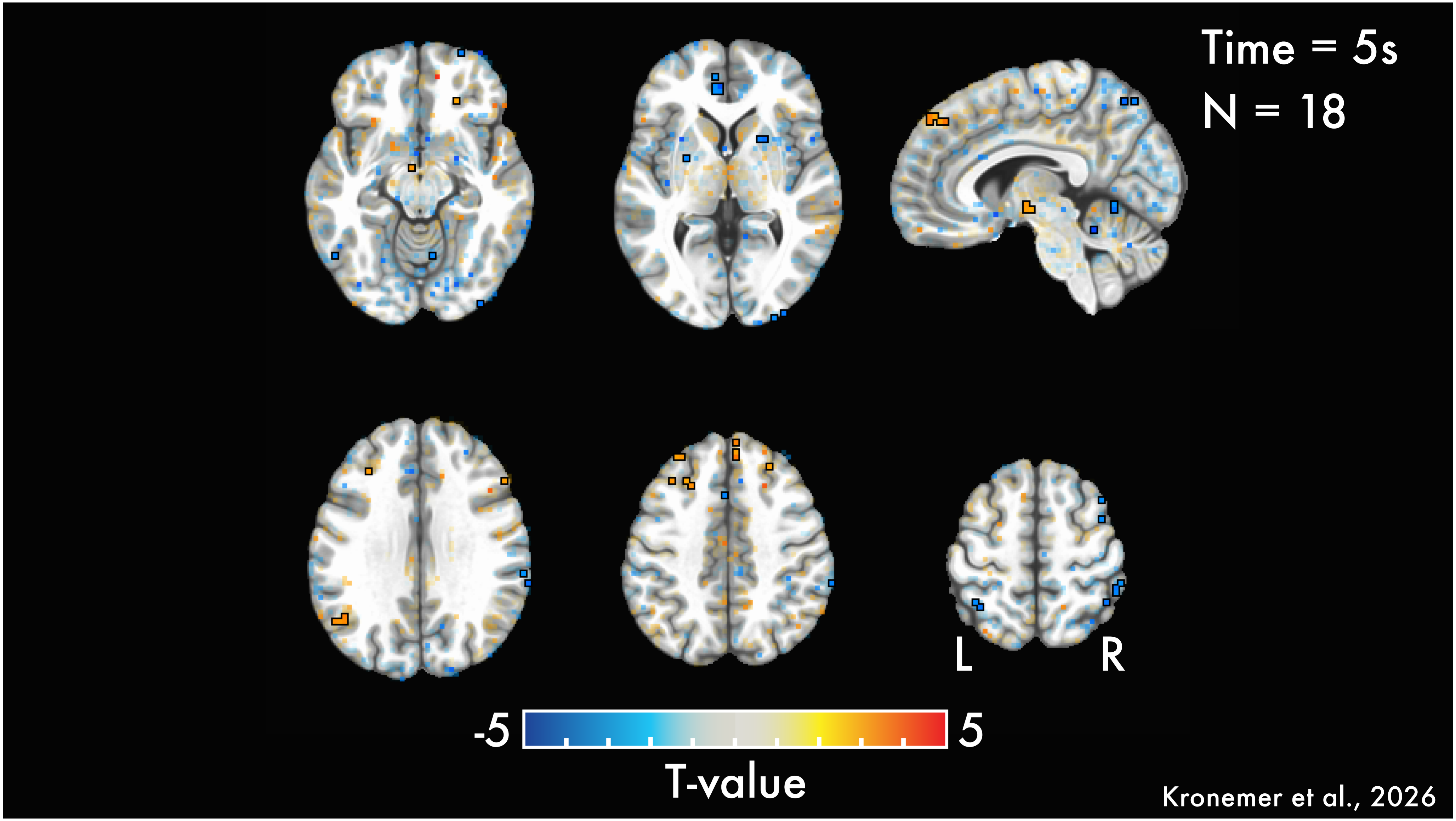
Time = 5s
N = 18
L
R
-5
5
T-value
Kronemer et al., 2026

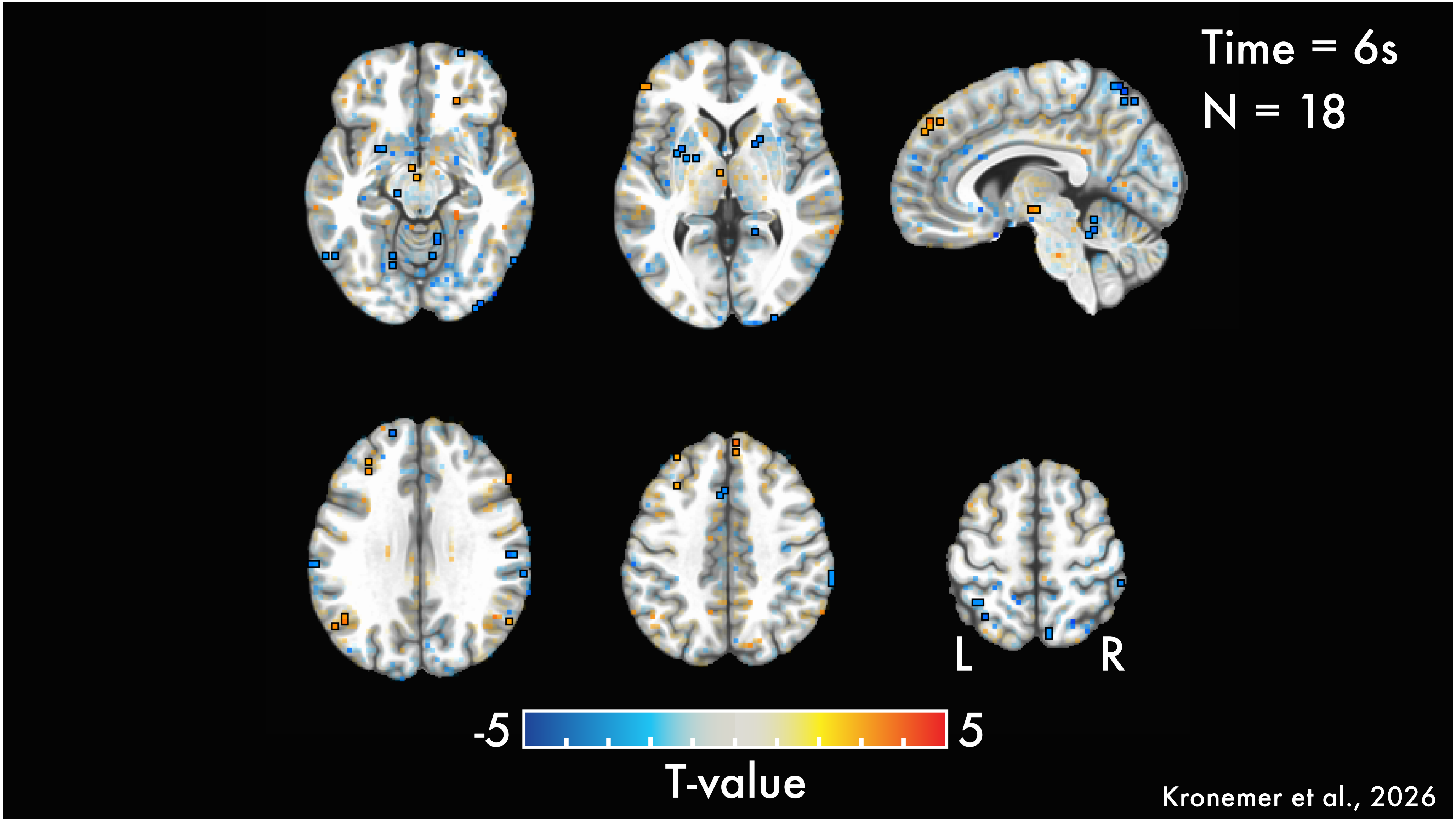

Time = 6s
N = 18
L
R
-5
5
T-value
Kronemer et al., 2026

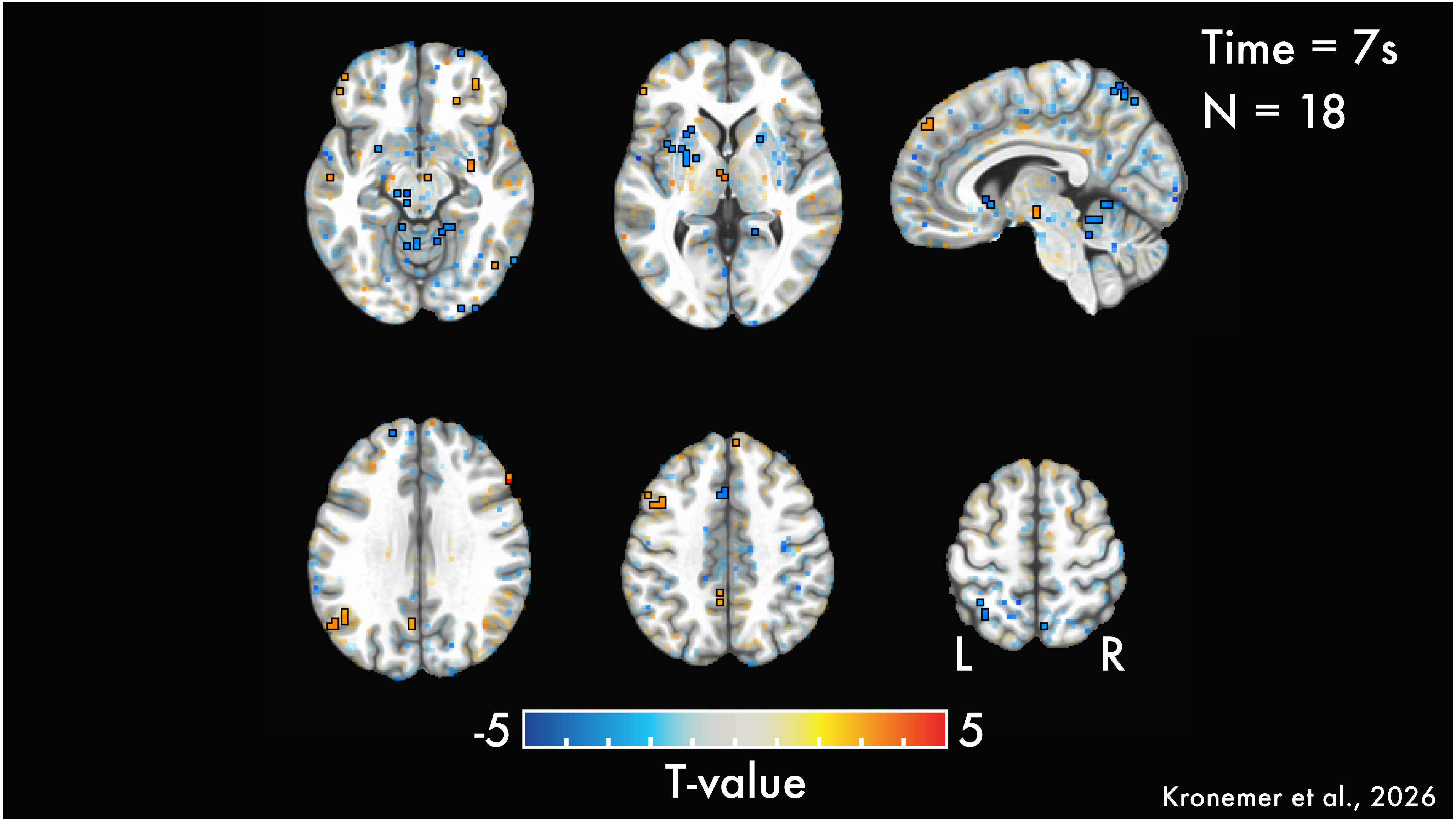
Time = 7s
N = 18
L
R
-5
5
T-value
Kronemer et al., 2026

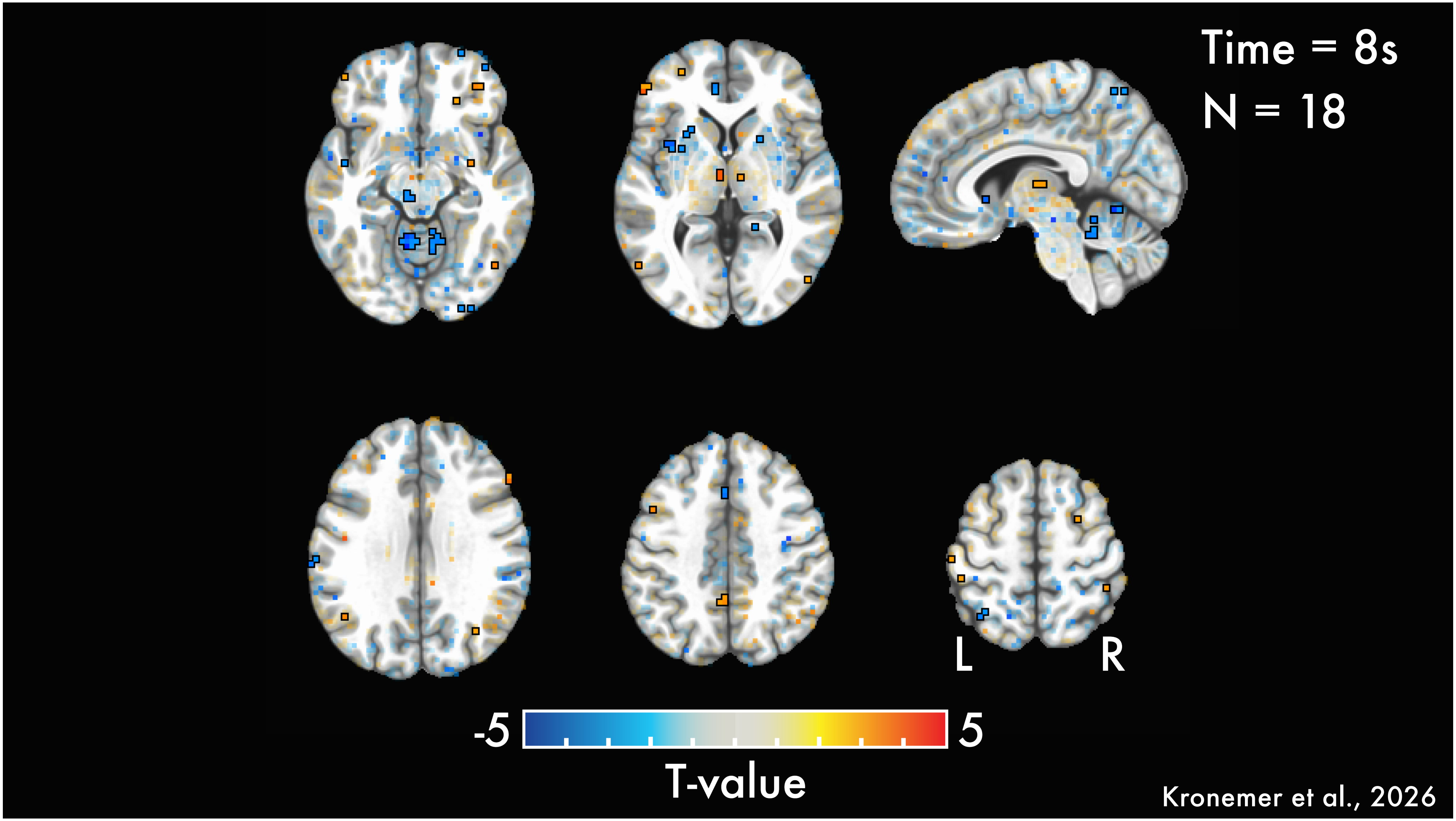

Time = 8s
N = 18
L
R
-5
5
T-value
Kronemer et al., 2026

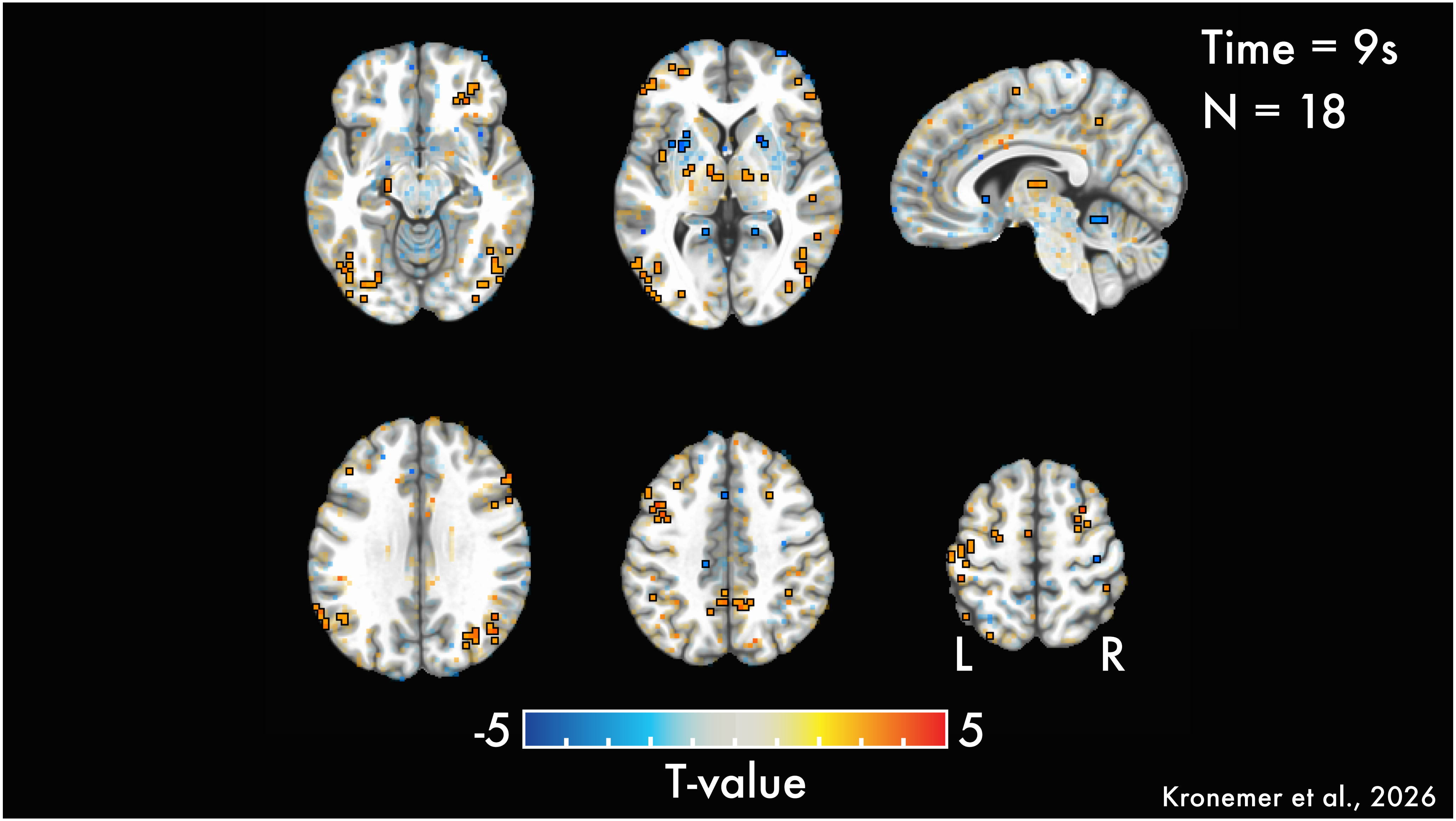
Time = 9s
N = 18
L
R
-5
5
T-value
Kronemer et al., 2026

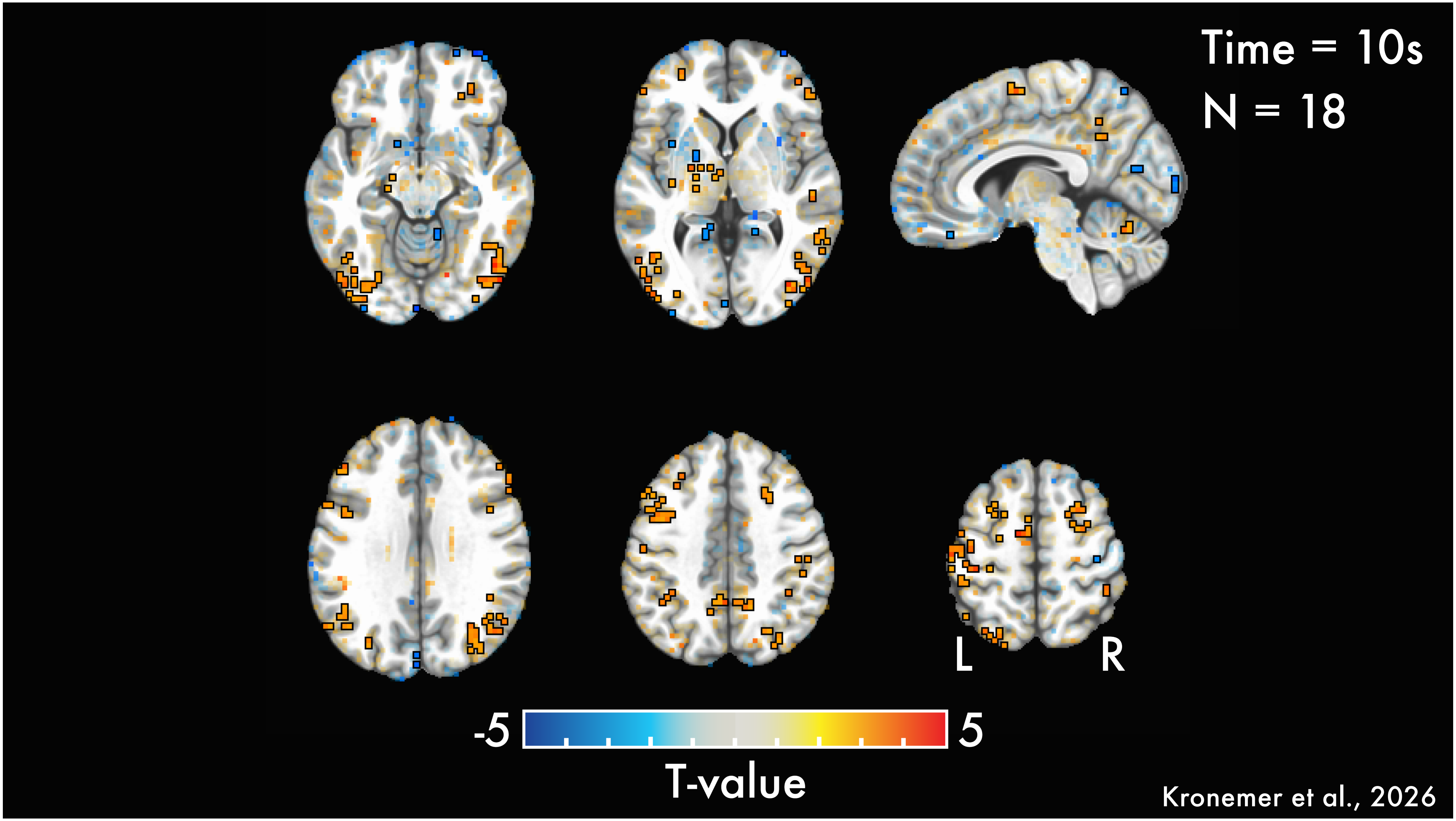
Time = 10s
N = 18
L
R
-5
5
T-value
Kronemer et al., 2026

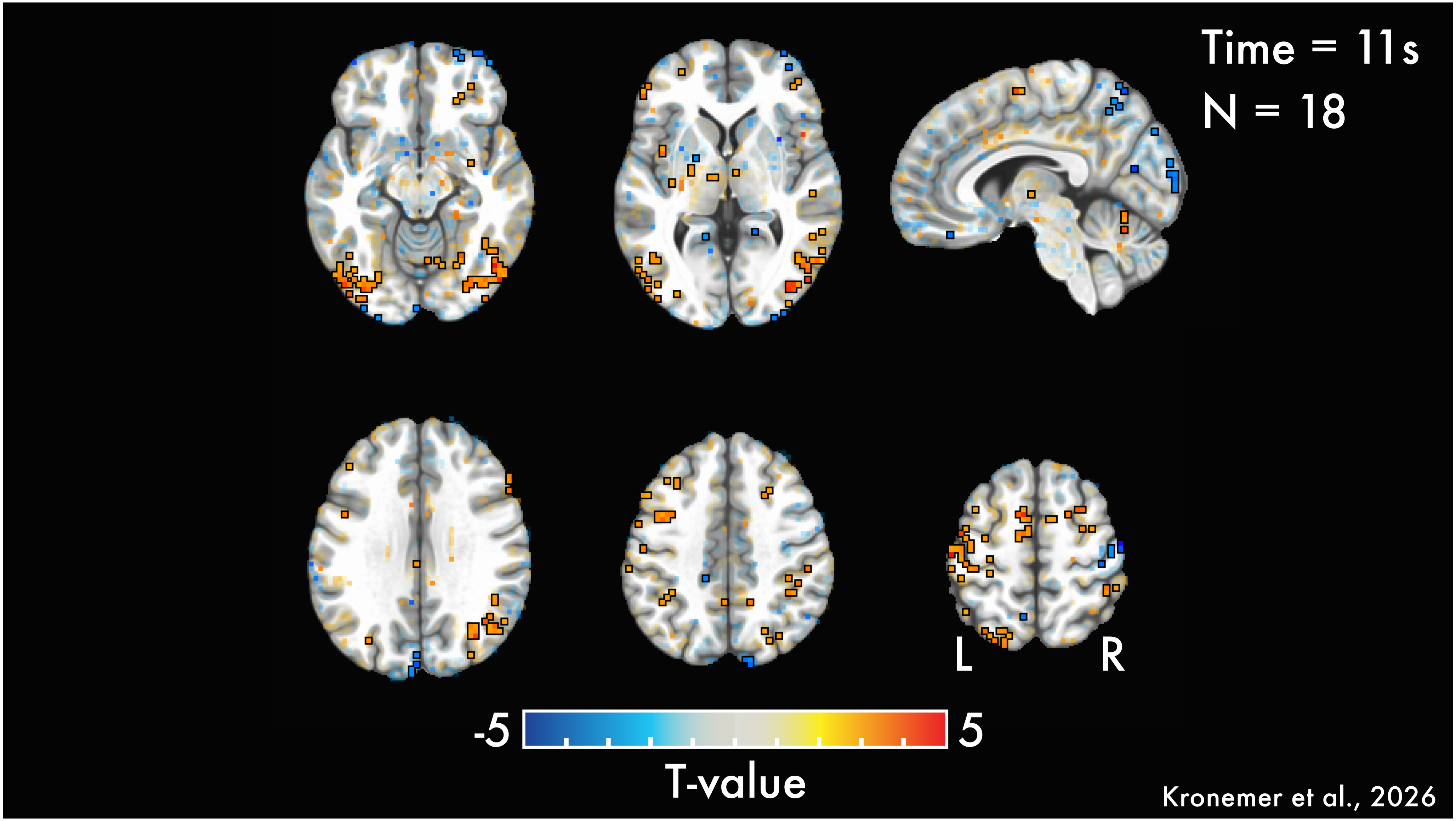

Time = 11s
N = 18
L
R
-5
5
T-value
Kronemer et al., 2026

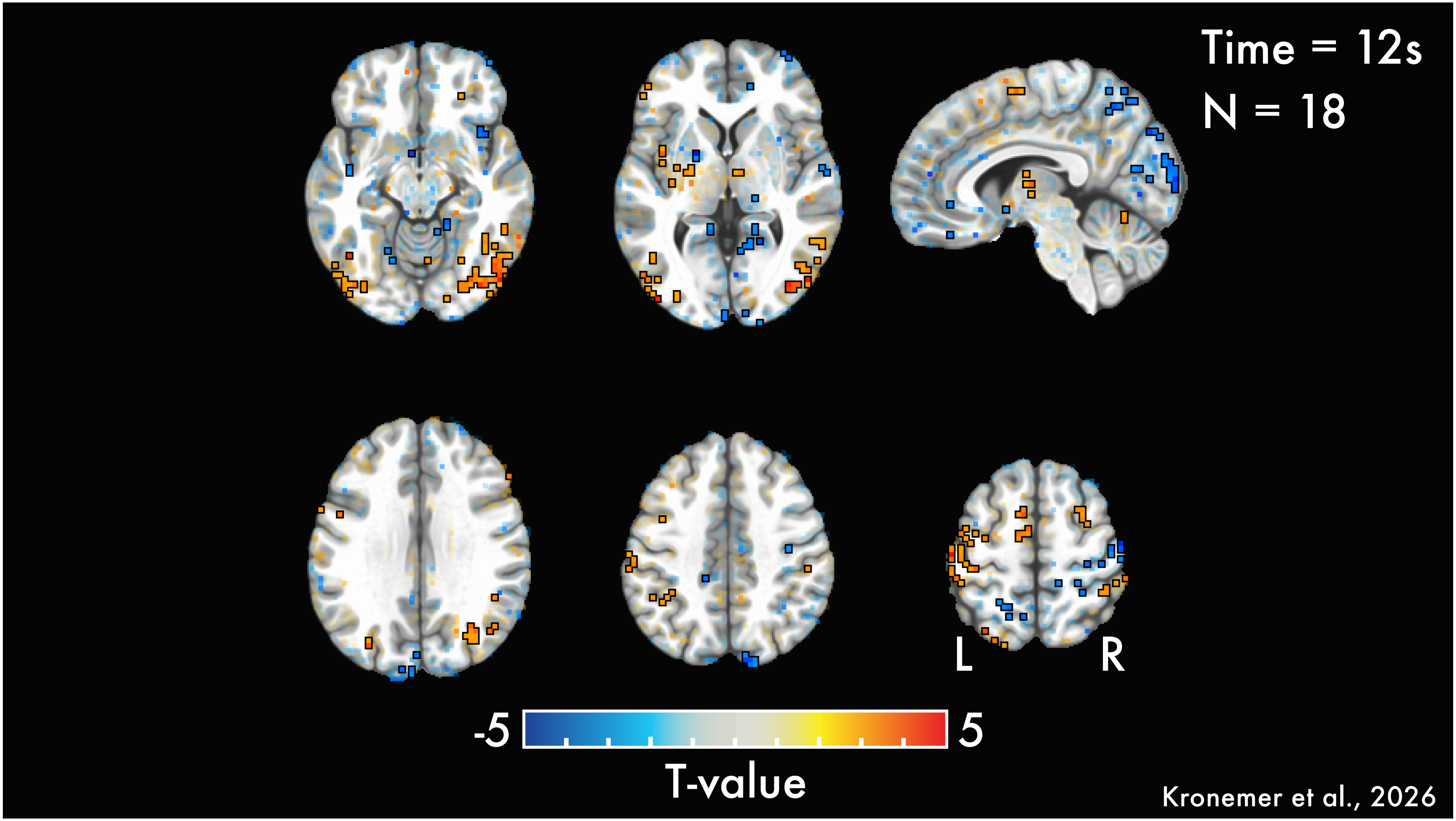
Time = 12s
N = 18
L
R
-5
5
T-value
Kronemer et al., 2026

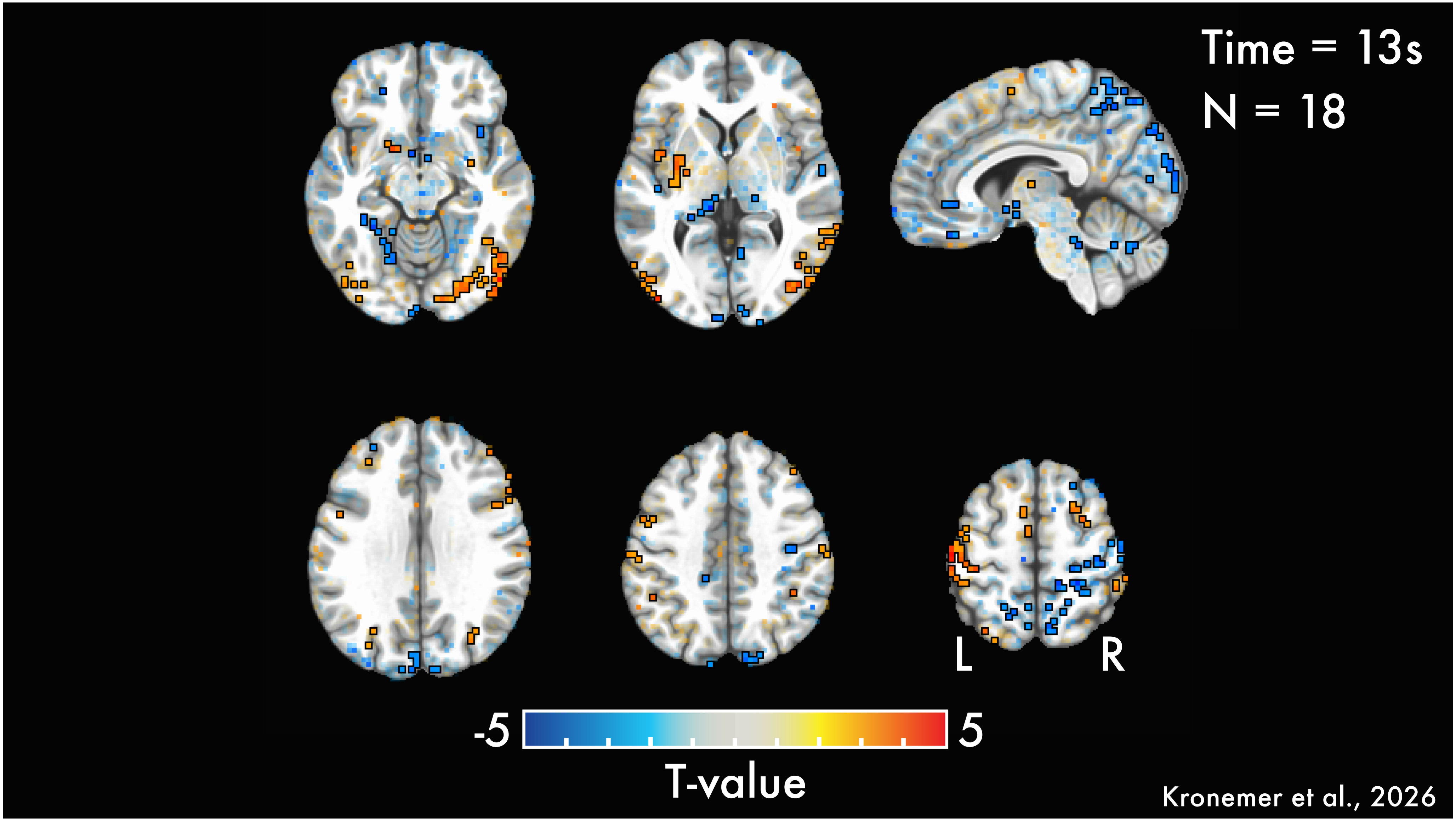
Time = 13s
N = 18
L
R
-5
5
T-value
Kronemer et al., 2026

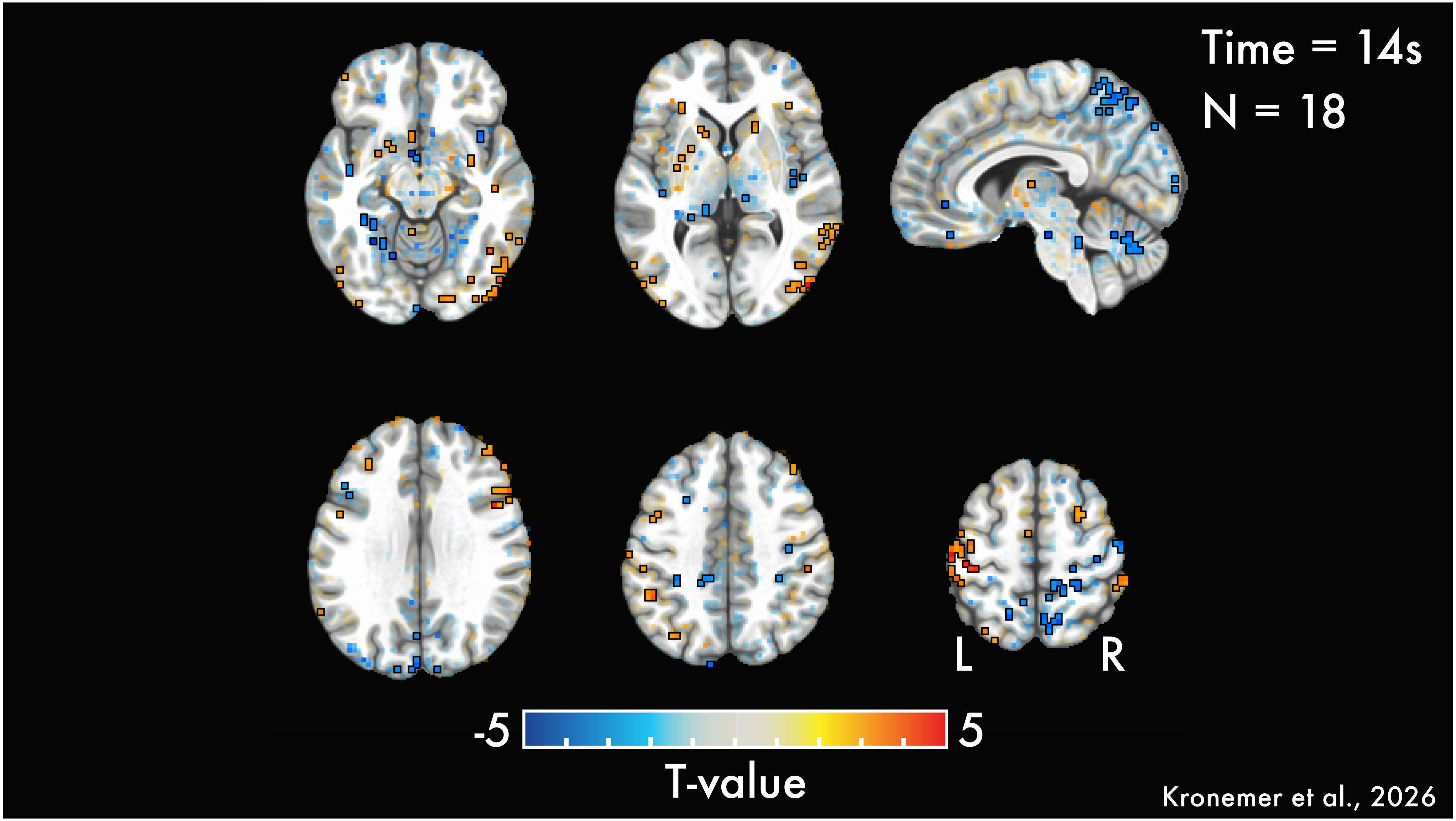
Time = 14s
N = 18
L
R
-5
5
T-value
Kronemer et al., 2026

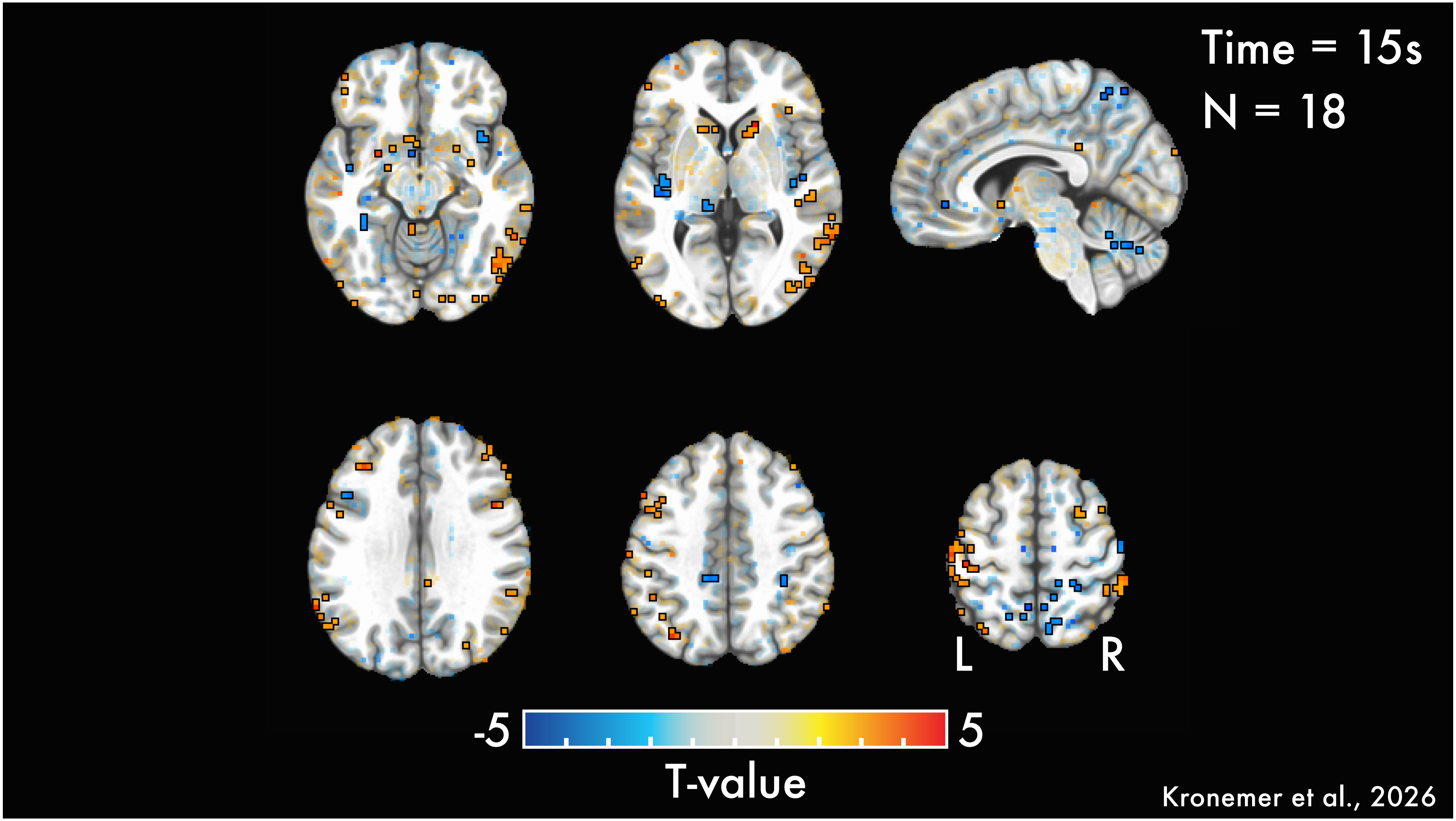
Time = 15s
N = 18
L
R
-5
5
T-value
Kronemer et al., 2026

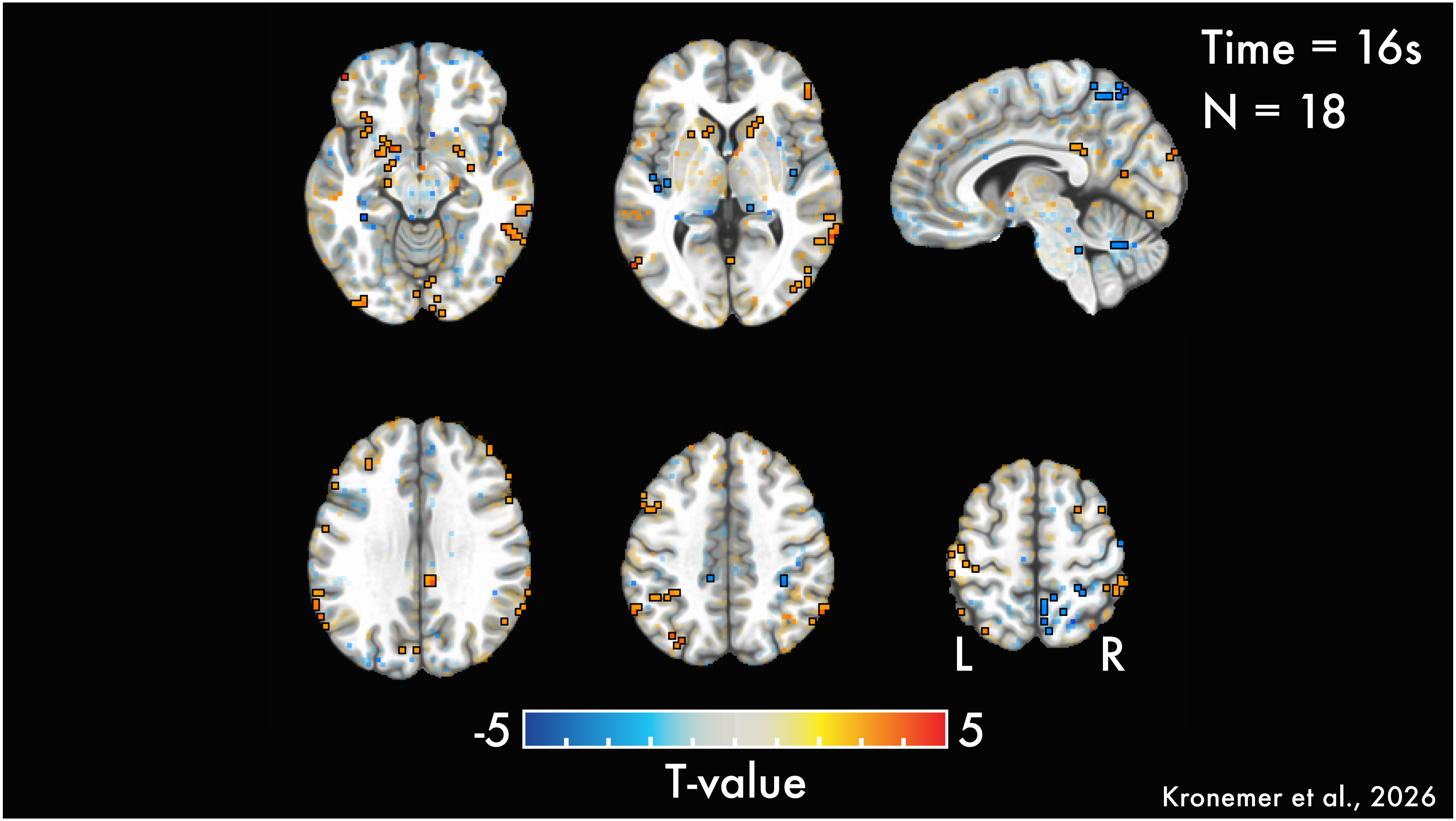
Time = 16s
N = 18
L
R
-5
5
T-value
Kronemer et al., 2026

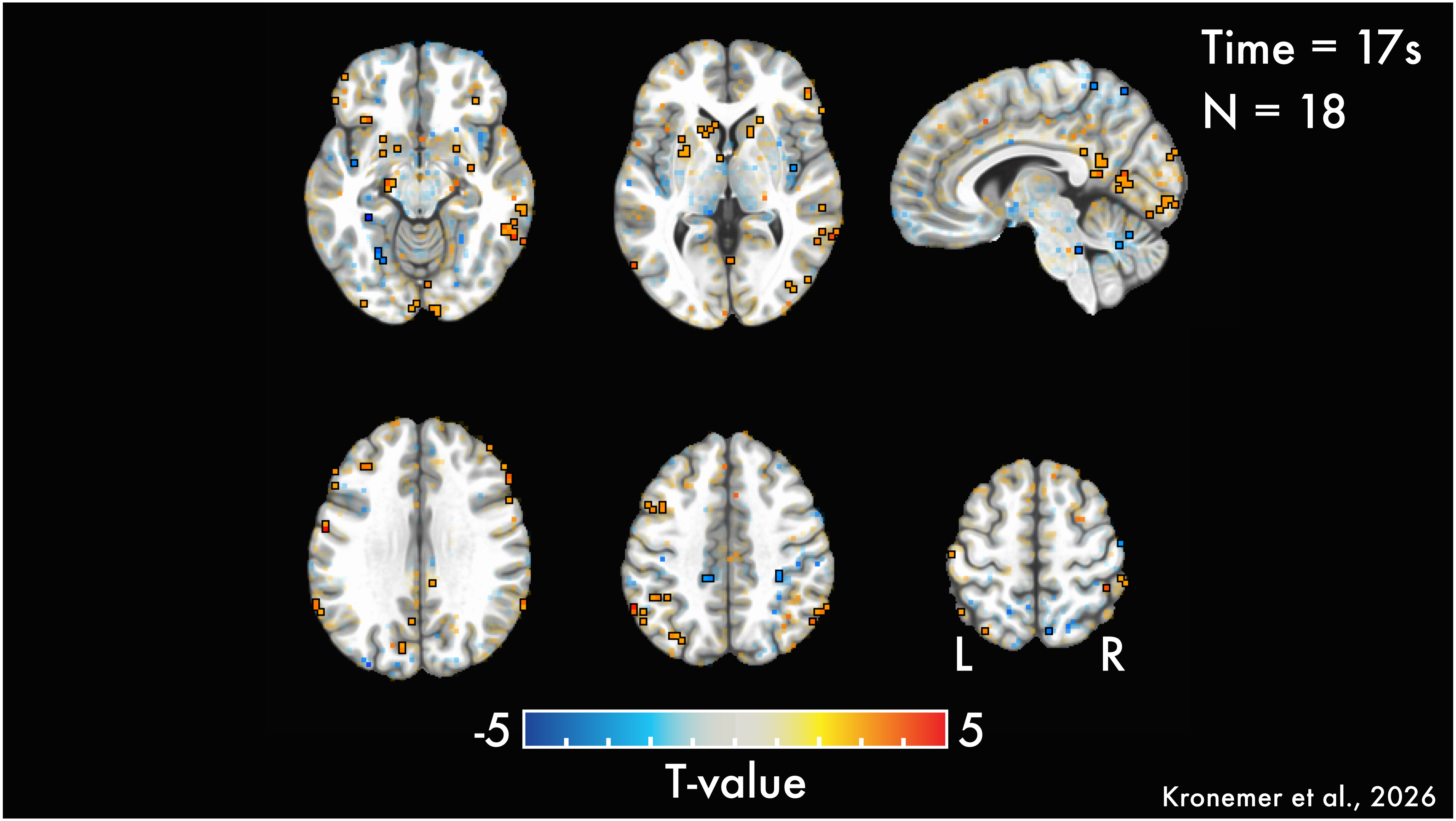
Time = 17s
N = 18
L
R
-5
5
T-value
Kronemer et al., 2026

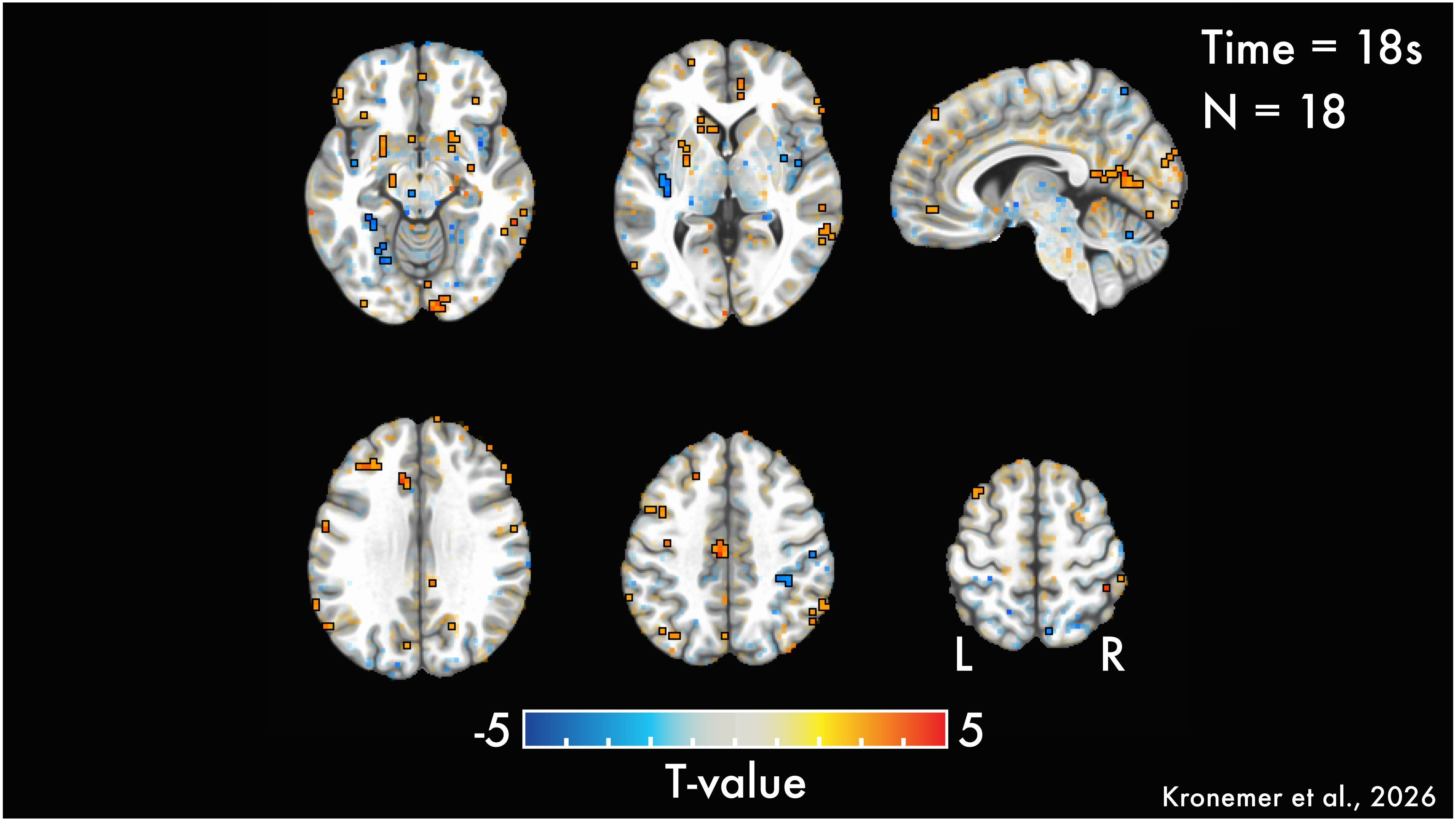

Time = 18s
N = 18
L
R
-5
5
T-value
Kronemer et al., 2026

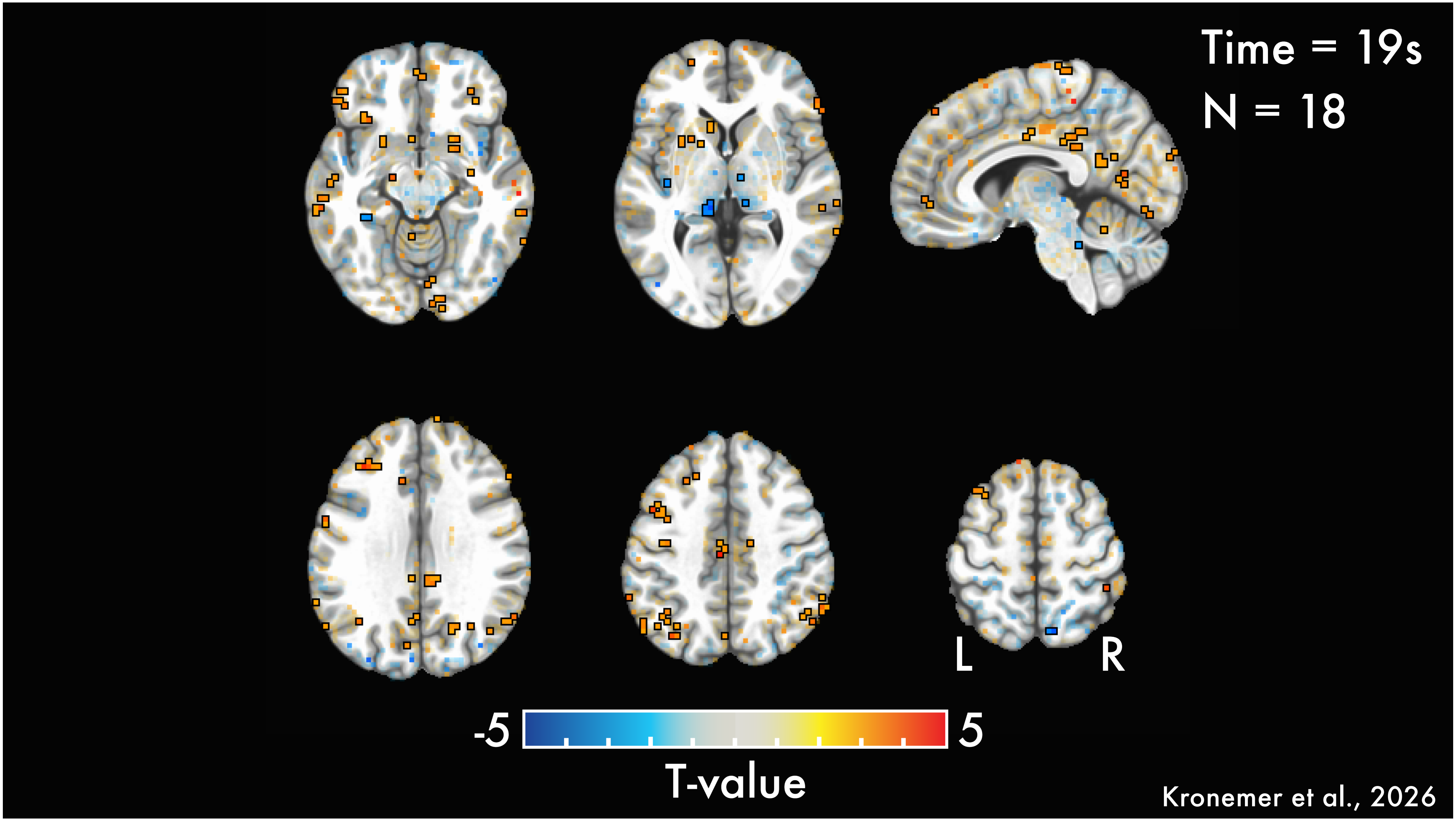

Time = 19s
N = 18
L
R
-5
5
T-value
Kronemer et al., 2026

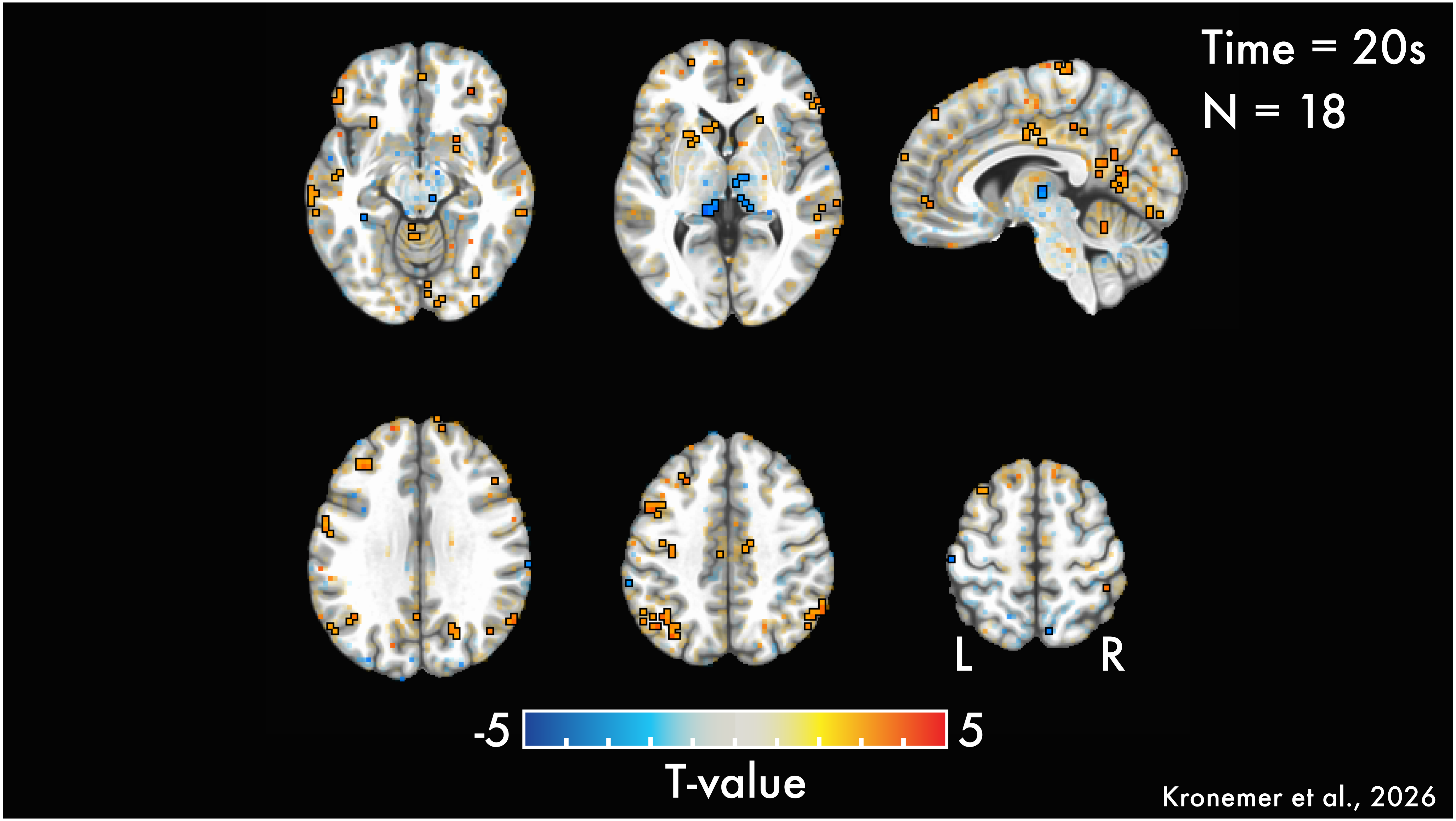

Time = 20s
N = 18
L
R
-5
5
T-value
Kronemer et al., 2026

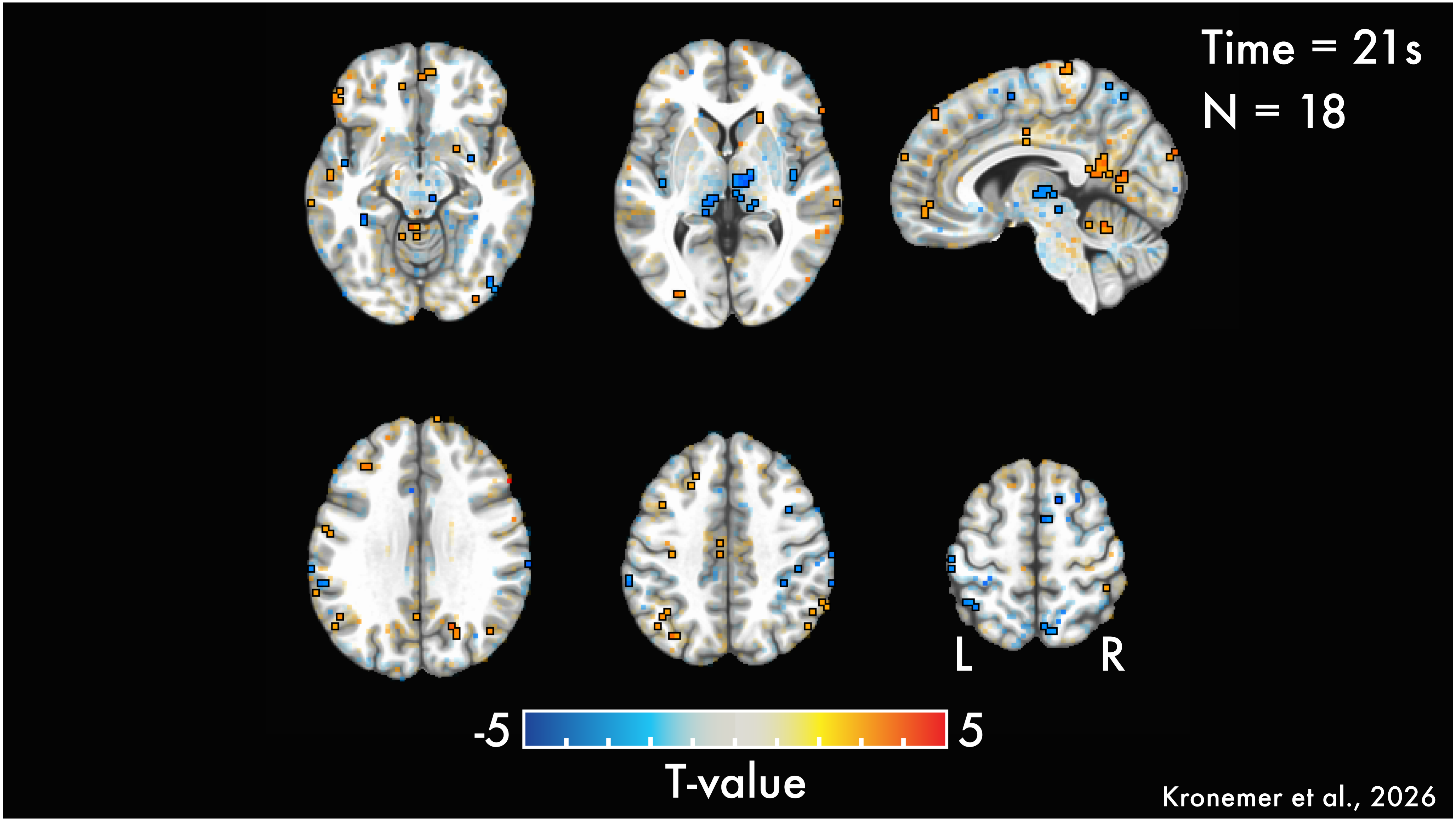
Time = 21s
N = 18
L
R
-5
5
T-value
Kronemer et al., 2026

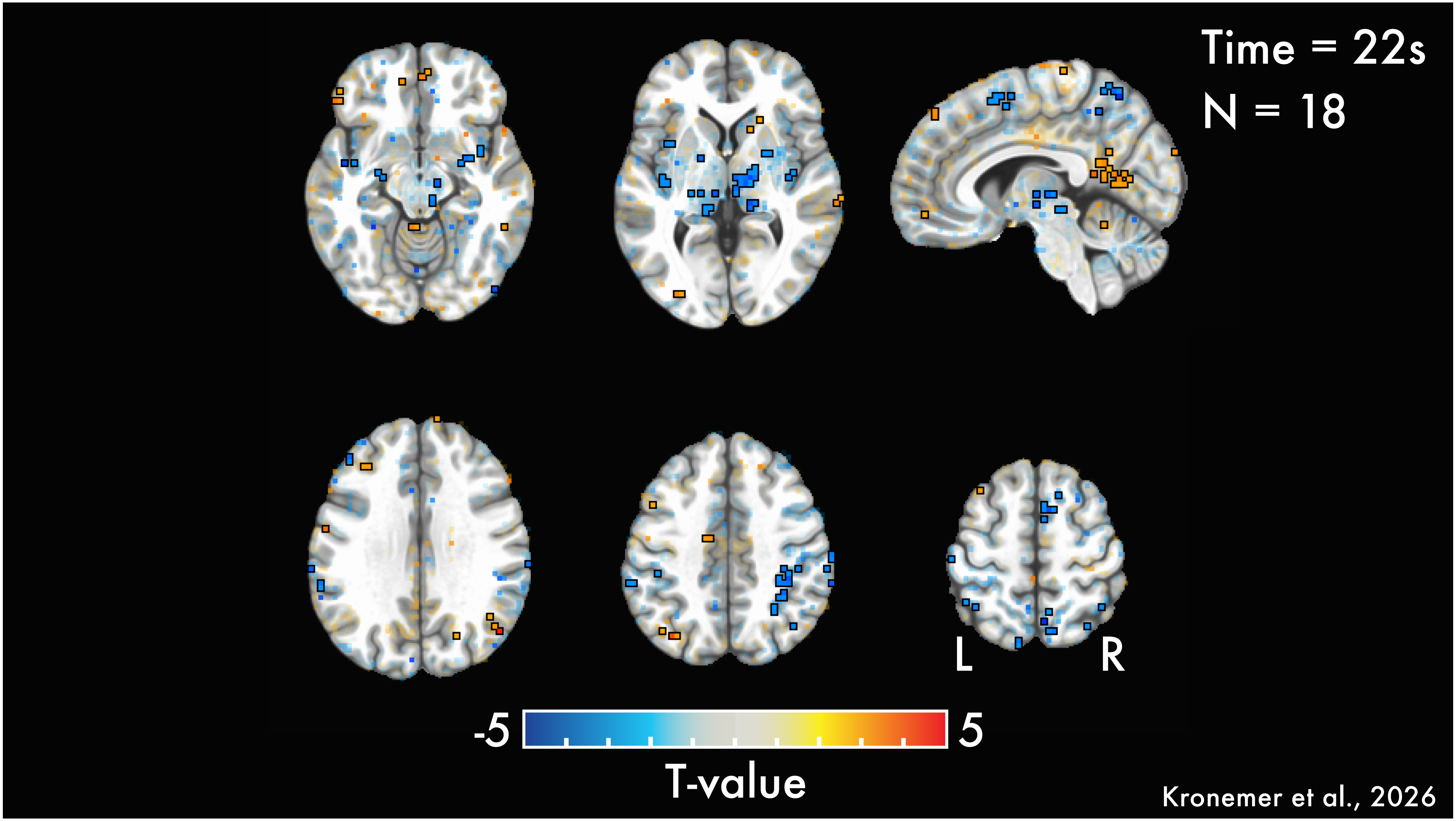
Time = 22s
N = 18
L
R
-5
5
T-value
Kronemer et al., 2026

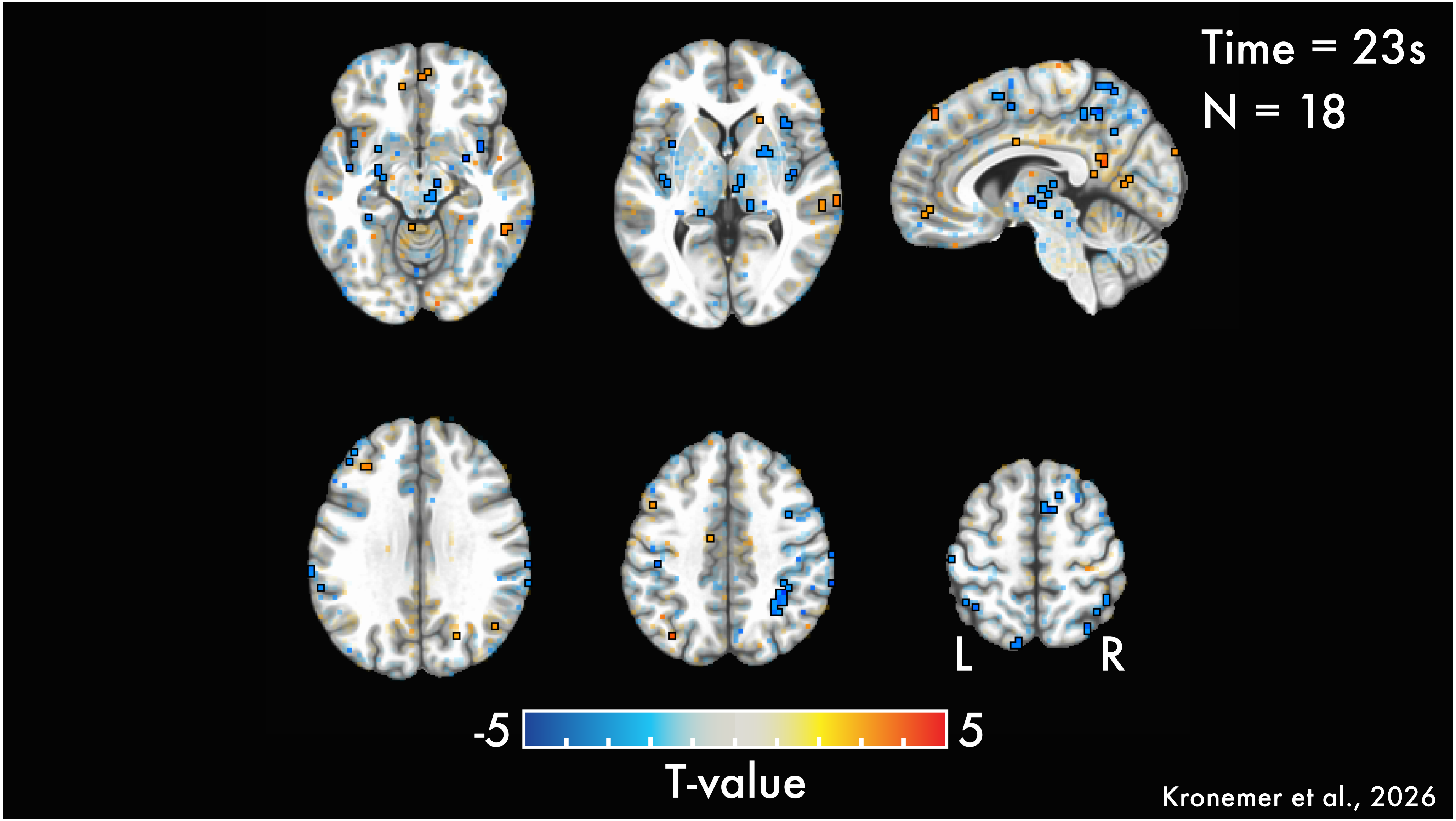

Time = 23s
N = 18
L
R
-5
5
T-value
Kronemer et al., 2026

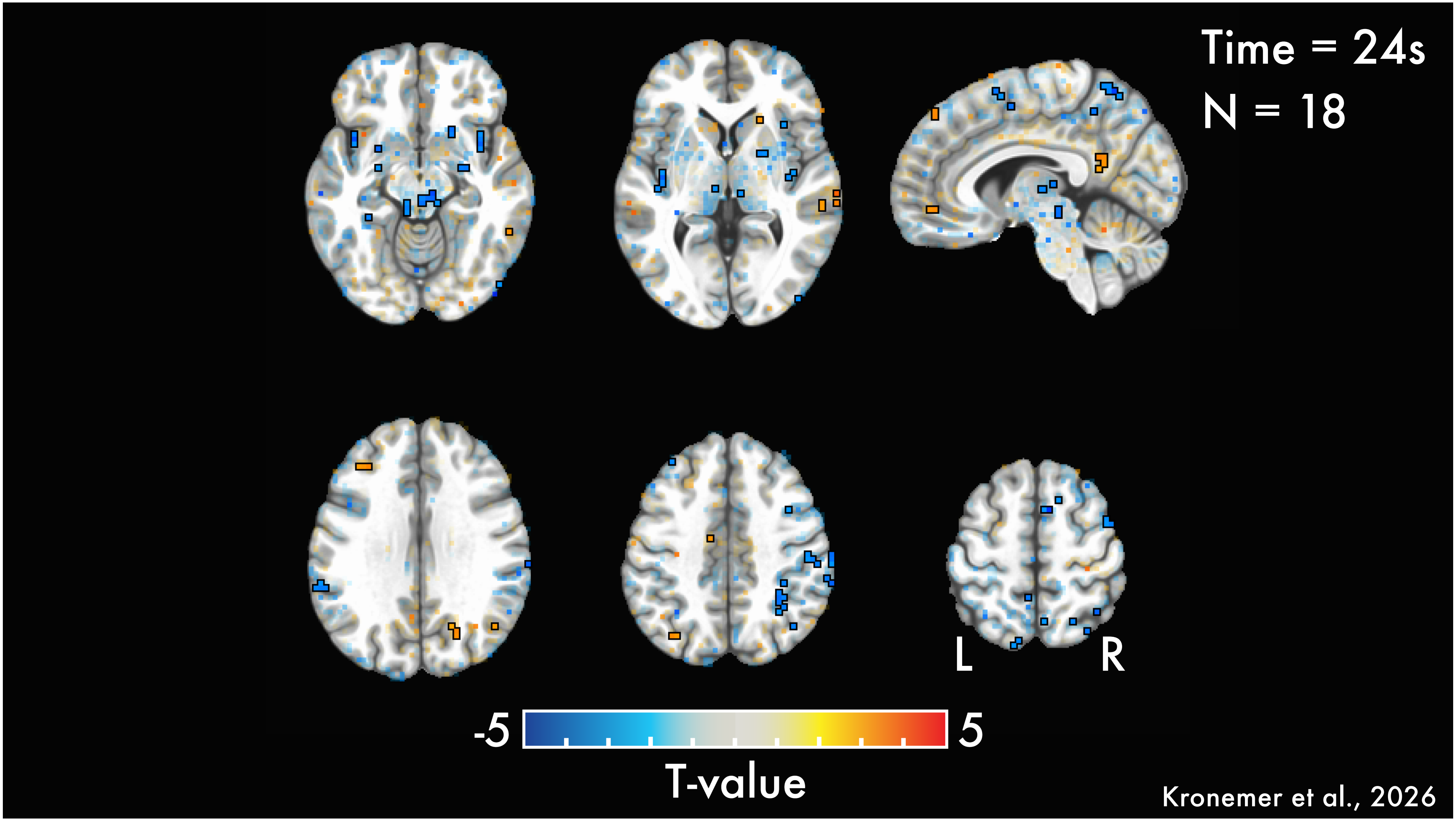

Time = 24s
N = 18
L
R
-5
5
T-value
Kronemer et al., 2026

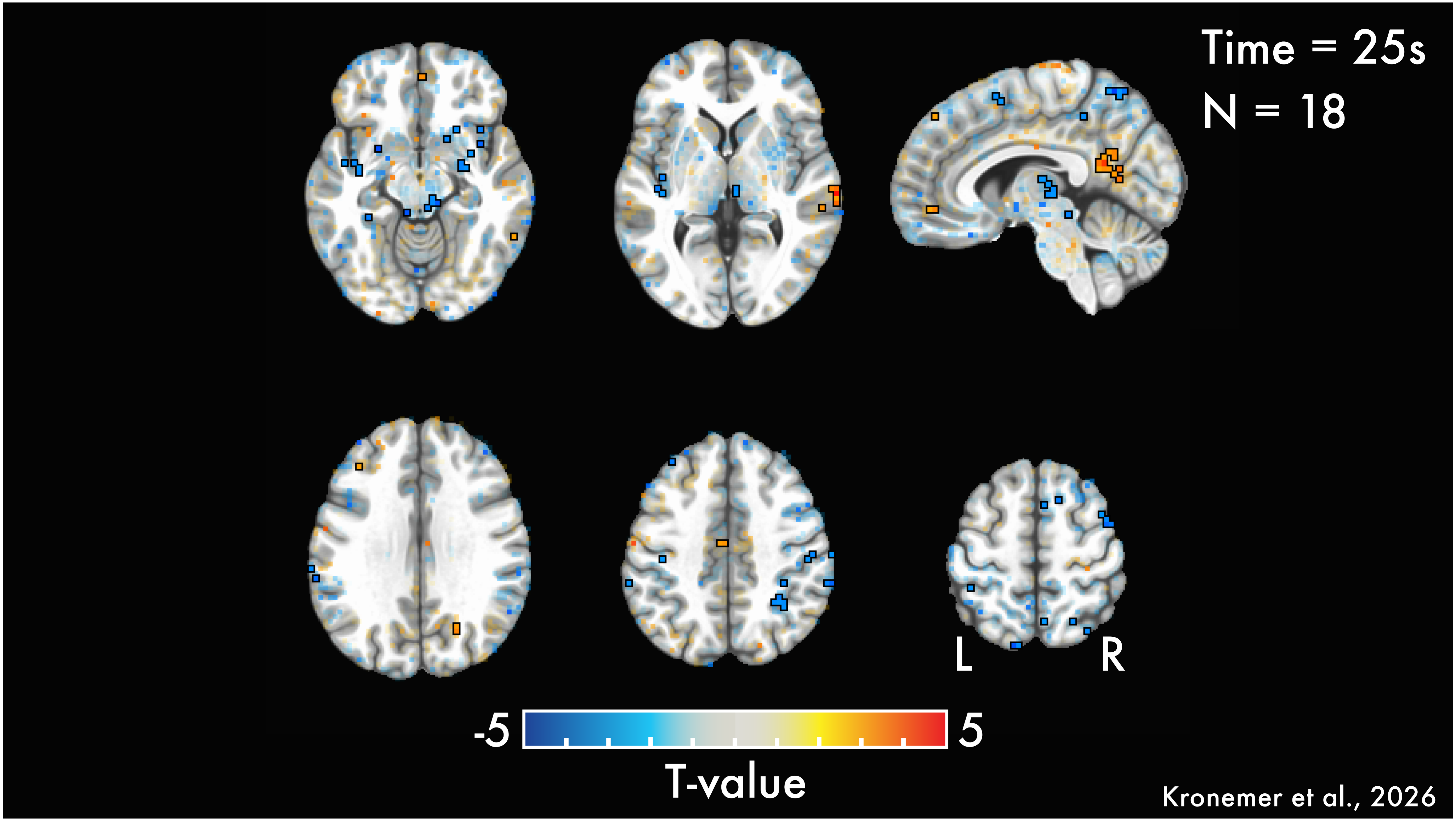
Time = 25s
N = 18
L
R
-5
5
T-value
Kronemer et al., 2026
